# Identification of the active mechanism of aminoglycoside entry in *V. cholerae* through characterization of sRNA *ctrR,* regulating carbohydrate utilization and transport

**DOI:** 10.1101/2023.07.19.549712

**Authors:** Sebastian A. Pierlé, Manon Lang, Rocío López-Igual, Evelyne Krin, Dominique Fourmy, Sean P. Kennedy, Marie-Eve Val, Zeynep Baharoglu, Didier Mazel

## Abstract

The possible active entry of aminoglycosides in bacterial cells has been debated since the development of this antibiotic family. Here we report the identification of their active transport mechanism in *Vibrio* species. We combined genome-wide transcriptional analysis and fitness screens to identify alterations driven by treatment of *V. cholerae* with sub-minimum inhibitory concentrations (sub-MIC) of the aminoglycoside tobramycin. RNA-seq data showed downregulation of the small non-coding RNA *ncRNA586* during such treatment, while Tn-seq revealed that inactivation of this sRNA was associated with improved fitness in the presence of tobramycin. This sRNA is located near sugar transport genes and previous work on a homologous region in *Vibrio tasmaniensis* suggested that this sRNA stabilizes gene transcripts for carbohydrate transport and utilization, as well as phage receptors. The role for *ncRNA586*, hereafter named *ctrR*, in the transport of both carbohydrates and aminoglycosides, was further investigated. Flow cytometry on cells treated with a fluorescent aminoglycoside confirmed the role of *ctrR* and of carbohydrate transporters in differential aminoglycoside entry. Despite sequence diversity, *ctrR* showed functional conservation across the Vibrionales. This system in directly modulated by carbon sources, suggesting regulation by carbon catabolite repression, a widely conserved mechanism in Gram-negative bacteria, priming future research on aminoglycoside uptake by sugar transporters in other bacterial species.

## Introduction

Antibiotics revolutionized medicine, extending life expectancies, and reducing child mortality. The first limit to their efficacy is the entry inside the bacterial cell, and several mechanisms have been involved as first lines of defense by keeping the antibiotics out of the cell wall, by altering efflux or permeability. While the entry mechanisms have been identified for many antibiotic families, the mechanism of entry for aminoglycosides (AG) into bacterial cells has been a subject of debate for decades, and various non-exclusive mechanisms that are dependent or independent of the proton motive force (PMF) have been described^10–12^. The idea of AG entry into the cytoplasm through active transport systems, normally used for low-affinity transporters devoted to other molecules such as carbohydrates has been previously proposed, but to our knowledge never confirmed^12, 13^.

*Vibrio cholerae* is a Gram-negative bacterium and the etiological cause of cholera. *V. cholerae* has demonstrated the capacity to resist multiple antibiotics, including aminoglycosides (AGs) during different pandemics, notably during 2010 in Haiti ^1^. Moreover, sub-minimum inhibitory concentrations (sub-MICs) of a variety of antibiotics, including AGs, have been shown to trigger the SOS response and mutagenesis in this bacterium ^2–4^. These low-doses have been commonly found in the environment and shown to exert selective pressure upon the bacteria present ^5, 6^. They have been shown to modify global genes expression (visible in numerous transcriptome studies in the presence of antibiotics, e.g. ^7–9)^. Evidence supports a role for sublethal doses in the development of resistance ^10, 11^, notably by producing stress that can influence mutation rates and adaptive responses ^3, 12, 13^.

Here we have combined two high-throughput approaches upon sub-MIC treatment with the AG tobramycin, in order to identify genetic elements relevant to the adaptation of *V. cholerae* to sub-MICs AGs. RNA-seq was used to identify transcriptional alterations, and a high-throughput transposon (Tn) insertion mutants screen was used to define the fitness of *V. cholerae* during growth in presence of AGs. We identified a small non-coding RNA (sRNA) that exhibited a significant transcriptional downregulation in presence of tobramycin, and whose disruption by Tn insertion provided a significant fitness advantage. This sRNA, located in the 5’ untranslated region of the *malK* gene, was recently partially characterized in *Vibrio tasmaniensis* (*vsr217*). Its expression was shown to be induced by maltose, under the dependence of MalT regulation, allowing the full expression of *malK*, one of the components of a maltose ABC transport. *vsr217,* conserved across vibrios, was also demonstrated *t*o act *in trans* to repress the *fbp gene*, whose product is involved in gluconeogenesis ^14^.

In this work, we have been able to link this sRNA to the differential entry of AGs through the regulation of the expression of a subset of its sugar transporter targets. We decided to name this sRNA *ctrR* (<u>c</u>arbohydrate transport regulating <u>R</u>NA). We found that the cAMP receptor protein (CRP), responsible of activating transcription of target genes following binding to cAMP ^15^, regulates *ctrR* expression through binding to its promoter region. Thus this sRNA is part of the carbon catabolite repression (CCR) regulon, a regulatory mechanism by which the expression of genes required for the utilization of different sources of carbon is orchestrated ^16^.

The description of *ctrR* as a sRNA associated with the CCR, that regulates AG entry in the Vibrionales represents a new mechanism of active uptake of AGs, via carbohydrates transporters. Although no *ctrR* homolog has been found in other genera, the general conservation of CCR in bacteria, including in ESKAPE pathogens ^17^, and the fact that preliminary data show that AGs may use the same transporters in other species, opens the possibility of exploiting such a conserved molecular mechanism as a potential drug target.

## Results

### Genome-wide screens reveal a sRNA as significantly affected by tobramycin treatment

We used whole genome comparative transcriptional profiling to screen to for transcriptional differences triggered by treatment with low doses of AGs. *V. cholerae* cells were grown in the presence or absence of 0.02 µg/ml tobramycin (2% of the MIC). An intergenic region, identified as a sRNA (ncRNA586), displayed a high transcriptional activity during growth in rich media and was the ninth on the list of molecules with the highest transcription during exponential phase (Table S1). This region was previously identified as ncRNA by transcriptional screening and massively parallel sequencing ^18–20^. In presence of tobramycin, 102 genes presented a significant alteration in transcription (Table S1), including the chaperonins *groEL/ES* ^21, 22^ (Figure 1A & Table S1). Notably, these significant alterations in transcription activity occurred at AG doses that do not appear to impact growth ^3, 23^, indicating stress induction even in the presence of non-lethal doses of AGs. The region corresponding to the ncRNA586 was significantly downregulated (5-fold) (Figure 1B). Genes involved in carbon source utilization and transport, including genes of the *VC1820-1827* Phosphotransferase System (PTS) cluster involved in fructose and mannose transport/metabolism ^23^, maltose metabolism and transport genes *malEFG* (VCA0943-945), *malQ* (VCA0014) and *malS* (VCA0860), and the maltoporin *lamB* (VCA1028) were also found to be downregulated.

**Figure 1:**
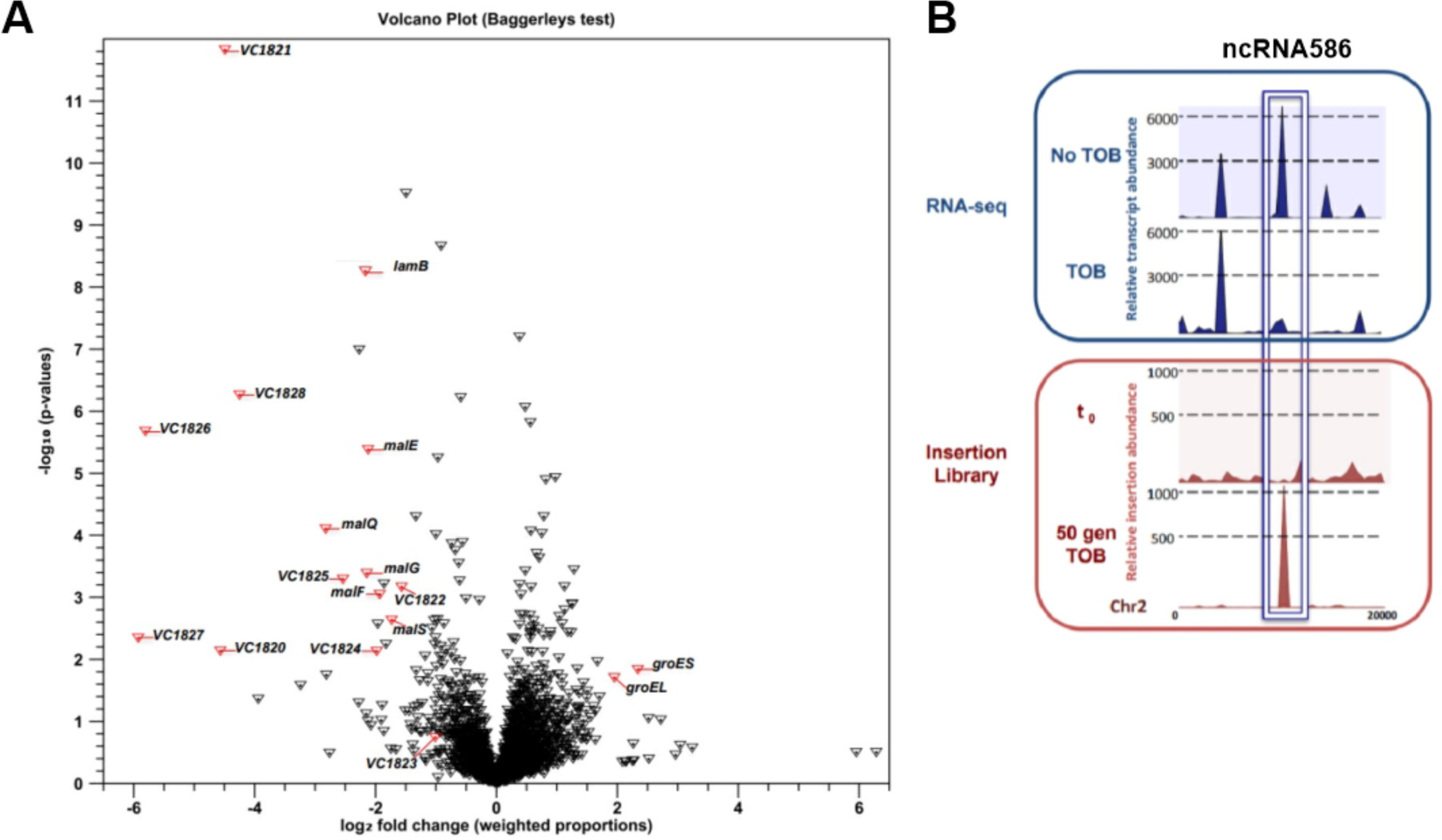
Identification of ncRNA586 and carbohydrate-related genes involved in tobramycin response. **A.** Volcano plot representing differential transcription of all *V. cholerae* genes under sub-MIC tobramycin growth. Each gene is represented by a inverted triangle. The x-axis indicates log2 fold change and -log10 p-value is plotted on the y-axis. Genes of interest are highlighted in red with their locus tags/gene names. **B**. Region of chromosome 2 where *ncRNA586* is encoded (white rectangle). The upper panel shows a histogram of transcriptional activity without tobramycin (No TOB) and treated with sub-MIC tobramycin (TOB). The lower panel displays histograms representing mapping of our fitness screen, at time point zero (t0) and after 50 generations passage in sub-MIC tobramycin (50 gen TOB).

A complementary high-throughput Tn insertion screen was performed in parallel. A library of 100 000 independent Tn insertion mutants was grown out for 50 generations in medium containing an identical 0.02 µg/ml of tobramycin (2% of the MIC) as in the transcriptional screen. The initial insertion library (t0) and the population obtained after 50 generations of tobramycin exposure (t50) were sequenced. The proportions of insertions obtained between t0 and t50 allowed us to identify genes for which inactivation provides a fitness advantage (enriched at t50) or disadvantage (depleted at t50). The insertional inactivation mutants of *ncRNA586* were the most enriched (highest fitness score) following tobramycin treatment (Figure 1B). This significant fitness advantage mimicked downregulation of this sRNA during growth of the WT control under similar sub-MIC selection. The involvement of this sRNA in AG response was further explored.

### *ncRNA586* deletion enable better tolerance to AG treatment

A Δ*ncRNA586* strain was constructed to test its role in fitness in the presence of tobramycin. Growth curves for the Δ*ncRNA586* mutant evidenced a decrease in susceptibility to tobramycin compared to the wild-type (WT) strain (Figure 2A). The mutant strain presents the same phenotype on solid medium where growth on 1.5 µg/ml of tobramycin is observed at dilutions 3 orders of magnitude greater than the WT (Figure 2B). Treatment with three other AGs (kanamycin, gentamicin and neomycin) resulted in similar results, suggesting that the mechanism that enables Δ*ncRNA586* to resist tobramycin applies to this class of antibiotics (Figure S1). Tolerance was not observed when challenged with representatives from five other antibiotic classes (β-lactam: ampicillin/carbenicillin; fluoroquinolone: ciprofloxacin; antifolate: trimethoprim; transcription blocker: rifampicin; and another protein synthesis blocker: chloramphenicol; Figure S1). These results suggested that antibiotic tolerance in the Δ*ncRNA586* strain is specific to AGs. Since *ncRNA586* is involved in the susceptibility to AGs, we looked for targets to explain its modulating role in response to AGs.

**Figure 2:**
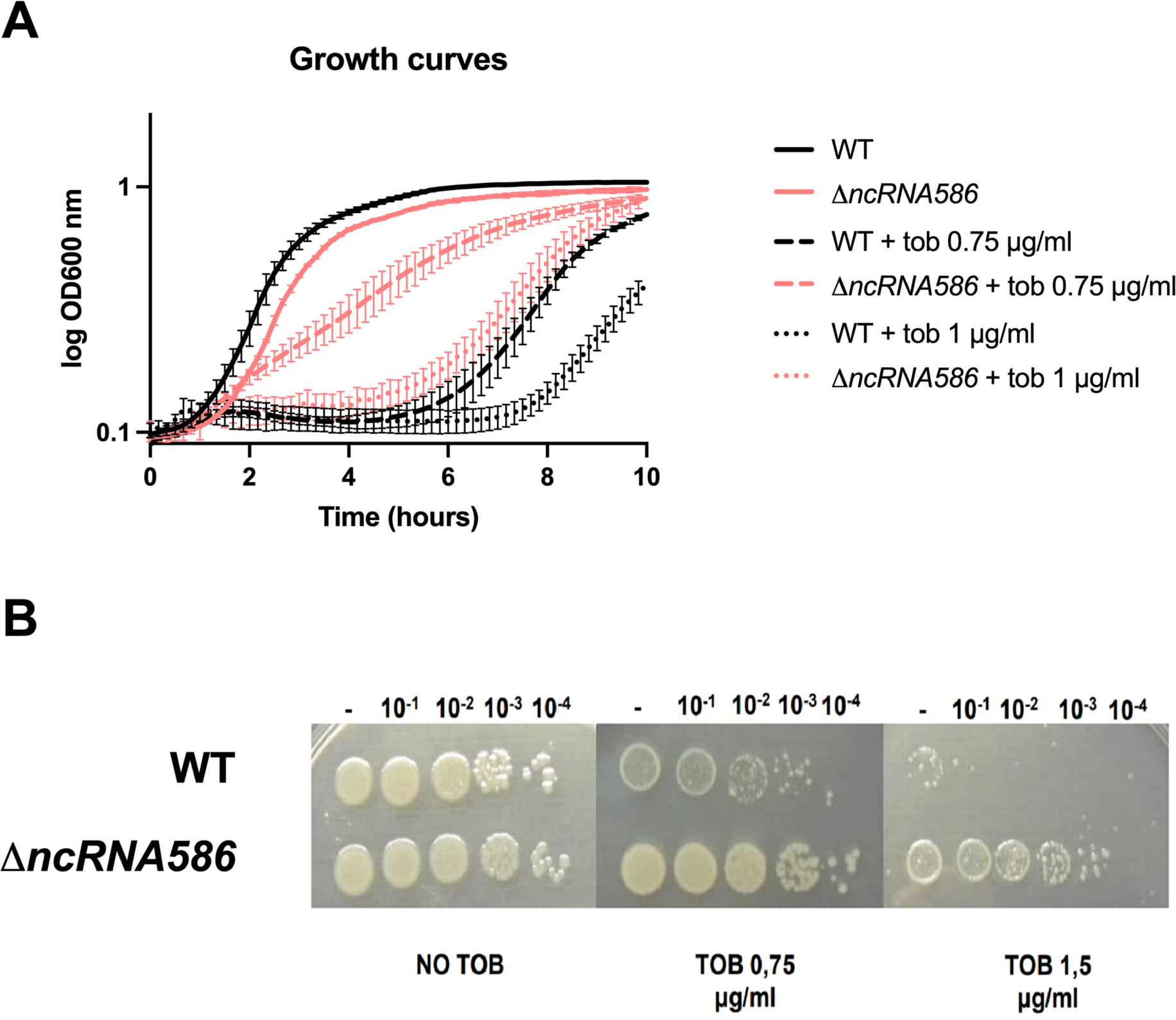
*ncRNA586* deletion decreases susceptibility to tobramycin. **A.** Growth curves of WT and Δ*ncRNA586* strains treated or not with 0.75 or 1 μg/ml tobramycin for 10 hours. **B.** Serial dilutions of WT and Δ*ncRNA586* spotted on plates containing or not 0.75 or 1.5 μg/ml of tobramycin.

### Bioinformatic analysis reveal carbohydrate transport genes as putative hybridization targets for *ncRNA586*

The homolog of *ncRNA586* in *V. tasmaniensis*, *vsr217*, was found to be highly induced during growth in thalassic conditions^24^. It was shown to regulate *malK*, one of the components of an ABC maltose transporter by forming a long *vsr217-malK* transcript)^14^. We confirmed that the *ncRNA586-malK* transcription was conserved in *V. cholerae* and was functionally homologous in also forming a long transcript (Figure S2).

RNA-seq data (Figure 1A) demonstrated that genes involved in the metabolism/transport of carbohydrates were downregulated, seemingly commensurate with downregulation of *ncRNA586*, in the presence of tobramycin.

This observation inspired us to perform mRNA target predictions for *ncRNA586* using a number of complementary bioinformatic tools. We used bioinformatic tools sTarPicker^25^ and RNApredator^26^ to detect RNA-protein interactions, and IntaRNA^27^ to probe for RNA-RNA interaction. Aggregated results revealed that the majority of potential *ncRNA586* regulatory targets are involved in transport or metabolism of different carbohydrates (Table S1). Top putative targets (probability score>0.9) included genes whose RNA-seq analysis indicated down-regulation during AG treatment: *VC1821* (subunit of VC1020-21 PTS), *VC1827* (*manA,* mannose isomerase, operon with PTS *VC1826*), *malE* (VCA0943) and *lamB* (VCA1028) (Table S2).

These data highlighted a link between this *ncRNA586* and carbohydrate transporters mediating AG response. We therefore named this sRNA in *V. cholerae* “*ctrR*” for <u>c</u>arbohydrate transport regulating <u>R</u>NA.

### *CtrR* and *vsr21*7 have common AG response properties

*The ctrR* sequence is conserved across the Vibrionaceae ^14^. We used RNAalifold ^28^ to predict the secondary structure of potential homologs. This analysis revealed high secondary degree of structural conservation despite sequence divergence (Figure S2B). Then we applied CopraRNA ^27^, in a comparative genomics approach, to compute whole genome target predictions for *ctrR* homologs. Three candidate homologous sRNA sequences from three distinct *Vibrio* species: *V. cholerae*, *V. parahaemolyticus* and *V. tasmaniensis (vsr217)* were used as inputs. This analysis revealed that putative targets for *V. parahaemolyticus* and *V. tasmaniensis* included all homologous genes previously identified as putative targets for *ctrR in V. cholerae,* which were shown to be downregulated under tobramycin treatment (Table S2).

We next performed an orthologous expression experiment in order to ascertain whether *vsr217* involvement in AG response was comparable to that seen in *V. cholerae*. We overexpressed *ctrR (*p*ctrR+)* and *vsr217 (*p*vsr217+)* on separate plasmids and tested those plasmids in both *V. cholerae* or *V. tasmaniensis* (Figure 3A). In *V. cholerae*, *ctrR* overexpression resulted in increased sensitivity to tobramycin, contrary to the deletion which resulted in better growth (Figure 2A). This was not, however, the case for *vsr217* overexpression (Figure 3B). Similar results were obtained for *V. tasmaniensis,* where the overexpression of its native *vsr217* also sensitized the bacteria to tobramycin and *ctrR* overexpression showed no discernable phenotype (Figure 3C). Collectively, these data suggest species-specific coevolution of this sRNA with a conserved mode of action.

**Figure 3:**
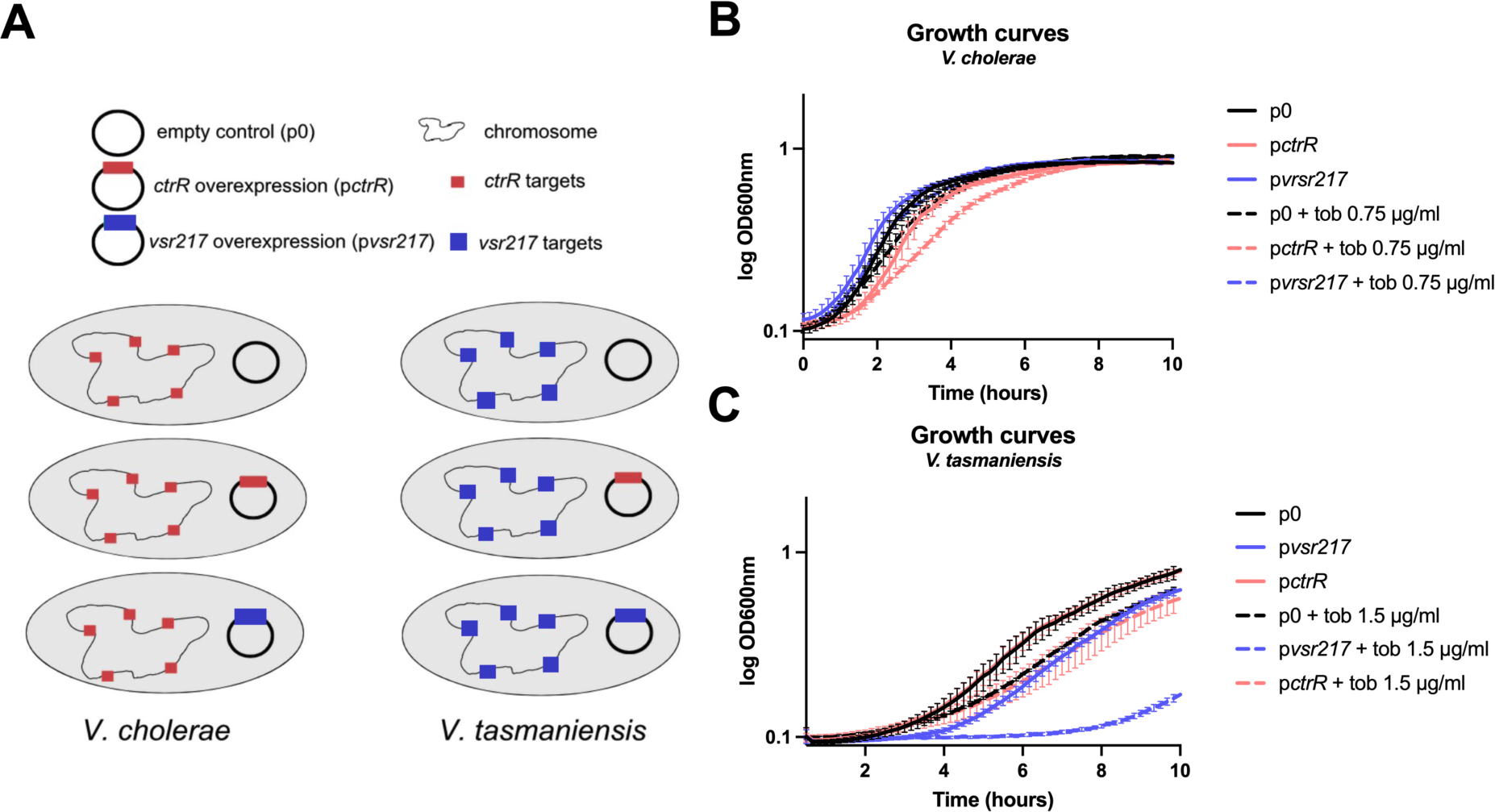
Species specific effect of *ctrR*. **A.** Experimental strategy. Overexpression of *ctrR (*p*ctrR), vsr217 (*p*vsr217)* and an empty control plasmid (p0) was performed in *V. cholerae* or *V. tasmaniensis*. **B.** Growth of *V. cholerae* overexpressing or not *ctrR* or *vsr217*, treated or not with tobramycin. **C.** Growth of *V. tasmaniensis* overexpressing or not *ctrR* or *vsr217*, treated or not with tobramycin.

### *CtrR* and carbohydrates transporters are involved in differential AG entry

Since *ctrR* appears to regulate carbohydrate transporters, we hypothesized that these transport systems could be involved in facilitating the entry of AG-class of antibiotics into *V. cholerae*. We tested this hypothesis using a Cy5 labeled AG, Neo-Cy5 ^29^, and flow cytometry to detect fluorescence intensities in *ctrR* deletion or overexpression strains. The Δ*ctrR* strain showed a lower percentage of fluorescent cells compared to the WT strain, reflecting a reduced entry of Neo-Cy5, consistent with the fact that this mutant is less susceptible to AGs (Figure 4A). Percentage of fluorescent cells in the *pctrR+* strain overexpressing *ctrR,* more susceptible to tobramycin, was higher than in the empty plasmid control (p0) (Figure 4B). During the course of the experiment, we noted that cell morphology, as measured by forward scatter (FSC) and side scatter (SSC) was altered in a higher percentage of cells in the *pctrR*+ overexpression population (41.2%) versus the control strain (21.09%) (Figure S3). This is potentially due to membrane alterations related to the surface proteins, such as carbohydrate transporters targeted by the sRNA. Collectively, these results support that differential entry of AGs in *V. cholerae* can be modulated by *ctrR*.

**Figure 4:**
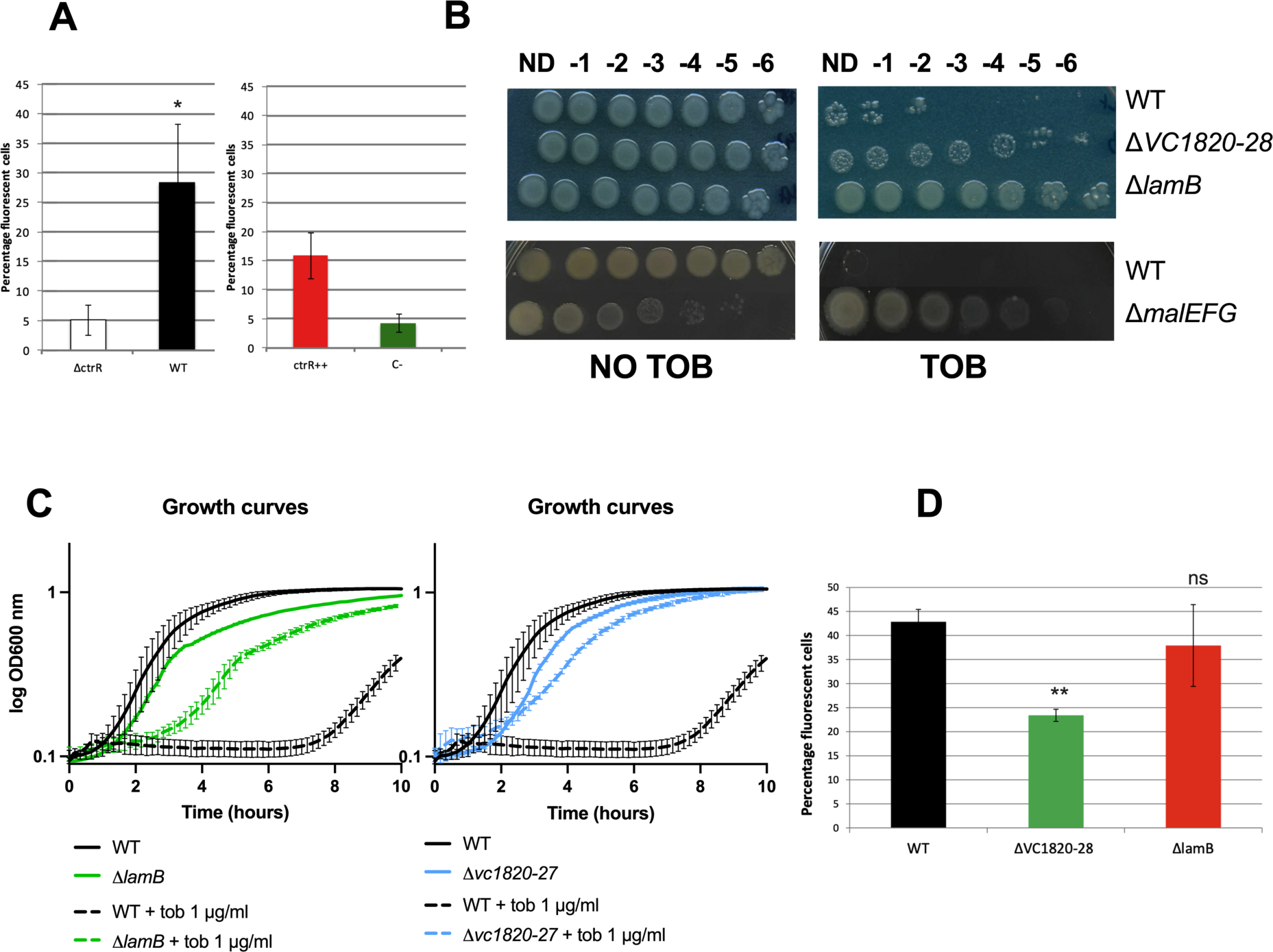
*ctrR* and carbohydrates transporters are involved in AG susceptibility and impact AG entry. **A.** Uptake of Neo-cy5 in the WT strain comparing to Δ*ctrR*, and the p0 strain compared to p*ctrR,* expressed in percentage of fluorescent cells in the population. The two experiments were independent. **B.** Serial dilution of the WT strain, ΔVC1820-27, Δ*lamB*, and Δ*malEFG* on plate containing or not 1.5 μg/ml of tobramycin. **C.** Growth curves of WT strain, ΔVC1820-27, Δ*lamB* in the presence or not of tobramycin. **D.** Uptake of Neo-cy5 in the WT strain compared to ΔVC1820-27 and Δ*lamB*, expressed in percentage of fluorescent cells in the population.

The three top predicted targets encode different types of transport systems, which led us to question whether these transport systems could affect AG entry. We constructed deletion strains to test functionality: Δ*VC1820-27* (cluster containing four PTS), Δ*lamB* (maltoporin), and Δ*malEFG* (ABC type maltose transporter). All three mutants showed improved resistance to tobramycin on solid medium (Figure 4B) as well as an increase MIC (Etest) of tobramycin (Table 1). It should be noted that the *malEFG* deletion itself is deleterious for growth. The MIC values for different families of antibiotics was also measured in the Δ*VC1820-27* and *ΔlamB* strains: ciprofloxacin, trimethoprim and carbenicillin. Gentamicin was also tested as an additional AG. Increased MICs in the deletion strain was only observed for AGs (Table 1).

**Table 1:** MIC to different antibiotic (µg/ml)

|  | tobramycin | gentamicin | ciprofloxacin | trimethoprim | carbenicillin |
| --- | --- | --- | --- | --- | --- |
| *V. cholerae* WT | 1 | 0.5 | 0.002 | 0.5 | 1.5 |
| $\Delta VC1820-27$ | 2.5 | 2.5 | 0.002 | 0.4 | 1.5 |
| $\Delta lamB$ | 1.5 | 2 | 0.002 | 0.25 | 1.5 |
| $\Delta malEFG$ | 1.5 | | | | |
|  | sugar 1% |  |  |  |  |
| MIC tobramycin | glucose | mannose | maltose |  |  |
| *V. cholerae* | 3.5 | 2.5 | 1.5 |  |  |
| *E. coli* | 2.75 | 2 | 1.75 |  |  |
| *P. aeruginosa* | 3 | 1.5 | 1.25 |  |  |
| *K. pneumoniae* | 2.75 | 1.25 | 1.25 |  |  |
| *A. baumannii* | 2.5 | 1.75 | 1.5 |  |  |

The Δ*VC1820-27* and Δ*lamB* mutants also showed improved resistance to tobramycin in liquid medium by performing growth curves in the presence or absence of tobramycin (the Δ*malEFG* mutant could not be tested due to aggregate formation in these conditions) (Figure 4C). This result indicates that these transporters, and predicted targets of *ctrR*, are involved in the susceptibility to AGs.

To confirm their involvement in AG entry, we treated the mutants with Neo-Cy5 and assessed the fluorescence level by flow cytometry analysis. The Δ*VC1820-27* strain registered half the level of percentage of fluorescent cells compared to the WT after 15 minutes of treatment with Neo-Cy5 (Figure 4D). Interestingly, the Δ*lamB* strain showed no differences in fluorescence. We believe that the predicted size of the LamB porin channel, able to transport molecules of up to 600 Daltons in size ^27^ could transport tobramycin (467 Da) and potentially neomycin (614 Da). However, it is likely that the conjugated Neo-Cy5 molecule (667 Da) is too large. The Δ*malEFG* strain was once again not analyzed through flow cytometry as its rugose morphology resulted in irregular fluorescence readings. Taken together, these data support the hypothesis that differential AG entry is mediated by *ctrR* regulation of carbohydrate transporters.

### *ctrR* regulates AG entry by regulation of its target

We investigated the mechanism behind the *ctrR* regulation of its top predicted target *manA* (VC1827, in operon with the VC1826 PTS transporter). ManA, a mannose isomerase, is implicated in perosamine synthesis ^30^, which is essential for O-antigen synthesis. ManA is also the receptor for phages ICP1 and ICP3 where expression determines phage sensitivity. If *ctrR* regulates *manA*, then *ctrR* overexpression should affect sensitivity to phage infection. Serial dilutions of ICP1 and ICP3 phages stocks were spotted on lawns of control (p0) and overexpression (*pctrR+*) strains. We observed a 2-log difference in sensitivity in the overexpressing strain relative to the control (Figure 5A). These results were similar for both phages, indicating that more phage receptors were available upon *ctrR* overexpression.

**Figure 5.**
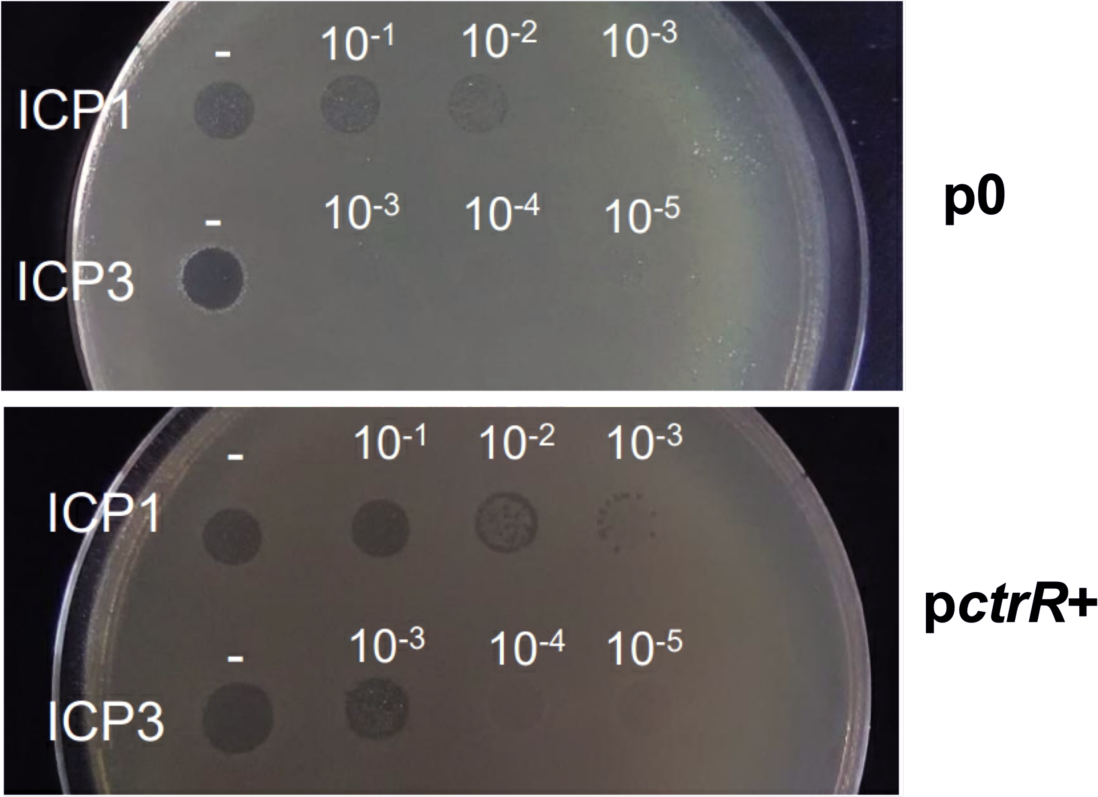
*ctrR* acts on *manA* mRNA transcription or stabilization. Susceptibility of the empty vector strain (p0) compared to the one overexpressing *ctrR* (p*ctrR*+) to ICP1/3 phages through serially diluted spots.

This result suggests that *ctrR* overexpression promotes a higher level of *manA* (VC1827) transcript, potentially by stabilization of the target mRNA as in the *vsr217* homolog ^14^.

As *ctrR* overexpression appeared to stabilize *manA* transcripts, we investigated potential *ctrR-manA* mRNA base-pairing. Secondary structure prediction for *ctrR* revealed that the stretch of nucleotides (152-161) was predicted to interact with putative targets, mostly single stranded, with the exception of three residues that participate in formation of a short stem (Figure S4A). We introduced two point-mutations in our overexpression vector (guanine > cytosine at positions 157 & 158) in the *ctrR* region predicted to base-pair with targets *VC1827*, *malE* and *lamB*. The resulting plasmid and strain are referred to as p*ctrR*157 and *ctrR*157 respectively (Figure S4B). The introduction of the p*ctrR*157 abolished the tobramycin susceptibility phenotype induced by the unaltered *pctrR+* (Figure S4C). It is worth noting that the 3’ region of the *vsr217* homolog is necessary for its own stabilization, that include these two bases after alignment of the sequences ^14^.

### *ctrR* is regulated by CCR

The link between *ctrR* and sugar metabolism systems suggested that *ctrR* might be regulated by the cAMP-CRP complex. As with *ctrR* binding we initially used bioinformatic tool; a virtual footprint tool (PRODORIC) ^31^, identified motifs as belonging to a CRP binding box with matching pattern scores between 4.18 and 5.12 (max = 10), when compared to known CRP binding sites (Table S3). The Softberry tool ^32^ also identified a CRP binding-box (score 10, Table S3). Based on the favorable bioinformatic result, we investigated the level of *ctrR* relative to the presence or absence of CRP. When CRP was deleted, dRT-PCR measured a greater than 4-fold decrease of *ctrR*, arguing for a role of CRP in the regulation of its expression (Figure 6A).

**Figure 6:**
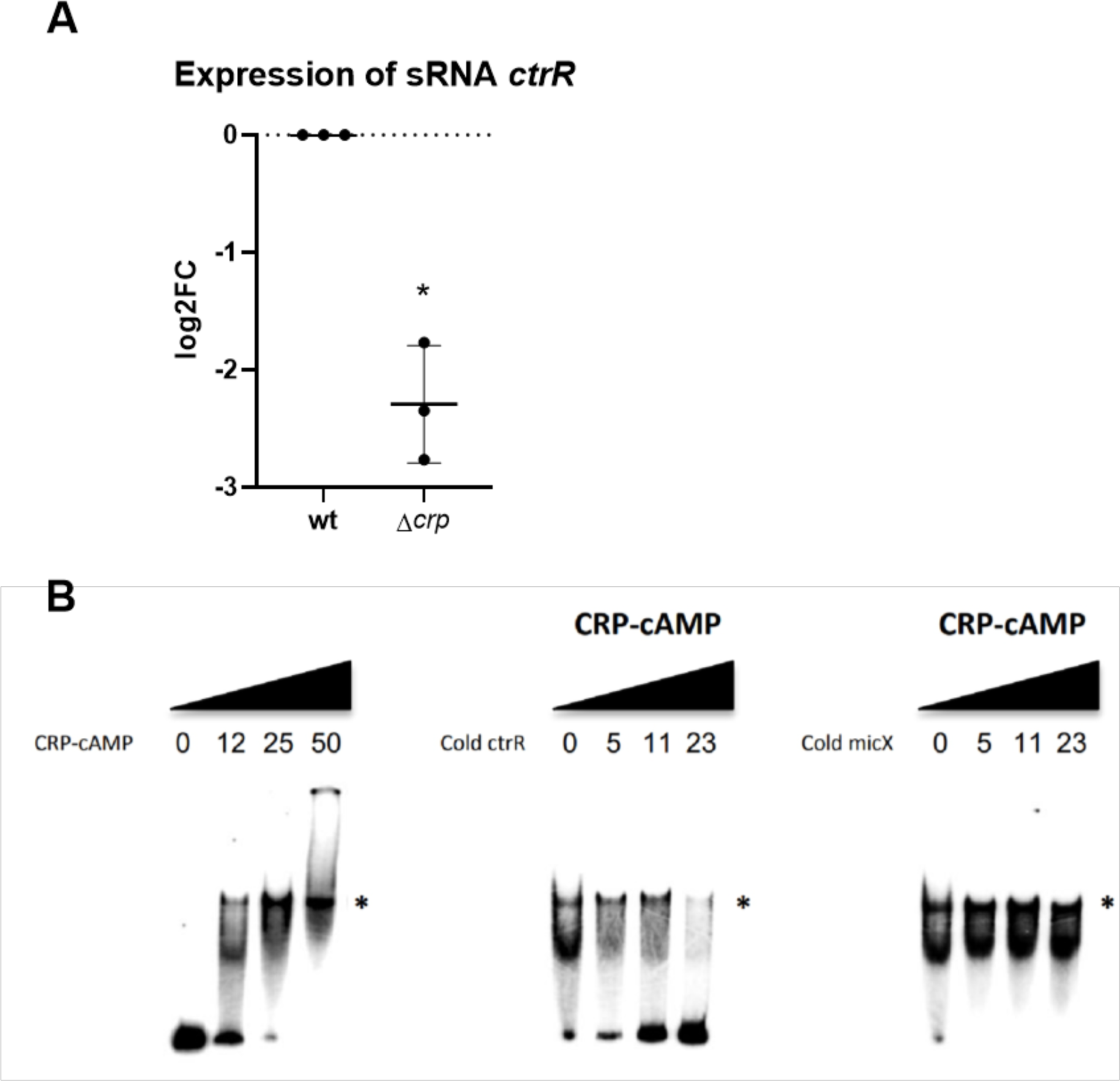
CRP regulates *ctrR*. **A.** dRT-PCR measurements of *ctrR* on *V. cholerae* WT compared to *Δcrp* total RNA (log2FC compared to WT). **B.** Binding of cAMP+CRP on *ctrR* promoter (left), depending on the concentrations (in μM indicated above the bands) (shift is highlighted by the *). Competitive binding with cold *ctrR* PCR product (middle) and competition with *micX*’s promoter (right) were used as controls.

We then performed an electrophoretic mobility shift assay (EMSA) on *the* ctrR promoter region (500 bp amplicon containing the predicted binding boxes) using the cAMP-CRP complex. Figure 6B shows that CRP is able to bind and produce a shift in the observed size of the labeled *ctrR* region. Addition of unlabeled competitor *ctrR* is seen to titrate away this binding in a specific manner compared to the negative control sRNA *micX* promoter, itself a known to regulate a surface transporter ^33^.

CRP regulation of *ctrR*, together with *ctrR* regulation of different carbon transporters, led us to possibility that different carbon sources could affect AG entry in *V. cholerae.* The MIC of tobramycin was tested in the presence of glucose, maltose or mannose. The baseline 3.5 µg/ml MIC of tobramycin obtained for glucose was found to be 1.5 µg/ml for maltose and 2.5 µg/ml for mannose (1% of sugar and bactotryptone). Due to conservation of CCR across a large number of bacterial species, we performed the same assay in other Gram-negative ESKAPE pathogens ^17^ *Klebsiella pneumoniae*, *Acinetobacter baumannii* and *Pseudomonas aeruginosa*, and in *Escherichia coli.* In the four tested pathogens, the MIC was also the highest in glucose, and the lowest in maltose (Table 1).

## Discussion

AG entry mechanisms in bacteria have been discussed around several non-exclusive hypotheses ^34–37^. While PMF-dependent uptake is now a recognized mechanism, AG entry into the cytoplasm through active transport systems has been proposed without experimental evidence ^38^. As a matter of fact, apart from the AG modifying genes, most of the genetic mutations associated with AG resistance are either found on ribosome associated genes, or in genes of the respiratory complexes which decrease respiration and PMF ^39, 40^.

The study of the *V. cholerae* genetic response to sub-MIC tobramycin allowed us to identify the sRNA c*trR* as an actor in optimal growth and adaptation to AG antibiotics. We further showed that its influence on AG susceptibility is linked to target carbohydrate transporter systems. c*trR* (this study) and its *V. tasmaniensis* homologue ^14^ were found to regulate mRNA levels of genes encoding for carbohydrate transporter proteins. The fact that *ctrR* regulates expression of these transporters and the link with AG susceptibility prompted us to measure AG uptake by the bacterial cells in their absence, leading to the discovery of a new mechanism of active AG uptake enabled by sugar transporters in the Vibrionaceae. This additional route of entry for AGs does not challenge the PMF-dependent uptake, and these two ways of entry could be linked ^41^. The fact that AG resistance through mutations in sugar transporters hasn’t previously been identified is likely due to the fact that AGs are transported by a variety of sugar transporters, which thus show redundancy in AG uptake. For instance, novel genome editing techniques helped dissect the PTS system in *V. cholerae*, revealing high redundancy and little specificity for carbon transport systems ^42^. Hence, inactivation of one transporter is not expected to yield a high AG resistance. Moreover, inactivation of sugar transporters could have a strong negative impact on fitness which could explain that such mutants are not selected in nature.

AGs contain an amino-sugar moiety, which could explain their uptake by sugar transporters. In this scenario, bacterial sugar transporters would be mistaking AGs for carbon sources, triggering CCR and facilitating antibiotic entry. The link between cAMP-CRP regulated CCR and antibiotic resistance in Gram-negative bacteria was first suggested in 1978, when mutants in genes *cya* and *crp* were selected for their resistance to fosfomycin and the AG streptomycin. Moreover, it was observed that glucose decreased the killing effect of AGs in *E. coli* ^38, 43^, and that carbon source affect AG susceptibility in *Pseudomonas aeruginosa* ^44^ or *E. coli* ^45^, through an effect on PMF. However, there was no evidence regarding a potential CRP-linked route of entry for AGs in bacteria. The fact that the c*trR* RNA, which regulates sugar transporters’ mRNA levels, is also under the dependence of CCR, constitutes such a link in Vibrionaceae.

*ctrR* was identified here under antibiotic stress conditions. A number of sRNAs have been shown to modulate antibiotic tolerance by base pairing with mRNAs encoding functions relevant for resistance such as drug efflux pumps (*dsrA* in *E. coli*), metabolic enzymes (*glmY* in *E. coli* and *Salmonella* spp.), or transport proteins (*micF* in *E. coli* and *Salmonella* spp.) ^46^. *ryhB* sRNA is involved in AG resistance in response to low levels of iron availability ^47^.

*ctrR* is conserved among vibrios, despite species-specific co-evolution with targets. The fast pace at which sRNA sequences change is one of several factors that make systematic studies of bacterial sRNAs challenging. As proven by the *ctrR* example, high degrees of intra- and inter-species polymorphism yield low sequence similarity, especially as compared to protein-coding genes. Some of the known constraints for sRNA evolution include a rho-independent terminator, double-stranded regions that allow for stable secondary structure, an unstructured seed region, where the sRNA base pairs with its target and, typically, an environmentally regulated promoter ^48^. Following target validation, the search for homologs of this sRNA in additional bacterial species, inside and out of the Vibrionaceae yielded no results through a sequence identity search, which contributes to significant challenges in tracing sRNAs across evolutionary distances ^34^. Despite the apparent lack of *ctrR* homologs outside the *Vibrionaceae*, the conservation of this mechanism for AG transport in different gram-negative bacteria should be further explored.

Interestingly, a recent study conducted on the Gram-positive bacterium *Bacillus subtilis* provides another example of RNA dependent regulation of sugar transporters. The study discovered an RNA thermometer that regulates the translation of glycerol permease, involving differential structuring of the 5’ UTR region of the transporter’s mRNA in response to external temperature ^49^. This illustrates the potential for RNA-dependent regulation mechanisms in addition to general catabolite control, adding another layer of complexity to the regulation of sugar transporters and their impact on AG uptake.

Collectively, the functional characteristics we describe for *ctrR* include species specificity, environmental regulation, and carbon source dependence, making this mechanism an attractive target for development of novel therapeutic strategies against gram-negative pathogens. This study uncovers an alternative entry route for AGs in the Vibrionaceae and possibly opens the door for much needed, novel, CCR related therapeutic strategies that will aid in the face of the current antibiotic pipeline decline.

## Methods

### Bacterial strains, plasmids, primers

Strains, plasmids and primers used in the study are presented in Table S3.

The *Vibrio cholerae* O1 biovar El Tor N16961 was used in this study. *A. baumannii*, *E. coli*, *P. aeruginosa*, and *V. cholerae* were grown on MH (Muller Hinton) medium, supplemented when appropriated with 0.4% of glucose, maltose or mannose. Overexpression mutants were grown in MH medium supplemented with carbenicillin for plasmid maintenance (100 µg/ml) and 0.2% arabinose to induce the promoter for pBAD24 (The effect of arabinose on antibiotic susceptibility was tested and deemed neutral).

### Allelic exchange and mutant constructions

Gene deletions were constructed using derivatives of an of an R6K γ-ori-based suicide vector, pSW7848 as described in ^50^. Briefly, we assembled through Gibson 500 bases pair homologous regions upstream and downstream of the gene of interest with the *aph* resistance gene for kanamycin surrounded by *frt* sites into the pSW7848 plasmid. Amplification of upstream and downstream regions for strain Δ*VC1820-27* was performed using oligos 5vc181, 6vc181, 7vc181 and 8vc181. Deletion of *ctrR* was performed as described ^18^. Amplification of upstream and downstream regions for strain Δ*lamB* was performed using primers 5lamB, 6lamB, 7lamB and 8lamB. Amplification of upstream and downstream regions for strain Δ*malEFG* was performed using primers 5malop, 6malop, 7malop and 8malop. These plasmids were transformed into π3813 cells, after colony PCR and plasmid extraction were transformed into β3914 donor cells^50^. Mutagenesis was then performed through conjugation for 24 hours between the target strain and the plasmid containing β3914 strain. Once deletion mutants were validated by appropriate PCR, conjugation with a strain containing plasmid pMP108 encoding the *frt* specific flippase^51^ was performed for 4 hours in order to excise the resistance cassette.

### Overexpression of ctrR and vsr217 from V. cholerae and V. tasmaniensis

Overexpression of *ctrR* and *vsr217* were achieve by PCR amplification using respectively primers 4347/4348 and ML435/436 and cloning into a pBAD24. For *ctrR*157, point mutation was introduced using primer ML466. Expression of genes were realized by addition of 0.2% of arabinose.

### RNA-seq

The MIC of tobramycin during growth in MH medium was determined to be 1 μg/ml.

Overnight cultures of the O1 biovar El Tor N16961 *V. cholerae* strain were diluted 100× and grown in Mueller-Hinton medium until an OD_600nm_ 0.4, with or without tobramycin 0.02 µg/ml (2% of the MIC). Samples were collected and total RNA was extracted as previously described ^18, 52^. Directional libraries were prepared using the TruSeq Stranded mRNA Sample preparation kit (20020595) following the manufacturer’s instructions (Illumina). 51-bp Single Read sequences were generated on the Hiseq2000 sequencer according to manufacturer’s instructions (Illumina). Reads were processed, quality checked, mapped, counted and normalized as previously described ^53^. Additionally, histograms of transcriptional activity across different chromosomes and conditions were compared in order to identify transcriptional differences in previously unannotated regions of the genome. For this, the same mapping tool used for RNA-seq was used in combination with CLC’s visualization track tool (CLC Bio). The data for this RNA-seq study has been submitted in the GenBank Sequence Read Archive (SRA) under project number: PRJNA506714.

### RNA *in silico* analyses

RNA structure was analyzed using the RNA folding program RNAfold ^54^. sTarPicker, RNApredator, IntaRNA and CopraRNA were used as tools for sRNA target prediction ^25–27^. Putative targets were considered when two different algorithms identified them as likely targets. RNApredator, IntaRNA and sTarPicker were used to determine potential targets within the *V. cholerae* genome. CopraRNA was then applied for conserved target identification using 3 homologous sRNA sequences from 3 distinct organisms: the *V. parahaemolyticus* (NC_004603), *V. tasmaniensis* (NC_011753) and *V. mimicus* (NZ_CP014042) genome sequences. Custom parameters were used for CopaRNA and IntaRNA in order to search to extract both sequences around the stop or start codon. Finally, RNAalifold ^28^ was used to determine secondary structure conservation between *ctrR* and *Vsr217* with default settings.

### High-throughput mutant screen

The MIC of tobramycin during growth in MH medium was determined to be 1 μg/ml.

A saturated mariner mutant library was generated by conjugation of plasmid pSC189 from *E. coli* to *V. cholerae* as previously described ^55^. Briefly, pSC189 ^56^ was delivered from *E. coli* strain 7257 (β2163 pSC189::spec, laboratory collection) into *V. cholerae*. Conjugation was performed for 2 h on 0.45 µM filters, and treated as previously described ^57^. After validation, the libraries were passaged for 50 generations with or without 0.02 µg/ml of tobramycin (2% of the MIC). Sequencing libraries were prepared using Agilent’s sureselect XT2 Kit with custom RNA baits designed to hybridize the edges of the Mariner transposon. A total of 12 cycles were used for library amplification. Agilent’s 2100 bioanalzyer was used to verify the size of the pooled libraries and their concentration. Ion Torrent Ion PGM sequencing technology was used producing 150bp long reads. Reads were then filtered through transposon mapping to ensure the presence of an informative transposon/genome junction using a previously described mapping algorithm ^55, 57^. Detection of at least 10 nucleotides of the transposon sequence were considered sufficient to retain a read. Informative reads were extracted, mapped and counted. Fitness scores were then calculated according to van Opijnen ^58^.

### Digital RT-PCR

RNA extraction and digital RT-PCR were performed as described ^21^. Cultures were diluted 1000X and grown in MH medium until OD_600nm_ of 0.4, and pellet were resuspended in 1:1.5 TRIzol (Invitrogen). Then, 300 µl of chloroform was added for 5 minutes and samples were centrifuged. A volume of 1:1 of 70% ethanol was mixed with the upper phase before transfer on column (RNeasy Mini kit, Qiagen), and RNA was extracted according to manufacturer directions. After purification, DNase treatment was realized with the Turbo DNA-free kit (Ambion) according to the manufacturer’s instruction. qRT-PCR reactions were prepared with 1 μl of RNA samples using the qScript XLT 1-Step RT-qPCR ToughMix (Quanta Biosciences, Gaithersburg, MD, USA) within Oppale chips during 10 minutes at 50°C before PCR, on the Naica Geode. Primers and probes are listed in Table S3. Image acquisition was performed using the Naica Prism3 reader. Results were acquiered and analyzed using Crystal Reader and Crystal Miner softwares. Values were normalized against expression of the housekeeping gene *gyrA* ^59^.

Primers and probes used for amplification are ML454/455/456 for *ctrR* (Table S3).

### Determination of antibiotic susceptibility

Antibiotic susceptibility was assessed in MH medium. The Tecan infinite 200 was used to measure bacterial growth in liquid media. Cultures were diluted 200X in 200 µl of culture medium per well. Plates were incubated using the following conditions kinetic run time of 20 hours. Experiments were done in biological triplicates.

MIC were determined using Etest (Biomérieux). Briefly, overnight cultures were diluted 20X in PBS and 300 µl were plated in appropriated medium. Plates were dried 10 minutes and Etest was added.

Susceptibility on plate was determined by serial dilution of bacterial overnight cultures, 5 µl of each dilution were spotted on MH plates supplemented or not with tobramycin.

### Neocy5 uptake

Experiments were performed as described ^60^. Briefly, overnight cultures were diluted 100 fold MOPS Rich medium. When the bacterial strains reached an OD of 0.25, cells were incubated them with 0.4 µM of Cy5 labeled Neomycin for 15’at 37**°**C. Then, 20 µl of the incubated culture were used for flow cytometry, diluting them in 200 μl of PBS before reading. Flow cytometry experiments were performed as previously described ^61^. For each experiment, 50 000 events were counted on the Miltenyi MACSquant device. The Y3 fluorescence channel was then used to measure fluorescence intensity.

### Phage infections

Overnight cultures of the *pctrR*^+^ strain and a control empty plasmid (p0) were diluted 100 times in LB medium. When these cultures reached an OD_600nm_ of 0.15, 10 mM CaCl_2_ were added to the cultures. When these cultures reached an OD_600nm_ of 0.3, 10 μl diluted of cultures were diluted in 2 ml of PBS and plated on MH plates containing 10 mM CaCl_2_. Serial dilutions of phages ICP1 (10^-1^ to 10^-3^) and ICP3 (10^-1^ to 10^-5^) (kindly provided by Andrew Camilli) were spotted on the plates to measure the infectivity of these phages. Experiments were performed in triplicate.

### CRP EMSA

The promoter of *ctrR* was predicted with Bprom (Softberry). A 500-bp DNA fragment including the promoter region of *ctrR* was amplified from *V. cholerae* N16961 genomic DNA by PCR using Dreamtaq (Fermentas) and oligonucleotides crp500F and crp500R (one of which was labeled with [γ32P]ATP by use of T4 polynucleotide kinase (NEB)). The PCR product was purified with the Nucleospin Gel and PCR clean-up kit (Macherey-Nagel). The binding of cAMP-CRP (kindly provided by Annie Kolb) to the 500-bp DNA fragment was performed as previously described ^62^. In order to test the specificity of CRP binding, non-radioactive *ctrR* and *micX*’s promoter region were used as positive and negative controls for competitive binding. Similarly, to *ctrR*, a 500 bp amplicon containing *micX*’s promoter region was used.

### Statistical analysis

F-test determine whether the variances were equal or different between conditions. For conditions with equal variance, Student’s t-test was used. For conditions with different variances, Welch correction was applied. One-way ANOVA or two-way ANOVA were used for multiple comparisons, with Bonferroni correction, to determine the statistical differences between groups. ∗∗∗∗ means p<0.0001, ∗∗∗ means p<0.001, ∗∗ means p<0.01, ∗ means p<0.05. Number of replicates for each experiment was n=3. For growth curves, averages with standard deviations were plotted. For logarithmic values, means and geometric means were plotted.

## Authors contributions

Conceptualization, S.A.P., Z.B. and D.M.; Methodology, S.A.P., M. L., M. E. V., Z.B., R. L.-I, D.F. and D.M.; Investigation, S.A.P., M. L., R.L-I., E.K., and Z.B.; Writing – Original Draft, S.A.P., M. L.,; Writing – Review & Editing, S.A.P., M. L., Z.B., S. P. K., D.F. and D.M.; Funding Acquisition, D.M., Z.B.; Resources, S. P. K., M. E. V.

## Aknowledgments

We thank Ivan Imaz for his help with the bioinformatics analysis, Manas Sabeti for helpful discussions, and Andy Camilli for providing us with phages ICP1 and ICP3. We thank Odile Sismeiro and Jean-Yves Coppee for RNA-seq analysis. This work was supported by the Institut Pasteur, the Centre National de la Recherche Scientifique (CNRS-UMR 3525), EU-PLASWIRES 612146/FP7-FET-Proactive (R.L.-I. salary), the Fondation pour la Recherche Médicale (Grant No. DBF20160635736), ANR Unibac (ANR-17-CE13-0010-01) and the Fondation pour la Recherche Médicale (équipe FRM 202103012569). S.A.P was the recipient of a long-term post-doctoral fellowship from Roux-Cantarini foundation.

