## Supplemental figures and tables for "Identification of the active mechanism of aminoglycoside entry in *V. cholerae* through characterization of sRNA *ctrR,* regulating carbohydrate utilization and transport"

### Supplementary figures:

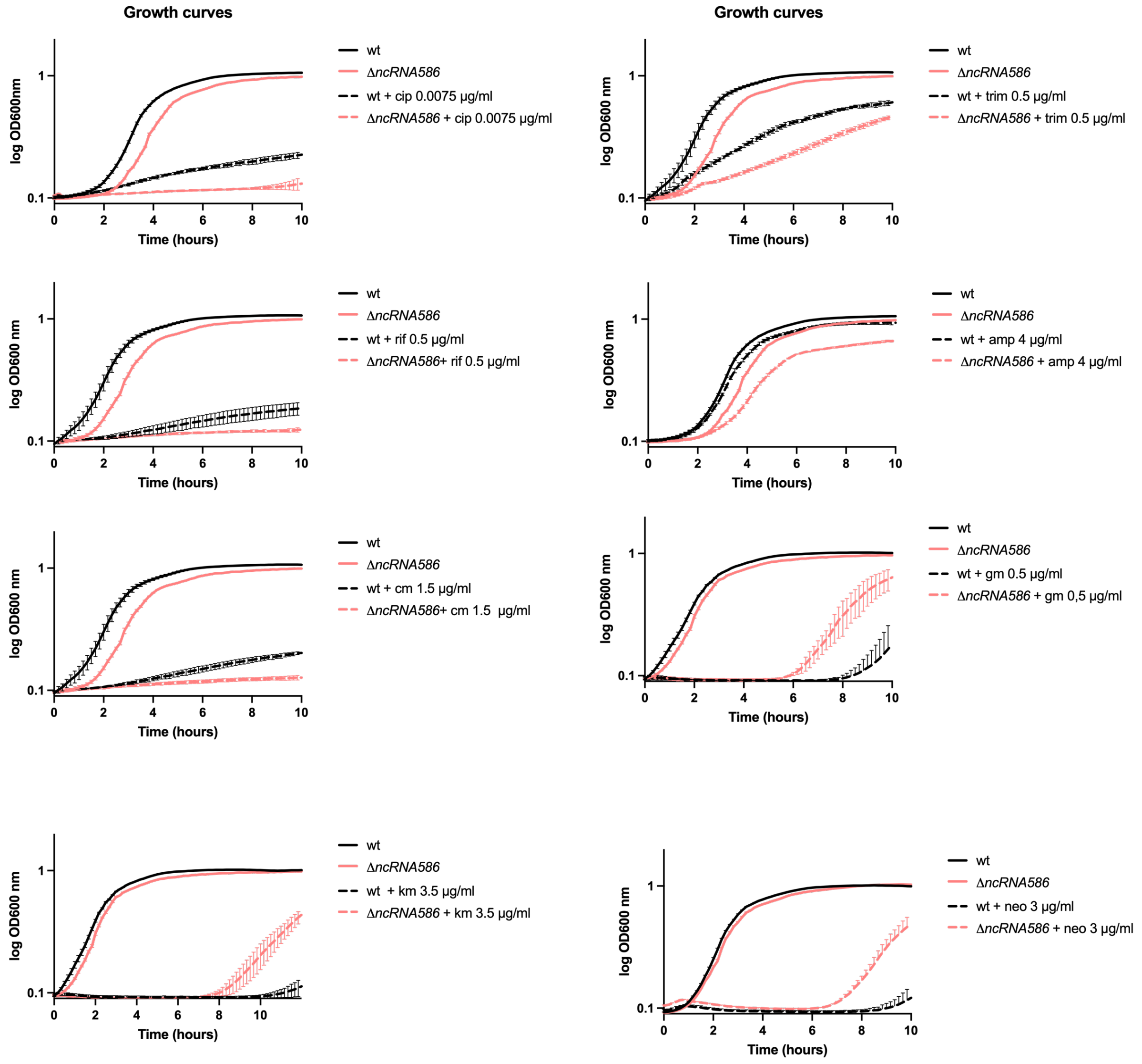

**Figure S1: ncRNA586 effect on antibiotic susceptibility is specific to AGs.** Growth in presence of antibiotics from five different classes (ciprofloxacin, rifampicin, chloramphenicol trimethoprim and ampicillin), or different aminoglycosides (gentamicin, kanamycin and and neomycin 3 µg/ml)**.**

A

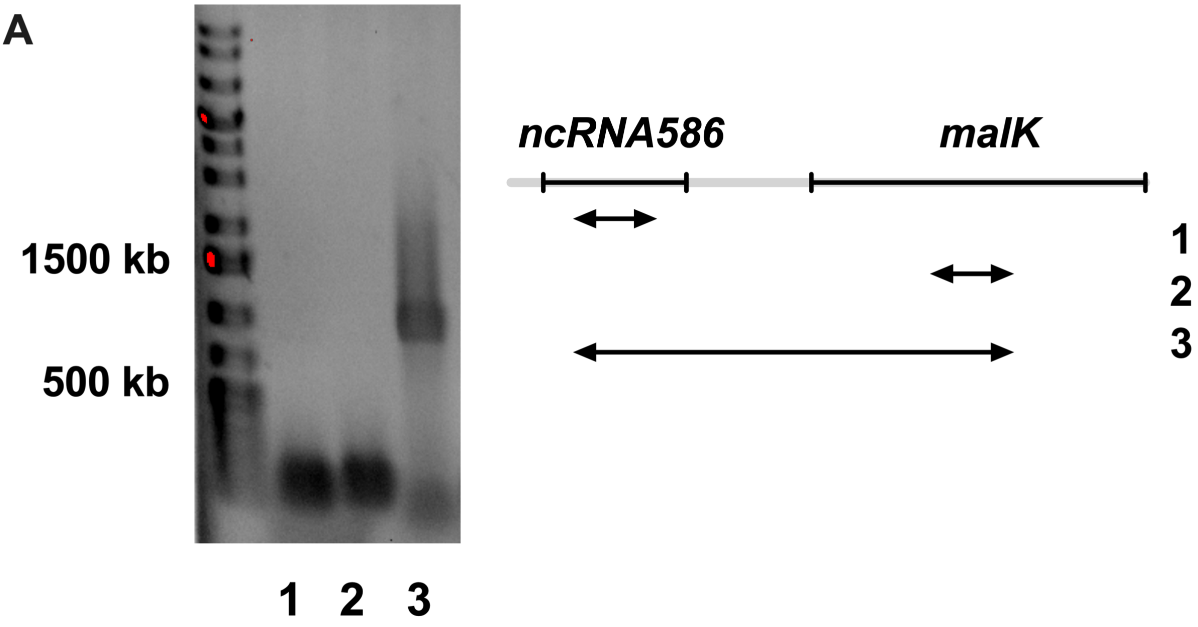

B

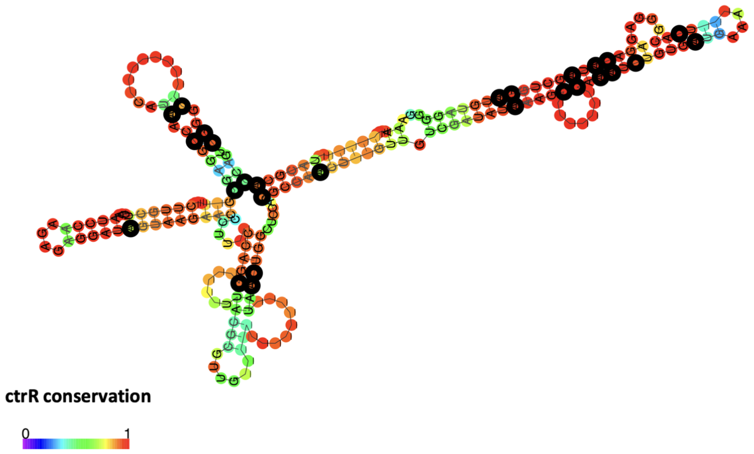

**Figure S2: A.** PCR on total *V. cholerae* RNA (following retrotranscription to obtain cDNA), using primers that bind *ncRNA586* (expected 88 pb amplicon) (1), *malK* (expected 162 pb amplicon) (2) and that bind *ncRNA586* in 5' and *malK* in 3' (expected 968 pb amplicon) (3). **B.** Conserved secondary structure prediction between Vibrio species (*V. cholerae*, *V. mimicus*, *V. parahaemolyticus*, *V. vulnificus* and *V. tasmaniensis*). Base pairing probabilities are shown as a gradient of colors, green being the lowest (0) and red being the highest (1).

A

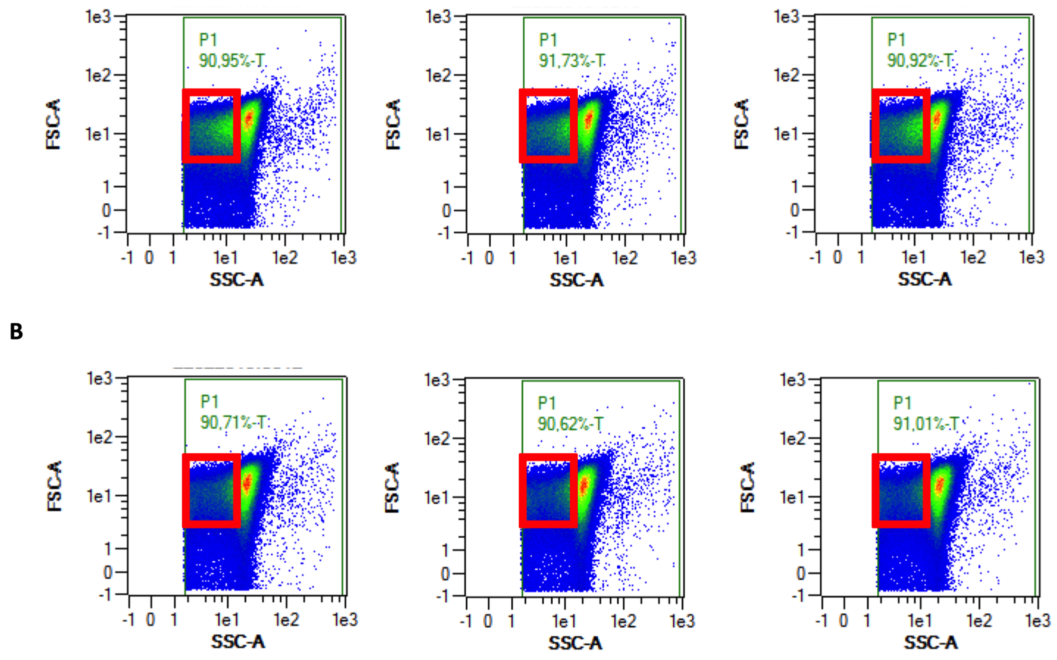

**Figure S3**: Altered cell morphology in the p*ctrR*+ strain (**A**) when compared to a control strain p0 (**B**). FSC (Forward scatter, X-axis) and SCC (Side scatter, Y-axis) parameters through FACS are shown. The populations of interest are highlighted by a red rectangle. Distribution of the p*ctrR*+ population reveals smaller cells that might be damaged by the antibiotic treatment as well as overexpression of the construct.

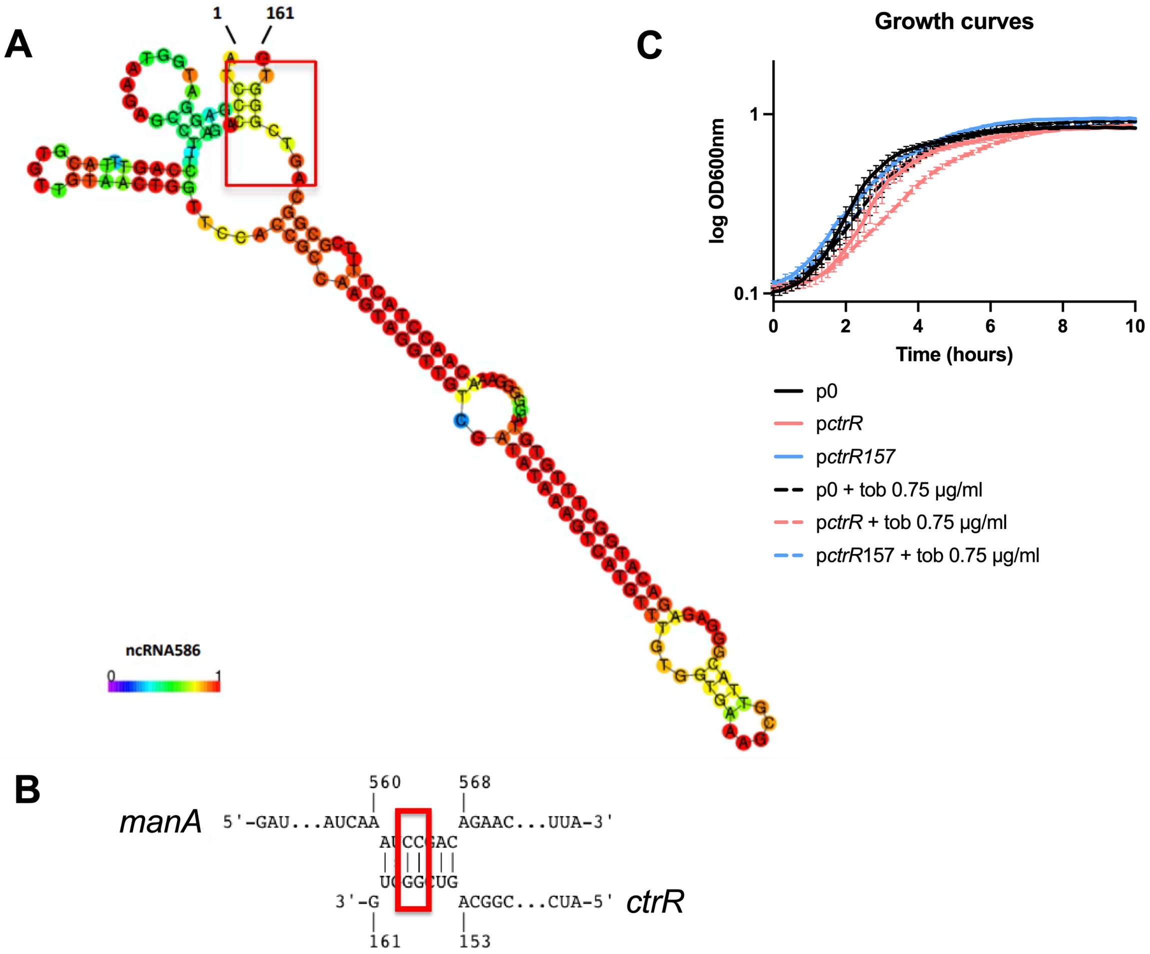

**Figure S4: *ctrR* in predicted to base-paired with *manA* mRNA. A.** Secondary structure of *ncRNA586* as predicted by RNAfold. The red square highlights the region predicted to interact with its putative targets. The first (1) and last (161) bases of ncRNA586 are shown. **B.** Region of interaction between *ctrR* and one of its targets (*manA*). The red square highlights the two nucleotides that were modified (GG to CC for *ctrR*, CC to GG for *manA*) to test the interaction between *ctrR* and *manA*. **C**. Growth curve of *V. cholerae* carrying the empty vector (p0), overexpressing *ctrR* (p*ctrR*+) or overexpressing *ctrR157* (p*ctr157*+), treated or not with tobramycin.

**Supplementary tables:**

**Table S1**: Differential genes expression of *V. cholerae* WT not treated vs upon sub-MIC tobramycin treatment.

| Feature ID | Experiment - Fold Change (original values) | Experiment - Difference (normalized values) | Experiment - Fold Change (normalized values) | Baggerley's test: tob vs nt normalized values - Weighted proportions difference | Baggerley's test: tob vs nt normalized values - Weighted proportions fold change | Baggerley's test: tob vs nt normalized values - Test statistic | Baggerley's test: tob vs nt normalized values - P-value |
|---|---|---|---|---|---|---|---|
| VC1821 | -19,363472 | -228,37867 | -22,503274 | -0,0002229 | -22,503274 | -7,0741151 | 1,5044E-12 |
| tRNA-Arg-6 | -2,6773828 | -28,059447 | -2,820095 | -2,738E-05 | -2,820095 | -6,2936553 | 3,101E-10 |
| eno | -1,7718251 | -920,41821 | -1,8876542 | -0,0008982 | -1,8876542 | -5,9821566 | 2,2021E-09 |
| lamB | -5,5323333 | -7697,9146 | -4,4727473 | -0,0075117 | -4,4727473 | -5,8368821 | 5,3191E-09 |
| tatA | 1,31832691 | 74,2071977 | 1,30224351 | 7,2412E-05 | 1,30224351 | 5,40688872 | 6,4131E-08 |
| ctrR | -5,0071993 | -8922,9306 | -4,8225695 | -0,0087071 | -4,8225695 | -5,3207913 | 1,0332E-07 |
| VC1828 | -17,722882 | -202,11306 | -19,025931 | -0,0001972 | -19,025931 | -5,0081088 | 5,497E-07 |
| VC0411 | -1,5469315 | -106,26349 | -1,5062801 | -0,0001037 | -1,5062801 | -4,9877325 | 6,1094E-07 |
| VCA0392 | 1,36422326 | 49,4908324 | 1,38926508 | 4,8294E-05 | 1,38926508 | 4,91865774 | 8,7142E-07 |
| VCA0514 | 1,4757237 | 45,7738674 | 1,47545574 | 4,4667E-05 | 1,47545574 | 4,80565047 | 1,5425E-06 |
| VC1826 | -41,303681 | -429,72539 | -56,016582 | -0,0004193 | -56,016582 | -4,7415543 | 2,1209E-06 |
| malE | -4,8547625 | -5455,0394 | -4,3579838 | -0,0053231 | -4,3579838 | -4,6018507 | 4,1877E-06 |
| VC2143 | -1,6927887 | -453,00959 | -1,957477 | -0,0004421 | -1,957477 | -4,5375177 | 5,6922E-06 |
| VCA0508 | 1,93829384 | 52,3459875 | 1,96478916 | 5,108E-05 | 1,96478916 | 4,38059815 | 1,1836E-05 |
| VC1055 | 1,79783414 | 54,180986 | 1,76097705 | 5,287E-05 | 1,76097705 | 4,36222403 | 1,2875E-05 |
| VC1920 | 1,75674528 | 28,3058133 | 1,72043221 | 2,7621E-05 | 1,72043221 | 4,0530588 | 5,0553E-05 |
| VC2283 | -2,5331897 | -56,001947 | -2,5186418 | -5,465E-05 | -2,5186418 | -4,0526493 | 5,0642E-05 |
| malQ | -5,6337395 | -424,7325 | -7,0837657 | -0,0004145 | -7,0837657 | -3,9462297 | 7,9393E-05 |
| VC0034 | 1,49359784 | 26,4417742 | 1,48228684 | 2,5802E-05 | 1,48228684 | 3,92597573 | 8,638E-05 |
| VCA1115 | 1,78907795 | 22,4085518 | 1,68125291 | 2,1866E-05 | 1,68125291 | 3,90457856 | 9,4391E-05 |
| VCA0741 | -2,0626387 | -1258,2909 | -2,0014183 | -0,0012279 | -2,0014183 | -3,8945341 | 9,8389E-05 |
| VC1148 | -1,5212746 | -69,229599 | -1,4749997 | -6,755E-05 | -1,4749997 | -3,8238475 | 0,00013139 |
| VC1461 | -1,6044993 | -19,326977 | -1,6688235 | -1,886E-05 | -1,6688235 | -3,8119492 | 0,00013788 |
| VC0410 | -1,5077954 | -166,42139 | -1,6036364 | -0,0001624 | -1,6036364 | -3,7487745 | 0,0001777 |
| recR | 1,62912273 | 39,9368531 | 1,58690879 | 3,8971E-05 | 1,58690879 | 3,72285166 | 0,00019699 |
| rpoB | 1,62965718 | 59,6922162 | 1,63018373 | 5,8248E-05 | 1,63018373 | 3,67865961 | 0,00023447 |
| VC0412 | -1,5530355 | -38,175071 | -1,540191 | -3,725E-05 | -1,540191 | -3,6285265 | 0,00028505 |
| VC0590 | 2,58207946 | 23,3935206 | 2,43163544 | 2,2828E-05 | 2,43163544 | 3,56484738 | 0,00036407 |
| VCA0043 | 1,40967129 | 26,4412204 | 1,39173256 | 2,5802E-05 | 1,39173256 | 3,55408991 | 0,00037929 |
| malG | -3,6932035 | -540,23669 | -4,4272186 | -0,0005272 | -4,4272186 | -3,5307464 | 0,00041439 |
| VC1825 | -4,880914 | -532,53668 | -5,809387 | -0,0005197 | -5,809387 | -3,4733019 | 0,0005141 |
| VC0413 | -1,5414845 | -55,461669 | -1,5256144 | -5,412E-05 | -1,5256144 | -3,4560546 | 0,00054815 |
| VCA1027 | -3,2348871 | -1003,731 | -3,6302852 | -0,0009794 | -3,6302852 | -3,4263311 | 0,0006118 |

| | | | | | | | |
|---|---|---|---|---|---|---|---|
| VC2516 | 1,32035719 | 29,5489571 | 1,30401388 | 2,8834E-05 | 1,30401388 | 3,42026696 | 0,0006256 |
| VCA0821 | 2,21836566 | 46,9006208 | 2,1804536 | 4,5766E-05 | 2,1804536 | 3,39686891 | 0,00068162 |
| VC1822 | -2,6836448 | -61,213993 | -2,9691724 | -5,973E-05 | -2,9691724 | -3,3941259 | 0,00068849 |
| VC2165 | 1,52381205 | 19,5963465 | 1,48657436 | 1,9122E-05 | 1,48657436 | 3,39182225 | 0,0006943 |
| malF | -3,0997594 | -886,67389 | -3,8109529 | -0,0008652 | -3,8109529 | -3,3192973 | 0,00090245 |
| VC2545 | 1,35086212 | 83,3290417 | 1,32454743 | 8,1313E-05 | 1,32454743 | 3,31529084 | 0,00091548 |
| VC1453 | -1,4029312 | -28,022544 | -1,4184456 | -2,734E-05 | -1,4184456 | -3,2730326 | 0,00106401 |
| VCA1030 | -1,2233121 | -36,157522 | -1,2166633 | -3,528E-05 | -1,2166633 | -3,2564343 | 0,00112822 |
| VCA0447 | 2,43715047 | 54,3436203 | 2,38600639 | 5,3029E-05 | 2,38600639 | 3,21946501 | 0,00128431 |
| VC0589 | 2,6095319 | 15,3656161 | 2,38164008 | 1,4994E-05 | 2,38164008 | 3,21327082 | 0,00131233 |
| VC1227 | 2,32873297 | 8,90285015 | 2,18750667 | 8,6875E-06 | 2,18750667 | 3,15442728 | 0,00160814 |
| VCA0845 | 1,33099403 | 23,3934036 | 1,31832948 | 2,2827E-05 | 1,31832948 | 3,10439214 | 0,00190671 |
| VC0124 | 1,3854802 | 24,9307326 | 1,36879618 | 2,4328E-05 | 1,36879618 | 3,09969919 | 0,00193718 |
| pepN | 1,53475792 | 17,4131638 | 1,46606873 | 1,6992E-05 | 1,46606873 | 3,09470715 | 0,00197008 |
| VCA1021 | 2,05679324 | 60,405597 | 2,06105071 | 5,8944E-05 | 2,06105071 | 3,08236827 | 0,00205362 |
| tRNA-Arg-5 | -1,8815859 | -16,858395 | -1,965382 | -1,645E-05 | -1,965382 | -3,0505012 | 0,00228461 |
| malS | -2,7980561 | -46,012991 | -3,3265528 | -4,49E-05 | -3,3265528 | -3,0421981 | 0,00234859 |
| VC2658 | -1,8964391 | -494,57805 | -2,039557 | -0,0004826 | -2,039557 | -3,0366588 | 0,00239218 |
| VC0749 | 1,58192393 | 17,4646201 | 1,52682852 | 1,7042E-05 | 1,52682852 | 3,0361316 | 0,00239636 |
| VCA0185 | 2,35702594 | 8,3602428 | 2,11911688 | 8,158E-06 | 2,11911688 | 2,99981697 | 0,00270143 |
| VCA0719 | -1,6815755 | -10,688001 | -1,8274187 | -1,043E-05 | -1,8274187 | -2,9994125 | 0,00270502 |
| VCA0015 | -3,5568826 | -119,22846 | -3,8992606 | -0,0001163 | -3,8992606 | -2,996435 | 0,00273158 |
| VC1053 | 1,3316118 | 24,692115 | 1,30429226 | 2,4095E-05 | 1,30429226 | 2,99049141 | 0,0027853 |
| VC0566 | 1,51849946 | 47,7747126 | 1,52895546 | 4,6619E-05 | 1,52895546 | 2,97249506 | 0,00295391 |
| VC2659 | -1,9583217 | -580,40422 | -2,0921688 | -0,0005664 | -2,0921688 | -2,9497977 | 0,00317984 |
| clpP | 1,51601393 | 41,0174361 | 1,47508168 | 4,0025E-05 | 1,47508168 | 2,94677727 | 0,00321106 |
| VCA1051 | 1,68965645 | 13,9753111 | 1,61570408 | 1,3637E-05 | 1,61570408 | 2,94158552 | 0,00326538 |
| VC1358 | 1,84614103 | 82,7890871 | 1,85519627 | 8,0786E-05 | 1,85519627 | 2,91072246 | 0,00360597 |
| VC1318 | 1,14151958 | 3119,35546 | 1,48261114 | 0,00304389 | 1,48261114 | 2,90811696 | 0,00363615 |
| VC0856 | 2,35673015 | 20,0990983 | 2,26828199 | 1,9613E-05 | 2,26828199 | 2,9044133 | 0,00367944 |
| VCA0449 | 2,41577406 | 10,4358746 | 2,35381739 | 1,0183E-05 | 2,35381739 | 2,90156947 | 0,003713 |
| VC1039 | 1,51446013 | 22,096447 | 1,47677705 | 2,1562E-05 | 1,47677705 | 2,88698511 | 0,00388954 |
| VC0031 | 1,97041441 | 12,6175898 | 1,84783081 | 1,2312E-05 | 1,84783081 | 2,86840888 | 0,00412544 |
| rbfA | 1,78945058 | 12,6880829 | 1,71961153 | 1,2381E-05 | 1,71961153 | 2,86472627 | 0,00417371 |
| VCA0697 | -1,8501559 | -27,496735 | -2,0148304 | -2,683E-05 | -2,0148304 | -2,8392134 | 0,00452251 |
| VC2515 | 1,20365716 | 38,3174799 | 1,21024709 | 3,7391E-05 | 1,21024709 | 2,8377594 | 0,00454316 |
| VC1827 | -48,042461 | -309,48169 | -60,798769 | -0,000302 | -60,798769 | -2,836712 | 0,0045581 |
| VC1226 | 1,54145305 | 13,9667314 | 1,47413715 | 1,3629E-05 | 1,47413715 | 2,83371474 | 0,00460105 |
| VC0087 | 1,22385381 | 27,8312529 | 1,24545054 | 2,7158E-05 | 1,24545054 | 2,82139723 | 0,00478151 |
| bioD | -1,5782473 | -210,82852 | -1,6351922 | -0,0002057 | -1,6351922 | -2,7794144 | 0,00544572 |
| VCA0013 | -2,7467449 | -433,79188 | -3,5492039 | -0,0004233 | -3,5492039 | -2,7605233 | 0,00577091 |
| VC1329 | -1,8398097 | -10,282259 | -2,025514 | -1,003E-05 | -2,025514 | -2,7553274 | 0,00586336 |
| VCA0899 | -1,8887646 | -98,189142 | -1,8941489 | -9,581E-05 | -1,8941489 | -2,7371457 | 0,00619751 |
| tRNA-Cys-1 | -1,7705194 | -19,779537 | -1,7831641 | -1,93E-05 | -1,7831641 | -2,695712 | 0,00702386 |
| VC1820 | -19,725955 | -281,33844 | -23,716402 | -0,0002745 | -23,716402 | -2,6795462 | 0,00737224 |

| | | | | | | | |
|---|---|---|---|---|---|---|---|
| VC1824 | -3,5808263 | -92,311399 | -3,9446926 | -9,008E-05 | -3,9446926 | -2,6764072 | 0,00744164 |
| VC0467 | 1,52652336 | 15,5941857 | 1,43741487 | 1,5217E-05 | 1,43741487 | 2,67272294 | 0,00752386 |
| VCA0539 | 1,71006983 | 25,1752055 | 1,69871817 | 2,4566E-05 | 1,69871817 | 2,66544789 | 0,00768861 |
| VCA0435 | 1,56517431 | 11,7873616 | 1,50117763 | 1,1502E-05 | 1,50117763 | 2,66194442 | 0,00776909 |
| VCA0180 | -1,6635557 | -69,381914 | -1,6696277 | -6,77E-05 | -1,6696277 | -2,65482 | 0,00793511 |
| rplN | 1,31789716 | 129,807795 | 1,13459172 | 0,00012667 | 1,13459172 | 2,6416223 | 0,00825103 |
| VC2336 | -1,7972188 | -21,329456 | -1,9120579 | -2,081E-05 | -1,9120579 | -2,6145902 | 0,00893348 |
| glgC (NC_002506 637318..638582) | -2,1074099 | -10,469394 | -2,265715 | -1,022E-05 | -2,265715 | -2,6139926 | 0,00894912 |
| VC2657 | -1,8521956 | -639,97119 | -1,8571014 | -0,0006245 | -1,8571014 | -2,6088571 | 0,00908456 |
| tuf (NC_002505 381739..382964) | 1,60930422 | 2818,7452 | 2,04616388 | 0,00275056 | 2,04616388 | 2,5897515 | 0,00960456 |
| VCA0718 | -1,5595379 | -18,596698 | -1,6192086 | -1,815E-05 | -1,6192086 | -2,5726285 | 0,01009297 |
| VCA0303 | -1,5110229 | -9,3410064 | -1,6108945 | -9,115E-06 | -1,6108945 | -2,5606509 | 0,01044765 |
| VC1970 | -1,4502547 | -14,933655 | -1,5120525 | -1,457E-05 | -1,5120525 | -2,5472092 | 0,01085886 |
| VC0018 | 3,30839329 | 372,038406 | 3,18137506 | 0,00036304 | 3,18137506 | 2,5427013 | 0,01099996 |
| VC2279 | 1,3042042 | 17,7209051 | 1,27532986 | 1,7292E-05 | 1,27532986 | 2,53643534 | 0,01119877 |
| VC1971 | -1,8237828 | -37,111673 | -1,8518919 | -3,621E-05 | -1,8518919 | -2,508023 | 0,0121409 |
| VC2662 | 1,72525926 | 130,754243 | 1,71594198 | 0,00012759 | 1,71594198 | 2,50557885 | 0,01222516 |
| VC1414 | 1,69435889 | 33,7642584 | 1,5828845 | 3,2947E-05 | 1,5828845 | 2,48055332 | 0,0131179 |
| VC1592 | -1,82957 | -8,7931449 | -1,990787 | -8,58E-06 | -1,990787 | -2,479248 | 0,013166 |
| VC2490 | 2,7886045 | 12,2674362 | 2,52576421 | 1,1971E-05 | 2,52576421 | 2,4470094 | 0,01440475 |
| VCA0391 | 1,44810362 | 12,7710551 | 1,36939432 | 1,2462E-05 | 1,36939432 | 2,44404582 | 0,01452361 |
| groES (NC_002506 765801..766132) | 5,18920602 | 90,0721523 | 5,06015749 | 8,7893E-05 | 5,06015749 | 2,44004752 | 0,01468538 |
| VCA0513 | 1,44695114 | 27,7143369 | 1,38531364 | 2,7044E-05 | 1,38531364 | 2,43897096 | 0,01472919 |
| VCA0538 | 1,85632519 | 9,95240366 | 1,70089245 | 9,7116E-06 | 1,70089245 | 2,43364417 | 0,01494772 |
| gmhA | 1,23255984 | 19,8154547 | 1,21989293 | 1,9336E-05 | 1,21989293 | 2,42674236 | 0,0152351 |
| VCA0130 | -2,3912329 | -38,833039 | -2,5112807 | -3,789E-05 | -2,5112807 | -2,424006 | 0,01535038 |
| VC0960 | 1,31076842 | 22,0857247 | 1,26528705 | 2,1551E-05 | 1,26528705 | 2,4210555 | 0,01547554 |
| VCA0213 | -1,5070304 | -12,931415 | -1,5825345 | -1,262E-05 | -1,5825345 | -2,3964747 | 0,01655367 |
| VCA0237 | -2,234224 | -23,237529 | -2,1951282 | -2,268E-05 | -2,1951282 | -2,3825086 | 0,01719516 |
| VC1462 | -1,376459 | -10,63497 | -1,4300561 | -1,038E-05 | -1,4300561 | -2,3762368 | 0,01749027 |
| VC2760 | 2,22433445 | 5,55488972 | 2,02487608 | 5,4205E-06 | 2,02487608 | 2,3761955 | 0,01749223 |
| VC0468 | 1,56425682 | 10,0205308 | 1,46029645 | 9,7781E-06 | 1,46029645 | 2,37157551 | 0,01771246 |
| glpK | -5,3429341 | -340,81794 | -7,0292696 | -0,0003326 | -7,0292696 | -2,3653572 | 0,01801274 |
| VC0179 | 1,76839953 | 143,033084 | 1,8591634 | 0,00013957 | 1,8591634 | 2,36002396 | 0,01827379 |
| glyA | 1,86478674 | 48,5913794 | 1,8427964 | 4,7416E-05 | 1,8427964 | 2,33736226 | 0,01942041 |
| hscA | 1,46162671 | 13,3164406 | 1,39704437 | 1,2994E-05 | 1,39704437 | 2,33681354 | 0,01944892 |
| VC2290 | 1,173456 | 38,2151396 | 1,16855783 | 3,7291E-05 | 1,16855783 | 2,33180233 | 0,01971113 |
| groEL (NC_002506 766120..767753) | 3,89334862 | 50,4438616 | 3,86299493 | 4,9224E-05 | 3,86299493 | 2,32908793 | 0,01985446 |
| VC0078 | -1,2147975 | -44,407613 | -1,2169286 | -4,333E-05 | -1,2169286 | -2,3238202 | 0,02013517 |
| VCA0450 | 2,22051234 | 7,29823974 | 2,12048939 | 7,1217E-06 | 2,12048939 | 2,31905077 | 0,02039233 |
| VC1125 | -1,2776362 | -21,640794 | -1,2913363 | -2,112E-05 | -1,2913363 | -2,3145981 | 0,02063497 |
| truB | 1,74600735 | 9,41012216 | 1,66224329 | 9,1825E-06 | 1,66224329 | 2,31383847 | 0,02067662 |
| VC2681 | 1,65421565 | 16,1899375 | 1,57616116 | 1,5798E-05 | 1,57616116 | 2,30399529 | 0,02122295 |
| tRNA-Arg-8 | -1,9230673 | -67,9442 | -1,8930064 | -6,63E-05 | -1,8930064 | -2,3024775 | 0,02130831 |

| | | | | | | | |
|---|---|---|---|---|---|---|---|
| VCA0175 | 1,5433579 | 11,1200335 | 1,4412616 | 1,0851E-05 | 1,4412616 | 2,29236403 | 0,02188468 |
| VC0335 | -1,3282667 | -13,632138 | -1,3811202 | -1,33E-05 | -1,3811202 | -2,2914812 | 0,02193563 |
| VC1174 | -2,0357739 | -4,2194731 | -2,4048198 | -4,117E-06 | -2,4048198 | -2,285358 | 0,02229188 |
| VC0488 | -1,637462 | -6,3754337 | -1,7450903 | -6,221E-06 | -1,7450903 | -2,2784752 | 0,02269832 |
| VC0285 | -2,0685945 | -27,527032 | -2,1991485 | -2,686E-05 | -2,1991485 | -2,2747119 | 0,02292326 |
| aceE | 2,67755842 | 67,5233686 | 2,71353731 | 6,589E-05 | 2,71353731 | 2,25730585 | 0,02398902 |
| VC1300 | -1,1991746 | -22,776556 | -1,181776 | -2,223E-05 | -1,181776 | -2,2553285 | 0,02411275 |
| VC2292 | 1,21308629 | 38,1669118 | 1,25030007 | 3,7244E-05 | 1,25030007 | 2,25523278 | 0,02411877 |
| VC0428 | -1,7373401 | -210,39748 | -1,7387476 | -0,0002053 | -1,7387476 | -2,2524938 | 0,02429113 |
| VCA1114 | 1,63438387 | 8,26506584 | 1,51925952 | 8,0651E-06 | 1,51925952 | 2,25145605 | 0,0243567 |
| VC0329 | 1,6748806 | 112,194429 | 1,70216287 | 0,00010948 | 1,70216287 | 2,25084757 | 0,02439524 |
| VCA0728 | 1,3046898 | 24,4232623 | 1,33285006 | 2,3832E-05 | 1,33285006 | 2,2442333 | 0,02481743 |
| VC0478 | -1,3122824 | -416,69647 | -1,4552789 | -0,0004066 | -1,4552789 | -2,2422671 | 0,02494416 |
| yajC | 1,37005871 | 64,1814323 | 1,36924435 | 6,2629E-05 | 1,36924435 | 2,2422412 | 0,02494585 |
| infB | 1,77203663 | 74,5916721 | 1,77391883 | 7,2787E-05 | 1,77391883 | 2,22900536 | 0,02581362 |
| VC2418 | 1,45846738 | 10,8127491 | 1,35956835 | 1,0551E-05 | 1,35956835 | 2,22336533 | 0,02619121 |
| VC2614 | -1,536415 | -553,06109 | -1,5379807 | -0,0005397 | -1,5379807 | -2,2223207 | 0,0262617 |
| VCA0748 | -7,6847321 | -156,5616 | -9,453909 | -0,0001528 | -9,453909 | -2,2178985 | 0,02656182 |
| hslU | 2,20242698 | 43,264259 | 2,12008374 | 4,2218E-05 | 2,12008374 | 2,21151317 | 0,02700038 |
| VC0302 | -1,6417205 | -9,6581605 | -1,681951 | -9,425E-06 | -1,681951 | -2,2108322 | 0,0270475 |
| 23Sb | 1,10210201 | 3640,34897 | 1,37918471 | 0,00355228 | 1,37918471 | 2,20616525 | 0,02737248 |
| VC0651 | -1,4051662 | -35,766579 | -1,4131447 | -3,49E-05 | -1,4131447 | -2,1972963 | 0,02799934 |
| VC1418 | 1,514894 | 17,0765907 | 1,46277278 | 1,6663E-05 | 1,46277278 | 2,19249175 | 0,02834406 |
| VCA1023 | 1,4333769 | 9,99982363 | 1,38108263 | 9,7579E-06 | 1,38108263 | 2,19121973 | 0,02843593 |
| tRNA-Arg-3 | -1,6469166 | -23,97476 | -1,6682042 | -2,339E-05 | -1,6682042 | -2,1885793 | 0,02862748 |
| VC2037 | -1,7304751 | -169,20692 | -1,9473592 | -0,0001651 | -1,9473592 | -2,1769444 | 0,02948477 |
| ilvH | 1,68391745 | 18,719692 | 1,6656308 | 1,8267E-05 | 1,6656308 | 2,17351333 | 0,02974175 |
| VC1486 | 1,3526151 | 11,7034916 | 1,30798241 | 1,142E-05 | 1,30798241 | 2,16458438 | 0,03041957 |
| VC1425 | 1,58429526 | 83,2516423 | 1,60337514 | 8,1238E-05 | 1,60337514 | 2,16347809 | 0,0305045 |
| VC1178 | 1,45699663 | 12,720047 | 1,36081442 | 1,2412E-05 | 1,36081442 | 2,15496485 | 0,03116463 |
| groEL (NC_002505 2832599..2834274) | 2,65353996 | 503,538714 | 2,50520429 | 0,00049136 | 2,50520429 | 2,14921518 | 0,03161739 |
| clpX | 1,21678111 | 48,9529537 | 1,22857976 | 4,7769E-05 | 1,22857976 | 2,14449788 | 0,03199306 |
| VC1624 | 2,91647046 | 21,2111641 | 2,83002402 | 2,0698E-05 | 2,83002402 | 2,13702586 | 0,03259596 |
| xerD | 1,49137104 | 9,31208791 | 1,38698081 | 9,0868E-06 | 1,38698081 | 2,12819127 | 0,03332127 |
| hisG | 1,94686331 | 9,00225761 | 1,78669677 | 8,7845E-06 | 1,78669677 | 2,12634672 | 0,03347445 |
| VC1849 | 1,59180432 | 128,945133 | 1,50421764 | 0,00012583 | 1,50421764 | 2,11442232 | 0,03447926 |
| VC1623 | 2,66655342 | 25,5133797 | 2,58088621 | 2,4896E-05 | 2,58088621 | 2,1012731 | 0,03561707 |
| VCA0784 | -1,4440538 | -72,888025 | -1,4750385 | -7,112E-05 | -1,4750385 | -2,1011207 | 0,03563043 |
| VCA0451 | 2,33592658 | 3,81660758 | 2,30056711 | 3,7243E-06 | 2,30056711 | 2,09848936 | 0,03586199 |
| VC2741 | 1,27818238 | 12,4339801 | 1,26461408 | 1,2133E-05 | 1,26461408 | 2,08778856 | 0,03681695 |
| VC0753 | 1,34981663 | 11,2421265 | 1,29662482 | 1,097E-05 | 1,29662482 | 2,08714661 | 0,03687493 |
| VCA0174 | 1,4065942 | 10,3612658 | 1,32396095 | 1,0111E-05 | 1,32396095 | 2,08164649 | 0,03737481 |
| VC2458 | 1,52232286 | 28,9296896 | 1,51151992 | 2,823E-05 | 1,51151992 | 2,08001022 | 0,03752466 |
| VC0558 | -1,2246664 | -18,989572 | -1,2122755 | -1,853E-05 | -1,2122755 | -2,0789556 | 0,03762147 |
| tRNA-Ser-2 | 1,19226375 | 20,0664576 | 1,19973156 | 1,9581E-05 | 1,19973156 | 2,07780509 | 0,03772736 |

| | | | | | | | |
|---|---|---|---|---|---|---|---|
| VC2656 | -1,6700879 | -652,04472 | -1,6353826 | -0,0006363 | -1,6353826 | -2,0775774 | 0,03774837 |
| VC1476 | 1,31784786 | 11,7565174 | 1,27758468 | 1,1472E-05 | 1,27758468 | 2,07334142 | 0,03814056 |
| VCA0954 | -1,3582477 | -90,081558 | -1,3876683 | -8,79E-05 | -1,3876683 | -2,0732837 | 0,03814593 |
| VC0244 | 1,5402497 | 391,618748 | 1,43211626 | 0,00038214 | 1,43211626 | 2,07090198 | 0,03836801 |
| VC2699 | -1,8742954 | -196,71138 | -1,9340438 | -0,000192 | -1,9340438 | -2,0631323 | 0,03910013 |
| VC0408 | -1,3784465 | -32,77847 | -1,3512646 | -3,199E-05 | -1,3512646 | -2,0615387 | 0,03925173 |
| VCA0337 | 1,36838683 | 10,7328996 | 1,3024072 | 1,0473E-05 | 1,3024072 | 2,05652014 | 0,03973245 |
| VC1831 | -1,4106654 | -12,342888 | -1,5045512 | -1,204E-05 | -1,5045512 | -2,0533482 | 0,04003886 |
| metG | 1,27550101 | 17,2348174 | 1,25998211 | 1,6818E-05 | 1,25998211 | 2,05310256 | 0,04006268 |
| VCA0463 | 1,27160843 | 13,1385856 | 1,23903449 | 1,2821E-05 | 1,23903449 | 2,05138897 | 0,04022913 |
| VC0556 | 1,57295698 | 15,5692817 | 1,50500582 | 1,5193E-05 | 1,50500582 | 2,0494285 | 0,04042029 |
| groES (NC_002505 2834287..2834618) | 3,32607223 | 701,923371 | 3,26404636 | 0,00068494 | 3,26404636 | 2,04807087 | 0,04055312 |
| VC0787 | 3,1881259 | 2,90100195 | 2,81452116 | 2,8308E-06 | 2,81452116 | 2,03468149 | 0,04188298 |
| VC2001 | -1,8802938 | -72,741735 | -1,8453324 | -7,098E-05 | -1,8453324 | -2,0346702 | 0,04188411 |
| VCA0785 | -1,4403737 | -6,1879477 | -1,5717254 | -6,038E-06 | -1,5717254 | -2,0315078 | 0,04220356 |
| VC2566 | -1,1944352 | -11,09216 | -1,2817417 | -1,082E-05 | -1,2817417 | -2,027079 | 0,04265438 |
| VCA0964 | -1,327913 | -13,377538 | -1,3700049 | -1,305E-05 | -1,3700049 | -2,0205151 | 0,04333003 |
| VCA0623 | 1,86191244 | 74,8001935 | 1,84571623 | 7,2991E-05 | 1,84571623 | 2,01875841 | 0,04351239 |
| VC1117 | 1,92507862 | 21,6619808 | 1,82364182 | 2,1138E-05 | 1,82364182 | 2,01513528 | 0,04389055 |
| VC2047 | -1,3160996 | -11,540565 | -1,3698397 | -1,126E-05 | -1,3698397 | -2,0148417 | 0,04392129 |
| glpC | -11,790012 | -243,36015 | -15,351582 | -0,0002375 | -15,351582 | -2,0128209 | 0,04413356 |
| VC0654 | -1,5934075 | -213,59842 | -1,6358319 | -0,0002084 | -1,6358319 | -2,0117805 | 0,04424313 |
| VC0178 | 1,64638782 | 95,6621012 | 1,72545602 | 9,3348E-05 | 1,72545602 | 2,00906839 | 0,04452995 |
| VC2387 | 1,69349339 | 74,8648562 | 1,72342822 | 7,3054E-05 | 1,72342822 | 2,00791204 | 0,04465268 |
| VC1315 | -1,6749812 | -65,299271 | -1,7992075 | -6,372E-05 | -1,7992075 | -2,0064883 | 0,04480424 |
| rpsL | 1,63297433 | 194,922008 | 1,46309713 | 0,00019021 | 1,46309713 | 2,00149757 | 0,04533887 |
| lpxC | -1,1922332 | -124,75223 | -1,3404949 | -0,0001217 | -1,3404949 | -2,0002437 | 0,04547401 |
| VC2675 | 2,14838862 | 24,4162168 | 2,0417823 | 2,3826E-05 | 2,0417823 | 1,99926206 | 0,04558008 |
| VCA0108 | 1,69551346 | 6,20528421 | 1,54463367 | 6,0552E-06 | 1,54463367 | 1,99610915 | 0,04592209 |
| VC1546 | 2,14872554 | 5,13477867 | 2,00142675 | 5,0106E-06 | 2,00142675 | 1,99560105 | 0,04597742 |
| VCA0486 | 1,61757293 | 68,3450974 | 1,64789915 | 6,6692E-05 | 1,64789915 | 1,99521774 | 0,04601922 |
| VCA1075 | -1,3761006 | -237,847 | -1,5833506 | -0,0002321 | -1,5833506 | -1,987913 | 0,04682139 |
| VC1689 | -1,8140398 | -23,883859 | -2,0076066 | -2,331E-05 | -2,0076066 | -1,9855609 | 0,04708216 |
| VCA0173 | 1,25212292 | 14,8061237 | 1,20130828 | 1,4448E-05 | 1,20130828 | 1,98481566 | 0,04716503 |
| VCA0084 | 1,37887185 | 9,09551024 | 1,33653563 | 8,8755E-06 | 1,33653563 | 1,98248684 | 0,04742482 |
| VC1803 | 1,45067579 | 66,4591451 | 1,46387539 | 6,4851E-05 | 1,46387539 | 1,97544547 | 0,04821768 |
| VC1972 | -1,6591748 | -42,944017 | -1,6633922 | -4,191E-05 | -1,6633922 | -1,9730509 | 0,04848981 |
| VC2291 | 1,2141614 | 44,1457344 | 1,20733757 | 4,3078E-05 | 1,20733757 | 1,97218285 | 0,04858878 |
| VC0165 | 1,48709706 | 35,7383178 | 1,46519175 | 3,4874E-05 | 1,46519175 | 1,96269014 | 0,04968227 |
| VCA0301 | -1,2228854 | -9,8895566 | -1,3034207 | -9,65E-06 | -1,3034207 | -1,9626866 | 0,04968265 |
| glpA | -3,9940265 | -139,00832 | -4,8527036 | -0,0001356 | -4,8527036 | -1,9541925 | 0,05067853 |
| VC0482 | 1,8772836 | 4,75317086 | 1,72889973 | 4,6382E-06 | 1,72889973 | 1,95161643 | 0,05098381 |
| VC0188 | 2,35995384 | 24,8483088 | 2,24173938 | 2,4247E-05 | 2,24173938 | 1,95035264 | 0,05113417 |
| VC2367 | 1,26750736 | 14,715559 | 1,24434518 | 1,436E-05 | 1,24434518 | 1,94982848 | 0,05119661 |
| VC1173 | -2,0691113 | -3,1453837 | -2,3451322 | -3,069E-06 | -2,3451322 | -1,947934 | 0,05142291 |

| | | | | | | | |
|---|---|---|---|---|---|---|---|
| VC2145 | -1,5106643 | -124,06416 | -1,5255962 | -0,0001211 | -1,5255962 | -1,9453925 | 0,05172781 |
| VCA0107 | 1,79565061 | 5,14387957 | 1,64529647 | 5,0194E-06 | 1,64529647 | 1,94021947 | 0,05235307 |
| VC0698 | 1,28034835 | 17,9130521 | 1,24424426 | 1,748E-05 | 1,24424426 | 1,93767908 | 0,05266244 |
| VC0553 | 1,34112065 | 10,1348375 | 1,28045793 | 9,8897E-06 | 1,28045793 | 1,93375172 | 0,05314371 |
| VC0360 | 1,72540562 | 257,186263 | 1,51351539 | 0,00025096 | 1,51351539 | 1,93199474 | 0,05336023 |
| VC1301 | -1,5048981 | -141,04255 | -1,5141467 | -0,0001376 | -1,5141467 | -1,9312414 | 0,05345325 |
| VC0416 | -1,1857872 | -15,135706 | -1,2292481 | -1,477E-05 | -1,2292481 | -1,9308278 | 0,0535044 |
| VC0842 | 2,23450028 | 11,8693125 | 2,18588099 | 1,1582E-05 | 2,18588099 | 1,91558497 | 0,05541802 |
| VCA0763 | 1,54068733 | 6,41972231 | 1,47021273 | 6,2644E-06 | 1,47021273 | 1,91470762 | 0,05552985 |
| VC0487 | -3,9929544 | -67,715885 | -3,7135147 | -6,608E-05 | -3,7135147 | -1,9123223 | 0,05583494 |
| VCA0356 | 1,32057139 | 13,7491355 | 1,27688963 | 1,3417E-05 | 1,27688963 | 1,91002569 | 0,05612995 |
| VCA0488 | 1,31721401 | 39,3423763 | 1,34719437 | 3,8391E-05 | 1,34719437 | 1,90979661 | 0,05615948 |
| VC1435 | -1,4870839 | -15,726886 | -1,5646031 | -1,535E-05 | -1,5646031 | -1,9043228 | 0,05686821 |
| infC | 1,3890364 | 2517,86515 | 1,85122065 | 0,00245695 | 1,85122065 | 1,90418185 | 0,05688659 |
| VC0286 | -2,0813651 | -8,9612313 | -2,4539063 | -8,744E-06 | -2,4539063 | -1,9041316 | 0,0568931 |
| tsf | 1,92363942 | 231,107843 | 1,70058471 | 0,00022552 | 1,70058471 | 1,90234863 | 0,05712567 |
| VC2483 | 1,79125303 | 12,877297 | 1,73552875 | 1,2566E-05 | 1,73552875 | 1,90058046 | 0,05735706 |
| rpmC | 2,84330405 | 1529,77677 | 3,0549319 | 0,00149277 | 3,0549319 | 1,89722282 | 0,05779861 |
| VC1266 | 2,79617742 | 7,30838749 | 2,57793314 | 7,1316E-06 | 2,57793314 | 1,8965473 | 0,05788779 |
| VC1477 | 1,45182907 | 145,292219 | 1,49806218 | 0,00014178 | 1,49806218 | 1,88663102 | 0,05921002 |
| VCA0946 | -2,1462452 | -251,23113 | -2,5583312 | -0,0002452 | -2,5583312 | -1,8830981 | 0,05968716 |
| VCA0098 | 2,1271751 | 5,88956275 | 1,93838401 | 5,7471E-06 | 1,93838401 | 1,88304618 | 0,05969417 |
| VCA0367 | 1,33270847 | 25,0243799 | 1,36084913 | 2,4419E-05 | 1,36084913 | 1,88172175 | 0,05987385 |
| VC1810 | -1,204506 | -15,873555 | -1,2480877 | -1,549E-05 | -1,2480877 | -1,8769529 | 0,0605246 |
| VC0985 | 2,53731319 | 247,824984 | 2,50977684 | 0,00024183 | 2,50977684 | 1,8766581 | 0,06056502 |
| VCA0493 | 1,38869222 | 22,2046767 | 1,37886201 | 2,1668E-05 | 1,37886201 | 1,87615197 | 0,06063447 |
| VC1093 | 1,45285352 | 12,3300044 | 1,36128126 | 1,2032E-05 | 1,36128126 | 1,8753986 | 0,06073796 |
| tgt | 1,70872219 | 21,012172 | 1,64695611 | 2,0504E-05 | 1,64695611 | 1,86754882 | 0,06182507 |
| pdhR | 1,48426221 | 15,99464 | 1,47292932 | 1,5608E-05 | 1,47292932 | 1,86342316 | 0,0624028 |
| rrmJ | 1,42346718 | 25,856638 | 1,39604174 | 2,5231E-05 | 1,39604174 | 1,86152873 | 0,06266962 |
| aceF | 2,49836967 | 71,3985547 | 2,5620976 | 6,9671E-05 | 2,5620976 | 1,85470887 | 0,06363788 |
| VCA0131 | -1,6046885 | -7,718443 | -1,778265 | -7,532E-06 | -1,778265 | -1,8536793 | 0,0637851 |
| VC2251 | 1,45138215 | 110,375941 | 1,41195771 | 0,00010771 | 1,41195771 | 1,84983482 | 0,06433741 |
| VC1593 | -1,8363188 | -36,17006 | -2,0031691 | -3,53E-05 | -2,0031691 | -1,8417676 | 0,06550921 |
| VCA0754a | 1,91017691 | 4,02510548 | 1,83156484 | 3,9277E-06 | 1,83156484 | 1,84091885 | 0,06563351 |
| VC0134 | -1,7179173 | -83,740031 | -1,6502518 | -8,171E-05 | -1,6502518 | -1,8405266 | 0,06569106 |
| VCA0998 | 1,70031784 | 5,63297422 | 1,60628132 | 5,4967E-06 | 1,60628132 | 1,84009218 | 0,06575476 |
| VC2469 | 1,13871758 | 17,3468627 | 1,13873191 | 1,6927E-05 | 1,13873191 | 1,83742472 | 0,06614727 |
| flgE | -1,3034275 | -102,14868 | -1,4066412 | -9,968E-05 | -1,4066412 | -1,8362942 | 0,06631422 |
| VC0274 | -1,3644321 | -59,006939 | -1,3883901 | -5,758E-05 | -1,3883901 | -1,8339282 | 0,0666647 |
| rpsS | 2,47062551 | 1514,81475 | 2,79890532 | 0,00147817 | 2,79890532 | 1,83271189 | 0,0668455 |
| VC1159 | 1,470562 | 25,3722367 | 1,40840159 | 2,4758E-05 | 1,40840159 | 1,83170952 | 0,06699476 |
| VC0407 | -1,2248325 | -21,315785 | -1,1967484 | -2,08E-05 | -1,1967484 | -1,8307678 | 0,06713525 |
| frr | 1,27526418 | 37,8278091 | 1,30791515 | 3,6913E-05 | 1,30791515 | 1,82957485 | 0,06731362 |
| VCA0948 | -1,4329974 | -52,698872 | -1,4548851 | -5,142E-05 | -1,4548851 | -1,8283238 | 0,06750106 |

| | | | | | | | |
|---|---|---|---|---|---|---|---|
| VC0711 | 2,19944883 | 46,7202829 | 2,14614466 | 4,559E-05 | 2,14614466 | 1,82799885 | 0,06754978 |
| VC0748 | 1,36831275 | 14,6559961 | 1,3360398 | 1,4301E-05 | 1,3360398 | 1,82523239 | 0,06796603 |
| glpD | -2,2800455 | -37,962391 | -2,8435934 | -3,704E-05 | -2,8435934 | -1,8241379 | 0,06813127 |
| rplK | 1,73561267 | 1008,68279 | 1,81110305 | 0,00098428 | 1,81110305 | 1,8224396 | 0,06838838 |
| VC0565 | -1,2314942 | -19,187356 | -1,236057 | -1,872E-05 | -1,236057 | -1,8205183 | 0,06868016 |
| VCA1026 | -2,3849787 | -159,34269 | -2,7186592 | -0,0001555 | -2,7186592 | -1,8187253 | 0,06895343 |
| VC1741 | 3,77470963 | 2,23601345 | 3,1107336 | 2,1819E-06 | 3,1107336 | 1,81836231 | 0,06900884 |
| VC0414 | -1,4181059 | -30,568516 | -1,4291089 | -2,983E-05 | -1,4291089 | -1,8169912 | 0,06921852 |
| recO | 1,46333881 | 6,89434998 | 1,37961172 | 6,7276E-06 | 1,37961172 | 1,81647215 | 0,06929803 |
| VC1637 | 1,69675493 | 13,2710278 | 1,60482387 | 1,295E-05 | 1,60482387 | 1,81073394 | 0,07018212 |
| VCA0941 | 1,34293461 | 18,028265 | 1,31844035 | 1,7592E-05 | 1,31844035 | 1,809232 | 0,07041504 |
| VCA0399 | 1,29346802 | 11,7302275 | 1,25450773 | 1,1446E-05 | 1,25450773 | 1,80726 | 0,0707218 |
| rplL | 2,00381906 | 278,332206 | 1,8190106 | 0,0002716 | 1,8190106 | 1,7999748 | 0,07186471 |
| tRNA-Arg-7 | -1,8430161 | -18,545413 | -1,7977562 | -1,81E-05 | -1,7977562 | -1,7979673 | 0,07218227 |
| VCA0047 | -1,2683286 | -19,286046 | -1,3385098 | -1,882E-05 | -1,3385098 | -1,7960503 | 0,07248657 |
| VC0606 | -1,4632146 | -5,6690967 | -1,5684286 | -5,532E-06 | -1,5684286 | -1,7903189 | 0,07340272 |
| VC1545 | 2,14962315 | 3,23373926 | 1,98681173 | 3,1555E-06 | 1,98681173 | 1,79030698 | 0,07340463 |
| secG | -1,3891918 | -515,24984 | -1,4986041 | -0,0005028 | -1,4986041 | -1,7893243 | 0,07356268 |
| VC2571 | 1,34434139 | 157,459246 | 1,17268695 | 0,00015365 | 1,17268695 | 1,78894726 | 0,07362337 |
| VCA0742 | -1,8281099 | -125,88801 | -1,8002231 | -0,0001228 | -1,8002231 | -1,7865052 | 0,07401755 |
| VC0472 | 1,52650037 | 19,1050487 | 1,48599698 | 1,8643E-05 | 1,48599698 | 1,78273995 | 0,07462872 |
| VC0554 | 1,33769223 | 8,90063994 | 1,27006888 | 8,6853E-06 | 1,27006888 | 1,78236661 | 0,07468952 |
| VC0142a | 2,0651703 | 3,13931083 | 2,01783374 | 3,0634E-06 | 2,01783374 | 1,78225422 | 0,07470784 |
| VC0841 | 1,61688969 | 24,6503633 | 1,59755388 | 2,4054E-05 | 1,59755388 | 1,78195379 | 0,07475684 |
| VCA0097 | 1,49157679 | 7,60576474 | 1,42590299 | 7,4218E-06 | 1,42590299 | 1,77950211 | 0,07515753 |
| rplX | 1,35698132 | 251,635902 | 1,22946051 | 0,00024555 | 1,22946051 | 1,77714995 | 0,07554365 |
| glpQ | -3,6906652 | -94,051052 | -4,4505492 | -9,178E-05 | -4,4505492 | -1,770606 | 0,07662633 |
| VC2166 | 1,36574669 | 9,30525088 | 1,29484624 | 9,0801E-06 | 1,29484624 | 1,76408301 | 0,07771806 |
| VC1801 | 2,78021779 | 2,40874556 | 2,50921713 | 2,3505E-06 | 2,50921713 | 1,76289943 | 0,07791751 |
| VCA0795 | 1,78614729 | 4,19640138 | 1,64987138 | 4,0949E-06 | 1,64987138 | 1,75712382 | 0,07889676 |
| VC0067 | 1,55477438 | 5,87714178 | 1,43233187 | 5,735E-06 | 1,43233187 | 1,75452688 | 0,07934032 |
| tolC | 1,20729411 | 33,949059 | 1,22083379 | 3,3128E-05 | 1,22083379 | 1,75424831 | 0,07938804 |
| rplJ | 2,03278155 | 607,404459 | 1,78463948 | 0,00059271 | 1,78463948 | 1,75308806 | 0,07958699 |
| VCA0109 | 1,84025675 | 4,10050967 | 1,66275975 | 4,0013E-06 | 1,66275975 | 1,74982039 | 0,08014936 |
| VC1843 | 1,75840356 | 112,623711 | 1,82128978 | 0,0001099 | 1,82128978 | 1,74965461 | 0,08017803 |
| VC0656 | -1,6527101 | -25,964844 | -1,6539754 | -2,534E-05 | -1,6539754 | -1,7492402 | 0,08024957 |
| VC1715 | 1,33060988 | 8,23127274 | 1,28122269 | 8,0321E-06 | 1,28122269 | 1,74478792 | 0,08102184 |
| VC1010 | 1,29454188 | 29,2964748 | 1,29754799 | 2,8588E-05 | 1,29754799 | 1,74188091 | 0,08152936 |
| VC0243 | 1,4898145 | 365,797196 | 1,42046165 | 0,00035695 | 1,42046165 | 1,74134612 | 0,08162299 |
| fabG | 1,21200278 | 34,3447341 | 1,2489172 | 3,3514E-05 | 1,2489172 | 1,74110025 | 0,08166608 |
| VC0862 | -1,4264505 | -22,644922 | -1,5776183 | -2,21E-05 | -1,5776183 | -1,7405666 | 0,08175964 |
| VC1842 | 1,67397193 | 26,8080706 | 1,64640847 | 2,616E-05 | 1,64640847 | 1,73969101 | 0,08191338 |
| VC0886 | 2,26830362 | 9,84210419 | 2,04210485 | 9,604E-06 | 2,04210485 | 1,7392305 | 0,0819943 |
| VC2035 | -1,5898133 | -13,499554 | -1,5841329 | -1,317E-05 | -1,5841329 | -1,7387079 | 0,08208623 |
| dnaK | 2,30232491 | 273,477229 | 2,24928856 | 0,00026686 | 2,24928856 | 1,73287875 | 0,08311728 |

| | | | | | | | |
|---|---|---|---|---|---|---|---|
| VC1498 | 2,21549559 | 26,0513775 | 2,14553809 | 2,5421E-05 | 2,14553809 | 1,73278166 | 0,08313458 |
| VCA0445 | 1,61993402 | 12,0424681 | 1,53665339 | 1,1751E-05 | 1,53665339 | 1,72841745 | 0,08391348 |
| VC0164 | 1,61262856 | 19,1354602 | 1,5715202 | 1,8673E-05 | 1,5715202 | 1,72772143 | 0,08403826 |
| VCA0324 | 1,25598605 | 17,6671163 | 1,26150415 | 1,724E-05 | 1,26150415 | 1,7276162 | 0,08405712 |
| VCA0858 | -1,4636012 | -21,097994 | -1,6201041 | -2,059E-05 | -1,6201041 | -1,7274712 | 0,08408313 |
| VC0731 | 1,47227215 | 189,474541 | 1,45235335 | 0,00018489 | 1,45235335 | 1,72395354 | 0,0847163 |
| VC0671 | 1,15628474 | 24,0550802 | 1,15659528 | 2,3473E-05 | 1,15659528 | 1,72354428 | 0,08479021 |
| VC0457 | 1,29936462 | 9,12107836 | 1,24303729 | 8,9004E-06 | 1,24303729 | 1,721912 | 0,08508551 |
| VC1264 | 3,12113444 | 19,813138 | 2,95383482 | 1,9334E-05 | 2,95383482 | 1,71773597 | 0,08584487 |
| tRNA-Leu-1 | -1,9047988 | -74,599612 | -1,8218072 | -7,279E-05 | -1,8218072 | -1,7174404 | 0,08589883 |
| VCA0017 | -1,8581022 | -13,865655 | -1,9351459 | -1,353E-05 | -1,9351459 | -1,7122873 | 0,08684377 |
| VCA0924 | 1,60403049 | 5,15720053 | 1,46686808 | 5,0324E-06 | 1,46686808 | 1,71117629 | 0,08704862 |
| VCA0052 | -1,8376276 | -79,717129 | -1,892172 | -7,779E-05 | -1,892172 | -1,7104608 | 0,08718076 |
| pfkA | -2,3064603 | -513,32466 | -2,4664304 | -0,0005009 | -2,4664304 | -1,7089782 | 0,08745506 |
| VCA0079 | -1,1914476 | -25,568288 | -1,1988082 | -2,495E-05 | -1,1988082 | -1,708438 | 0,08755515 |
| VC0982 | 1,13662655 | 33,596112 | 1,18422111 | 3,2783E-05 | 1,18422111 | 1,70252303 | 0,08865741 |
| VC1415 | -1,8829491 | -8,8747339 | -1,9101826 | -8,66E-06 | -1,9101826 | -1,7024053 | 0,08867948 |
| VC0577 | -1,1847054 | -35,906887 | -1,1654095 | -3,504E-05 | -1,1654095 | -1,7016971 | 0,08881223 |
| VC1419 | 1,47095018 | 8,38441559 | 1,39729811 | 8,1816E-06 | 1,39729811 | 1,70162321 | 0,08882608 |
| tig | 1,34595906 | 77,7350185 | 1,33735813 | 7,5855E-05 | 1,33735813 | 1,70150479 | 0,08884833 |
| VC1316 | -1,2844216 | -30,474451 | -1,3243579 | -2,974E-05 | -1,3243579 | -1,6999267 | 0,08914476 |
| VC1265 | 2,70606411 | 7,30430125 | 2,47939559 | 7,1276E-06 | 2,47939559 | 1,6964945 | 0,08979235 |
| VC0863 | -1,9380995 | -67,291651 | -2,1708093 | -6,566E-05 | -2,1708093 | -1,6948519 | 0,0901036 |
| VCA0676 | 5,76295842 | 40,0150855 | 5,72135566 | 3,9047E-05 | 5,72135566 | 1,69250969 | 0,09054893 |
| rplV | 2,69122276 | 718,021596 | 2,51550245 | 0,00070065 | 2,51550245 | 1,69145077 | 0,09075081 |
| VC0180 | 1,39034243 | 58,1338968 | 1,50064314 | 5,6728E-05 | 1,50064314 | 1,69034368 | 0,09096228 |
| VC0657 | -1,4004316 | -37,41631 | -1,3896682 | -3,651E-05 | -1,3896682 | -1,6867612 | 0,09164934 |
| VC1483 | 1,41587544 | 73,2805178 | 1,40411062 | 7,1508E-05 | 1,40411062 | 1,68663515 | 0,09167359 |
| VC1444 | 1,18876732 | 14,9089694 | 1,19087545 | 1,4548E-05 | 1,19087545 | 1,68267179 | 0,09243869 |
| VC1793 | 1,91886923 | 3,44304636 | 1,75436639 | 3,3598E-06 | 1,75436639 | 1,6819528 | 0,09257804 |
| VC0480 | 1,25792608 | 42,0832065 | 1,29531848 | 4,1065E-05 | 1,29531848 | 1,68145279 | 0,09267508 |
| VCA0535 | -1,53435 | -22,329197 | -1,5954252 | -2,179E-05 | -1,5954252 | -1,679521 | 0,0930506 |
| VC0815 | 1,62322698 | 7,94825137 | 1,52321823 | 7,756E-06 | 1,52321823 | 1,6782286 | 0,09330254 |
| VC0854 | 1,96160202 | 130,684633 | 2,01406733 | 0,00012752 | 2,01406733 | 1,67687757 | 0,09356651 |
| hisD | 1,78915409 | 4,82141879 | 1,64492253 | 4,7048E-06 | 1,64492253 | 1,67647578 | 0,09364509 |
| VC2038 | -1,4810048 | -3,9776828 | -1,6130073 | -3,881E-06 | -1,6130073 | -1,6731701 | 0,09429385 |
| VC2205 | -1,3233977 | -31,14768 | -1,3061824 | -3,039E-05 | -1,3061824 | -1,6731307 | 0,09430166 |
| VCA0677 | 6,57408387 | 44,6844261 | 6,58208421 | 4,3603E-05 | 6,58208421 | 1,66659756 | 0,09559455 |
| VC0479 | 1,31924433 | 80,570472 | 1,32255022 | 7,8621E-05 | 1,32255022 | 1,66356122 | 0,09620022 |
| VC0926 | 1,51857485 | 5,04848723 | 1,44706061 | 4,9264E-06 | 1,44706061 | 1,66343227 | 0,09622599 |
| rpsC | 2,76693795 | 502,461166 | 2,7143213 | 0,00049031 | 2,7143213 | 1,66338639 | 0,09623519 |
| VC0750 | 1,48907294 | 25,8863097 | 1,48904024 | 2,526E-05 | 1,48904024 | 1,66325886 | 0,09626072 |
| VC1641 | 2,27692088 | 66,985965 | 2,28662875 | 6,5365E-05 | 2,28662875 | 1,66319354 | 0,09627378 |
| VC0655 | -1,5676125 | -81,210986 | -1,504717 | -7,925E-05 | -1,504717 | -1,6615624 | 0,09660061 |
| VCA0516 | -3,2236812 | -395,90884 | -3,7470241 | -0,0003863 | -3,7470241 | -1,6614751 | 0,09661812 |

| | | | | | | | |
|---|---|---|---|---|---|---|---|
| VCA0622 | 2,54312084 | 4,30341875 | 2,25308973 | 4,1993E-06 | 2,25308973 | 1,66039977 | 0,09683411 |
| VC2137 | -1,1393056 | -14,291496 | -1,1368075 | -1,395E-05 | -1,1368075 | -1,6568971 | 0,09754033 |
| VC1185 | -1,467137 | -6,4304675 | -1,5624739 | -6,275E-06 | -1,5624739 | -1,6558866 | 0,09774483 |
| rpmD | 1,90896985 | 508,849426 | 1,77692656 | 0,00049654 | 1,77692656 | 1,65468537 | 0,09798839 |
| rplB | 2,31678104 | 326,869204 | 2,21476587 | 0,00031896 | 2,21476587 | 1,65395291 | 0,09813714 |
| fliP | -1,248955 | -17,340094 | -1,2658666 | -1,692E-05 | -1,2658666 | -1,6537145 | 0,0981856 |
| VC0834 | 2,071476 | 8,85876909 | 1,96614965 | 8,6445E-06 | 1,96614965 | 1,65358468 | 0,098212 |
| VC0348 | -1,0628172 | -82,406619 | -1,2036053 | -8,041E-05 | -1,2036053 | -1,648157 | 0,09932054 |
| glpT | -3,5862811 | -118,58143 | -4,3654798 | -0,0001157 | -4,3654798 | -1,6457113 | 0,09982329 |
| oppD | 1,39942051 | 10,26266 | 1,32401751 | 1,0014E-05 | 1,32401751 | 1,64308034 | 0,10036636 |
| VC1135 | 1,78460479 | 8,80269152 | 1,72721013 | 8,5897E-06 | 1,72721013 | 1,64167934 | 0,10065655 |
| rpmI | 1,40381269 | 703,421611 | 1,53685536 | 0,0006864 | 1,53685536 | 1,63679615 | 0,10167317 |
| VC1231 | 1,63602327 | 4,57382713 | 1,47928825 | 4,4632E-06 | 1,47928825 | 1,62868831 | 0,10337907 |
| VC1532 | -1,9197553 | -143,08542 | -2,0854931 | -0,0001396 | -2,0854931 | -1,6263242 | 0,10388077 |
| VC1114 | -1,3442198 | -23,527809 | -1,3170181 | -2,296E-05 | -1,3170181 | -1,6253152 | 0,10409548 |
| rplO | 1,85786794 | 442,876356 | 1,68800364 | 0,00043216 | 1,68800364 | 1,62356214 | 0,10446935 |
| VC0319 | -1,1414379 | -12,579094 | -1,149724 | -1,227E-05 | -1,149724 | -1,6212832 | 0,10495697 |
| VCA0333 | 1,37937717 | 83,5627583 | 1,32474176 | 8,1541E-05 | 1,32474176 | 1,61900068 | 0,10544719 |
| leuS | 1,23514152 | 10,8094635 | 1,20858293 | 1,0548E-05 | 1,20858293 | 1,61865197 | 0,10552221 |
| VC1768 | 1,46295629 | 38,4170057 | 1,52715057 | 3,7488E-05 | 1,52715057 | 1,61692625 | 0,10589425 |
| VC0172 | 1,5082717 | 8,46884138 | 1,44876412 | 8,264E-06 | 1,44876412 | 1,61525269 | 0,10625604 |
| VC1313 | -1,2229994 | -6,591756 | -1,3027553 | -6,432E-06 | -1,3027553 | -1,6124587 | 0,10686222 |
| VC1548 | 2,11378223 | 3,95419861 | 1,97710037 | 3,8585E-06 | 1,97710037 | 1,61155214 | 0,1070595 |
| VCA0887 | 1,47427105 | 5,49909933 | 1,37353205 | 5,3661E-06 | 1,37353205 | 1,61130067 | 0,10711425 |
| rplR | 1,73141909 | 313,212939 | 1,53389489 | 0,00030564 | 1,53389489 | 1,61112334 | 0,10715291 |
| VCA0135 | -1,5788874 | -11,678105 | -1,7730338 | -1,14E-05 | -1,7730338 | -1,6074432 | 0,10795727 |
| VC1110 | 1,11638514 | 25,7134345 | 1,15611511 | 2,5091E-05 | 1,15611511 | 1,6061262 | 0,10824625 |
| secY | 1,85756992 | 439,822486 | 1,67965418 | 0,00042918 | 1,67965418 | 1,60543759 | 0,10839761 |
| mtlR | -1,2107506 | -7,6265564 | -1,2747317 | -7,442E-06 | -1,2747317 | -1,6035167 | 0,10882067 |
| VC2517 | 1,29646616 | 18,7666403 | 1,31141775 | 1,8313E-05 | 1,31141775 | 1,60168529 | 0,1092253 |
| VC2167 | 1,31979361 | 9,00697474 | 1,24879377 | 8,7891E-06 | 1,24879377 | 1,60139857 | 0,10928872 |
| flgL | -1,2800045 | -25,754351 | -1,3056322 | -2,513E-05 | -1,3056322 | -1,5997261 | 0,10965943 |
| VC0835 | 2,09945062 | 7,42805295 | 2,00712487 | 7,2484E-06 | 2,00712487 | 1,59750006 | 0,11015437 |
| VC0092 | 1,73190257 | 17,1930995 | 1,61410858 | 1,6777E-05 | 1,61410858 | 1,5939126 | 0,1109557 |
| VC1217 | 2,16685085 | 10,8909739 | 1,98510336 | 1,0628E-05 | 1,98510336 | 1,59047943 | 0,11172686 |
| VC0873 | 1,43257929 | 9,94992955 | 1,38355295 | 9,7092E-06 | 1,38355295 | 1,58987949 | 0,11186203 |
| VCA0319 | 1,56903862 | 6,04708786 | 1,5044852 | 5,9008E-06 | 1,5044852 | 1,58514906 | 0,11293253 |
| VC2033 | -2,3528233 | -1067,3585 | -2,6785239 | -0,0010415 | -2,6785239 | -1,5834707 | 0,11331429 |
| VC0695 | 1,81544702 | 4,1648422 | 1,64307072 | 4,0641E-06 | 1,64307072 | 1,58329351 | 0,11335464 |
| VCA0490 | 1,3354128 | 9,31683379 | 1,27646015 | 9,0914E-06 | 1,27646015 | 1,58265755 | 0,1134996 |
| VCA0487 | 1,40841733 | 23,8672073 | 1,41135043 | 2,329E-05 | 1,41135043 | 1,5817958 | 0,11369628 |
| VCA0397 | 1,20602282 | 14,1640251 | 1,1776966 | 1,3821E-05 | 1,1776966 | 1,57801967 | 0,11456114 |
| VC0191 | -1,3994991 | -4,2932456 | -1,4784945 | -4,189E-06 | -1,4784945 | -1,5768865 | 0,11482171 |
| VC1343 | -1,4205022 | -48,99052 | -1,458701 | -4,781E-05 | -1,458701 | -1,5760985 | 0,1150032 |
| rplP | 2,79635043 | 639,186331 | 2,71124616 | 0,00062372 | 2,71124616 | 1,56949902 | 0,11653178 |

| | | | | | | | |
|---|---|---|---|---|---|---|---|
| dcuC | -1,4951866 | -130,2113 | -1,6224268 | -0,0001271 | -1,6224268 | -1,5672442 | 0,11705768 |
| hscB | 1,39491035 | 5,90967949 | 1,32104166 | 5,7667E-06 | 1,32104166 | 1,56598079 | 0,11735317 |
| VCA0471 | 1,45202658 | 29,1787062 | 1,44474271 | 2,8473E-05 | 1,44474271 | 1,56459556 | 0,11767786 |
| VCA0607 | 1,32554043 | 20,2126909 | 1,30831297 | 1,9724E-05 | 1,30831297 | 1,56205014 | 0,11827625 |
| VC1544 | 1,92313091 | 9,59363133 | 1,82775252 | 9,3615E-06 | 1,82775252 | 1,56192877 | 0,11830484 |
| VCA0457 | 1,52859712 | 4,49751346 | 1,44129933 | 4,3887E-06 | 1,44129933 | 1,5617291 | 0,11835187 |
| VC0486 | -4,5476602 | -96,634176 | -4,2195305 | -9,43E-05 | -4,2195305 | -1,5616243 | 0,11837662 |
| rpsE | 1,76664173 | 178,316367 | 1,69294985 | 0,000174 | 1,69294985 | 1,55858731 | 0,11909416 |
| VC1547 | 2,16397363 | 3,46844636 | 2,07431053 | 3,3845E-06 | 2,07431053 | 1,55248231 | 0,12054693 |
| VCA0752 | 1,96260125 | 36,2780874 | 1,98110713 | 3,54E-05 | 1,98110713 | 1,55098698 | 0,12090485 |
| VC1640 | 1,85554363 | 239,295327 | 1,61584998 | 0,00023351 | 1,61584998 | 1,54796947 | 0,12162974 |
| VCA0049 | -1,2867386 | -11,788668 | -1,391187 | -1,15E-05 | -1,391187 | -1,5468915 | 0,12188948 |
| VC0890 | 1,36528893 | 9,918612 | 1,29920435 | 9,6787E-06 | 1,29920435 | 1,54300368 | 0,12282995 |
| VCA0099 | 1,38420625 | 6,8354541 | 1,29268378 | 6,6701E-06 | 1,29268378 | 1,54132612 | 0,12323749 |
| VC0082 | 1,48161694 | 4,70852257 | 1,40430597 | 4,5946E-06 | 1,40430597 | 1,54122561 | 0,12326194 |
| VCA0189 | 1,28296245 | 8,00446976 | 1,21855585 | 7,8108E-06 | 1,21855585 | 1,5380896 | 0,12402676 |
| VC1753 | -1,1903668 | -6,4735238 | -1,2767218 | -6,317E-06 | -1,2767218 | -1,5363968 | 0,12444114 |
| VCA1042 | 1,2263167 | 16,9106484 | 1,19356145 | 1,6502E-05 | 1,19356145 | 1,53352516 | 0,12514657 |
| VCA0021 | 1,38076562 | 6,11907836 | 1,2922971 | 5,971E-06 | 1,2922971 | 1,52997948 | 0,12602186 |
| VC0409 | -1,301852 | -370,69584 | -1,3421429 | -0,0003617 | -1,3421429 | -1,5288744 | 0,12629568 |
| VC2084 | -2,5994322 | -165,78393 | -2,5204342 | -0,0001618 | -2,5204342 | -1,5283834 | 0,12641746 |
| VCA0236 | -1,9464979 | -126,74371 | -1,7842886 | -0,0001237 | -1,7842886 | -1,5282507 | 0,12645037 |
| VCA0898 | 2,15484316 | 32,6220511 | 2,11143113 | 3,1833E-05 | 2,11143113 | 1,52730986 | 0,12668406 |
| VC2601 | -1,3115392 | -17,48418 | -1,3394325 | -1,706E-05 | -1,3394325 | -1,5270218 | 0,12675567 |
| tRNA-Gln-2 | -1,9691201 | -2,4670803 | -2,1129513 | -2,407E-06 | -2,1129513 | -1,526934 | 0,12677748 |
| VC1460 | -1,3627509 | -4,295594 | -1,4414421 | -4,192E-06 | -1,4414421 | -1,5264708 | 0,1268927 |
| tRNA-Leu-11 | -1,3995691 | -21,29233 | -1,3954886 | -2,078E-05 | -1,3954886 | -1,519618 | 0,12860708 |
| VC1642 | 2,5740921 | 86,8512984 | 2,52402195 | 8,475E-05 | 2,52402195 | 1,51886702 | 0,12879606 |
| VC1368 | 1,22221049 | 12,3643047 | 1,1646097 | 1,2065E-05 | 1,1646097 | 1,51867183 | 0,12884517 |
| tRNA-Asn-2 | -1,4083549 | -194,54574 | -1,6110362 | -0,0001898 | -1,6110362 | -1,5170123 | 0,12926366 |
| VC1767 | 1,39542853 | 13,0515417 | 1,38966181 | 1,2736E-05 | 1,38966181 | 1,51648214 | 0,12939753 |
| VC1134 | 1,82392941 | 5,08429197 | 1,68545059 | 4,9613E-06 | 1,68545059 | 1,51543494 | 0,12966236 |
| VCA0048 | -1,265561 | -23,296135 | -1,3104364 | -2,273E-05 | -1,3104364 | -1,5152525 | 0,12970852 |
| VC0228 | 1,44628814 | 41,9847393 | 1,50578257 | 4,0969E-05 | 1,50578257 | 1,51505386 | 0,12975885 |
| rpsQ | 2,65142058 | 546,383959 | 2,54454289 | 0,00053317 | 2,54454289 | 1,51104124 | 0,13077803 |
| VC2013 | -1,179074 | -265,95476 | -1,4035632 | -0,0002595 | -1,4035632 | -1,5086584 | 0,13138619 |
| VCA0859 | -1,4710977 | -43,236807 | -1,6000433 | -4,219E-05 | -1,6000433 | -1,5076234 | 0,13165099 |
| VC1069 | 1,62958036 | 4,10602996 | 1,54681059 | 4,0067E-06 | 1,54681059 | 1,50006979 | 0,13359638 |
| VCA0656 | -1,7234527 | -33,846733 | -1,9969886 | -3,303E-05 | -1,9969886 | -1,4971796 | 0,13434659 |
| VC0512 | 1,22867375 | 10,5973143 | 1,22724637 | 1,0341E-05 | 1,22724637 | 1,49680209 | 0,13444482 |
| VC0186 | 1,17465111 | 22,112482 | 1,19850679 | 2,1578E-05 | 1,19850679 | 1,4958266 | 0,13469892 |
| tRNA-Phe-2 | -1,7505002 | -51,890155 | -1,7262627 | -5,063E-05 | -1,7262627 | -1,4921534 | 0,13565903 |
| VC1367 | -1,3461459 | -18,041229 | -1,4312981 | -1,76E-05 | -1,4312981 | -1,4903042 | 0,13614435 |
| VC1463 | -1,3384976 | -5,3267371 | -1,497131 | -5,198E-06 | -1,497131 | -1,4900828 | 0,13620252 |
| VC2735 | 1,82250704 | 15,4982271 | 1,73203101 | 1,5123E-05 | 1,73203101 | 1,49000417 | 0,13622323 |

| | | | | | | | |
|---|---|---|---|---|---|---|---|
| VCA0947 | -1,3958309 | -42,857671 | -1,457354 | -4,182E-05 | -1,457354 | -1,489924 | 0,13624431 |
| glnL | -1,236651 | -4,9736079 | -1,3495197 | -4,853E-06 | -1,3495197 | -1,4898629 | 0,13626035 |
| VC1451 | 1,64637248 | 3,53755233 | 1,52893295 | 3,452E-06 | 1,52893295 | 1,48986044 | 0,13626099 |
| def (NC_002505 44278..44828) | 1,1067917 | 14,1051077 | 1,11067102 | 1,3764E-05 | 1,11067102 | 1,48964949 | 0,13631648 |
| VC0485 | -2,2772073 | -799,52347 | -2,6476808 | -0,0007802 | -2,6476808 | -1,4873129 | 0,13693226 |
| VC0810 | 1,34677076 | 6,40000053 | 1,26027927 | 6,2452E-06 | 1,26027927 | 1,48694649 | 0,137029 |
| VC1543 | 1,87453643 | 9,2669664 | 1,7950105 | 9,0428E-06 | 1,7950105 | 1,48361521 | 0,1379111 |
| VC1062 | 1,62618808 | 7,64798621 | 1,49003395 | 7,463E-06 | 1,49003395 | 1,4803959 | 0,13876769 |
| pyrG | -1,2735393 | -109,88359 | -1,3244899 | -0,0001072 | -1,3244899 | -1,4797277 | 0,138946 |
| tpiA | -1,4128176 | -212,95676 | -1,6338184 | -0,0002078 | -1,6338184 | -1,4778637 | 0,13944431 |
| ilvM | 2,62594703 | 16,0188313 | 2,52382839 | 1,5631E-05 | 2,52382839 | 1,47454562 | 0,14033484 |
| VCA0302 | -1,1798292 | -5,991928 | -1,2740898 | -5,847E-06 | -1,2740898 | -1,471946 | 0,14103551 |
| moaD | 1,53228764 | 6,59716409 | 1,48497321 | 6,4376E-06 | 1,48497321 | 1,47067278 | 0,1413797 |
| VC2099 | 1,22840818 | 31,2807609 | 1,26816129 | 3,0524E-05 | 1,26816129 | 1,47051282 | 0,141423 |
| VCA0338 | 1,31526488 | 6,9690712 | 1,23036972 | 6,8005E-06 | 1,23036972 | 1,46953376 | 0,14168812 |
| VC0199 | 1,69382746 | 4,14069573 | 1,57221151 | 4,0405E-06 | 1,57221151 | 1,46791733 | 0,14212674 |
| VC2118 | -1,1901697 | -11,760208 | -1,2098018 | -1,148E-05 | -1,2098018 | -1,4671064 | 0,14234715 |
| fliR | -1,2589516 | -6,963414 | -1,3387184 | -6,795E-06 | -1,3387184 | -1,4668065 | 0,14242874 |
| pgk | -1,6793732 | -433,27498 | -1,9178489 | -0,0004228 | -1,9178489 | -1,4664703 | 0,14252027 |
| rplT | 1,37210376 | 256,912856 | 1,2479167 | 0,0002507 | 1,2479167 | 1,46516721 | 0,14287537 |
| VCA0608 | 1,46200991 | 27,8962352 | 1,44736496 | 2,7221E-05 | 1,44736496 | 1,46271048 | 0,14354671 |
| VC2213 | -1,1262734 | -247,34637 | -1,3657093 | -0,0002414 | -1,3657093 | -1,4617775 | 0,14380227 |
| rpmJ (NC_002505 2754084..2754238) | 1,7151991 | 170,804574 | 1,60531669 | 0,00016667 | 1,60531669 | 1,46083443 | 0,14406095 |
| VC2086 | -2,2666524 | -95,386412 | -2,2064167 | -9,308E-05 | -2,2064167 | -1,4589496 | 0,14457907 |
| VCA0730 | 1,28558279 | 10,4609548 | 1,24139628 | 1,0208E-05 | 1,24139628 | 1,45882086 | 0,14461446 |
| VC2615 | -3,4227639 | -27,280187 | -3,6434169 | -2,662E-05 | -3,6434169 | -1,45263 | 0,1463266 |
| VC2172 | 1,39687064 | 8,99181316 | 1,32116314 | 8,7743E-06 | 1,32116314 | 1,45135047 | 0,14668237 |
| VCA0729 | 1,35257671 | 10,7779444 | 1,28539814 | 1,0517E-05 | 1,28539814 | 1,44877248 | 0,14740119 |
| rpsO | 1,62291949 | 620,582618 | 1,59667138 | 0,00060557 | 1,59667138 | 1,4472373 | 0,14783057 |
| VC1406 | 1,35919076 | 6,40220539 | 1,30189186 | 6,2473E-06 | 1,30189186 | 1,44622656 | 0,14811372 |
| VC0446 | 1,17929664 | 10,7229153 | 1,13903531 | 1,0464E-05 | 1,13903531 | 1,44606626 | 0,14815866 |
| sucC | -2,4305736 | -148,35649 | -2,4037903 | -0,0001448 | -2,4037903 | -1,4455606 | 0,14830057 |
| VC0430 | 1,20302302 | 27,9459243 | 1,19179434 | 2,727E-05 | 1,19179434 | 1,44457473 | 0,14857745 |
| VC1443 | 2,04651683 | 11,2658981 | 1,84207237 | 1,0993E-05 | 1,84207237 | 1,44173411 | 0,14937748 |
| VC0524 | 1,22371575 | 8,87356562 | 1,18832353 | 8,6589E-06 | 1,18832353 | 1,43979147 | 0,14992645 |
| VC0336 | -2,3417585 | -383,81185 | -2,3584209 | -0,0003745 | -2,3584209 | -1,4394049 | 0,15003591 |
| VCA0907 | 2,02149188 | 36,9388016 | 2,03961004 | 3,6045E-05 | 2,03961004 | 1,43849902 | 0,15029259 |
| VC2048 | -1,250143 | -9,769016 | -1,285507 | -9,533E-06 | -1,285507 | -1,4384438 | 0,15030825 |
| VC1478 | -1,5694939 | -34,759567 | -1,5988204 | -3,392E-05 | -1,5988204 | -1,4376265 | 0,15054011 |
| pntB | -1,3653262 | -132,48359 | -1,5022688 | -0,0001293 | -1,5022688 | -1,4372085 | 0,1506588 |
| rplW | 2,12456368 | 257,347576 | 2,03896453 | 0,00025112 | 2,03896453 | 1,43703274 | 0,15070875 |
| VC2654 | 1,23592316 | 18,3923933 | 1,21913349 | 1,7947E-05 | 1,21913349 | 1,43608049 | 0,15097951 |
| VC1479 | 1,18702095 | 24,286227 | 1,19700294 | 2,3699E-05 | 1,19700294 | 1,43500793 | 0,1512849 |
| VC0125 | 1,21642224 | 16,5619545 | 1,19619099 | 1,6161E-05 | 1,19619099 | 1,43493719 | 0,15130506 |
| VCA0300 | -1,2818743 | -18,300174 | -1,3231907 | -1,786E-05 | -1,3231907 | -1,433694 | 0,15165969 |

| | | | | | | | |
|---|---|---|---|---|---|---|---|
| VCA0911 | 3,20100548 | 1,91521797 | 2,88073029 | 1,8689E-06 | 2,88073029 | 1,43252201 | 0,15199455 |
| VC0504 | -1,1978988 | -9,153477 | -1,2105026 | -8,932E-06 | -1,2105026 | -1,4320809 | 0,15212072 |
| VC1182 | 1,34079993 | 15,2569565 | 1,29863768 | 1,4888E-05 | 1,29863768 | 1,43191128 | 0,15216929 |
| rpsD | 1,31645394 | 398,759187 | 1,30949592 | 0,00038911 | 1,30949592 | 1,43035912 | 0,15261406 |
| VCA0921 | -1,2586262 | -26,214771 | -1,3038546 | -2,558E-05 | -1,3038546 | -1,4283184 | 0,15320031 |
| VCA0653 | -1,6285323 | -21,840856 | -1,9085611 | -2,131E-05 | -1,9085611 | -1,4281618 | 0,15324537 |
| VC2293 | 1,19274205 | 45,3054055 | 1,1737808 | 4,4209E-05 | 1,1737808 | 1,42487778 | 0,15419261 |
| VC0013 | -1,1055634 | -12,091227 | -1,1185529 | -1,18E-05 | -1,1185529 | -1,4248044 | 0,15421379 |
| VC0395 | 1,39233048 | 37,7503777 | 1,3692057 | 3,6837E-05 | 1,3692057 | 1,42434473 | 0,1543468 |
| VC1021 | 1,23376989 | 7,67122241 | 1,19291478 | 7,4856E-06 | 1,19291478 | 1,42289909 | 0,15476546 |
| VC1449 | 2,28502561 | 50,8381625 | 2,30609761 | 4,9608E-05 | 2,30609761 | 1,42281215 | 0,15479072 |
| VCA0342 | 1,28177433 | 7,54177375 | 1,22228044 | 7,3593E-06 | 1,22228044 | 1,42188881 | 0,15505857 |
| VC2062 | -1,1350707 | -13,01375 | -1,1312036 | -1,27E-05 | -1,1312036 | -1,4214209 | 0,15519446 |
| VC0334 | -1,1579331 | -6,2173416 | -1,2428256 | -6,067E-06 | -1,2428256 | -1,4210804 | 0,15529345 |
| VCA0094 | -1,4503534 | -3,0943933 | -1,5559464 | -3,02E-06 | -1,5559464 | -1,4209884 | 0,15532016 |
| VC0171 | 1,50414211 | 17,7859793 | 1,47175116 | 1,7356E-05 | 1,47175116 | 1,41925583 | 0,15582451 |
| VCA0433 | 1,506367 | 16,8593391 | 1,4741068 | 1,6451E-05 | 1,4741068 | 1,41713108 | 0,1564447 |
| VC0607 | -1,643158 | -2,34297 | -1,7989026 | -2,286E-06 | -1,7989026 | -1,4164393 | 0,15664699 |
| VC1456 | 1,33644234 | 6,36374205 | 1,28173327 | 6,2098E-06 | 1,28173327 | 1,41638524 | 0,1566628 |
| VC1883 | 1,31321822 | 6,66318029 | 1,25560239 | 6,502E-06 | 1,25560239 | 1,41545318 | 0,15693572 |
| VC2103 | 1,76266123 | 3,93747474 | 1,56923018 | 3,8422E-06 | 1,56923018 | 1,413907 | 0,15738927 |
| VC1099 | 1,62738454 | 12,1298293 | 1,45488139 | 1,1836E-05 | 1,45488139 | 1,41332209 | 0,1575611 |
| VC0786 | 1,87520833 | 2,79778342 | 1,6244781 | 2,7301E-06 | 1,6244781 | 1,41320811 | 0,15759459 |
| VCA1039 | 1,31856289 | 8,2817979 | 1,24754692 | 8,0814E-06 | 1,24754692 | 1,41216973 | 0,15790004 |
| nagB | -1,158386 | -13,256877 | -1,1727168 | -1,294E-05 | -1,1727168 | -1,4120338 | 0,15794007 |
| VC0652 | -1,4080514 | -22,560689 | -1,4316415 | -2,201E-05 | -1,4316415 | -1,4114034 | 0,15812577 |
| VC2565 | -1,3636731 | -21,297114 | -1,4488651 | -2,078E-05 | -1,4488651 | -1,4113468 | 0,15814242 |
| VCA0806 | -1,2226043 | -13,895921 | -1,2690935 | -1,356E-05 | -1,2690935 | -1,4108018 | 0,15830311 |
| VCA0908 | 2,06515605 | 26,8229133 | 2,03587579 | 2,6174E-05 | 2,03587579 | 1,40540532 | 0,15990087 |
| VCA0446 | 2,47463776 | 495,838358 | 2,72477335 | 0,00048384 | 2,72477335 | 1,4051101 | 0,15998864 |
| VC1588 | 1,74995905 | 5,60859568 | 1,66521363 | 5,4729E-06 | 1,66521363 | 1,40497468 | 0,16002888 |
| VCA0056 | 1,53168273 | 3,71839027 | 1,42848042 | 3,6284E-06 | 1,42848042 | 1,4029384 | 0,16063526 |
| VC0458 | 1,25243307 | 7,18592827 | 1,20069092 | 7,0121E-06 | 1,20069092 | 1,4021511 | 0,16087017 |
| VC2221 | -1,9886949 | -1148,7239 | -1,6677315 | -0,0011209 | -1,6677315 | -1,4004436 | 0,16138061 |
| VC0927 | 1,35898381 | 4,67851143 | 1,32408598 | 4,5653E-06 | 1,32408598 | 1,3990142 | 0,16180878 |
| VC2476 | 1,26370084 | 12,7066088 | 1,22194702 | 1,2399E-05 | 1,22194702 | 1,39868062 | 0,16190884 |
| VC2598 | -1,1799948 | -13,825246 | -1,1828089 | -1,349E-05 | -1,1828089 | -1,3972641 | 0,16233424 |
| VC0843 | 1,65689795 | 5,78392145 | 1,582627 | 5,644E-06 | 1,582627 | 1,39480161 | 0,16307575 |
| tRNA-Leu-10 | -1,4820537 | -64,665933 | -1,4512878 | -6,31E-05 | -1,4512878 | -1,3939354 | 0,16333721 |
| VC1964 | 1,32877675 | 20,3557962 | 1,3248462 | 1,9863E-05 | 1,3248462 | 1,39161486 | 0,16403913 |
| VC1001 | 1,19696382 | 10,7575479 | 1,16291622 | 1,0497E-05 | 1,16291622 | 1,39142514 | 0,1640966 |
| VC2061 | -1,2008532 | -17,704509 | -1,2009856 | -1,728E-05 | -1,2009856 | -1,3910252 | 0,16421785 |
| VC0754 | 1,33515313 | 7,57577819 | 1,31609322 | 7,3925E-06 | 1,31609322 | 1,38928251 | 0,16474692 |
| VC0419 | -1,1087054 | -9,8451494 | -1,1413655 | -9,607E-06 | -1,1413655 | -1,3856657 | 0,16584905 |
| VC0666 | 1,37272202 | 11,8225695 | 1,30018591 | 1,1537E-05 | 1,30018591 | 1,38563575 | 0,16585823 |

| | | | | | | | |
|---|---|---|---|---|---|---|---|
| VC1060 | 1,17213508 | 8,76935196 | 1,15739237 | 8,5572E-06 | 1,15739237 | 1,38542958 | 0,16592121 |
| VCA0458 | 1,31636258 | 5,94271029 | 1,25760894 | 5,7989E-06 | 1,25760894 | 1,38541767 | 0,16592484 |
| tRNA-Arg-4 | -1,6491953 | -13,39608 | -1,6481355 | -1,307E-05 | -1,6481355 | -1,3851549 | 0,16600516 |
| VC0422 | -1,1304919 | -8,1399681 | -1,1705192 | -7,943E-06 | -1,1705192 | -1,3851214 | 0,16601541 |
| clpS | 1,36394225 | 43,2716134 | 1,36161039 | 4,2225E-05 | 1,36161039 | 1,38392463 | 0,16638162 |
| VC1678 | 1,40893365 | 10,586383 | 1,35804808 | 1,033E-05 | 1,35804808 | 1,3820594 | 0,16695353 |
| glmU | 1,14196106 | 10,0137615 | 1,13743119 | 9,7715E-06 | 1,13743119 | 1,38100725 | 0,16727678 |
| VCA0088 | 2,3452544 | 17,3529839 | 2,15543377 | 1,6933E-05 | 2,15543377 | 1,38092576 | 0,16730188 |
| VCA0652 | 1,5842812 | 17,6531198 | 1,53559546 | 1,7226E-05 | 1,53559546 | 1,38008747 | 0,16755978 |
| pntA | -1,3407711 | -94,485399 | -1,3913006 | -9,22E-05 | -1,3913006 | -1,3800336 | 0,16757634 |
| VC0356 | -1,5833004 | -146,68681 | -1,5459129 | -0,0001431 | -1,5459129 | -1,3797222 | 0,16767228 |
| VCA0299 | -1,2901242 | -14,493607 | -1,3321169 | -1,414E-05 | -1,3321169 | -1,3773613 | 0,16840065 |
| VCA0897 | 2,03305662 | 20,0639745 | 2,00728918 | 1,9579E-05 | 2,00728918 | 1,37364282 | 0,16955268 |
| VC1791 | 1,47407963 | 4,51787991 | 1,4010111 | 4,4086E-06 | 1,4010111 | 1,37344556 | 0,16961392 |
| VC2627 | 1,29050421 | 10,8705214 | 1,23778496 | 1,0608E-05 | 1,23778496 | 1,37300795 | 0,16974993 |
| VC1864 | 2,04525176 | 5,51853183 | 1,80251791 | 5,385E-06 | 1,80251791 | 1,37290873 | 0,1697808 |
| VC1314 | -1,4629743 | -28,792396 | -1,6114126 | -2,81E-05 | -1,6114126 | -1,3726182 | 0,16987116 |
| VC2386 | 1,50129665 | 78,3753751 | 1,56671757 | 7,6479E-05 | 1,56671757 | 1,37254185 | 0,16989487 |
| VCA0200 | 1,27712821 | 43,8280205 | 1,38265887 | 4,2768E-05 | 1,38265887 | 1,37146106 | 0,17023133 |
| era | 1,31283823 | 12,0086196 | 1,26967311 | 1,1718E-05 | 1,26967311 | 1,3709355 | 0,17039512 |
| VC0557 | 1,32887104 | 44,5442794 | 1,36160392 | 4,3467E-05 | 1,36160392 | 1,36888407 | 0,17103559 |
| serB | 1,47169266 | 9,8651079 | 1,39966335 | 9,6265E-06 | 1,39966335 | 1,36769706 | 0,17140699 |
| VC1859 | 1,84708591 | 6,73419137 | 1,63725539 | 6,5713E-06 | 1,63725539 | 1,36642714 | 0,17180501 |
| VC2154 | 1,09392103 | 14,5642177 | 1,1057393 | 1,4212E-05 | 1,1057393 | 1,36532753 | 0,17215018 |
| glpX | -1,4010457 | -13,134664 | -1,401629 | -1,282E-05 | -1,401629 | -1,3619825 | 0,17320345 |
| VC2420 | 1,25814126 | 13,4503855 | 1,24472644 | 1,3125E-05 | 1,24472644 | 1,36184943 | 0,17324548 |
| VC0885 | 1,61529301 | 5,52729132 | 1,48424122 | 5,3936E-06 | 1,48424122 | 1,36165249 | 0,17330763 |
| VC0850 | 1,3609312 | 8,27846889 | 1,29845715 | 8,0782E-06 | 1,29845715 | 1,36023315 | 0,17375622 |
| VCA1031 | -1,4935664 | -18,83805 | -1,4687316 | -1,838E-05 | -1,4687316 | -1,3589812 | 0,17415261 |
| VC1141 | 1,33254401 | 36,1303565 | 1,29250786 | 3,5256E-05 | 1,29250786 | 1,35841829 | 0,17433109 |
| fliS | -1,3012155 | -33,128664 | -1,2886921 | -3,233E-05 | -1,2886921 | -1,3581225 | 0,17442489 |
| VCA0901 | -1,3147888 | -3,6006869 | -1,4095835 | -3,514E-06 | -1,4095835 | -1,3550516 | 0,17540115 |
| VC0812 | 1,41445617 | 11,1640964 | 1,34053855 | 1,0894E-05 | 1,34053855 | 1,35459239 | 0,17554751 |
| VC1087 | -1,2129492 | -10,355903 | -1,2848418 | -1,011E-05 | -1,2848418 | -1,3539741 | 0,1757447 |
| VCA1097 | -1,6933753 | -3,4498312 | -1,7361341 | -3,366E-06 | -1,7361341 | -1,352863 | 0,17609945 |
| VCA0489 | 1,20972729 | 18,4107775 | 1,21904356 | 1,7965E-05 | 1,21904356 | 1,35269093 | 0,17615446 |
| VC0245 | 1,50306061 | 242,215837 | 1,41454461 | 0,00023636 | 1,41454461 | 1,3526312 | 0,17617353 |
| VC1124 | -1,1969623 | -5,2129867 | -1,264467 | -5,087E-06 | -1,264467 | -1,3514858 | 0,17653992 |
| tRNA-Ser-5 | -1,1791155 | -22,685952 | -1,1626479 | -2,214E-05 | -1,1626479 | -1,3513679 | 0,17657768 |
| VC0156 | 1,76052029 | 55,338025 | 1,82829612 | 5,3999E-05 | 1,82829612 | 1,34899228 | 0,17733952 |
| VC0181 | 1,31687622 | 32,353482 | 1,3749016 | 3,1571E-05 | 1,3749016 | 1,348192 | 0,17759669 |
| tRNA-Ser-4 | 1,15172595 | 10,133213 | 1,14725729 | 9,8881E-06 | 1,14725729 | 1,34791553 | 0,1776856 |
| VCA0314 | 1,21809895 | 12,5615573 | 1,20529193 | 1,2258E-05 | 1,20529193 | 1,34393046 | 0,17897093 |
| flgH | -1,2725401 | -19,603158 | -1,3339505 | -1,913E-05 | -1,3339505 | -1,342605 | 0,17939996 |
| VC0832 | 2,08827601 | 6,80762397 | 1,96527528 | 6,6429E-06 | 1,96527528 | 1,34234229 | 0,17948509 |

| | | | | | | | |
|---|---|---|---|---|---|---|---|
| VCA0910 | 2,81255497 | 1,63501855 | 2,42858232 | 1,5955E-06 | 2,42858232 | 1,34187704 | 0,17963593 |
| VC0105 | 1,32613097 | 13,8695572 | 1,2533457 | 1,3534E-05 | 1,2533457 | 1,34175048 | 0,17967698 |
| VCA0329a | 1,33214513 | 5,51213488 | 1,24398739 | 5,3788E-06 | 1,24398739 | 1,34090949 | 0,17994989 |
| VC2058 | -1,2601753 | -46,939521 | -1,2366044 | -4,58E-05 | -1,2366044 | -1,3374085 | 0,18108943 |
| VC0054 | 1,36015524 | 5,78666726 | 1,29473039 | 5,6467E-06 | 1,29473039 | 1,33685791 | 0,18126907 |
| VCA1079 | 1,21707207 | 23,3670996 | 1,22777326 | 2,2802E-05 | 1,22777326 | 1,33651999 | 0,18137944 |
| VC1838 | 1,11987632 | 12,0317185 | 1,13703036 | 1,1741E-05 | 1,13703036 | 1,33643537 | 0,18140707 |
| rbsC | -1,4512366 | -4,5843185 | -1,6905382 | -4,473E-06 | -1,6905382 | -1,3352597 | 0,18179142 |
| VC0833 | 1,89445816 | 11,2823519 | 1,86365583 | 1,1009E-05 | 1,86365583 | 1,3343705 | 0,18208253 |
| VC0166 | 1,49052365 | 20,5479644 | 1,49155179 | 2,0051E-05 | 1,49155179 | 1,33407831 | 0,18217828 |
| tRNA-Ala-5 | -1,5949838 | -2,4925935 | -1,6227375 | -2,432E-06 | -1,6227375 | -1,3324859 | 0,18270061 |
| VC0039 | 1,27147856 | 6,09036266 | 1,21487096 | 5,943E-06 | 1,21487096 | 1,33138471 | 0,18306252 |
| VC1823 | -1,688044 | -14,473223 | -2,0167296 | -1,412E-05 | -2,0167296 | -1,3308121 | 0,1832509 |
| nrdB | 1,74432034 | 45,0371619 | 1,74748394 | 4,3948E-05 | 1,74748394 | 1,3293928 | 0,18371849 |
| ruvB | 1,35758559 | 5,47969234 | 1,30095934 | 5,3471E-06 | 1,30095934 | 1,32934189 | 0,18373526 |
| metQ | 1,92086279 | 66,536508 | 1,99394888 | 6,4927E-05 | 1,99394888 | 1,32918992 | 0,1837854 |
| VC1100 | 1,43824456 | 5,74640452 | 1,30989175 | 5,6074E-06 | 1,30989175 | 1,32650968 | 0,18467098 |
| VC2206 | -1,2236061 | -63,723012 | -1,2776712 | -6,218E-05 | -1,2776712 | -1,3261825 | 0,18477933 |
| VC1874 | -1,3966028 | -3,0131632 | -1,4826162 | -2,94E-06 | -1,4826162 | -1,3256223 | 0,18496488 |
| VCA0592 | -1,1524875 | -5,6433008 | -1,2299192 | -5,507E-06 | -1,2299192 | -1,3212232 | 0,18642701 |
| dnaA | -1,1662248 | -17,430107 | -1,1828059 | -1,701E-05 | -1,1828059 | -1,3189462 | 0,18718718 |
| VCA0675 | 2,12234337 | 20,7077343 | 2,01722397 | 2,0207E-05 | 2,01722397 | 1,31838893 | 0,18737357 |
| VC0688 | 1,28463247 | 6,6840051 | 1,19295266 | 6,5223E-06 | 1,19295266 | 1,31648896 | 0,18801003 |
| VC0776 | 2,95395435 | 1,25582203 | 2,69408732 | 1,2254E-06 | 2,69408732 | 1,3144357 | 0,18869967 |
| VCA0334 | 1,4195225 | 10,3814023 | 1,35566218 | 1,013E-05 | 1,35566218 | 1,31331116 | 0,18907819 |
| VCA0190 | 1,22293107 | 7,37221242 | 1,16887574 | 7,1939E-06 | 1,16887574 | 1,31230848 | 0,18941611 |
| VC0137 | -1,2031306 | -12,445877 | -1,2744539 | -1,214E-05 | -1,2744539 | -1,3118782 | 0,1895613 |
| VC1688 | -1,5982437 | -2,0388509 | -1,7828603 | -1,99E-06 | -1,7828603 | -1,3117486 | 0,18960502 |
| VC1450 | 2,1188158 | 20,3702423 | 2,12290499 | 1,9877E-05 | 2,12290499 | 1,31121218 | 0,18978618 |
| VC1863 | 1,7673599 | 7,10058235 | 1,59545298 | 6,9288E-06 | 1,59545298 | 1,30907006 | 0,1905107 |
| VC0058 | 1,21575032 | 8,46023747 | 1,17284866 | 8,2556E-06 | 1,17284866 | 1,30849827 | 0,19070441 |
| VC0876 | 1,37433749 | 4,69512526 | 1,27670154 | 4,5815E-06 | 1,27670154 | 1,30840173 | 0,19073714 |
| rpsA | 1,71848319 | 1137,53052 | 1,99114468 | 0,00111001 | 1,99114468 | 1,30768156 | 0,19098143 |
| VCA0230 | 2,48582486 | 4,53503486 | 2,2818 | 4,4253E-06 | 2,2818 | 1,307124 | 0,19117068 |
| murP | 3,16729511 | 8,71970395 | 2,63575437 | 8,5088E-06 | 2,63575437 | 1,30709549 | 0,19118037 |
| VC0121 | 1,50572905 | 10,1268344 | 1,44522056 | 9,8819E-06 | 1,44522056 | 1,30616108 | 0,19149784 |
| VCA0082 | 1,67110908 | 2,73056698 | 1,52528208 | 2,6645E-06 | 1,52528208 | 1,30535862 | 0,19177081 |
| aspS | 1,20988488 | 11,154507 | 1,18875515 | 1,0885E-05 | 1,18875515 | 1,30258691 | 0,19271588 |
| thrA | 1,91412119 | 13,0148174 | 1,84498985 | 1,27E-05 | 1,84498985 | 1,30151526 | 0,19308223 |
| VCA0477 | 1,35424029 | 14,5409747 | 1,32737252 | 1,4189E-05 | 1,32737252 | 1,30009546 | 0,19356833 |
| VC1802 | 1,93787321 | 2,12440969 | 1,72011903 | 2,073E-06 | 1,72011903 | 1,29893932 | 0,1939648 |
| VC1044 | 1,37436193 | 6,02776095 | 1,26443271 | 5,8819E-06 | 1,26443271 | 1,29801821 | 0,19428113 |
| VC0026 | 1,18791745 | 7,91386293 | 1,1526262 | 7,7224E-06 | 1,1526262 | 1,29746908 | 0,19446988 |
| VC1740 | 2,19820433 | 3,28519489 | 1,94542141 | 3,2057E-06 | 1,94542141 | 1,29745373 | 0,19447518 |
| VCA0637 | -1,3646818 | -14,059644 | -1,4090112 | -1,372E-05 | -1,4090112 | -1,2964584 | 0,19481769 |

| | | | | | | | |
|---|---|---|---|---|---|---|---|
| rplA | 1,69246692 | 355,976693 | 1,53717968 | 0,00034737 | 1,53717968 | 1,2944926 | 0,19549539 |
| VC0320 | -1,1337272 | -7,1870648 | -1,1684559 | -7,013E-06 | -1,1684559 | -1,2942378 | 0,19558334 |
| VC1045 | 1,40993341 | 9,82719731 | 1,31838162 | 9,5895E-06 | 1,31838162 | 1,29140205 | 0,19656436 |
| VC1625 | 2,91569446 | 8,73185351 | 2,80536499 | 8,5206E-06 | 2,80536499 | 1,29013023 | 0,19700553 |
| VCA0883 | 1,36095584 | 4,23605906 | 1,30131759 | 4,1336E-06 | 1,30131759 | 1,28993872 | 0,19707198 |
| VCA0627 | 1,71127588 | 2,52740556 | 1,5604089 | 2,4663E-06 | 1,5604089 | 1,28824132 | 0,19766202 |
| VC0247 | 1,44469834 | 230,460233 | 1,36288169 | 0,00022489 | 1,36288169 | 1,28777475 | 0,19782446 |
| VC0608 | 2,86508318 | 24,2987229 | 2,81734477 | 2,3711E-05 | 2,81734477 | 1,28730322 | 0,19798871 |
| VCA0732 | 1,84601984 | 29,2431296 | 1,82986965 | 2,8536E-05 | 1,82986965 | 1,28708217 | 0,19806574 |
| VCA0581 | 1,32856486 | 4,96135661 | 1,25044447 | 4,8413E-06 | 1,25044447 | 1,28702565 | 0,1980854 |
| VC0891 | 1,49289418 | 7,35042553 | 1,39304349 | 7,1726E-06 | 1,39304349 | 1,28643028 | 0,198293 |
| VC2492 | 1,36819345 | 6,59277129 | 1,31127861 | 6,4333E-06 | 1,31127861 | 1,2854176 | 0,19864645 |
| VCA0113 | 1,61395087 | 3,57282569 | 1,46761062 | 3,4864E-06 | 1,46761062 | 1,28403715 | 0,19912901 |
| uvrB | 1,34285864 | 4,70343761 | 1,26429473 | 4,5897E-06 | 1,26429473 | 1,28336568 | 0,19936404 |
| VC1454 | -1,288449 | -7,4235155 | -1,3303791 | -7,244E-06 | -1,3303791 | -1,2824817 | 0,19967378 |
| VC0357 | -1,6150321 | -55,27196 | -1,5347069 | -5,393E-05 | -1,5347069 | -1,2817864 | 0,19991768 |
| VC0459 | 1,24228312 | 5,88033211 | 1,20503663 | 5,7381E-06 | 1,20503663 | 1,28078472 | 0,20026934 |
| VC0869 | 1,59968209 | 4,56698819 | 1,53835289 | 4,4565E-06 | 1,53835289 | 1,28018123 | 0,20048147 |
| VC0836 | 1,76755598 | 5,66706358 | 1,707234 | 5,53E-06 | 1,707234 | 1,27825193 | 0,20116068 |
| VC1899 | 1,48978678 | 6,12100598 | 1,37678084 | 5,9729E-06 | 1,37678084 | 1,27817875 | 0,20118647 |
| VC2098 | 1,25645013 | 20,6749091 | 1,26225455 | 2,0175E-05 | 1,26225455 | 1,27806438 | 0,2012268 |
| VCA0678 | 3,09801392 | 58,8322026 | 3,10873947 | 5,7409E-05 | 3,10873947 | 1,27804706 | 0,20123291 |
| VC0029 | 1,96154107 | 10,0819148 | 1,83827809 | 9,838E-06 | 1,83827809 | 1,27686648 | 0,20164947 |
| VC1912 | -1,1712541 | -20,38592 | -1,1598139 | -1,989E-05 | -1,1598139 | -1,2758549 | 0,2020069 |
| VCA0482 | 1,41785243 | 4,84245528 | 1,37845704 | 4,7253E-06 | 1,37845704 | 1,26929932 | 0,20433438 |
| acpS | 1,29067209 | 29,1756535 | 1,29133806 | 2,847E-05 | 1,29133806 | 1,26904814 | 0,20442398 |
| sspA | -1,1786618 | -19,498816 | -1,145103 | -1,903E-05 | -1,145103 | -1,266627 | 0,20528875 |
| VCA0938 | -1,3345429 | -5,1500786 | -1,3723901 | -5,025E-06 | -1,3723901 | -1,2659682 | 0,2055245 |
| VC2266 | -1,267562 | -6,5657877 | -1,2991832 | -6,407E-06 | -1,2991832 | -1,2656898 | 0,20562419 |
| VCr025 | -1,1490794 | -7758,7819 | -1,3940015 | -0,0075711 | -1,3940015 | -1,2645896 | 0,20601858 |
| VC2297 | 1,50465089 | 12,0226237 | 1,3756495 | 1,1732E-05 | 1,3756495 | 1,26277847 | 0,20666888 |
| VCA0610 | -1,3110612 | -27,450027 | -1,4237151 | -2,679E-05 | -1,4237151 | -1,2626689 | 0,20670829 |
| VC2124 | -1,1830725 | -15,710766 | -1,2117059 | -1,533E-05 | -1,2117059 | -1,2624882 | 0,20677324 |
| VC1095 | 1,30050414 | 6,43607624 | 1,22756627 | 6,2804E-06 | 1,22756627 | 1,26231774 | 0,20683452 |
| VC1195 | 1,73387767 | 29,7609966 | 1,73028888 | 2,9041E-05 | 1,73028888 | 1,25900872 | 0,20802727 |
| VCA1029 | -1,4459742 | -33,998173 | -1,5787167 | -3,318E-05 | -1,5787167 | -1,2583582 | 0,20826234 |
| VC2282 | -1,2777403 | -3,5949093 | -1,3438783 | -3,508E-06 | -1,3438783 | -1,2578884 | 0,20843216 |
| VC1578 | 1,60904423 | 499,833669 | 1,505837 | 0,00048774 | 1,505837 | 1,25667477 | 0,2088715 |
| VC1137 | 1,73219867 | 9,76468079 | 1,71019272 | 9,5285E-06 | 1,71019272 | 1,2565494 | 0,20891693 |
| VCA0913 | 2,29076888 | 2,1869483 | 2,15272211 | 2,134E-06 | 2,15272211 | 1,25602123 | 0,20910834 |
| VC2491 | 1,45298257 | 4,93283828 | 1,34153863 | 4,8135E-06 | 1,34153863 | 1,25520359 | 0,20940492 |
| VC0417 | -1,1084986 | -7,6975974 | -1,146084 | -7,511E-06 | -1,146084 | -1,2538001 | 0,20991473 |
| VC1070 | 1,51925715 | 3,26685758 | 1,43748995 | 3,1878E-06 | 1,43748995 | 1,25352242 | 0,2100157 |
| VC0373 | -1,2078885 | -7,533946 | -1,2524723 | -7,352E-06 | -1,2524723 | -1,2511408 | 0,21088315 |
| VC2555 | 1,28852403 | 7,58882441 | 1,20648557 | 7,4052E-06 | 1,20648557 | 1,25003281 | 0,21128761 |

| | | | | | | | |
|---|---|---|---|---|---|---|---|
| VC1373 | 1,60014681 | 2,72572518 | 1,47195157 | 2,6598E-06 | 1,47195157 | 1,24948765 | 0,21148683 |
| truD | 1,31799169 | 4,96572719 | 1,25836196 | 4,8456E-06 | 1,25836196 | 1,24900736 | 0,21166243 |
| VCA0814 | 1,27855469 | 15,4345179 | 1,25608928 | 1,5061E-05 | 1,25608928 | 1,24809837 | 0,21199509 |
| VC0025 | 1,28792159 | 8,79555356 | 1,21644925 | 8,5828E-06 | 1,21644925 | 1,24797177 | 0,21204145 |
| VC0469 | 1,43014857 | 3,81847977 | 1,31451613 | 3,7261E-06 | 1,31451613 | 1,24767153 | 0,21215142 |
| VC1254 | 1,59408699 | 29,9243637 | 1,57983579 | 2,92E-05 | 1,57983579 | 1,24646619 | 0,21259336 |
| hisH | 1,70173956 | 13,992467 | 1,67797811 | 1,3654E-05 | 1,67797811 | 1,24598496 | 0,21277 |
| VC1985 | 1,51582375 | 13,7907522 | 1,39231842 | 1,3457E-05 | 1,39231842 | 1,2451731 | 0,2130682 |
| hemH (NC_002505 1264415..1265562) | -1,383013 | -22,17521 | -1,4737439 | -2,164E-05 | -1,4737439 | -1,2447 | 0,21324208 |
| VC0055 | 1,37812794 | 4,2174981 | 1,29059977 | 4,1155E-06 | 1,29059977 | 1,24170423 | 0,21434575 |
| VC1758 | 1,54991836 | 7,43941624 | 1,44318179 | 7,2594E-06 | 1,44318179 | 1,24112518 | 0,21455958 |
| rplF | 1,59896872 | 357,282165 | 1,42454612 | 0,00034864 | 1,42454612 | 1,24081861 | 0,21467285 |
| VC1330 | -1,2936301 | -2,795528 | -1,4469411 | -2,728E-06 | -1,4469411 | -1,2376778 | 0,2158356 |
| VCA1024 | -1,1717379 | -5,0157343 | -1,2745556 | -4,894E-06 | -1,2745556 | -1,2371229 | 0,21604151 |
| VC2454 | -1,17711 | -4,2083973 | -1,2756666 | -4,107E-06 | -1,2756666 | -1,236689 | 0,21620261 |
| VCA1113 | 1,63708406 | 7,72715329 | 1,58156366 | 7,5402E-06 | 1,58156366 | 1,23605545 | 0,21643802 |
| rpmE2 | 1,40459126 | 4,91401722 | 1,31171382 | 4,7951E-06 | 1,31171382 | 1,23555525 | 0,21662399 |
| VCA0172 | 1,2866673 | 5,69048766 | 1,19612432 | 5,5528E-06 | 1,19612432 | 1,23475116 | 0,21692317 |
| VCA0238 | -1,3643006 | -4,0763681 | -1,3829614 | -3,978E-06 | -1,3829614 | -1,2344919 | 0,2170197 |
| gltB | 1,63133696 | 4,48353741 | 1,50973092 | 4,3751E-06 | 1,50973092 | 1,23390744 | 0,21723745 |
| VC0765 | -1,4700044 | -55,347634 | -1,4379755 | -5,401E-05 | -1,4379755 | -1,2338561 | 0,21725659 |
| VC1695 | 1,35203296 | 4,14755298 | 1,27868664 | 4,0472E-06 | 1,27868664 | 1,23360471 | 0,21735028 |
| VC1844 | 1,54639368 | 86,614196 | 1,58378207 | 8,4519E-05 | 1,58378207 | 1,23354756 | 0,21737163 |
| VC0771 | 1,76739046 | 2,23901985 | 1,58573844 | 2,1849E-06 | 1,58573844 | 1,23353157 | 0,21737756 |
| VCA0332 | 1,39381669 | 79,1268438 | 1,23760144 | 7,7213E-05 | 1,23760144 | 1,23337346 | 0,21743651 |
| VC2655 | 1,47695633 | 6,16813261 | 1,4351886 | 6,0189E-06 | 1,4351886 | 1,23332633 | 0,2174541 |
| VC1837 | 1,16625769 | 17,7866077 | 1,18088178 | 1,7356E-05 | 1,18088178 | 1,23228277 | 0,21784354 |
| VC0420 | -1,1374295 | -5,6974681 | -1,1949549 | -5,56E-06 | -1,1949549 | -1,2321476 | 0,217894 |
| VC1025 | 1,53391386 | 7,33537978 | 1,48587148 | 7,1579E-06 | 1,48587148 | 1,23057447 | 0,21848212 |
| vmrA | -1,2036679 | -4,0582487 | -1,2839071 | -3,96E-06 | -1,2839071 | -1,2302189 | 0,2186152 |
| VCA0588 | -1,1459882 | -4,7086813 | -1,2398241 | -4,595E-06 | -1,2398241 | -1,2298566 | 0,21875085 |
| VC0719 | 1,35917806 | 8,66592675 | 1,30407694 | 8,4563E-06 | 1,30407694 | 1,22901354 | 0,21906679 |
| VC1770 | 1,32591237 | 12,7749791 | 1,31099812 | 1,2466E-05 | 1,31099812 | 1,22704768 | 0,21980474 |
| VC1987 | 1,21023805 | 6,5476542 | 1,16568217 | 6,3893E-06 | 1,16568217 | 1,22589463 | 0,22023837 |
| VC2188 | -1,2055713 | -96,000644 | -1,3110098 | -9,368E-05 | -1,3110098 | -1,2254178 | 0,22041791 |
| surE | 1,28711845 | 5,91571282 | 1,23200085 | 5,7726E-06 | 1,23200085 | 1,2250443 | 0,22055859 |
| dapA | 1,26878322 | 46,0188567 | 1,28125238 | 4,4906E-05 | 1,28125238 | 1,22477809 | 0,22065892 |
| hslO | 1,61378304 | 7,49100955 | 1,47707426 | 7,3098E-06 | 1,47707426 | 1,22419399 | 0,22087912 |
| VC1771 | 1,3379561 | 6,84100565 | 1,25690247 | 6,6755E-06 | 1,25690247 | 1,22337917 | 0,22118656 |
| VC2507 | -1,1926164 | -5,4393026 | -1,2998049 | -5,308E-06 | -1,2998049 | -1,223115 | 0,2212863 |
| VCA0396 | 1,20321474 | 11,7754597 | 1,15681947 | 1,1491E-05 | 1,15681947 | 1,22105865 | 0,22206387 |
| VCA0058 | 1,19371196 | 7,23888073 | 1,14718708 | 7,0638E-06 | 1,14718708 | 1,22013386 | 0,22241418 |
| VC2059 | -1,1754104 | -77,539389 | -1,2134948 | -7,566E-05 | -1,2134948 | -1,2184769 | 0,22304285 |
| VC0552 | 1,20471965 | 7,03167677 | 1,15115973 | 6,8616E-06 | 1,15115973 | 1,21753781 | 0,2233997 |
| VC2049 | -1,5904991 | -43,828043 | -1,5265261 | -4,277E-05 | -1,5265261 | -1,216989 | 0,22360845 |

| | | | | | | | |
|---|---|---|---|---|---|---|---|
| tRNA-His-1 | -1,3657278 | -9,5084764 | -1,3885706 | -9,278E-06 | -1,3885706 | -1,2166872 | 0,22372332 |
| leuD | 1,32830562 | 9,80597137 | 1,27750043 | 9,5688E-06 | 1,27750043 | 1,21667762 | 0,22372696 |
| VC1464 | -1,1998172 | -28,595113 | -1,1716241 | -2,79E-05 | -1,1716241 | -1,2158497 | 0,22404225 |
| glnA | -1,3868243 | -174,99045 | -1,4651252 | -0,0001708 | -1,4651252 | -1,2132173 | 0,2250468 |
| VC1279 | -1,4259369 | -13,021091 | -1,4652817 | -1,271E-05 | -1,4652817 | -1,2078332 | 0,22711149 |
| VC0238 | 1,2235571 | 41,6285943 | 1,32313804 | 4,0622E-05 | 1,32313804 | 1,20693657 | 0,22745662 |
| VCA0347 | 1,21112713 | 6,28162738 | 1,16706287 | 6,1297E-06 | 1,16706287 | 1,2053401 | 0,22807208 |
| VCA0372 | -1,4489682 | -23,455136 | -1,4149875 | -2,289E-05 | -1,4149875 | -1,2025745 | 0,22914109 |
| tuf (NC_002505 335305..336530) | 1,24016763 | 666,024964 | 1,40521774 | 0,00064991 | 1,40521774 | 1,19919212 | 0,2304533 |
| VC0184 | 1,5595466 | 3,0945689 | 1,42305336 | 3,0197E-06 | 1,42305336 | 1,19839692 | 0,23076257 |
| VC0170 | 1,67008136 | 5,34561831 | 1,47819526 | 5,2163E-06 | 1,47819526 | 1,19839644 | 0,23076279 |
| VC2429 | 1,11563107 | 9,36440873 | 1,10763297 | 9,1379E-06 | 1,10763297 | 1,19783077 | 0,23098295 |
| VC1457 | 1,27540774 | 7,53266452 | 1,22975467 | 7,3504E-06 | 1,22975467 | 1,19604711 | 0,23167823 |
| VC0167 | 1,26717917 | 5,49253644 | 1,21121237 | 5,3597E-06 | 1,21121237 | 1,19572588 | 0,23180359 |
| VCA0909 | 1,86255224 | 1,9894877 | 1,65410346 | 1,9414E-06 | 1,65410346 | 1,19493626 | 0,23211198 |
| flgJ | -1,2514724 | -25,517143 | -1,2979291 | -2,49E-05 | -1,2979291 | -1,194079 | 0,23244715 |
| murQ | 6,21657164 | 9,04667967 | 4,80139809 | 8,8278E-06 | 4,80139809 | 1,19283948 | 0,23293233 |
| VCA0167 | 1,14925339 | 20,1270672 | 1,17186489 | 1,964E-05 | 1,17186489 | 1,19180989 | 0,23333584 |
| nhaR | 1,39186608 | 7,05845582 | 1,3157105 | 6,8877E-06 | 1,3157105 | 1,19137923 | 0,23350481 |
| VCA0507 | -1,4109011 | -22,246246 | -1,3858221 | -2,171E-05 | -1,3858221 | -1,1908451 | 0,23371446 |
| VCA0726 | -1,2713963 | -4,0785081 | -1,306349 | -3,98E-06 | -1,306349 | -1,1900931 | 0,23400983 |
| VC1139 | 1,5490664 | 13,8338091 | 1,51036597 | 1,3499E-05 | 1,51036597 | 1,1898669 | 0,23409878 |
| tRNA-Leu-4 | -1,4112384 | -12,682727 | -1,4083589 | -1,238E-05 | -1,4083589 | -1,1873442 | 0,23509196 |
| mdoG | 1,23650237 | 12,0202173 | 1,20351097 | 1,1729E-05 | 1,20351097 | 1,18718279 | 0,23515559 |
| VCA1086 | -1,2590239 | -3,3413293 | -1,3270829 | -3,26E-06 | -1,3270829 | -1,1869866 | 0,23523295 |
| VCA0258 | 1,62906452 | 2,6463003 | 1,4310353 | 2,5823E-06 | 1,4310353 | 1,18643134 | 0,23545206 |
| VCA0298 | -1,120272 | -4,8181713 | -1,2157375 | -4,702E-06 | -1,2157375 | -1,1863441 | 0,23548647 |
| tRNA-Gly-2 | -1,5125354 | -25,791458 | -1,4757332 | -2,517E-05 | -1,4757332 | -1,1855173 | 0,23581305 |
| VC2019 | -1,3137272 | -49,305853 | -1,3264072 | -4,811E-05 | -1,3264072 | -1,1853762 | 0,23586879 |
| VC0204 | 2,65416096 | 5,3222375 | 2,22746436 | 5,1935E-06 | 2,22746436 | 1,18220967 | 0,2371226 |
| VCA0756 | 1,80868676 | 2,06321285 | 1,58307841 | 2,0133E-06 | 1,58307841 | 1,18202959 | 0,23719401 |
| VCA0480 | 1,44282242 | 3,58341399 | 1,38603234 | 3,4967E-06 | 1,38603234 | 1,18187194 | 0,23725657 |
| VC2412 | 1,50959075 | 104,319695 | 1,47434529 | 0,0001018 | 1,47434529 | 1,1815903 | 0,23736835 |
| potA | 1,31329013 | 17,1986712 | 1,31161698 | 1,6783E-05 | 1,31161698 | 1,18052237 | 0,23779259 |
| VCA0229 | 2,37809677 | 1,34987744 | 2,23398182 | 1,3172E-06 | 2,23398182 | 1,17819063 | 0,23872067 |
| 16Sf | -1,2157726 | -27,454571 | -1,1921799 | -2,679E-05 | -1,1921799 | -1,1781615 | 0,2387323 |
| rnc | 1,27256346 | 10,0070296 | 1,24251362 | 9,7649E-06 | 1,24251362 | 1,17717116 | 0,23912727 |
| tRNA-Phe-3 | -1,5460292 | -8,3329878 | -1,5926373 | -8,131E-06 | -1,5926373 | -1,176909 | 0,23923191 |
| VCA0835 | -1,1645359 | -4,5511695 | -1,2257242 | -4,441E-06 | -1,2257242 | -1,1767394 | 0,23929957 |
| VC2060 | -1,1820969 | -24,751422 | -1,1558564 | -2,415E-05 | -1,1558564 | -1,1758217 | 0,2396662 |
| VC0355 | -1,5604713 | -53,349894 | -1,4670705 | -5,206E-05 | -1,4670705 | -1,1753534 | 0,23985343 |
| VCA0376 | 1,81546141 | 1,61555581 | 1,79296011 | 1,5765E-06 | 1,79296011 | 1,17304741 | 0,24077684 |
| VCA0745 | -2,1320243 | -80,867471 | -2,6113704 | -7,891E-05 | -2,6113704 | -1,1717926 | 0,24128037 |
| VC2723 | -1,1413503 | -5,1434009 | -1,1951685 | -5,019E-06 | -1,1951685 | -1,1712859 | 0,24148391 |
| VC0421 | -1,1100992 | -6,8835564 | -1,1419025 | -6,717E-06 | -1,1419025 | -1,1696966 | 0,24212313 |

| | | | | | | | |
|---|---|---|---|---|---|---|---|
| VCA0123 | 1,26705863 | 4,94129202 | 1,20887047 | 4,8218E-06 | 1,20887047 | 1,16922039 | 0,24231489 |
| VC1101 | 1,40821737 | 4,04567144 | 1,27556446 | 3,9478E-06 | 1,27556446 | 1,16602837 | 0,24360302 |
| VC0916 | 2,26695812 | 1,37393409 | 2,12958338 | 1,3407E-06 | 2,12958338 | 1,16578969 | 0,24369953 |
| VC0089 | -1,3601761 | -2,2885544 | -1,4922686 | -2,233E-06 | -1,4922686 | -1,1645179 | 0,24421426 |
| lpxD | 1,22565552 | 37,4209498 | 1,21870767 | 3,6516E-05 | 1,21870767 | 1,16251338 | 0,24502704 |
| VC1113 | 8,40583059 | 19,1450746 | 8,22933509 | 1,8682E-05 | 8,22933509 | 1,16109759 | 0,2456023 |
| VCA0764 | 1,30557151 | 7,79747582 | 1,22002241 | 7,6088E-06 | 1,22002241 | 1,16086581 | 0,24569653 |
| VCA0349 | 1,29100896 | 4,82102423 | 1,20530119 | 4,7044E-06 | 1,20530119 | 1,16037296 | 0,24589703 |
| uppP | 1,26131521 | 4,95361793 | 1,19887668 | 4,8338E-06 | 1,19887668 | 1,15936244 | 0,24630852 |
| VC2083 | -1,3351928 | -2,5479181 | -1,4246901 | -2,486E-06 | -1,4246901 | -1,1570786 | 0,24724028 |
| xseA | -1,3210058 | -13,404725 | -1,3507496 | -1,308E-05 | -1,3507496 | -1,1568081 | 0,24735083 |
| VCA0912 | 2,1785413 | 3,97472956 | 2,03255228 | 3,8786E-06 | 2,03255228 | 1,15665001 | 0,24741545 |
| VC0734 | 2,07319032 | 1,76494971 | 1,80198442 | 1,7223E-06 | 1,80198442 | 1,15659562 | 0,24743766 |
| VC0845 | 1,46037762 | 10,7815318 | 1,41905157 | 1,0521E-05 | 1,41905157 | 1,15623564 | 0,24758486 |
| VCA0184 | -1,4744769 | -688,28444 | -1,4049911 | -0,0006716 | -1,4049911 | -1,1562131 | 0,24759406 |
| VC1903 | 1,23698538 | 6,84469885 | 1,17764865 | 6,6791E-06 | 1,17764865 | 1,15570862 | 0,2478004 |
| VCA0545 | 1,36242885 | 11,1670693 | 1,31289208 | 1,0897E-05 | 1,31289208 | 1,15415153 | 0,24843811 |
| VC0730 | -1,3688864 | -4,4901322 | -1,4250261 | -4,382E-06 | -1,4250261 | -1,1539788 | 0,24850894 |
| VCA0951 | 1,49145217 | 18,7317331 | 1,46359393 | 1,8279E-05 | 1,46359393 | 1,15333279 | 0,24877387 |
| VCA0443 | 1,4344734 | 6,53793364 | 1,36018747 | 6,3798E-06 | 1,36018747 | 1,15319962 | 0,24882852 |
| VCA0227 | 2,49424319 | 7,63203881 | 2,35338437 | 7,4474E-06 | 2,35338437 | 1,15297941 | 0,24891892 |
| rplM | 1,24483377 | 806,277974 | 1,50883334 | 0,00078677 | 1,50883334 | 1,15279988 | 0,24899262 |
| VC0358 | -1,4813247 | -32,346252 | -1,4261738 | -3,156E-05 | -1,4261738 | -1,1473255 | 0,2512472 |
| rplD | 1,75245736 | 161,572094 | 1,69459585 | 0,00015766 | 1,69459585 | 1,14729355 | 0,25126037 |
| VC2204 | -1,2923119 | -32,520605 | -1,2650949 | -3,173E-05 | -1,2650949 | -1,1464821 | 0,2515958 |
| VCA1074 | 1,31672987 | 4,15160953 | 1,23592611 | 4,0512E-06 | 1,23592611 | 1,14638995 | 0,25163388 |
| VC0961 | 1,20966796 | 10,4316491 | 1,18526155 | 1,0179E-05 | 1,18526155 | 1,1458957 | 0,25183836 |
| aroB | 1,30864605 | 10,3830197 | 1,25355015 | 1,0132E-05 | 1,25355015 | 1,14561437 | 0,25195482 |
| VC2269 | -1,1168009 | -11,460415 | -1,1805662 | -1,118E-05 | -1,1805662 | -1,1446491 | 0,25235459 |
| VC2159 | 1,18065236 | 25,3921163 | 1,19344909 | 2,4778E-05 | 1,19344909 | 1,14350341 | 0,25282969 |
| hfq | -1,0637216 | -97,853755 | -1,2241267 | -9,549E-05 | -1,2241267 | -1,1431728 | 0,25296694 |
| VCA0444 | 1,36106749 | 7,32581015 | 1,29544663 | 7,1486E-06 | 1,29544663 | 1,14207251 | 0,25342394 |
| VC2050 | -1,5964674 | -88,066726 | -1,5211767 | -8,594E-05 | -1,5211767 | -1,1408613 | 0,2539277 |
| VC2116 | 1,38851442 | 16,2564695 | 1,33402882 | 1,5863E-05 | 1,33402882 | 1,14057809 | 0,25404561 |
| VC0296 | 1,37788914 | 46,7555741 | 1,39828366 | 4,5624E-05 | 1,39828366 | 1,1393583 | 0,25455382 |
| VC1149 | 1,32659094 | 9,88008899 | 1,24669515 | 9,6411E-06 | 1,24669515 | 1,13829995 | 0,25499531 |
| VCA0163 | -1,0977253 | -5,8084235 | -1,2034632 | -5,668E-06 | -1,2034632 | -1,1379273 | 0,2551509 |
| VC0604 | 1,34158759 | 23,9454287 | 1,28303729 | 2,3366E-05 | 1,28303729 | 1,13732678 | 0,25540178 |
| VC0928 | 1,50436992 | 6,55224687 | 1,48482871 | 6,3937E-06 | 1,48482871 | 1,13658759 | 0,25571079 |
| VC0365 | 1,36074815 | 15,8495338 | 1,35233004 | 1,5466E-05 | 1,35233004 | 1,13633417 | 0,2558168 |
| VC2340 | -1,1825872 | -3,4551351 | -1,2843184 | -3,372E-06 | -1,2843184 | -1,1358482 | 0,25602015 |
| VC0271 | 2,13233662 | 19,2230831 | 1,90234114 | 1,8758E-05 | 1,90234114 | 1,13535605 | 0,25622625 |
| tRNA-Asn-4 | -1,380469 | -99,67388 | -1,4242762 | -9,726E-05 | -1,4242762 | -1,1337366 | 0,25690516 |
| VCA0736 | -1,115471 | -5,8596969 | -1,1575937 | -5,718E-06 | -1,1575937 | -1,1331604 | 0,25714694 |
| VC1037 | 1,44778067 | 5,68874055 | 1,3228934 | 5,5511E-06 | 1,3228934 | 1,13287455 | 0,257267 |

| | | | | | | | |
|---|---|---|---|---|---|---|---|
| VC2520 | 1,30184548 | 5,68230631 | 1,23021552 | 5,5448E-06 | 1,23021552 | 1,13287015 | 0,25726884 |
| frsA | 1,36393857 | 3,79159526 | 1,25422488 | 3,6999E-06 | 1,25422488 | 1,13262365 | 0,25737238 |
| VC2391 | 1,22860232 | 32,8699355 | 1,25425791 | 3,2075E-05 | 1,25425791 | 1,13162083 | 0,25779395 |
| asnC | 1,21344067 | 12,4483144 | 1,1649206 | 1,2147E-05 | 1,1649206 | 1,13089721 | 0,25809841 |
| VC0838 | 1,89257258 | 7,29299722 | 1,85004949 | 7,1166E-06 | 1,85004949 | 1,12998123 | 0,25848421 |
| VC2360 | 1,55499642 | 9,01238113 | 1,52847172 | 8,7944E-06 | 1,52847172 | 1,12984249 | 0,25854268 |
| VCA0266 | 1,68572883 | 2,31397253 | 1,45064144 | 2,258E-06 | 1,45064144 | 1,12983957 | 0,25854387 |
| VC1841 | 1,58306568 | 36,9351224 | 1,62921358 | 3,6042E-05 | 1,62921358 | 1,12962608 | 0,2586339 |
| VC1886 | 1,34806173 | 3,41256054 | 1,2847213 | 3,33E-06 | 1,2847213 | 1,12952831 | 0,25867508 |
| VC0823 | 1,80521248 | 50,7450614 | 1,84273815 | 4,9517E-05 | 1,84273815 | 1,12950879 | 0,25868334 |
| potB | 1,35028952 | 13,3399979 | 1,34214562 | 1,3017E-05 | 1,34214562 | 1,12712871 | 0,25968813 |
| tRNA-Gly-1 | -1,6139958 | -29,180906 | -1,5532246 | -2,847E-05 | -1,5532246 | -1,126857 | 0,25980303 |
| VC0097 | 1,14620618 | 7,23550497 | 1,12423065 | 7,0605E-06 | 1,12423065 | 1,12673491 | 0,25985462 |
| VCA0479 | 1,31059466 | 14,2494598 | 1,2891819 | 1,3905E-05 | 1,2891819 | 1,12665907 | 0,25988672 |
| VC1638 | 1,42853669 | 2,78608399 | 1,35689261 | 2,7187E-06 | 1,35689261 | 1,12501575 | 0,2605824 |
| VC1043 | -1,1563919 | -146,12669 | -1,3612217 | -0,0001426 | -1,3612217 | -1,1249171 | 0,26062421 |
| VC2514 | 1,11598897 | 11,7711852 | 1,09677187 | 1,1486E-05 | 1,09677187 | 1,12454165 | 0,26078336 |
| tRNA-Pro-2 | -1,4879052 | -79,321208 | -1,4811284 | -7,74E-05 | -1,4811284 | -1,1232749 | 0,26132084 |
| VC2352 | -1,2540473 | -82,414227 | -1,3496562 | -8,042E-05 | -1,3496562 | -1,1229521 | 0,26145787 |
| VCA0448 | 2,03729755 | 44,3316832 | 2,00814348 | 4,3259E-05 | 2,00814348 | 1,12282996 | 0,2615098 |
| VC1271 | -1,2409648 | -2,7893755 | -1,3544266 | -2,722E-06 | -1,3544266 | -1,1223716 | 0,26170451 |
| minE | 1,34929614 | 3,83633201 | 1,24553354 | 3,7435E-06 | 1,24553354 | 1,12180695 | 0,26194458 |
| VCA0398 | 1,32832014 | 5,51901579 | 1,22596936 | 5,3855E-06 | 1,22596936 | 1,12151027 | 0,26207076 |
| VC1177 | 1,31947659 | 6,68220614 | 1,23745116 | 6,5206E-06 | 1,23745116 | 1,12039567 | 0,26254524 |
| VCA0204 | -1,3420313 | -9,7517019 | -1,4412063 | -9,516E-06 | -1,4412063 | -1,1144518 | 0,26508547 |
| rpsN | 1,41116047 | 127,352007 | 1,27861205 | 0,00012427 | 1,27861205 | 1,11377559 | 0,26537553 |
| VC0048 | 1,52820976 | 22,847532 | 1,47772699 | 2,2295E-05 | 1,47772699 | 1,11311819 | 0,26565774 |
| VC1233 | 1,39268865 | 4,58158155 | 1,31457409 | 4,4707E-06 | 1,31457409 | 1,11267004 | 0,26585022 |
| VCA0124 | 1,15441283 | 13,9602842 | 1,16924686 | 1,3623E-05 | 1,16924686 | 1,11241305 | 0,26596066 |
| VC1572 | 1,8350006 | 1,54778246 | 1,72378611 | 1,5103E-06 | 1,72378611 | 1,11079744 | 0,26665559 |
| nhaA | 1,34535621 | 3,67416955 | 1,28507587 | 3,5853E-06 | 1,28507587 | 1,11026817 | 0,26688353 |
| rhtB | -1,1695695 | -7,5977258 | -1,2876173 | -7,414E-06 | -1,2876173 | -1,1088408 | 0,26749894 |
| VC2164 | 1,26597345 | 4,94135991 | 1,18083168 | 4,8218E-06 | 1,18083168 | 1,10870604 | 0,26755706 |
| VC0483 | 1,33288977 | 16,3940362 | 1,31044984 | 1,5997E-05 | 1,31044984 | 1,10817793 | 0,26778503 |
| VC0242 | 1,33377019 | 106,307781 | 1,27655411 | 0,00010374 | 1,27655411 | 1,10800467 | 0,26785987 |
| VCA0475 | 1,13543114 | 25,4498813 | 1,13743375 | 2,4834E-05 | 1,13743375 | 1,10794996 | 0,26788348 |
| VC0515 | 1,74154048 | 9,54143172 | 1,70870896 | 9,3106E-06 | 1,70870896 | 1,10657938 | 0,2684759 |
| VC0150 | 1,13064555 | 28,3105406 | 1,08604935 | 2,7626E-05 | 1,08604935 | 1,10621082 | 0,26863534 |
| VC0821 | 2,21073141 | 25,6236563 | 2,20936924 | 2,5004E-05 | 2,20936924 | 1,10619403 | 0,26864262 |
| VCA0400 | 1,30487646 | 14,9266964 | 1,29731432 | 1,4566E-05 | 1,29731432 | 1,10440521 | 0,26941747 |
| VC1112 | 9,68537801 | 39,2741517 | 9,43998623 | 3,8324E-05 | 9,43998623 | 1,10400227 | 0,26959224 |
| yieM | 1,36630049 | 5,3191887 | 1,32032 | 5,1905E-06 | 1,32032 | 1,10358866 | 0,26977167 |
| rplS | 1,5398522 | 129,721389 | 1,33988415 | 0,00012658 | 1,33988415 | 1,10283684 | 0,27009809 |
| flgG | -1,1838783 | -20,747406 | -1,2138133 | -2,025E-05 | -1,2138133 | -1,1028017 | 0,27011335 |
| VCA0459 | 1,27768384 | 4,11730399 | 1,21776321 | 4,0177E-06 | 1,21776321 | 1,10130153 | 0,27076549 |

| | | | | | | | |
|---|---|---|---|---|---|---|---|
| VCA0504 | 1,25192311 | 4,73011959 | 1,18671086 | 4,6157E-06 | 1,18671086 | 1,10075764 | 0,2710022 |
| modB | -1,199094 | -3,4081567 | -1,2872735 | -3,326E-06 | -1,2872735 | -1,0998848 | 0,27138237 |
| trpA | -1,1047866 | -4,9388893 | -1,1775186 | -4,819E-06 | -1,1775186 | -1,0990634 | 0,27174045 |
| pheS | 1,2958606 | 6,41212078 | 1,23614276 | 6,257E-06 | 1,23614276 | 1,09808767 | 0,27216625 |
| VCA0432 | 1,31096135 | 14,040141 | 1,31028726 | 1,37E-05 | 1,31028726 | 1,09637345 | 0,27291545 |
| VC2222 | 1,3399948 | 3,70930648 | 1,24183472 | 3,6196E-06 | 1,24183472 | 1,09564083 | 0,27323603 |
| fliD | -1,1770248 | -15,798684 | -1,2059998 | -1,542E-05 | -1,2059998 | -1,0934745 | 0,27418557 |
| VC1509 | -1,2160714 | -3,2060952 | -1,3194645 | -3,129E-06 | -1,3194645 | -1,0922587 | 0,27471943 |
| VC1983 | -1,0842637 | -7,0428282 | -1,1196557 | -6,872E-06 | -1,1196557 | -1,0921481 | 0,27476806 |
| fur | -1,1513244 | -121,53327 | -1,3306122 | -0,0001186 | -1,3306122 | -1,0914614 | 0,27507 |
| VC0091 | 1,44458065 | 3,31667358 | 1,30669678 | 3,2364E-06 | 1,30669678 | 1,09097285 | 0,27528488 |
| rplQ | 1,3214016 | 75,0715681 | 1,16506705 | 7,3255E-05 | 1,16506705 | 1,090743 | 0,27538603 |
| VC1769 | 1,36166807 | 17,3478383 | 1,36603216 | 1,6928E-05 | 1,36603216 | 1,0873087 | 0,27690044 |
| VCA0933 | -1,3470741 | -379,56487 | -1,5054463 | -0,0003704 | -1,5054463 | -1,085985 | 0,2774857 |
| VC0037 | 1,24360933 | 29,2290462 | 1,28292105 | 2,8522E-05 | 1,28292105 | 1,08593285 | 0,27750875 |
| VC0840 | 1,71994939 | 14,3418985 | 1,70082149 | 1,3995E-05 | 1,70082149 | 1,0844276 | 0,27817533 |
| VCA0421 | 1,2325977 | 5,02060078 | 1,16918899 | 4,8991E-06 | 1,16918899 | 1,08388371 | 0,27841641 |
| VC1111 | 2,24788394 | 3,24702131 | 2,21939192 | 3,1685E-06 | 2,21939192 | 1,08317003 | 0,27873303 |
| VC2294 | 1,11493592 | 26,7073059 | 1,13448856 | 2,6061E-05 | 1,13448856 | 1,08245588 | 0,27905007 |
| VCA0199 | 1,29217974 | 30,9797985 | 1,33174132 | 3,023E-05 | 1,33174132 | 1,08225306 | 0,27914014 |
| pspB | 1,33266947 | 6,38003 | 1,28595266 | 6,2257E-06 | 1,28595266 | 1,08204777 | 0,27923136 |
| VC0028 | 1,98899686 | 11,1991713 | 1,95351716 | 1,0928E-05 | 1,95351716 | 1,0799059 | 0,28018416 |
| VC0951 | 1,21932459 | 5,43083895 | 1,153884 | 5,2995E-06 | 1,153884 | 1,07891631 | 0,28062508 |
| VC1862 | 1,66713116 | 2,02942473 | 1,47223111 | 1,9803E-06 | 1,47223111 | 1,07840287 | 0,28085405 |
| VC0980 | 1,44655746 | 4,39289591 | 1,31237038 | 4,2866E-06 | 1,31237038 | 1,07797998 | 0,28104274 |
| VC1717 | 1,26728767 | 14,6608464 | 1,24664431 | 1,4306E-05 | 1,24664431 | 1,07774105 | 0,2811494 |
| tRNA-Met-5 | -1,2071324 | -4,4168022 | -1,2607326 | -4,31E-06 | -1,2607326 | -1,0776858 | 0,28117405 |
| VCA0955 | -1,2300466 | -9,946256 | -1,3008569 | -9,706E-06 | -1,3008569 | -1,0774499 | 0,28127936 |
| VC0977 | 2,13555966 | 14,991466 | 1,91574639 | 1,4629E-05 | 1,91574639 | 1,07659992 | 0,2816591 |
| VCA0171 | 1,3086424 | 4,04107135 | 1,21124101 | 3,9433E-06 | 1,21124101 | 1,07617893 | 0,28184728 |
| VCA0205 | -1,4741176 | -186,62497 | -1,7950998 | -0,0001821 | -1,7950998 | -1,0756243 | 0,28209536 |
| VCA1072 | -3,184438 | -43,623736 | -3,354453 | -4,257E-05 | -3,354453 | -1,0754801 | 0,28215991 |
| VC2046 | -1,1776646 | -5,8090229 | -1,2376297 | -5,668E-06 | -1,2376297 | -1,0746033 | 0,28255242 |
| VCA0201 | 1,22824124 | 34,3058656 | 1,29954125 | 3,3476E-05 | 1,29954125 | 1,07391454 | 0,28286103 |
| VC1075 | -1,1820184 | -3,2165613 | -1,2712001 | -3,139E-06 | -1,2712001 | -1,0734372 | 0,28307504 |
| VCA0051 | -1,1760665 | -15,116448 | -1,2411998 | -1,475E-05 | -1,2411998 | -1,0733104 | 0,28313191 |
| VC0514 | 1,76642912 | 12,0001346 | 1,74820794 | 1,171E-05 | 1,74820794 | 1,072677 | 0,28341613 |
| VC1686 | 1,19215542 | 11,6493175 | 1,19602752 | 1,1368E-05 | 1,19602752 | 1,07235276 | 0,28356168 |
| oppB | 1,33529029 | 8,2533111 | 1,24530612 | 8,0537E-06 | 1,24530612 | 1,06966957 | 0,28476812 |
| VC0187 | 1,16648274 | 5,93027753 | 1,13729947 | 5,7868E-06 | 1,13729947 | 1,06907875 | 0,28503423 |
| VC2385 | 1,28583252 | 44,5455935 | 1,33047982 | 4,3468E-05 | 1,33047982 | 1,06766672 | 0,28567093 |
| tRNA-Thr-6 | -1,4766974 | -22,247622 | -1,4500569 | -2,171E-05 | -1,4500569 | -1,0675945 | 0,28570352 |
| VCA0985 | -1,3639123 | -3,9461978 | -1,5214912 | -3,851E-06 | -1,5214912 | -1,0674933 | 0,28574921 |
| VC1880 | -1,167832 | -6,1019533 | -1,2267738 | -5,954E-06 | -1,2267738 | -1,0660928 | 0,28638173 |
| VC0399 | -1,0736977 | -5,3400411 | -1,1525485 | -5,211E-06 | -1,1525485 | -1,0655368 | 0,28663311 |

| | | | | | | | |
|---|---|---|---|---|---|---|---|
| VC1963 | 1,21467734 | 8,26748412 | 1,18020711 | 8,0675E-06 | 1,18020711 | 1,06536262 | 0,28671191 |
| VC1455 | -1,2293032 | -30,729107 | -1,1802828 | -2,999E-05 | -1,1802828 | -1,0653016 | 0,28673953 |
| 23Sh | 1,06915797 | 1579,79609 | 1,24272814 | 0,00154158 | 1,24272814 | 1,06422331 | 0,28722761 |
| rpsB | 1,62524647 | 196,757134 | 1,33404626 | 0,000192 | 1,33404626 | 1,06340993 | 0,28759617 |
| VC0976 | 1,30458878 | 22,7422852 | 1,25820474 | 2,2192E-05 | 1,25820474 | 1,06308262 | 0,28774456 |
| VC0406 | -1,1177195 | -6,4604621 | -1,1520172 | -6,304E-06 | -1,1520172 | -1,0623258 | 0,28808785 |
| VC1481 | 1,2881433 | 55,1932327 | 1,26205581 | 5,3858E-05 | 1,26205581 | 1,06214538 | 0,28816976 |
| ksgA | 1,2120309 | 14,2792907 | 1,17449801 | 1,3934E-05 | 1,17449801 | 1,06143879 | 0,28849058 |
| VCA0551 | 1,2224081 | 18,3315418 | 1,20261009 | 1,7888E-05 | 1,20261009 | 1,06010825 | 0,28909542 |
| VCA1071 | -3,0842521 | -53,332773 | -3,1631736 | -5,204E-05 | -3,1631736 | -1,0592626 | 0,28948027 |
| VCA1022 | 1,38852897 | 2,64249951 | 1,3310716 | 2,5786E-06 | 1,3310716 | 1,05923286 | 0,28949379 |
| VC1661 | 1,22298443 | 12,139765 | 1,22283276 | 1,1846E-05 | 1,22283276 | 1,05832277 | 0,28990838 |
| VC1063 | 1,4652687 | 13,247062 | 1,42932554 | 1,2927E-05 | 1,42932554 | 1,05818558 | 0,28997091 |
| VC0559 | -1,1136175 | -5,3381907 | -1,1501268 | -5,209E-06 | -1,1501268 | -1,0574574 | 0,29030294 |
| VC0975 | 1,17882806 | 11,9512609 | 1,13907113 | 1,1662E-05 | 1,13907113 | 1,05711231 | 0,29046038 |
| rpsH | 1,53754338 | 400,091863 | 1,39952742 | 0,00039041 | 1,39952742 | 1,05678261 | 0,29061088 |
| VC2363 | 1,71096654 | 20,7651656 | 1,69226963 | 2,0263E-05 | 1,69226963 | 1,05650049 | 0,2907397 |
| VC0295 | 1,38597099 | 36,2684997 | 1,39895387 | 3,5391E-05 | 1,39895387 | 1,05639582 | 0,29078748 |
| VCA0125 | -1,1213732 | -15,327827 | -1,1053996 | -1,496E-05 | -1,1053996 | -1,0549884 | 0,29143066 |
| VC1374 | 1,61354216 | 3,57864895 | 1,49682056 | 3,4921E-06 | 1,49682056 | 1,05479191 | 0,29152056 |
| tRNA-Thr-2 | -1,1861389 | -2,8380461 | -1,3002995 | -2,769E-06 | -1,3002995 | -1,0542844 | 0,29175275 |
| VCA0636 | -1,2568084 | -10,841905 | -1,2763043 | -1,058E-05 | -1,2763043 | -1,0539057 | 0,29192612 |
| VC1721 | 1,18861125 | 5,85558678 | 1,13476009 | 5,7139E-06 | 1,13476009 | 1,05308107 | 0,29230388 |
| VC0820 | 1,80365381 | 6,22340867 | 1,69816086 | 6,0729E-06 | 1,69816086 | 1,05285882 | 0,29240576 |
| VC1539a | -1,3478366 | -21,903435 | -1,4937962 | -2,137E-05 | -1,4937962 | -1,050546 | 0,29346719 |
| galM | -1,6432854 | -155,75155 | -1,9100844 | -0,000152 | -1,9100844 | -1,0484325 | 0,29443945 |
| VC0828 | -1,3077858 | -12,386839 | -1,3186261 | -1,209E-05 | -1,3186261 | -1,0470901 | 0,29505806 |
| VCA0575 | 1,14714295 | 7,72032058 | 1,13727121 | 7,5336E-06 | 1,13727121 | 1,04705171 | 0,29507577 |
| VC2000 | -1,9478899 | -1560,8708 | -1,6185649 | -0,0015231 | -1,6185649 | -1,0467763 | 0,29520283 |
| VCA0336 | 1,37816166 | 2,75671266 | 1,30496522 | 2,69E-06 | 1,30496522 | 1,04604794 | 0,29553894 |
| VC1839 | 1,12160534 | 7,83451238 | 1,12104191 | 7,645E-06 | 1,12104191 | 1,04505195 | 0,295999 |
| VC2150 | 1,33734442 | 3,24603035 | 1,2522267 | 3,1675E-06 | 1,2522267 | 1,04431016 | 0,29634195 |
| acpP | 1,0618922 | 570,345142 | 1,26338121 | 0,00055655 | 1,26338121 | 1,04320033 | 0,29685556 |
| VC1267 | 1,85696653 | 5,61541667 | 1,70035059 | 5,4796E-06 | 1,70035059 | 1,04261378 | 0,29712724 |
| gltA | 2,71058477 | 136,924114 | 2,42699184 | 0,00013361 | 2,42699184 | 1,04202678 | 0,29739934 |
| VC1790 | -1,370915 | -20,267618 | -1,3424371 | -1,978E-05 | -1,3424371 | -1,0416622 | 0,29756837 |
| VC2568 | 1,3396977 | 35,9980318 | 1,3486465 | 3,5127E-05 | 1,3486465 | 1,04110655 | 0,29782617 |
| VC1866 | -1,4951043 | -751,24259 | -1,4446331 | -0,0007331 | -1,4446331 | -1,0397641 | 0,29844957 |
| VC1354 | -1,2108047 | -2,9940303 | -1,2733144 | -2,922E-06 | -1,2733144 | -1,0391851 | 0,2987187 |
| VC0439 | -1,2483208 | -8,9968118 | -1,3252352 | -8,779E-06 | -1,3252352 | -1,0391031 | 0,29875685 |
| rimM | 1,49320248 | 183,688256 | 1,24868877 | 0,00017924 | 1,24868877 | 1,03745872 | 0,29952217 |
| VC0246 | 1,43016821 | 129,887346 | 1,35269726 | 0,00012675 | 1,35269726 | 1,03688425 | 0,29978985 |
| VCA0655 | -1,2771535 | -8,1425571 | -1,4957102 | -7,946E-06 | -1,4957102 | -1,0367703 | 0,299843 |
| cysB | 1,30878471 | 25,7648097 | 1,26120843 | 2,5142E-05 | 1,26120843 | 1,03498689 | 0,30067509 |
| VCA0087 | 2,41484869 | 12,9364594 | 2,36698006 | 1,2624E-05 | 2,36698006 | 1,03494937 | 0,30069264 |

| | | | | | | | |
|---|---|---|---|---|---|---|---|
| radC | -1,4375863 | -6,5448897 | -1,5451141 | -6,387E-06 | -1,5451141 | -1,0343166 | 0,30098824 |
| VC2171 | -1,2536445 | -3,4951544 | -1,3252198 | -3,411E-06 | -1,3252198 | -1,0335379 | 0,30135228 |
| VCA0585 | 1,33221157 | 3,24303152 | 1,24662751 | 3,1646E-06 | 1,24662751 | 1,03346222 | 0,30138768 |
| VCA0584 | 1,32181257 | 3,86688929 | 1,20254089 | 3,7733E-06 | 1,20254089 | 1,03285737 | 0,30167069 |
| VC2210 | 1,68057881 | 5,41216763 | 1,60202122 | 5,2812E-06 | 1,60202122 | 1,03283102 | 0,30168304 |
| VC1064 | 1,13471875 | 29,507271 | 1,12894849 | 2,8793E-05 | 1,12894849 | 1,03187829 | 0,30212918 |
| VC0880 | 2,11264313 | 2,2730487 | 2,00204718 | 2,2181E-06 | 2,00204718 | 1,03149999 | 0,30230647 |
| VC1236 | 1,4070317 | 2,62756632 | 1,31136225 | 2,564E-06 | 1,31136225 | 1,03047757 | 0,30278592 |
| VC2641 | 1,38693139 | 2,72515171 | 1,29851858 | 2,6592E-06 | 1,29851858 | 1,03043335 | 0,30280666 |
| VC2224 | -1,1657772 | -2,9322494 | -1,2744807 | -2,861E-06 | -1,2744807 | -1,030335 | 0,30285283 |
| VC0906 | 1,49787599 | 3,06484134 | 1,37698449 | 2,9907E-06 | 1,37698449 | 1,02988577 | 0,30306368 |
| VC0959 | 1,23076009 | 57,7124938 | 1,19244557 | 5,6316E-05 | 1,19244557 | 1,02947083 | 0,30325852 |
| prpB | 1,67223394 | 2,22427214 | 1,43156347 | 2,1705E-06 | 1,43156347 | 1,02906934 | 0,30344713 |
| VC0386 | 2,24453185 | 28,2980287 | 2,25743408 | 2,7613E-05 | 2,25743408 | 1,02754977 | 0,30416173 |
| glmM | 1,12771239 | 12,8912933 | 1,1275038 | 1,2579E-05 | 1,1275038 | 1,02384114 | 0,30591036 |
| VCA0401 | 1,29890316 | 11,2732047 | 1,27391283 | 1,1E-05 | 1,27391283 | 1,02311166 | 0,30625511 |
| VCA1044 | -1,2750471 | -33,55247 | -1,3024154 | -3,274E-05 | -1,3024154 | -1,0218187 | 0,30686678 |
| fadI | 1,56507746 | 2,86667326 | 1,392629 | 2,7973E-06 | 1,392629 | 1,02178935 | 0,30688066 |
| VC2366 | 1,10760773 | 12,9210107 | 1,10813066 | 1,2608E-05 | 1,10813066 | 1,02098398 | 0,30726208 |
| VC1362 | -1,2674752 | -2,1134957 | -1,3934044 | -2,062E-06 | -1,3934044 | -1,020879 | 0,3073118 |
| glyS | 1,27835099 | 10,4392272 | 1,25057034 | 1,0187E-05 | 1,25057034 | 1,02083919 | 0,30733067 |
| VC1799 | 1,63541116 | 1,79049497 | 1,48103487 | 1,7472E-06 | 1,48103487 | 1,02051652 | 0,3074836 |
| VC1260 | 1,41015477 | 2,82674944 | 1,29571556 | 2,7584E-06 | 1,29571556 | 1,01891512 | 0,30824331 |
| VC1966 | -1,1083666 | -4,3755701 | -1,1707325 | -4,27E-06 | -1,1707325 | -1,0161055 | 0,30957921 |
| VCA0922 | -1,109971 | -4,1339699 | -1,1815608 | -4,034E-06 | -1,1815608 | -1,0159609 | 0,30964804 |
| tarA | 1,54823761 | 3,23736227 | 1,42890286 | 3,159E-06 | 1,42890286 | 1,0134102 | 0,31086431 |
| VCA0517 | -2,3335049 | -190,51346 | -2,5813756 | -0,0001859 | -2,5813756 | -1,0131827 | 0,31097296 |
| VCA0057 | 1,30235904 | 3,16769924 | 1,24141854 | 3,0911E-06 | 1,24141854 | 1,01171823 | 0,31167284 |
| VC1521 | -1,1257741 | -5,0152038 | -1,2154502 | -4,894E-06 | -1,2154502 | -1,0117103 | 0,31167663 |
| VC0157 | 1,40361473 | 2,73132567 | 1,285234 | 2,6653E-06 | 1,285234 | 1,01131171 | 0,31186731 |
| tRNA-Gly-7 | -1,4163864 | -7,6074654 | -1,4196355 | -7,423E-06 | -1,4196355 | -1,0112108 | 0,31191559 |
| VC0040 | 1,20105997 | 12,1922703 | 1,19378563 | 1,1897E-05 | 1,19378563 | 1,01045127 | 0,31227919 |
| VC1470 | 1,16434077 | 5,69967661 | 1,12692301 | 5,5618E-06 | 1,12692301 | 1,01016003 | 0,31241866 |
| VC2369 | -1,0941122 | -4,7987884 | -1,1518366 | -4,683E-06 | -1,1518366 | -1,0078996 | 0,31350269 |
| VCA0241 | -1,6812725 | -2,5555007 | -1,8933491 | -2,494E-06 | -1,8933491 | -1,0069516 | 0,31395808 |
| ruvA | 1,34926527 | 4,25694983 | 1,29973852 | 4,154E-06 | 1,29973852 | 1,00598293 | 0,31442383 |
| VC1599 | -1,1092735 | -5,7927026 | -1,2056218 | -5,653E-06 | -1,2056218 | -1,0042754 | 0,31524591 |
| VCA0465 | 1,21445196 | 6,85217834 | 1,15446419 | 6,6864E-06 | 1,15446419 | 1,00365137 | 0,31554673 |
| VCA0576 | 1,91873244 | 8,05262376 | 1,82846607 | 7,8578E-06 | 1,82846607 | 1,00330067 | 0,31571589 |
| VC1885 | 1,44075682 | 2,45003131 | 1,31733988 | 2,3908E-06 | 1,31733988 | 1,0032654 | 0,31573286 |
| VC2139 | -1,139571 | -22,043977 | -1,1400826 | -2,151E-05 | -1,1400826 | -1,0030174 | 0,31585251 |
| VC2177 | -1,1141698 | -5,1056585 | -1,1796956 | -4,982E-06 | -1,1796956 | -1,0024561 | 0,31612342 |
| VC0811 | 1,41689733 | 39,4538967 | 1,42701454 | 3,8499E-05 | 1,42701454 | 1,00161933 | 0,31652755 |
| VC2726 | -1,1212776 | -4,1674058 | -1,1744298 | -4,067E-06 | -1,1744298 | -1,0014671 | 0,31660108 |
| moaC | 1,37204066 | 3,44969314 | 1,31760665 | 3,3662E-06 | 1,31760665 | 1,00134533 | 0,31665993 |

| | | | | | | | |
|---|---|---|---|---|---|---|---|
| metN | 1,5336262 | 2,51552404 | 1,38109206 | 2,4547E-06 | 1,38109206 | 1,00120309 | 0,31672867 |
| VC0533 | 1,37477637 | 18,7644582 | 1,3018961 | 1,8311E-05 | 1,3018961 | 1,00093646 | 0,3168576 |
| 23Se | 1,03487991 | 12615,6156 | 1,09423534 | 0,01231043 | 1,09423534 | 0,99986855 | 0,3173742 |
| VC0814 | 1,39892454 | 11,9501847 | 1,34864667 | 1,1661E-05 | 1,34864667 | 0,99895559 | 0,31781625 |
| cadB | 79,8664375 | 105,842743 | 77,9336526 | 0,00010328 | 77,9336526 | 0,99860529 | 0,31798602 |
| VC0201 | 1,74895209 | 2,57157948 | 1,63053579 | 2,5094E-06 | 1,63053579 | 0,99825031 | 0,31815805 |
| atpC | -1,0016612 | -102,88813 | -1,1626375 | -0,0001004 | -1,1626375 | -0,9982377 | 0,31816415 |
| VC2381 | -1,1751453 | -3,7198169 | -1,2417597 | -3,63E-06 | -1,2417597 | -0,9981348 | 0,31821403 |
| VCA0966 | 1,38687104 | 6,68872324 | 1,27730106 | 6,5269E-06 | 1,27730106 | 0,99772343 | 0,31841354 |
| VC2161 | -1,2229129 | -13,617937 | -1,2625433 | -1,329E-05 | -1,2625433 | -0,9963596 | 0,31907553 |
| VC1138 | 1,58994541 | 8,39220769 | 1,57709696 | 8,1892E-06 | 1,57709696 | 0,9960893 | 0,31920682 |
| VC0281 | 65,61349 | 69,1291103 | 61,9550509 | 6,7457E-05 | 61,9550509 | 0,99538132 | 0,31955092 |
| VC0493 | 1,513257 | 29,718619 | 1,46574692 | 2,9E-05 | 1,46574692 | 0,99537286 | 0,319555 |
| 16Se | 1,0844878 | 9839,64627 | 1,1830185 | 0,00960161 | 1,1830185 | 0,9953495 | 0,31956636 |
| mukB | 1,20401841 | 4,5405581 | 1,15587313 | 4,4307E-06 | 1,15587313 | 0,99242103 | 0,32099222 |
| VCA0010 | -1,3545699 | -5,6562512 | -1,3790651 | -5,519E-06 | -1,3790651 | -0,9923049 | 0,32104884 |
| VC0162 | -1,6893232 | -34,984839 | -1,6220275 | -3,414E-05 | -1,6220275 | -0,9912427 | 0,32156716 |
| VCA0692 | 1,7117762 | 10,0695924 | 1,603459 | 9,826E-06 | 1,603459 | 0,98959858 | 0,32237041 |
| VCA0792 | -1,3501515 | -19,511777 | -1,3240405 | -1,904E-05 | -1,3240405 | -0,9878963 | 0,32320349 |
| VC1067 | -1,1271894 | -5,7613052 | -1,1749591 | -5,622E-06 | -1,1749591 | -0,9867223 | 0,32377885 |
| VC1098 | -1,4108235 | -75,560874 | -1,3367932 | -7,373E-05 | -1,3367932 | -0,9861193 | 0,32407463 |
| VCA0807 | -1,233564 | -10,223833 | -1,3682042 | -9,977E-06 | -1,3682042 | -0,983515 | 0,32535406 |
| VC0637 | 1,11579972 | 23,4059468 | 1,10050709 | 2,284E-05 | 1,10050709 | 0,98311682 | 0,32555001 |
| VCA0368 | 1,29735351 | 2,97511464 | 1,25797243 | 2,9031E-06 | 1,25797243 | 0,9829296 | 0,32564213 |
| VCA0569 | -1,1658735 | -14,717862 | -1,180488 | -1,436E-05 | -1,180488 | -0,9828557 | 0,32567849 |
| VC2725 | -1,1044422 | -5,3807909 | -1,1406413 | -5,251E-06 | -1,1406413 | -0,9825225 | 0,32584254 |
| VC0057 | 1,24861644 | 4,16543977 | 1,16641392 | 4,0647E-06 | 1,16641392 | 0,9797628 | 0,32720326 |
| VC2371 | -2,1958345 | -39,038209 | -1,944854 | -3,809E-05 | -1,944854 | -0,979496 | 0,32733498 |
| VC1163 | 1,50507882 | 1,91687376 | 1,3991248 | 1,8705E-06 | 1,3991248 | 0,97810747 | 0,32802121 |
| VC0351 | 1,21445967 | 4,69276483 | 1,1765933 | 4,5792E-06 | 1,1765933 | 0,97785703 | 0,32814508 |
| VC2700 | -6,4095216 | -23,110127 | -6,783909 | -2,255E-05 | -6,783909 | -0,9778249 | 0,32816101 |
| VCA0491 | 1,34845922 | 5,67299826 | 1,29425186 | 5,5358E-06 | 1,29425186 | 0,97732214 | 0,32840973 |
| VC1512 | -1,6570595 | -20,497979 | -1,7113671 | -2E-05 | -1,7113671 | -0,9765364 | 0,32879879 |
| VC0288 | -1,581715 | -5,4907433 | -1,9119742 | -5,358E-06 | -1,9119742 | -0,9759419 | 0,32909331 |
| VC1487 | 1,27824956 | 6,72326085 | 1,2522701 | 6,5606E-06 | 1,2522701 | 0,97520976 | 0,32945628 |
| VC1413 | 1,33221034 | 3,63461046 | 1,22355528 | 3,5467E-06 | 1,22355528 | 0,97414568 | 0,32998426 |
| VC2184 | -1,2884002 | -18,453812 | -1,2930064 | -1,801E-05 | -1,2930064 | -0,9728922 | 0,33060693 |
| VC0925 | 1,36650851 | 2,47030467 | 1,29274443 | 2,4105E-06 | 1,29274443 | 0,97275723 | 0,33067401 |
| VC1183 | 1,90121634 | 6,85885598 | 1,66988391 | 6,6929E-06 | 1,66988391 | 0,97101827 | 0,33153924 |
| VC1216 | -1,2997712 | -10,911541 | -1,3232254 | -1,065E-05 | -1,3232254 | -0,9704451 | 0,33182474 |
| VC2392 | -1,0582406 | -5,4486156 | -1,1221948 | -5,317E-06 | -1,1221948 | -0,9701693 | 0,33196215 |
| VC0353 | 1,17732539 | 5,1307705 | 1,1300893 | 5,0067E-06 | 1,1300893 | 0,96957796 | 0,33225693 |
| VC1800 | 1,8793305 | 1,22081449 | 1,6906332 | 1,1913E-06 | 1,6906332 | 0,96957739 | 0,33225722 |
| tRNA-Leu-7 | -1,3597859 | -12,069528 | -1,3891457 | -1,178E-05 | -1,3891457 | -0,9690719 | 0,33250939 |
| VCA0143 | -1,1764311 | -2,6596072 | -1,2665105 | -2,595E-06 | -1,2665105 | -0,9686126 | 0,33273853 |

| | | | | | | | |
|---|---|---|---|---|---|---|---|
| VCA0521 | -1,3732296 | -13,199908 | -1,3967462 | -1,288E-05 | -1,3967462 | -0,9674543 | 0,33331705 |
| VC1051 | 1,2605785 | 18,1507266 | 1,23857071 | 1,7712E-05 | 1,23857071 | 0,96700074 | 0,33354371 |
| VC2041 | -1,1104878 | -7,8576092 | -1,1020987 | -7,668E-06 | -1,1020987 | -0,9668494 | 0,33361936 |
| VC0831 | 1,6336747 | 4,55128933 | 1,55163207 | 4,4412E-06 | 1,55163207 | 0,96655717 | 0,3337655 |
| VC0839 | 1,85416494 | 7,17626526 | 1,78613085 | 7,0027E-06 | 1,78613085 | 0,96576211 | 0,3341633 |
| valS | 1,26514079 | 7,81157064 | 1,21342488 | 7,6226E-06 | 1,21342488 | 0,96481893 | 0,33463557 |
| VCA0294 | -1,1801762 | -8,2808768 | -1,2700168 | -8,081E-06 | -1,2700168 | -0,9647386 | 0,33467579 |
| VCA0621 | 1,58539947 | 10,0286187 | 1,4731735 | 9,786E-06 | 1,4731735 | 0,96281821 | 0,3356388 |
| VC2607 | 1,54794377 | 7,43893855 | 1,38981437 | 7,259E-06 | 1,38981437 | 0,96124562 | 0,33642875 |
| VC2643 | 1,74213817 | 1,38610521 | 1,56912105 | 1,3526E-06 | 1,56912105 | 0,95977506 | 0,33716847 |
| VCA0920 | -1,0803748 | -4,9954048 | -1,1307435 | -4,875E-06 | -1,1307435 | -0,9589568 | 0,33758056 |
| VC2149 | -1,1418463 | -41,166415 | -1,1744584 | -4,017E-05 | -1,1744584 | -0,9588323 | 0,33764327 |
| lolA | 1,19768769 | 7,41801071 | 1,16132321 | 7,2386E-06 | 1,16132321 | 0,9581941 | 0,33796494 |
| VCA1043 | 1,27663897 | 44,1720394 | 1,29261566 | 4,3103E-05 | 1,29261566 | 0,95779817 | 0,3381646 |
| VC0461 | 1,21116758 | 4,53736518 | 1,14451675 | 4,4276E-06 | 1,14451675 | 0,95777618 | 0,33817567 |
| VC2549 | 1,32297032 | 4,2010046 | 1,21616309 | 4,0994E-06 | 1,21616309 | 0,95618788 | 0,33897736 |
| VC1262 | 1,50629254 | 69,3387995 | 1,4151381 | 6,7661E-05 | 1,4151381 | 0,95433054 | 0,33991641 |
| VCA0498 | 1,17913188 | 28,8497747 | 1,19781463 | 2,8152E-05 | 1,19781463 | 0,95413394 | 0,34001591 |
| VC0117 | -1,1768664 | -6,6477647 | -1,2649275 | -6,487E-06 | -1,2649275 | -0,9534482 | 0,34036305 |
| VC0638 | 1,14095891 | 5,61924124 | 1,1139806 | 5,4833E-06 | 1,1139806 | 0,9534008 | 0,34038707 |
| VC1256 | 1,55620223 | 24,9912467 | 1,54052786 | 2,4387E-05 | 1,54052786 | 0,95334519 | 0,34041526 |
| VC2100 | -2,0960884 | -11,856781 | -2,0473411 | -1,157E-05 | -2,0473411 | -0,9525408 | 0,34082284 |
| VCA0520 | -1,4969378 | -20,538666 | -1,5000606 | -2,004E-05 | -1,5000606 | -0,9524399 | 0,34087397 |
| VCA0866 | 2,6417968 | 0,93638165 | 2,10390483 | 9,1373E-07 | 2,10390483 | 0,95228051 | 0,34095478 |
| VC2036 | 1,49231221 | 30,868724 | 1,43366816 | 3,0122E-05 | 1,43366816 | 0,95220814 | 0,34099151 |
| VC0610 | 1,26621771 | 3,60248758 | 1,21168037 | 3,5153E-06 | 1,21168037 | 0,95107766 | 0,34156499 |
| VCA0460 | 1,34621322 | 2,26896027 | 1,30649548 | 2,2141E-06 | 1,30649548 | 0,95106816 | 0,34156981 |
| VC0139 | -1,1300229 | -6,3308803 | -1,2132728 | -6,178E-06 | -1,2132728 | -0,9507669 | 0,34172276 |
| VCA0309 | 1,16696804 | 16,5170955 | 1,19615879 | 1,6118E-05 | 1,19615879 | 0,94930961 | 0,34246322 |
| VC1189 | -1,5384652 | -88,850309 | -1,5825535 | -8,67E-05 | -1,5825535 | -0,9488529 | 0,34269548 |
| VCA0044 | 1,24664314 | 3,5832227 | 1,18262759 | 3,4965E-06 | 1,18262759 | 0,94840857 | 0,34292154 |
| VCA0540 | -2,1850432 | -358,76025 | -2,1029397 | -0,0003501 | -2,1029397 | -0,9481831 | 0,34303629 |
| VC0952 | 1,167887 | 11,2972713 | 1,14714836 | 1,1024E-05 | 1,14714836 | 0,9468526 | 0,34371393 |
| VC0275 | 1,49803062 | 3,00461295 | 1,40396928 | 2,9319E-06 | 1,40396928 | 0,94646528 | 0,34391137 |
| VCA0997 | 1,34314851 | 2,44048468 | 1,27865832 | 2,3814E-06 | 1,27865832 | 0,94623111 | 0,34403076 |
| VCA0817 | 1,34492134 | 4,62956531 | 1,23706732 | 4,5176E-06 | 1,23706732 | 0,94589938 | 0,34419996 |
| VC2044 | 1,26085094 | 29,7917035 | 1,27351715 | 2,9071E-05 | 1,27351715 | 0,94587273 | 0,34421356 |
| VCA0679 | 2,10231012 | 78,6822202 | 2,17184018 | 7,6779E-05 | 2,17184018 | 0,94509283 | 0,34461153 |
| rpmF | 1,27326814 | 39,8851147 | 1,27123003 | 3,892E-05 | 1,27123003 | 0,944937 | 0,34469111 |
| VC2295 | 1,12159501 | 33,2845893 | 1,09714456 | 3,2479E-05 | 1,09714456 | 0,94430352 | 0,34501463 |
| VC1639 | 1,33892023 | 3,24158807 | 1,28682401 | 3,1632E-06 | 1,28682401 | 0,94306651 | 0,34564693 |
| VCA0168 | 1,1890877 | 6,54362692 | 1,149193 | 6,3853E-06 | 1,149193 | 0,94232658 | 0,3460255 |
| VCA0940 | -1,2474421 | -18,622679 | -1,4044164 | -1,817E-05 | -1,4044164 | -0,9419661 | 0,34621005 |
| VCA1035 | 1,99186677 | 6,58893953 | 1,89064563 | 6,4295E-06 | 1,89064563 | 0,94175203 | 0,34631967 |
| VC0202 | 1,87758319 | 2,5668851 | 1,76025402 | 2,5048E-06 | 1,76025402 | 0,94164964 | 0,3463721 |

| | | | | | | | |
|---|---|---|---|---|---|---|---|
| VC2361 | 1,41022548 | 567,626473 | 1,45943854 | 0,00055389 | 1,45943854 | 0,94109945 | 0,34665395 |
| VC1568 | 1,41863592 | 2,09046894 | 1,32878199 | 2,0399E-06 | 1,32878199 | 0,94096546 | 0,3467226 |
| VC2261 | 1,20756075 | 10,0146615 | 1,18718575 | 9,7724E-06 | 1,18718575 | 0,93850583 | 0,34798457 |
| fliF | -1,0689297 | -7,1174141 | -1,1033047 | -6,945E-06 | -1,1033047 | -0,9381876 | 0,34814805 |
| VCA0586 | -1,1951509 | -2,6642085 | -1,2549064 | -2,6E-06 | -1,2549064 | -0,9363108 | 0,34911323 |
| VC2769 | 1,22947122 | 76,3845321 | 1,18346635 | 7,4537E-05 | 1,18346635 | 0,93530019 | 0,34963369 |
| VC1200 | 1,1981783 | 4,96268281 | 1,12484351 | 4,8426E-06 | 1,12484351 | 0,93529232 | 0,34963771 |
| fumC | 1,56111182 | 5,32445058 | 1,48072387 | 5,1956E-06 | 1,48072387 | 0,93507585 | 0,34974925 |
| 16Sh | 1,21288012 | 1570,29384 | 1,62979901 | 0,00153231 | 1,62979901 | 0,93501636 | 0,34977994 |
| VC1181 | 1,26640791 | 3,59572901 | 1,17853749 | 3,5087E-06 | 1,17853749 | 0,93451867 | 0,35003645 |
| VCA1102 | -1,2238673 | -2,0284837 | -1,3349625 | -1,979E-06 | -1,3349625 | -0,9343424 | 0,35012736 |
| VC2370 | -1,6397432 | -13,004777 | -1,5820313 | -1,269E-05 | -1,5820313 | -0,9343071 | 0,35014555 |
| flgD | -1,142471 | -34,394521 | -1,1598407 | -3,356E-05 | -1,1598407 | -0,9339249 | 0,35034271 |
| VC0249 | 1,39977229 | 337,11208 | 1,34504263 | 0,00032896 | 1,34504263 | 0,93303278 | 0,35080308 |
| VC0385 | 2,6121295 | 16,6264574 | 2,66330487 | 1,6224E-05 | 2,66330487 | 0,93282952 | 0,35090806 |
| VC0027 | 1,75862389 | 16,2440824 | 1,74654244 | 1,5851E-05 | 1,74654244 | 0,9326826 | 0,35098394 |
| VC1201 | -1,2121385 | -28,733886 | -1,1712714 | -2,804E-05 | -1,1712714 | -0,93258 | 0,35103695 |
| nusG | 1,18783257 | 36,3279917 | 1,19952785 | 3,5449E-05 | 1,19952785 | 0,93239074 | 0,35113468 |
| VC1948 | 2,6228052 | 0,66174064 | 2,5575043 | 6,4573E-07 | 2,5575043 | 0,93228057 | 0,35119159 |
| VC0789 | 7,54232871 | 328,142614 | 7,7996639 | 0,0003202 | 7,7996639 | 0,93170856 | 0,35148723 |
| VC2241 | -1,3997387 | -69,984235 | -1,4514572 | -6,829E-05 | -1,4514572 | -0,9315507 | 0,35156879 |
| VC1904 | 1,32868548 | 66,0794375 | 1,29526212 | 6,4481E-05 | 1,29526212 | 0,93012398 | 0,35230696 |
| VC1126 | 1,28904453 | 15,4052717 | 1,29736031 | 1,5033E-05 | 1,29736031 | 0,93009006 | 0,35232451 |
| VCA0721 | 1,45968442 | 2,66089172 | 1,36455018 | 2,5965E-06 | 1,36455018 | 0,92730252 | 0,35376953 |
| VC0175 | 1,30594284 | 43,3025352 | 1,32730676 | 4,2255E-05 | 1,32730676 | 0,92600436 | 0,35444376 |
| VC1384 | -1,1607887 | -152,38011 | -1,2825112 | -0,0001487 | -1,2825112 | -0,9257869 | 0,35455678 |
| VCA0467 | 1,47319985 | 7,50789225 | 1,47970941 | 7,3263E-06 | 1,47970941 | 0,92573903 | 0,35458168 |
| VC1420 | 1,34178632 | 2,55520685 | 1,25147433 | 2,4934E-06 | 1,25147433 | 0,92531466 | 0,35480229 |
| rfaD | 1,16409789 | 16,1722493 | 1,17782333 | 1,5781E-05 | 1,17782333 | 0,92523861 | 0,35484184 |
| VCA0524 | -1,2219361 | -2,2413319 | -1,2909097 | -2,187E-06 | -1,2909097 | -0,9240399 | 0,35546556 |
| VCA1037 | 1,39752978 | 3,37038534 | 1,26055143 | 3,2889E-06 | 1,26055143 | 0,92320678 | 0,35589949 |
| VCA0055 | 1,47859878 | 2,01053357 | 1,32880097 | 1,9619E-06 | 1,32880097 | 0,92282253 | 0,35609973 |
| tRNA-Gly-5 | -1,1879442 | -7,1573255 | -1,2046356 | -6,984E-06 | -1,2046356 | -0,9213923 | 0,35684571 |
| VC0727 | 1,28967439 | 5,57608102 | 1,19986457 | 5,4412E-06 | 1,19986457 | 0,92038727 | 0,35737046 |
| VC0516 | 1,47626351 | 42,8307047 | 1,56318296 | 4,1795E-05 | 1,56318296 | 0,91971282 | 0,3577229 |
| VC0908 | 1,20619311 | 8,08410988 | 1,16592107 | 7,8885E-06 | 1,16592107 | 0,91970923 | 0,35772478 |
| VCA0366 | 1,27299432 | 3,21999533 | 1,19163848 | 3,1421E-06 | 1,19163848 | 0,91907103 | 0,35805845 |
| VC0168 | 1,51964336 | 96,063524 | 1,40409681 | 9,374E-05 | 1,40409681 | 0,91777068 | 0,35873899 |
| VC2766 | 1,21068651 | 63,2066922 | 1,14327239 | 6,1678E-05 | 1,14327239 | 0,91747968 | 0,3588914 |
| VCA0560 | -1,0732587 | -4,4553376 | -1,13432 | -4,348E-06 | -1,13432 | -0,9171683 | 0,3590545 |
| VC2694 | -1,0550291 | -4,0412476 | -1,1490024 | -3,943E-06 | -1,1490024 | -0,9168595 | 0,35921629 |
| VCA0248 | -1,2125742 | -8,2796729 | -1,3202215 | -8,079E-06 | -1,3202215 | -0,9167465 | 0,35927552 |
| flgK | -1,1024572 | -12,457808 | -1,1409283 | -1,216E-05 | -1,1409283 | -0,9166635 | 0,35931905 |
| VC1054 | 1,18781947 | 5,36666176 | 1,11418132 | 5,2368E-06 | 1,11418132 | 0,91650646 | 0,35940136 |
| VC0343 | 1,16509221 | 5,23821776 | 1,11272348 | 5,1115E-06 | 1,11272348 | 0,91569049 | 0,35982929 |

| | | | | | | | |
|---|---|---|---|---|---|---|---|
| VC1575 | 1,42501058 | 31,8005966 | 1,37468544 | 3,1031E-05 | 1,37468544 | 0,91428404 | 0,36056765 |
| VC2518 | 1,23395286 | 9,15223819 | 1,21537615 | 8,9308E-06 | 1,21537615 | 0,91364457 | 0,36090369 |
| VCA0643 | 2,04548893 | 0,88599724 | 1,91542998 | 8,6456E-07 | 1,91542998 | 0,91356278 | 0,36094667 |
| flhF | -1,0745738 | -7,8346249 | -1,1306206 | -7,645E-06 | -1,1306206 | -0,9132234 | 0,3611251 |
| maf (NC_002505 444542..445200) | -1,0997256 | -13,100448 | -1,1163134 | -1,278E-05 | -1,1163134 | -0,9127755 | 0,36136069 |
| VC0284 | 1,61267637 | 1,38300898 | 1,50157856 | 1,3496E-06 | 1,50157856 | 0,91208706 | 0,36172293 |
| VC1205 | 2,98079391 | 3,93895421 | 2,42620246 | 3,8437E-06 | 2,42620246 | 0,91049433 | 0,36256193 |
| VC0523 | -1,0678981 | -5,4038146 | -1,1156743 | -5,273E-06 | -1,1156743 | -0,9086445 | 0,36353785 |
| VCA0536 | -1,3633991 | -62,624234 | -1,3613039 | -6,111E-05 | -1,3613039 | -0,9069341 | 0,36444167 |
| VCA0183 | -1,1469479 | -2,5795759 | -1,2378439 | -2,517E-06 | -1,2378439 | -0,906919 | 0,36444968 |
| VC0161 | 1,39689374 | 4,16418429 | 1,29997667 | 4,0634E-06 | 1,29997667 | 0,90671243 | 0,36455894 |
| VCA0900 | -1,1651208 | -2,6380948 | -1,2317894 | -2,574E-06 | -1,2317894 | -0,9066272 | 0,364604 |
| rplE | 1,35491733 | 89,5222286 | 1,12800705 | 8,7357E-05 | 1,12800705 | 0,90648048 | 0,36468166 |
| sdhC | 2,68753349 | 69,0894933 | 2,44254972 | 6,7418E-05 | 2,44254972 | 0,90521442 | 0,36535187 |
| trmD | 1,56246861 | 126,367566 | 1,23296147 | 0,00012331 | 1,23296147 | 0,90508511 | 0,36542034 |
| aroK | 1,28961033 | 22,9016825 | 1,23824664 | 2,2348E-05 | 1,23824664 | 0,90492055 | 0,36550752 |
| VC0715 | 1,38131822 | 25,2882411 | 1,39406211 | 2,4676E-05 | 1,39406211 | 0,90482589 | 0,36555769 |
| VC1417 | 1,2980814 | 2,83819134 | 1,21655439 | 2,7695E-06 | 1,21655439 | 0,90312758 | 0,36645822 |
| VC1103 | 1,43126076 | 3,72807221 | 1,28445063 | 3,6379E-06 | 1,28445063 | 0,9007007 | 0,36774752 |
| VCA0843 | 1,58765112 | 9,7616711 | 1,4352512 | 9,5255E-06 | 1,4352512 | 0,89965944 | 0,36830158 |
| sbcB | 1,39495361 | 2,70091943 | 1,3014648 | 2,6356E-06 | 1,3014648 | 0,89952656 | 0,36837229 |
| VC0470 | 1,37576859 | 2,14857869 | 1,28688673 | 2,0966E-06 | 1,28688673 | 0,89923045 | 0,36852996 |
| VC1854 | -1,4767564 | -765,94822 | -1,3165796 | -0,0007474 | -1,3165796 | -0,8977741 | 0,36930607 |
| VC0035 | 1,25955846 | 3,41547153 | 1,17029241 | 3,3328E-06 | 1,17029241 | 0,89666274 | 0,36989894 |
| VC0746 | -1,2703567 | -9,1202934 | -1,2682371 | -8,9E-06 | -1,2682371 | -0,8961096 | 0,3701943 |
| VC2296 | 1,20129303 | 5,09082629 | 1,12796014 | 4,9677E-06 | 1,12796014 | 0,8956736 | 0,37042715 |
| VCA0101 | 1,22202784 | 4,794163 | 1,1270116 | 4,6782E-06 | 1,1270116 | 0,89489069 | 0,37084556 |
| VC1651 | 1,28989141 | 2,62055131 | 1,22678909 | 2,5572E-06 | 1,22678909 | 0,89481087 | 0,37088823 |
| tRNA-Gln-3 | -1,4570069 | -5,5826655 | -1,4996377 | -5,448E-06 | -1,4996377 | -0,8945196 | 0,37104402 |
| VC1860 | 1,20653863 | 25,3583356 | 1,2276588 | 2,4745E-05 | 1,2276588 | 0,89406003 | 0,37128983 |
| VC1393 | -1,0963129 | -3,556506 | -1,1617524 | -3,47E-06 | -1,1617524 | -0,8935108 | 0,37158374 |
| VC1437 | -1,2313913 | -12,411947 | -1,2667929 | -1,211E-05 | -1,2667929 | -0,8933156 | 0,37168819 |
| VC1979 | 1,40157186 | 2,17131727 | 1,27836444 | 2,1188E-06 | 1,27836444 | 0,89211217 | 0,37233285 |
| VCA0278 | 1,613779 | 43,8446777 | 1,54925102 | 4,2784E-05 | 1,54925102 | 0,89191379 | 0,37243921 |
| VC1798 | 1,33745034 | 2,22721098 | 1,27000033 | 2,1733E-06 | 1,27000033 | 0,89148186 | 0,37267076 |
| tnaA | 2,07919848 | 3,48203865 | 1,76159069 | 3,3978E-06 | 1,76159069 | 0,89129597 | 0,37277046 |
| VCA0315 | -1,3374597 | -3,7720668 | -1,4347419 | -3,681E-06 | -1,4347419 | -0,891294 | 0,37277154 |
| VCA1104 | -1,2783844 | -7,1315056 | -1,3772439 | -6,959E-06 | -1,3772439 | -0,8880309 | 0,37452416 |
| ushA | -1,2544423 | -87,920698 | -1,3686441 | -8,579E-05 | -1,3686441 | -0,8878765 | 0,37460727 |
| VC0225 | 1,27879384 | 2,9087664 | 1,19804887 | 2,8384E-06 | 1,19804887 | 0,88671898 | 0,37523025 |
| VC1452 | -1,1191788 | -6,9983 | -1,1190709 | -6,829E-06 | -1,1190709 | -0,8863575 | 0,37542497 |
| lysS | 1,2680681 | 17,5009193 | 1,24739969 | 1,7078E-05 | 1,24739969 | 0,8861558 | 0,37553362 |
| VCA0674 | 1,51766387 | 18,9201609 | 1,49910406 | 1,8462E-05 | 1,49910406 | 0,88523622 | 0,37602928 |
| VC1071 | 1,54348884 | 1,44303433 | 1,51613033 | 1,4081E-06 | 1,51613033 | 0,88412213 | 0,37663032 |
| fis | 1,38858858 | 38,6391985 | 1,42595849 | 3,7704E-05 | 1,42595849 | 0,88362167 | 0,37690055 |

| | | | | | | | |
|---|---|---|---|---|---|---|---|
| VCA0218 | 1,61782674 | 2,62899653 | 1,39072165 | 2,5654E-06 | 1,39072165 | 0,88291874 | 0,37728022 |
| VC2653 | 1,29060645 | 56,1191397 | 1,27946418 | 5,4762E-05 | 1,27946418 | 0,88246795 | 0,37752387 |
| VC0223 | -1,0869525 | -3,5904372 | -1,1556373 | -3,504E-06 | -1,1556373 | -0,8818782 | 0,37784273 |
| VC0541 | 1,6338817 | 12,8758916 | 1,64148868 | 1,2564E-05 | 1,64148868 | 0,88157545 | 0,3780065 |
| tmk | 1,23991138 | 2,97295543 | 1,19043941 | 2,901E-06 | 1,19043941 | 0,88058498 | 0,37854253 |
| ulaD | -1,527143 | -2,5870343 | -1,6754377 | -2,524E-06 | -1,6754377 | -0,8797647 | 0,37898683 |
| VC1716 | 1,2849667 | 5,02071373 | 1,20560999 | 4,8993E-06 | 1,20560999 | 0,87966587 | 0,37904038 |
| VC0567 | -1,0137422 | -30,227782 | -1,0821423 | -2,95E-05 | -1,0821423 | -0,87932 | 0,37922782 |
| VC0849 | 1,23966161 | 10,2211481 | 1,22047393 | 9,9739E-06 | 1,22047393 | 0,87926329 | 0,37925859 |
| VCA0565 | -1,1356944 | -9,1170379 | -1,1311384 | -8,896E-06 | -1,1311384 | -0,8788503 | 0,37948247 |
| VCA0582 | 1,26206422 | 4,79035603 | 1,2072661 | 4,6745E-06 | 1,2072661 | 0,87811355 | 0,37988214 |
| VC1510 | -1,6524177 | -38,523537 | -1,6878461 | -3,759E-05 | -1,6878461 | -0,8776689 | 0,38012344 |
| VC0248 | 1,42408353 | 163,294777 | 1,3046684 | 0,00015934 | 1,3046684 | 0,87681629 | 0,38058646 |
| VC0451 | -1,0803537 | -5,3521166 | -1,100566 | -5,223E-06 | -1,100566 | -0,8767841 | 0,38060397 |
| VC1672 | -1,1862899 | -2,1243821 | -1,2739564 | -2,073E-06 | -1,2739564 | -0,8762509 | 0,38089369 |
| VC1285 | 1,25993078 | 3,29748221 | 1,16735917 | 3,2177E-06 | 1,16735917 | 0,87400864 | 0,38211358 |
| VC1962 | 1,24945737 | 5,93140851 | 1,17762286 | 5,7879E-06 | 1,17762286 | 0,87203951 | 0,38318686 |
| VCA0986 | -1,3574836 | -10,993738 | -1,4273817 | -1,073E-05 | -1,4273817 | -0,8716985 | 0,38337294 |
| gltD (NC_002505 2542113..2543569) | 1,56539916 | 5,47322849 | 1,5155686 | 5,3408E-06 | 1,5155686 | 0,86862049 | 0,3850548 |
| VC1278 | -1,1043458 | -3,2145045 | -1,1707328 | -3,137E-06 | -1,1707328 | -0,8684214 | 0,38516373 |
| rpoZ | 1,13236027 | 19,4463444 | 1,14292693 | 1,8976E-05 | 1,14292693 | 0,86820907 | 0,38527993 |
| proA | 1,26613388 | 5,11295995 | 1,20687379 | 4,9893E-06 | 1,20687379 | 0,86606439 | 0,3864549 |
| flhB | -1,1630039 | -5,926856 | -1,2373404 | -5,783E-06 | -1,2373404 | -0,8651563 | 0,38695303 |
| VC0270 | 1,13236293 | 14,7891328 | 1,13017695 | 1,4431E-05 | 1,13017695 | 0,86425232 | 0,38744934 |
| VC2402 | -1,126369 | -3,3421351 | -1,1830556 | -3,261E-06 | -1,1830556 | -0,864062 | 0,38755386 |
| VC0256 | -1,3558672 | -18,611719 | -1,3102625 | -1,816E-05 | -1,3102625 | -0,8634353 | 0,38789823 |
| dapF | 1,12012866 | 6,39708169 | 1,09419714 | 6,2423E-06 | 1,09419714 | 0,86316403 | 0,38804732 |
| VCA0629 | -1,2696328 | -1,5468991 | -1,3823682 | -1,509E-06 | -1,3823682 | -0,8630364 | 0,38811751 |
| VC2107 | 1,2761001 | 9,69401794 | 1,2504577 | 9,4595E-06 | 1,2504577 | 0,86298892 | 0,38814361 |
| VC2625 | 1,19344684 | 4,23577894 | 1,12445161 | 4,1333E-06 | 1,12445161 | 0,86280329 | 0,38824566 |
| grxA | 1,2240142 | 34,8508644 | 1,23551446 | 3,4008E-05 | 1,23551446 | 0,86225938 | 0,38854484 |
| panC | 1,20098896 | 12,2375134 | 1,18347823 | 1,1941E-05 | 1,18347823 | 0,86192635 | 0,38872809 |
| VC1308 | 1,2196808 | 3,61571175 | 1,14696091 | 3,5282E-06 | 1,14696091 | 0,86169188 | 0,38885713 |
| VC0882 | 1,25860479 | 8,29554199 | 1,1960045 | 8,0949E-06 | 1,1960045 | 0,86058533 | 0,38946652 |
| VC2706 | 1,65668265 | 12,0750592 | 1,61904696 | 1,1783E-05 | 1,61904696 | 0,86020195 | 0,38967779 |
| tRNA-Met-4 | -1,3164737 | -8,649703 | -1,3380839 | -8,44E-06 | -1,3380839 | -0,8596863 | 0,38996202 |
| VCA0112 | 1,60019419 | 1,37564707 | 1,43606297 | 1,3424E-06 | 1,43606297 | 0,85949891 | 0,39006536 |
| VCA0867 | 1,85846808 | 182,193068 | 1,8777929 | 0,00017779 | 1,8777929 | 0,85867514 | 0,39051981 |
| rplI | 1,29399794 | 77,4674666 | 1,26550621 | 7,5593E-05 | 1,26550621 | 0,85832259 | 0,39071441 |
| VC2558 | 1,86452144 | 5,47686251 | 1,80012086 | 5,3444E-06 | 1,80012086 | 0,85807005 | 0,39085385 |
| VCA0743 | -1,3513328 | -21,476542 | -1,3631355 | -2,096E-05 | -1,3631355 | -0,857155 | 0,39135928 |
| VC0667 | 1,25366129 | 11,4829174 | 1,21111654 | 1,1205E-05 | 1,21111654 | 0,85694848 | 0,39147341 |
| VC1150 | 1,58003054 | 20,154279 | 1,46689539 | 1,9667E-05 | 1,46689539 | 0,85620412 | 0,39188494 |
| VC0255 | 1,20034326 | 62,2072307 | 1,28069191 | 6,0702E-05 | 1,28069191 | 0,85598808 | 0,39200441 |
| VC0203 | 1,56357537 | 1,55879976 | 1,42392876 | 1,5211E-06 | 1,42392876 | 0,85594538 | 0,39202802 |

| | | | | | | | |
|---|---|---|---|---|---|---|---|
| tolB | 1,46520579 | 116,525982 | 1,31237132 | 0,00011371 | 1,31237132 | 0,85530185 | 0,39238409 |
| VC1474 | 1,45470805 | 1,64196494 | 1,34876602 | 1,6022E-06 | 1,34876602 | 0,85524584 | 0,3924151 |
| VC2022 | 1,24751661 | 18,7234515 | 1,23390994 | 1,8271E-05 | 1,23390994 | 0,85441399 | 0,3928757 |
| pheT | 1,28008583 | 7,20703247 | 1,2103147 | 7,0327E-06 | 1,2103147 | 0,85301855 | 0,39364904 |
| VC1889 | 1,29453675 | 2,39356758 | 1,22498845 | 2,3357E-06 | 1,22498845 | 0,8521229 | 0,39414592 |
| aroE | 1,23900321 | 6,41585718 | 1,18638932 | 6,2606E-06 | 1,18638932 | 0,85167594 | 0,39439402 |
| VC1086 | -1,2776124 | -6,135244 | -1,3379712 | -5,987E-06 | -1,3379712 | -0,851531 | 0,39447449 |
| VCA0195 | -1,2194168 | -9,6915877 | -1,3563604 | -9,457E-06 | -1,3563604 | -0,8514408 | 0,3945246 |
| VCA0518 | -1,8683808 | -277,57304 | -2,252276 | -0,0002709 | -2,252276 | -0,8508783 | 0,394837 |
| VC0431 | -1,0920828 | -10,682885 | -1,0546082 | -1,042E-05 | -1,0546082 | -0,8503729 | 0,39511784 |
| fliI | -1,0715953 | -3,77267 | -1,1364861 | -3,681E-06 | -1,1364861 | -0,8503262 | 0,3951438 |
| VC1617 | 2,09432741 | 14,3903892 | 1,84790971 | 1,4042E-05 | 1,84790971 | 0,85030662 | 0,3951547 |
| VCA0765 | 1,29587592 | 7,23770241 | 1,20446687 | 7,0626E-06 | 1,20446687 | 0,85029879 | 0,39515904 |
| VC2119 | -1,0759061 | -5,513304 | -1,1301649 | -5,38E-06 | -1,1301649 | -0,8500782 | 0,39528162 |
| VC0827 | -1,2511434 | -274,95367 | -1,2975704 | -0,0002683 | -1,2975704 | -0,8491952 | 0,39577276 |
| VC1518 | -1,2089831 | -6,0729691 | -1,2970656 | -5,926E-06 | -1,2970656 | -0,8483639 | 0,3962354 |
| VCA0476 | 1,30646943 | 11,0366789 | 1,27523003 | 1,077E-05 | 1,27523003 | 0,84804175 | 0,39641477 |
| ribA | 1,35299834 | 26,2272566 | 1,35546757 | 2,5593E-05 | 1,35546757 | 0,84770891 | 0,39660015 |
| VC1490 | 1,24481004 | 9,28173769 | 1,17909914 | 9,0572E-06 | 1,17909914 | 0,84764377 | 0,39663644 |
| VC0692 | 1,20097811 | 3,55049654 | 1,1445666 | 3,4646E-06 | 1,1445666 | 0,84737373 | 0,39678688 |
| VC0784 | 2,37877366 | 10,7704999 | 2,00716686 | 1,051E-05 | 2,00716686 | 0,84710407 | 0,39693717 |
| VCA0340 | 1,2429655 | 2,98020744 | 1,17402831 | 2,9081E-06 | 1,17402831 | 0,84598994 | 0,39755838 |
| VCA0982 | 1,97329485 | 3,17579831 | 1,65030994 | 3,099E-06 | 1,65030994 | 0,84581676 | 0,39765502 |
| VC0617 | -2,1305573 | -7,2645978 | -2,2610342 | -7,089E-06 | -2,2610342 | -0,84573 | 0,39770345 |
| uvrA | 1,26033966 | 2,84731976 | 1,18214181 | 2,7784E-06 | 1,18214181 | 0,84439514 | 0,39844867 |
| VC1029 | 1,33922411 | 2,17885293 | 1,24466497 | 2,1261E-06 | 1,24466497 | 0,84408631 | 0,3986212 |
| VC0881 | 1,3000683 | 2,59743385 | 1,20095994 | 2,5346E-06 | 1,20095994 | 0,84349905 | 0,39894943 |
| VC0649 | -1,1304369 | -4,246649 | -1,1864721 | -4,144E-06 | -1,1864721 | -0,8434363 | 0,39898451 |
| tRNA-Arg-2 | -1,578537 | -1,6103368 | -1,5911404 | -1,571E-06 | -1,5911404 | -0,8428636 | 0,39930476 |
| VC1936 | 1,49975225 | 1,78990382 | 1,30450249 | 1,7466E-06 | 1,30450249 | 0,84233292 | 0,39960167 |
| VC0981 | 1,06938477 | 6,77842393 | 1,07960413 | 6,6144E-06 | 1,07960413 | 0,84190545 | 0,39984092 |
| VC1445 | 1,111781 | 10,7833867 | 1,1278599 | 1,0523E-05 | 1,1278599 | 0,8418796 | 0,39985539 |
| rplC | 1,53995054 | 194,39359 | 1,4169466 | 0,00018969 | 1,4169466 | 0,84185733 | 0,39986784 |
| VC2090 | 2,18188058 | 78,7392608 | 2,06868106 | 7,6834E-05 | 2,06868106 | 0,84182326 | 0,39988694 |
| VCA0666 | -1,1132213 | -2,6272248 | -1,1973051 | -2,564E-06 | -1,1973051 | -0,8412719 | 0,40019567 |
| VC1561 | 1,28468339 | 2,60212481 | 1,19806791 | 2,5392E-06 | 1,19806791 | 0,83871475 | 0,40162944 |
| VC1371 | -1,1145282 | -3,5880664 | -1,1759946 | -3,501E-06 | -1,1759946 | -0,8378073 | 0,402139 |
| VC1235 | 2,33365027 | 14,3343223 | 2,40665715 | 1,3988E-05 | 2,40665715 | 0,83700984 | 0,40258709 |
| VC1123 | -1,2065989 | -5,120055 | -1,2669715 | -4,996E-06 | -1,2669715 | -0,8369635 | 0,40261309 |
| rdgC | 1,32149038 | 2,57435599 | 1,1993362 | 2,5121E-06 | 1,1993362 | 0,83665274 | 0,40278783 |
| VC1496 | 1,12013165 | 32,1405155 | 1,1668158 | 3,1363E-05 | 1,1668158 | 0,83664848 | 0,40279023 |
| VC1616 | 1,88536808 | 18,4442907 | 1,6619443 | 1,7998E-05 | 1,6619443 | 0,83660741 | 0,40281334 |
| VC1140 | -1,0806394 | -4,0450178 | -1,1223925 | -3,947E-06 | -1,1223925 | -0,8365517 | 0,40284464 |
| alr (NC_002505 1393705..1394969) | -1,0964089 | -3,3071898 | -1,1672592 | -3,227E-06 | -1,1672592 | -0,8365202 | 0,40286234 |
| araD | -1,2721823 | -1,1487249 | -1,5090702 | -1,121E-06 | -1,5090702 | -0,8361837 | 0,4030516 |

| | | | | | | | |
|---|---|---|---|---|---|---|---|
| VC0291 | 1,34465869 | 18,8890202 | 1,35772338 | 1,8432E-05 | 1,35772338 | 0,83605118 | 0,40312617 |
| VC0513 | 1,63930135 | 5,2878535 | 1,58845979 | 5,1599E-06 | 1,58845979 | 0,835746 | 0,40329786 |
| VC0460 | 1,23702866 | 2,95350722 | 1,17098612 | 2,8821E-06 | 1,17098612 | 0,83538283 | 0,40350223 |
| guaA | 1,37569533 | 23,4761911 | 1,418793 | 2,2908E-05 | 1,418793 | 0,83447354 | 0,40401424 |
| VC2724 | -1,0959838 | -3,4456833 | -1,1439922 | -3,362E-06 | -1,1439922 | -0,8332238 | 0,40471856 |
| pgi | -1,5323073 | -52,255703 | -1,4376825 | -5,099E-05 | -1,4376825 | -0,8327788 | 0,40496951 |
| VC0714 | 1,43233865 | 40,4624969 | 1,49941348 | 3,9484E-05 | 1,49941348 | 0,83234832 | 0,40521242 |
| VC2244 | -1,0846459 | -8,3088524 | -1,1056618 | -8,108E-06 | -1,1056618 | -0,8323339 | 0,40522052 |
| rpsP | 1,3993814 | 85,1603472 | 1,19777968 | 8,31E-05 | 1,19777968 | 0,82921336 | 0,40698372 |
| VC0792 | 5,93616744 | 34,2323529 | 5,74218292 | 3,3404E-05 | 5,74218292 | 0,82888467 | 0,40716973 |
| VC2051 | -1,4078092 | -30,138831 | -1,3555371 | -2,941E-05 | -1,3555371 | -0,8284588 | 0,40741075 |
| bcp | 1,1230369 | 25,8523623 | 1,14203912 | 2,5227E-05 | 1,14203912 | 0,82814096 | 0,40759072 |
| VC2152 | 1,30101245 | 2,61454027 | 1,22604793 | 2,5513E-06 | 1,22604793 | 0,82772886 | 0,40782411 |
| VC0781 | -1,0656305 | -3,7429309 | -1,1296218 | -3,652E-06 | -1,1296218 | -0,8267246 | 0,40839317 |
| VC1563 | 1,36503025 | 1,84564147 | 1,28160444 | 1,801E-06 | 1,28160444 | 0,82667628 | 0,40842061 |
| VCA0323 | 1,1379877 | 13,5047081 | 1,13691264 | 1,3178E-05 | 1,13691264 | 0,82592402 | 0,40884722 |
| VC1436 | -1,2341862 | -27,340152 | -1,1794618 | -2,668E-05 | -1,1794618 | -0,8252633 | 0,40922217 |
| VC0997 | 1,09162111 | 19,9061119 | 1,11468134 | 1,9425E-05 | 1,11468134 | 0,82512911 | 0,40929835 |
| VC1566 | 1,46629466 | 1,37614593 | 1,39479594 | 1,3429E-06 | 1,39479594 | 0,82498464 | 0,40938034 |
| VC1434 | -1,1405029 | -11,275104 | -1,1254924 | -1,1E-05 | -1,1254924 | -0,8243103 | 0,40976329 |
| VC1448 | 1,71033508 | 1,94341223 | 1,70599706 | 1,8964E-06 | 1,70599706 | 0,8204013 | 0,41198742 |
| VCA0186 | 1,63722074 | 22,710245 | 1,48240475 | 2,2161E-05 | 1,48240475 | 0,81978901 | 0,41233645 |
| prfA | 1,14968465 | 6,40794536 | 1,11234492 | 6,2529E-06 | 1,11234492 | 0,81960793 | 0,41243968 |
| ubiE | 1,12815032 | 5,00829925 | 1,0934793 | 4,8871E-06 | 1,0934793 | 0,81910781 | 0,41272494 |
| VC1938 | 1,38934283 | 2,92786623 | 1,26702253 | 2,857E-06 | 1,26702253 | 0,81882285 | 0,41288753 |
| VCA0329 | -1,0649936 | -3,252611 | -1,1474501 | -3,174E-06 | -1,1474501 | -0,8185454 | 0,41304584 |
| VC2532 | -1,0618901 | -5,8641058 | -1,144302 | -5,722E-06 | -1,144302 | -0,8183628 | 0,41315009 |
| VC1364 | 1,13303878 | 6,09576169 | 1,11393526 | 5,9483E-06 | 1,11393526 | 0,81830087 | 0,41318544 |
| VC0548 | -1,0657434 | -129,51768 | -1,1930915 | -0,0001264 | -1,1930915 | -0,8180404 | 0,41333413 |
| VC2271 | -1,0621734 | -5,6248148 | -1,1278234 | -5,489E-06 | -1,1278234 | -0,8180386 | 0,41333515 |
| VCA0245 | -1,2237388 | -1,4610277 | -1,4805 | -1,426E-06 | -1,4805 | -0,8178645 | 0,41343459 |
| VC1353 | -1,153956 | -4,3476138 | -1,2528706 | -4,242E-06 | -1,2528706 | -0,8174031 | 0,41369816 |
| VCA0996 | 1,22323287 | 2,9154959 | 1,16484524 | 2,845E-06 | 1,16484524 | 0,81610436 | 0,41444049 |
| clpA | 1,24474526 | 23,8198127 | 1,23546363 | 2,3244E-05 | 1,23546363 | 0,81502829 | 0,41505617 |
| cysM | 1,31065227 | 2,40058018 | 1,2146768 | 2,3425E-06 | 1,2146768 | 0,81470229 | 0,41524277 |
| VC0049 | 1,41740854 | 45,1189365 | 1,38276886 | 4,4027E-05 | 1,38276886 | 0,81398178 | 0,41565544 |
| VC0445 | 1,22645245 | 12,9386813 | 1,18795913 | 1,2626E-05 | 1,18795913 | 0,8122601 | 0,41664244 |
| VC2416 | -1,2428437 | -20,164162 | -1,2957119 | -1,968E-05 | -1,2957119 | -0,8103815 | 0,41772101 |
| VC1919 | -1,4489675 | -828,17499 | -1,3152266 | -0,0008081 | -1,3152266 | -0,8096916 | 0,41811751 |
| VCA0363 | 1,50248625 | 1,3463846 | 1,38705329 | 1,3138E-06 | 1,38705329 | 0,80928301 | 0,41835241 |
| VC1002 | 1,15496356 | 4,44322381 | 1,10302159 | 4,3357E-06 | 1,10302159 | 0,80809349 | 0,4190368 |
| VC1764 | -1,3635635 | -19,486938 | -1,3151488 | -1,902E-05 | -1,3151488 | -0,8069245 | 0,41970999 |
| VCA0566 | 1,10928709 | 19,3986488 | 1,13649108 | 1,8929E-05 | 1,13649108 | 0,80674873 | 0,41981131 |
| VCA1036 | 1,36597713 | 3,26076439 | 1,23625825 | 3,1819E-06 | 1,23625825 | 0,80662798 | 0,41988088 |
| VC2039 | 1,09911672 | 7,24848853 | 1,10908858 | 7,0731E-06 | 1,10908858 | 0,80591162 | 0,42029384 |

| | | | | | | | |
|---|---|---|---|---|---|---|---|
| VC1865 | -1,6713551 | -92,730032 | -1,5501288 | -9,049E-05 | -1,5501288 | -0,805803 | 0,42035646 |
| VCA0478 | 1,22742379 | 10,1336676 | 1,20206492 | 9,8885E-06 | 1,20206492 | 0,80548686 | 0,42053881 |
| VC2078 | 1,40416133 | 20,3253432 | 1,41325973 | 1,9834E-05 | 1,41325973 | 0,80484896 | 0,42090688 |
| VC1873 | -1,2993755 | -1,3226499 | -1,3901728 | -1,291E-06 | -1,3901728 | -0,8048182 | 0,42092463 |
| VC1620 | 1,17048669 | 8,78948595 | 1,17407601 | 8,5769E-06 | 1,17407601 | 0,8032256 | 0,42184439 |
| VCA0192 | 1,28332183 | 8,60157299 | 1,26345971 | 8,3935E-06 | 1,26345971 | 0,8030206 | 0,42196287 |
| VCA0840 | 1,2425242 | 24,9517668 | 1,27853601 | 2,4348E-05 | 1,27853601 | 0,80259715 | 0,42220766 |
| sucA | -1,4913784 | -40,377998 | -1,4828066 | -3,94E-05 | -1,4828066 | -0,801522 | 0,42282957 |
| VC0232 | 1,155183 | 4,09829641 | 1,11017666 | 3,9992E-06 | 1,11017666 | 0,80122786 | 0,42299977 |
| VC2040 | 1,1168004 | 19,0662225 | 1,12420331 | 1,8605E-05 | 1,12420331 | 0,80094698 | 0,42316237 |
| VC2441 | 1,15663992 | 13,7069152 | 1,14814313 | 1,3375E-05 | 1,14814313 | 0,79973169 | 0,42386632 |
| VC1565 | 1,33125802 | 1,68404559 | 1,3029044 | 1,6433E-06 | 1,3029044 | 0,79955693 | 0,42396758 |
| VC1595 | -1,3937399 | -74,625973 | -1,4659254 | -7,282E-05 | -1,4659254 | -0,7993047 | 0,42411378 |
| VC2314 | -1,2701774 | -1,3905904 | -1,3610194 | -1,357E-06 | -1,3610194 | -0,7986872 | 0,42447187 |
| VC2362 | 1,41714496 | 12,8807079 | 1,40004781 | 1,2569E-05 | 1,40004781 | 0,79867211 | 0,42448062 |
| VCA0902 | -1,0689799 | -2,6554709 | -1,1779038 | -2,591E-06 | -1,1779038 | -0,7979403 | 0,42490517 |
| VC1696 | -1,2204562 | -16,689345 | -1,3727283 | -1,629E-05 | -1,3727283 | -0,7972025 | 0,42533347 |
| VC0914 | -1,0680098 | -3,094893 | -1,1627337 | -3,02E-06 | -1,1627337 | -0,7969407 | 0,42548548 |
| VC2722 | -1,0738297 | -3,1616825 | -1,1434037 | -3,085E-06 | -1,1434037 | -0,7966233 | 0,42566984 |
| VC0573 | 1,61006356 | 105,612933 | 1,53491722 | 0,00010306 | 1,53491722 | 0,79616235 | 0,42593773 |
| VCA0492 | 1,22540899 | 3,33758266 | 1,18895712 | 3,2568E-06 | 1,18895712 | 0,79473597 | 0,42676714 |
| VC1272 | 1,384139 | 4,19355312 | 1,30281728 | 4,0921E-06 | 1,30281728 | 0,79344222 | 0,42752026 |
| cysN | 1,98326982 | 5,11068097 | 1,93994104 | 4,987E-06 | 1,93994104 | 0,79234276 | 0,42816089 |
| VC1516 | -1,2267729 | -17,602333 | -1,2858952 | -1,718E-05 | -1,2858952 | -0,7923372 | 0,42816416 |
| VC1713 | 1,28484166 | 3,20285615 | 1,22002338 | 3,1254E-06 | 1,22002338 | 0,79193322 | 0,42839965 |
| VC2144 | -1,2077685 | -4,606702 | -1,2478467 | -4,495E-06 | -1,2478467 | -0,7905529 | 0,42920498 |
| VC2624 | 1,2138358 | 3,51574976 | 1,12565884 | 3,4307E-06 | 1,12565884 | 0,78963368 | 0,42974176 |
| VC1152 | 1,14101747 | 8,17452803 | 1,13837865 | 7,9768E-06 | 1,13837865 | 0,78935842 | 0,42990259 |
| VCA0378 | 1,95139555 | 0,73166194 | 1,78992817 | 7,1396E-07 | 1,78992817 | 0,78833978 | 0,43049802 |
| VC1480 | 1,15154855 | 24,5713993 | 1,15719563 | 2,3977E-05 | 1,15719563 | 0,78783955 | 0,43079061 |
| VC1068 | 1,37952553 | 1,85063484 | 1,30693857 | 1,8059E-06 | 1,30693857 | 0,78617791 | 0,4317633 |
| VCA0953 | 1,35175436 | 7,97487345 | 1,29971736 | 7,782E-06 | 1,29971736 | 0,7856232 | 0,43208832 |
| VC1621 | 1,39793939 | 21,1908326 | 1,34939901 | 2,0678E-05 | 1,34939901 | 0,7855497 | 0,43213138 |
| VC1676 | 1,28420292 | 4,10543882 | 1,22159557 | 4,0061E-06 | 1,22159557 | 0,78543129 | 0,43220077 |
| oadA-2 | 5,0745192 | 60,0580729 | 4,85355961 | 5,8605E-05 | 4,85355961 | 0,78540169 | 0,43221815 |
| qrr2 | -1,1479403 | -2,4407351 | -1,2174784 | -2,382E-06 | -1,2174784 | -0,7853705 | 0,43223643 |
| ubiA | -1,0650783 | -2,9165076 | -1,1514468 | -2,846E-06 | -1,1514468 | -0,7848042 | 0,43256843 |
| VC2501 | 1,20992303 | 6,96282055 | 1,14387901 | 6,7944E-06 | 1,14387901 | 0,78459481 | 0,4326912 |
| 5Sf | -1,0814986 | -9260,6815 | -1,0802818 | -0,0090367 | -1,0802818 | -0,7845208 | 0,43273462 |
| VC1431 | 1,10364488 | 7,25275708 | 1,10984803 | 7,0773E-06 | 1,10984803 | 0,78408837 | 0,43298829 |
| VC1215 | -1,0896886 | -10,066517 | -1,1043484 | -9,823E-06 | -1,1043484 | -0,7840239 | 0,43302615 |
| VC2727 | -1,0855511 | -4,4953654 | -1,14531 | -4,387E-06 | -1,14531 | -0,7836056 | 0,43327159 |
| VC2286 | 1,13241648 | 6,98312014 | 1,11834204 | 6,8142E-06 | 1,11834204 | 0,78270507 | 0,43380036 |
| VC0732 | 1,2163961 | 3,97929223 | 1,15701342 | 3,883E-06 | 1,15701342 | 0,78246349 | 0,43394227 |
| VC0816 | 1,47533549 | 1,51089934 | 1,31183495 | 1,4743E-06 | 1,31183495 | 0,78192228 | 0,43426028 |

| | | | | | | | |
|---|---|---|---|---|---|---|---|
| hemE | 1,10928036 | 5,3440808 | 1,07924147 | 5,2148E-06 | 1,07924147 | 0,78169225 | 0,43439549 |
| VC0800 | 5,09681895 | 139,524818 | 4,82421181 | 0,00013615 | 4,82421181 | 0,78161109 | 0,43444322 |
| cysS | 1,08273494 | 8,87957672 | 1,07462238 | 8,6648E-06 | 1,07462238 | 0,78147062 | 0,43452579 |
| VCA0243 | -1,2387187 | -1,2004176 | -1,4265245 | -1,171E-06 | -1,4265245 | -0,7802498 | 0,43524386 |
| VC0694 | -1,1023819 | -2,6389471 | -1,166273 | -2,575E-06 | -1,166273 | -0,7795318 | 0,43566653 |
| VC1268 | 1,26417696 | 2,78894507 | 1,20526153 | 2,7215E-06 | 1,20526153 | 0,7793598 | 0,43576783 |
| VC1649 | 1,54806517 | 76,6104382 | 1,61056663 | 7,4757E-05 | 1,61056663 | 0,77908538 | 0,43592946 |
| VCA1011 | -1,1851487 | -1,6918088 | -1,2714591 | -1,651E-06 | -1,2714591 | -0,7788208 | 0,43608533 |
| VC0259 | 1,09406115 | 21,3314522 | 1,1589561 | 2,0815E-05 | 1,1589561 | 0,77770263 | 0,43674439 |
| VC1513 | -1,4779742 | -31,46035 | -1,4893818 | -3,07E-05 | -1,4893818 | -0,7776036 | 0,43680277 |
| VC1584 | 1,5698741 | 1,22740655 | 1,39268781 | 1,1977E-06 | 1,39268781 | 0,77738535 | 0,43693148 |
| VC1365 | 1,44458539 | 4,70403211 | 1,39447889 | 4,5902E-06 | 1,39447889 | 0,77590804 | 0,43780333 |
| VC2453 | -1,1144746 | -6,1487813 | -1,2214292 | -6E-06 | -1,2214292 | -0,7755803 | 0,4379969 |
| VC1370 | 1,33480197 | 1,91762083 | 1,23331544 | 1,8712E-06 | 1,23331544 | 0,77524802 | 0,43819314 |
| VC0578 | 1,04393682 | 8,43347085 | 1,06432118 | 8,2295E-06 | 1,06432118 | 0,77503547 | 0,43831873 |
| VC1615 | 2,08006873 | 9,68599421 | 1,79292741 | 9,4517E-06 | 1,79292741 | 0,77482365 | 0,43844393 |
| potC | 1,2991631 | 8,78298268 | 1,26460443 | 8,5705E-06 | 1,26460443 | 0,77413095 | 0,43885339 |
| VC0797 | 5,05209317 | 159,678324 | 4,74698195 | 0,00015582 | 4,74698195 | 0,77353661 | 0,43920492 |
| VC1319 | 1,372209 | 104,646986 | 1,31750696 | 0,00010212 | 1,31750696 | 0,77304554 | 0,43949546 |
| VC1198 | 1,35977262 | 4,80962091 | 1,28571421 | 4,6933E-06 | 1,28571421 | 0,77276174 | 0,43966344 |
| VCA0225 | 1,18021145 | 5,72326839 | 1,12471758 | 5,5848E-06 | 1,12471758 | 0,77216667 | 0,44001575 |
| VC0241 | 1,18959987 | 68,5450431 | 1,15060921 | 6,6887E-05 | 1,15060921 | 0,77201667 | 0,44010461 |
| VC2661 | -1,0904064 | -4,13211 | -1,1226297 | -4,032E-06 | -1,1226297 | -0,7718319 | 0,440214 |
| VCA0427 | 1,29050153 | 12,6174342 | 1,25267898 | 1,2312E-05 | 1,25267898 | 0,77158132 | 0,44036247 |
| VC0616 | -2,1467499 | -5,5425076 | -2,2460727 | -5,408E-06 | -2,2460727 | -0,7708112 | 0,44081887 |
| VC1184 | 1,54369079 | 4,29835712 | 1,37655719 | 4,1944E-06 | 1,37655719 | 0,77061351 | 0,44093609 |
| VC2329 | 1,15396936 | 6,69880242 | 1,11800647 | 6,5367E-06 | 1,11800647 | 0,77053956 | 0,44097993 |
| VC2097 | 1,1537725 | 4,29608907 | 1,11646339 | 4,1922E-06 | 1,11646339 | 0,76923234 | 0,44175543 |
| VC2442 | 1,19335387 | 10,2417393 | 1,18682506 | 9,994E-06 | 1,18682506 | 0,76768893 | 0,44267205 |
| VC1792 | 1,50062053 | 1,29705937 | 1,35614036 | 1,2657E-06 | 1,35614036 | 0,76692254 | 0,44312761 |
| VC2236 | 1,25384558 | 2,47851444 | 1,17174713 | 2,4186E-06 | 1,17174713 | 0,76683125 | 0,44318188 |
| VC0846 | 1,34657574 | 1,61334255 | 1,27562295 | 1,5743E-06 | 1,27562295 | 0,76565459 | 0,44388189 |
| VC0794 | 4,7137795 | 133,021721 | 4,42271087 | 0,0001298 | 4,42271087 | 0,76531281 | 0,44408535 |
| VCA0283 | 1,17239491 | 20,8733162 | 1,17480112 | 2,0368E-05 | 1,17480112 | 0,76371053 | 0,4450398 |
| VC0289 | 1,40998891 | 3,08055313 | 1,2705974 | 3,006E-06 | 1,2705974 | 0,76302952 | 0,44544583 |
| VC2323 | 1,30389496 | 2,01478937 | 1,24552212 | 1,966E-06 | 1,24552212 | 0,76294501 | 0,44549622 |
| VC0889 | 1,30399593 | 7,8660348 | 1,23695009 | 7,6757E-06 | 1,23695009 | 0,76121394 | 0,44652933 |
| VC1872 | -1,1637956 | -1,7843899 | -1,2426808 | -1,741E-06 | -1,2426808 | -0,7610992 | 0,44659782 |
| cpxA | -1,0432597 | -3,9486813 | -1,1292548 | -3,853E-06 | -1,1292548 | -0,7610977 | 0,44659872 |
| VCA0029 | 2,25705709 | 53,9868974 | 2,14197393 | 5,2681E-05 | 2,14197393 | 0,76107447 | 0,44661264 |
| VC2316 | -1,2191151 | -1,3659729 | -1,3287543 | -1,333E-06 | -1,3287543 | -0,7606005 | 0,44689572 |
| fadB | 1,70285488 | 1,28323996 | 1,44559214 | 1,2522E-06 | 1,44559214 | 0,75906725 | 0,44781236 |
| rpmJ (NC_002505 940858..941024) | 1,34269389 | 5,35675573 | 1,27000008 | 5,2272E-06 | 1,27000008 | 0,75895733 | 0,44787812 |
| VC1808 | -1,179775 | -7,6161528 | -1,3124436 | -7,432E-06 | -1,3124436 | -0,7581083 | 0,44838621 |
| VCA0020 | 1,34192787 | 1,81466755 | 1,23571544 | 1,7708E-06 | 1,23571544 | 0,75761185 | 0,44868341 |

| | | | | | | | |
|---|---|---|---|---|---|---|---|
| VC1403 | 1,43480062 | 1,2538358 | 1,36002294 | 1,2235E-06 | 1,36002294 | 0,7575108 | 0,44874393 |
| VC1186 | -1,0641632 | -4,4551701 | -1,1474586 | -4,347E-06 | -1,1474586 | -0,7573688 | 0,44882894 |
| VC0795 | 4,53718952 | 168,068879 | 4,40528742 | 0,000164 | 4,40528742 | 0,75725166 | 0,44889916 |
| VC0668 | 1,39202043 | 2,2337293 | 1,24933633 | 2,1797E-06 | 1,24933633 | 0,75721136 | 0,44892328 |
| VCA0705 | -1,2449831 | -1,234528 | -1,3659545 | -1,205E-06 | -1,3659545 | -0,7568718 | 0,44912668 |
| VC0713 | 1,39804007 | 1106,72743 | 1,65866764 | 0,00107995 | 1,65866764 | 0,75670794 | 0,44922491 |
| VC0796 | 4,43161803 | 380,288887 | 4,43085037 | 0,00037109 | 4,43085037 | 0,75611449 | 0,44958061 |
| rpmG | 1,09440936 | 778,126843 | 1,35818059 | 0,0007593 | 1,35818059 | 0,75556375 | 0,44991085 |
| VC2745 | 1,31833044 | 1,75454101 | 1,24329689 | 1,7121E-06 | 1,24329689 | 0,75556011 | 0,44991301 |
| VC2668 | 1,31762231 | 6,88662363 | 1,2075136 | 6,72E-06 | 1,2075136 | 0,75482357 | 0,45035489 |
| VC0798 | 4,6899084 | 189,633707 | 4,37711831 | 0,00018505 | 4,37711831 | 0,75477271 | 0,45038542 |
| VCA0030 | 1,83774504 | 12,1620007 | 1,82354391 | 1,1868E-05 | 1,82354391 | 0,75370698 | 0,45102525 |
| VC0871 | -1,3527492 | -32,128738 | -1,2972344 | -3,135E-05 | -1,2972344 | -0,7522986 | 0,45187153 |
| gidB | 1,16590669 | 7,96241738 | 1,15522817 | 7,7698E-06 | 1,15522817 | 0,75209634 | 0,45199317 |
| VC1511 | -1,5406735 | -29,78586 | -1,562204 | -2,907E-05 | -1,562204 | -0,7520313 | 0,45203226 |
| VCA0715 | 1,27750603 | 2,07219158 | 1,20014101 | 2,0221E-06 | 1,20014101 | 0,75200512 | 0,45204801 |
| VC0801 | 4,70833461 | 77,9907342 | 4,26746816 | 7,6104E-05 | 4,26746816 | 0,75193403 | 0,45209078 |
| rpsU | -1,0923951 | -111,3344 | -1,2231504 | -0,0001086 | -1,2231504 | -0,7507695 | 0,45279143 |
| VC1252 | -1,0627882 | -4,9431326 | -1,1242687 | -4,824E-06 | -1,1242687 | -0,7492931 | 0,45368061 |
| VC1852 | -1,0775833 | -7,1299113 | -1,0817765 | -6,957E-06 | -1,0817765 | -0,7479806 | 0,45447186 |
| VCA0312 | -1,0383593 | -2,9386862 | -1,1354583 | -2,868E-06 | -1,1354583 | -0,7478241 | 0,45456628 |
| VC0440 | -1,1384931 | -8,5435649 | -1,177112 | -8,337E-06 | -1,177112 | -0,7472861 | 0,45489091 |
| VC0945 | 1,24425631 | 4,63689604 | 1,20634902 | 4,5247E-06 | 1,20634902 | 0,74692766 | 0,45510726 |
| rbn | 1,2390756 | 2,26536885 | 1,17869545 | 2,2106E-06 | 1,17869545 | 0,74660608 | 0,4553014 |
| VC0799 | 4,73118625 | 102,429609 | 4,237863 | 9,9952E-05 | 4,237863 | 0,74657171 | 0,45532218 |
| fmt | 1,12812766 | 4,39008604 | 1,08821128 | 4,2839E-06 | 1,08821128 | 0,74590414 | 0,45572535 |
| VC1250 | 1,17259925 | 4,22683191 | 1,10906779 | 4,1246E-06 | 1,10906779 | 0,74559213 | 0,45591387 |
| VC1765 | -1,2960566 | -28,472973 | -1,2352863 | -2,778E-05 | -1,2352863 | -0,745148 | 0,45618228 |
| rumA | 1,16810765 | 3,14466933 | 1,12483341 | 3,0686E-06 | 1,12483341 | 0,74448643 | 0,45658228 |
| VC0239 | 1,11493763 | 13,8073903 | 1,14294647 | 1,3473E-05 | 1,14294647 | 0,74413083 | 0,45679736 |
| VC1317 | 1,18176624 | 36,9062021 | 1,16100648 | 3,6013E-05 | 1,16100648 | 0,74396131 | 0,45689992 |
| VCA0673 | 1,33336978 | 8,58047408 | 1,28256129 | 8,3729E-06 | 1,28256129 | 0,74323282 | 0,45734077 |
| tRNA-His-2 | -1,3928847 | -17,706689 | -1,3418412 | -1,728E-05 | -1,3418412 | -0,7430713 | 0,45743855 |
| pyrE | -1,2050254 | -15,085492 | -1,1561821 | -1,472E-05 | -1,1561821 | -0,7429847 | 0,45749099 |
| VC0216 | -1,2536932 | -81,658241 | -1,3697404 | -7,968E-05 | -1,3697404 | -0,7420812 | 0,45803815 |
| lpxB | 1,26657504 | 1,97802059 | 1,20337308 | 1,9302E-06 | 1,20337308 | 0,74008403 | 0,45924904 |
| VC0263 | 1,1748064 | 3,42200828 | 1,11808378 | 3,3392E-06 | 1,11808378 | 0,73982426 | 0,45940666 |
| VC2274 | 1,25957009 | 4,25598268 | 1,21528353 | 4,153E-06 | 1,21528353 | 0,73965081 | 0,45951194 |
| VCA1096 | -1,2525584 | -1,2354901 | -1,3461372 | -1,206E-06 | -1,3461372 | -0,7394837 | 0,45961337 |
| VC2333 | -1,1676467 | -19,923117 | -1,2028784 | -1,944E-05 | -1,2028784 | -0,7386488 | 0,46012031 |
| VC1248 | -1,1928729 | -2,4848022 | -1,3112793 | -2,425E-06 | -1,3112793 | -0,7378432 | 0,46060979 |
| VC1514 | -1,4176788 | -37,500134 | -1,4160475 | -3,659E-05 | -1,4160475 | -0,7378275 | 0,46061935 |
| VC0337 | -1,0892675 | -2,3310948 | -1,1687122 | -2,275E-06 | -1,1687122 | -0,7375919 | 0,46076253 |
| VCA1045 | -1,2056925 | -4,1515889 | -1,3149922 | -4,051E-06 | -1,3149922 | -0,737408 | 0,46087427 |
| VC0074 | 1,23846219 | 2,68866866 | 1,18492192 | 2,6236E-06 | 1,18492192 | 0,73648858 | 0,46143344 |

| | | | | | | | |
|---|---|---|---|---|---|---|---|
| VC1819 | -1,2953405 | -0,9331086 | -1,4792803 | -9,105E-07 | -1,4792803 | -0,7356296 | 0,46195615 |
| VC1424 | -1,1073976 | -10,345073 | -1,1267948 | -1,009E-05 | -1,1267948 | -0,7348585 | 0,46242568 |
| VC1850 | 1,0826239 | 4,97302971 | 1,07499318 | 4,8527E-06 | 1,07499318 | 0,73432496 | 0,46275074 |
| cyaY | 1,13443186 | 7,35068763 | 1,12548103 | 7,1729E-06 | 1,12548103 | 0,73355267 | 0,46322145 |
| VCA0086 | 1,29518836 | 2,7862286 | 1,20008755 | 2,7188E-06 | 1,20008755 | 0,73338374 | 0,46332444 |
| VC1537 | -1,1520717 | -24,580868 | -1,1912482 | -2,399E-05 | -1,1912482 | -0,7331583 | 0,46346189 |
| VC0185 | 1,31419959 | 2,50822535 | 1,23010398 | 2,4475E-06 | 1,23010398 | 0,73285015 | 0,46364986 |
| VC0502 | 1,14803345 | 11,7375667 | 1,12452304 | 1,1454E-05 | 1,12452304 | 0,73158831 | 0,46441992 |
| VC1534 | 1,32749911 | 2,00015949 | 1,20110371 | 1,9518E-06 | 1,20110371 | 0,73155319 | 0,46444136 |
| VC2765 | 1,23475782 | 56,4688018 | 1,08311299 | 5,5103E-05 | 1,08311299 | 0,73101036 | 0,46477288 |
| VC1309 | 1,19815349 | 2,65344356 | 1,14376444 | 2,5893E-06 | 1,14376444 | 0,73064595 | 0,46499547 |
| VC2348 | -1,2880994 | -14,671102 | -1,3228303 | -1,432E-05 | -1,3228303 | -0,7304584 | 0,46511008 |
| VC1298 | 1,82639449 | 16,748918 | 1,62179904 | 1,6344E-05 | 1,62179904 | 0,73011053 | 0,46532267 |
| csrD | -1,1668514 | -2213,1391 | -1,2826502 | -0,0021596 | -1,2826502 | -0,7300375 | 0,46536727 |
| VC0614 | -1,6080429 | -1,7580107 | -1,7666064 | -1,715E-06 | -1,7666064 | -0,7293819 | 0,46576815 |
| VCA0308 | 1,16900704 | 20,5271927 | 1,21587477 | 2,0031E-05 | 1,21587477 | 0,72852875 | 0,46629002 |
| VCA0497 | 1,17689633 | 19,1418448 | 1,17964098 | 1,8679E-05 | 1,17964098 | 0,72837471 | 0,46638429 |
| rpsJ | 1,4491177 | 157,807618 | 1,25637312 | 0,00015399 | 1,25637312 | 0,72815176 | 0,46652072 |
| VCA0906 | -1,188999 | -1,504026 | -1,2664206 | -1,468E-06 | -1,2664206 | -0,7275838 | 0,46686842 |
| VC2298 | 1,27801942 | 18,9200816 | 1,21904137 | 1,8462E-05 | 1,21904137 | 0,7274274 | 0,46696423 |
| VC1951 | 1,39113924 | 51,1391058 | 1,42667766 | 4,9902E-05 | 1,42667766 | 0,72734617 | 0,46701397 |
| VC2215 | 1,23701998 | 2,16364143 | 1,18671883 | 2,1113E-06 | 1,18671883 | 0,72649367 | 0,46753622 |
| VC2380 | -1,1226491 | -2,3015781 | -1,1813897 | -2,246E-06 | -1,1813897 | -0,7260939 | 0,46778121 |
| VCA0658 | 1,59652357 | 7,97931139 | 1,40903473 | 7,7863E-06 | 1,40903473 | 0,72489658 | 0,46851553 |
| VC0367 | 1,25671214 | 209,555251 | 1,18718848 | 0,00020449 | 1,18718848 | 0,72428967 | 0,46888796 |
| VC1485 | 1,17112156 | 17,3218723 | 1,13716296 | 1,6903E-05 | 1,13716296 | 0,7242556 | 0,46890886 |
| VC1011 | 1,20613753 | 2,54713138 | 1,1473908 | 2,4855E-06 | 1,1473908 | 0,72421898 | 0,46893134 |
| VCA0053 | -1,4128978 | -106,44818 | -1,5674129 | -0,0001039 | -1,5674129 | -0,724106 | 0,46900071 |
| VC2105 | 1,45676247 | 7,2099002 | 1,31103541 | 7,0355E-06 | 1,31103541 | 0,72390343 | 0,46912507 |
| rpmE | 1,43106652 | 110,721764 | 1,18016529 | 0,00010804 | 1,18016529 | 0,72246108 | 0,47001108 |
| VCA0231 | 1,63634538 | 0,89151635 | 1,4846817 | 8,6995E-07 | 1,4846817 | 0,72230183 | 0,47010896 |
| VC2524 | 1,1530233 | 3,92851942 | 1,0960221 | 3,8335E-06 | 1,0960221 | 0,72200684 | 0,4702903 |
| VC0631 | 1,14336911 | 20,3704466 | 1,16487243 | 1,9878E-05 | 1,16487243 | 0,72174269 | 0,47045273 |
| VCA0362 | 1,20705856 | 5,47511647 | 1,14515725 | 5,3427E-06 | 1,14515725 | 0,72068404 | 0,47110396 |
| VCA0972 | 1,19523938 | 2,63341466 | 1,1406774 | 2,5697E-06 | 1,1406774 | 0,72054493 | 0,47118958 |
| VC2738 | -1,4229774 | -170,67088 | -1,5834491 | -0,0001665 | -1,5834491 | -0,7205362 | 0,47119493 |
| VC2749 | -1,0736202 | -2,3369239 | -1,1597035 | -2,28E-06 | -1,1597035 | -0,7200231 | 0,47151077 |
| rnhA | -1,0504706 | -2,6511372 | -1,1393449 | -2,587E-06 | -1,1393449 | -0,7197576 | 0,47167427 |
| VC2093 | 1,2853224 | 1,9721024 | 1,19139975 | 1,9244E-06 | 1,19139975 | 0,71884896 | 0,47223401 |
| VC1499 | 1,24258056 | 2,11558193 | 1,17719111 | 2,0644E-06 | 1,17719111 | 0,71870557 | 0,47232237 |
| VC2673 | 1,20044052 | 4,39823397 | 1,15653139 | 4,2918E-06 | 1,15653139 | 0,71822487 | 0,47261868 |
| VC0983 | 1,09758409 | 11,1872397 | 1,1145173 | 1,0917E-05 | 1,1145173 | 0,71795132 | 0,47278733 |
| tyrA | 1,25317455 | 3,52800988 | 1,14815524 | 3,4427E-06 | 1,14815524 | 0,71792762 | 0,47280195 |
| VC0924 | 1,24068595 | 2,16760024 | 1,17213809 | 2,1152E-06 | 1,17213809 | 0,71787349 | 0,47283532 |
| VC1835 | 1,37930901 | 194,601366 | 1,24343269 | 0,00018989 | 1,24343269 | 0,71716593 | 0,47327175 |

| | | | | | | | |
|---|---|---|---|---|---|---|---|
| VC1085 | -1,2326666 | -3,4213227 | -1,2965909 | -3,339E-06 | -1,2965909 | -0,7162926 | 0,47381072 |
| VCA0235 | -1,4550589 | -132,53877 | -1,3549014 | -0,0001293 | -1,3549014 | -0,7161194 | 0,47391764 |
| VC2319 | -1,1557384 | -1,8668322 | -1,2011063 | -1,822E-06 | -1,2011063 | -0,7153322 | 0,47440384 |
| VC0700 | 1,22072281 | 2,16477662 | 1,1707138 | 2,1124E-06 | 1,1707138 | 0,71466603 | 0,47481545 |
| VC0551 | -1,3190461 | -25,528377 | -1,3909516 | -2,491E-05 | -1,3909516 | -0,7132483 | 0,47569215 |
| cysE | 1,13435122 | 6,08632216 | 1,09679792 | 5,9391E-06 | 1,09679792 | 0,71311439 | 0,475775 |
| VC2650 | 1,26657785 | 9,32057012 | 1,18066646 | 9,0951E-06 | 1,18066646 | 0,71278894 | 0,47597641 |
| motB | -1,0881725 | -5,8409454 | -1,1286329 | -5,7E-06 | -1,1286329 | -0,7126955 | 0,47603421 |
| VCA1087 | -1,1685881 | -1,5133643 | -1,2518317 | -1,477E-06 | -1,2518317 | -0,7125593 | 0,47611851 |
| VCA0753 | 1,5563583 | 1,28376814 | 1,5362468 | 1,2527E-06 | 1,5362468 | 0,71167573 | 0,47666563 |
| VC1871 | -1,3609562 | -168,41567 | -1,4587001 | -0,0001643 | -1,4587001 | -0,7113982 | 0,47683755 |
| VC2569 | 1,37293451 | 9,65353595 | 1,3356533 | 9,42E-06 | 1,3356533 | 0,71103091 | 0,47706513 |
| VC0710 | -1,0979178 | -5,8291199 | -1,1174696 | -5,688E-06 | -1,1174696 | -0,7108269 | 0,47719153 |
| VC2201 | -1,0650357 | -7,3830371 | -1,0465751 | -7,204E-06 | -1,0465751 | -0,7100284 | 0,47768652 |
| VCA0388 | 1,14892076 | 3,44731853 | 1,10216295 | 3,3639E-06 | 1,10216295 | 0,70896109 | 0,47834865 |
| VC0613 | -1,5412824 | -1,7937489 | -1,6985649 | -1,75E-06 | -1,6985649 | -0,7085484 | 0,47860479 |
| ccrB | 1,1177501 | 11,8374452 | 1,10455379 | 1,1551E-05 | 1,10455379 | 0,70834142 | 0,47873329 |
| VC2667 | 1,4266768 | 5,705579 | 1,29687338 | 5,5676E-06 | 1,29687338 | 0,70815501 | 0,47884904 |
| flgI | -1,1242362 | -11,831065 | -1,1891147 | -1,154E-05 | -1,1891147 | -0,7080613 | 0,4789072 |
| VC1249 | -1,3528284 | -33,134635 | -1,4423864 | -3,233E-05 | -1,4423864 | -0,7078768 | 0,47902178 |
| VCA0659 | -1,2240517 | -9,6052284 | -1,2784787 | -9,373E-06 | -1,2784787 | -0,7077826 | 0,47908029 |
| VC2077 | 1,4505466 | 24,4983973 | 1,46035318 | 2,3906E-05 | 1,46035318 | 0,70717895 | 0,47945531 |
| 5Sh | -1,0723161 | -3354,9342 | -1,0299555 | -0,0032738 | -1,0299555 | -0,7071068 | 0,47950015 |
| VC1756 | 1,19845997 | 2,62173236 | 1,13570661 | 2,5583E-06 | 1,13570661 | 0,70695033 | 0,47959737 |
| metH | 1,21714453 | 2,29297633 | 1,15598977 | 2,2375E-06 | 1,15598977 | 0,70548616 | 0,48050777 |
| VC0106 | 1,23851332 | 2,04070381 | 1,17678428 | 1,9913E-06 | 1,17678428 | 0,70512712 | 0,48073116 |
| VC1574 | 1,42946055 | 51,4925971 | 1,35724072 | 5,0247E-05 | 1,35724072 | 0,70510579 | 0,48074443 |
| VC0429 | 1,25193327 | 2,04927417 | 1,17595245 | 1,9997E-06 | 1,17595245 | 0,70507663 | 0,48076258 |
| VCA0571 | 1,06510035 | 16,6752555 | 1,0739858 | 1,6272E-05 | 1,0739858 | 0,70473188 | 0,48097713 |
| VCA0767 | 1,21180258 | 2,9852291 | 1,11737164 | 2,913E-06 | 1,11737164 | 0,70459284 | 0,48106368 |
| VCA0501 | 1,18550906 | 5,13327696 | 1,14226667 | 5,0091E-06 | 1,14226667 | 0,70440142 | 0,48118285 |
| VC1882 | 1,23554739 | 2,14881401 | 1,16673183 | 2,0968E-06 | 1,16673183 | 0,70431947 | 0,48123387 |
| VC2433 | -1,3044418 | -24,722595 | -1,2412974 | -2,412E-05 | -1,2412974 | -0,7041328 | 0,48135014 |
| tRNA-Gly-6 | -1,3416324 | -33,423215 | -1,2702599 | -3,261E-05 | -1,2702599 | -0,7041278 | 0,48135322 |
| VC2470 | -1,1359306 | -13,817939 | -1,1717201 | -1,348E-05 | -1,1717201 | -0,703657 | 0,48164646 |
| VC1926 | -1,1941061 | -1,608424 | -1,3585892 | -1,57E-06 | -1,3585892 | -0,7030041 | 0,48205322 |
| VC2552 | 1,24942486 | 4,02277495 | 1,16206952 | 3,9255E-06 | 1,16206952 | 0,70069 | 0,48349653 |
| carB | 1,24431928 | 4,49553196 | 1,22401414 | 4,3868E-06 | 1,22401414 | 0,69974593 | 0,48408602 |
| VC0630 | -1,0189588 | -3,0563921 | -1,1145422 | -2,982E-06 | -1,1145422 | -0,6970947 | 0,48574358 |
| cmk | -1,1448462 | -6,9612485 | -1,1598399 | -6,793E-06 | -1,1598399 | -0,6950636 | 0,4870155 |
| VC0389 | 1,15442095 | 4,8809875 | 1,10448367 | 4,7629E-06 | 1,10448367 | 0,69494229 | 0,4870915 |
| VC2608 | 1,20696445 | 2,54900719 | 1,13471006 | 2,4873E-06 | 1,13471006 | 0,69467397 | 0,48725967 |
| VC2136 | -1,0364695 | -3,7598502 | -1,0893666 | -3,669E-06 | -1,0893666 | -0,6946011 | 0,48730536 |
| VC2080 | -1,044935 | -3,0793824 | -1,1101568 | -3,005E-06 | -1,1101568 | -0,6944597 | 0,48739401 |
| VC2255 | -1,0519283 | -4,7294027 | -1,0979557 | -4,615E-06 | -1,0979557 | -0,6942361 | 0,48753419 |

| | | | | | | | |
|---|---|---|---|---|---|---|---|
| VCA0359 | 1,18512254 | 2,56404248 | 1,13361443 | 2,502E-06 | 1,13361443 | 0,69405885 | 0,48764533 |
| VC0068 | 1,63781554 | 1,39220508 | 1,36875363 | 1,3585E-06 | 1,36875363 | 0,69349565 | 0,48799858 |
| VC1698 | -1,0595743 | -4,2864834 | -1,1973191 | -4,183E-06 | -1,1973191 | -0,69223 | 0,48879292 |
| VCA0662 | 1,20720899 | 5,69423119 | 1,13929582 | 5,5565E-06 | 1,13929582 | 0,69169847 | 0,48912673 |
| mgsA | -1,0587802 | -6,1999128 | -1,1528346 | -6,05E-06 | -1,1528346 | -0,6916436 | 0,48916122 |
| VC1906 | 1,24423506 | 2,49574386 | 1,14480677 | 2,4354E-06 | 1,14480677 | 0,69151568 | 0,48924156 |
| VC0547 | -1,1439316 | -18,110171 | -1,1236256 | -1,767E-05 | -1,1236256 | -0,6908298 | 0,48967253 |
| VC2747 | -1,1113692 | -1,7564442 | -1,1987625 | -1,714E-06 | -1,1987625 | -0,6901729 | 0,49008548 |
| VC1293 | 1,34521444 | 14,6594956 | 1,34178994 | 1,4305E-05 | 1,34178994 | 0,69016657 | 0,49008949 |
| VC2472 | 1,16330133 | 3,6837534 | 1,1180007 | 3,5946E-06 | 1,1180007 | 0,68986634 | 0,49027827 |
| lolB | -1,0632463 | -2,7967198 | -1,1202375 | -2,729E-06 | -1,1202375 | -0,6897918 | 0,49032519 |
| VC1035 | -1,1256658 | -1,5595452 | -1,2261605 | -1,522E-06 | -1,2261605 | -0,6894319 | 0,49055156 |
| VC1719 | 1,69738642 | 55,2485296 | 1,76974634 | 5,3912E-05 | 1,76974634 | 0,68930736 | 0,49062991 |
| VC0456 | 1,14033075 | 3,33473336 | 1,10683155 | 3,2541E-06 | 1,10683155 | 0,6885521 | 0,4911052 |
| VC1668 | 1,21347379 | 2,7053437 | 1,14312956 | 2,6399E-06 | 1,14312956 | 0,68800381 | 0,49145041 |
| glyQ | 1,19926899 | 6,76452168 | 1,15578444 | 6,6009E-06 | 1,15578444 | 0,68794178 | 0,49148947 |
| tRNA-Asp-5 | 1,2725227 | 8,37386422 | 1,21905655 | 8,1713E-06 | 1,21905655 | 0,68704379 | 0,49205517 |
| VC2207 | -1,0866179 | -36,743407 | -1,1378426 | -3,585E-05 | -1,1378426 | -0,6870433 | 0,49205547 |
| VC1666 | -1,1512902 | -1,4422069 | -1,2448605 | -1,407E-06 | -1,2448605 | -0,6869743 | 0,49209895 |
| rpsM | 1,17770018 | 77,6330795 | 1,06838056 | 7,5755E-05 | 1,06838056 | 0,68682975 | 0,49219004 |
| VC2365 | 1,1506143 | 4,69747056 | 1,1099764 | 4,5838E-06 | 1,1099764 | 0,68656022 | 0,49235993 |
| VCA0468 | 1,18916884 | 9,3494772 | 1,16693821 | 9,1233E-06 | 1,16693821 | 0,68626644 | 0,49254515 |
| VC0818 | 1,25366481 | 1,88686597 | 1,1814187 | 1,8412E-06 | 1,1814187 | 0,68612791 | 0,49263248 |
| VCA0470 | 1,2327295 | 4,83040202 | 1,16992545 | 4,7135E-06 | 1,16992545 | 0,68604455 | 0,49268505 |
| 16Sd | 1,04983716 | 456,318872 | 1,23482004 | 0,00044528 | 1,23482004 | 0,68588033 | 0,49278861 |
| VC1294 | 1,48645733 | 1,29382003 | 1,39219272 | 1,2625E-06 | 1,39219272 | 0,68584261 | 0,49281239 |
| tRNA-Ser-1 | 1,80461042 | 39,272826 | 1,91748133 | 3,8323E-05 | 1,91748133 | 0,6851942 | 0,49322144 |
| flgA | -1,0394796 | -4,3120726 | -1,1014826 | -4,208E-06 | -1,1014826 | -0,6849899 | 0,4933503 |
| infA | -1,2569805 | -42,708389 | -1,2525892 | -4,168E-05 | -1,2525892 | -0,6844126 | 0,4937147 |
| VC1102 | 1,33640244 | 1,9305086 | 1,20429969 | 1,8838E-06 | 1,20429969 | 0,68415131 | 0,49387965 |
| VC0145 | -1,0684458 | -6,0398582 | -1,0984446 | -5,894E-06 | -1,0984446 | -0,6835079 | 0,49428601 |
| VCA0506 | 1,15025346 | 4,06045201 | 1,10439696 | 3,9622E-06 | 1,10439696 | 0,68309578 | 0,49454635 |
| VC0114 | 1,27930087 | 5,73274645 | 1,22959344 | 5,5941E-06 | 1,22959344 | 0,6819474 | 0,49527225 |
| VCA0680 | 1,62809375 | 43,6926243 | 1,68685784 | 4,2636E-05 | 1,68685784 | 0,68049433 | 0,49619153 |
| VC1299 | 1,33308687 | 7,14876872 | 1,32190406 | 6,9758E-06 | 1,32190406 | 0,68041912 | 0,49623916 |
| VC1577 | 1,50121563 | 181,215745 | 1,3424404 | 0,00017683 | 1,3424404 | 0,67960315 | 0,4967558 |
| VCA0532 | 1,22916367 | 3,02965399 | 1,17115761 | 2,9564E-06 | 1,17115761 | 0,67909635 | 0,49707685 |
| VC0432 | -1,233232 | -69,175421 | -1,3099807 | -6,75E-05 | -1,3099807 | -0,6789323 | 0,49718082 |
| VC0079 | -1,1127014 | -10,592637 | -1,1325679 | -1,034E-05 | -1,1325679 | -0,6780005 | 0,49777139 |
| VC2252 | 1,09541066 | 8,13161457 | 1,07763276 | 7,9349E-06 | 1,07763276 | 0,67785941 | 0,49786087 |
| VC1210 | 1,22665989 | 2,98830955 | 1,13164182 | 2,916E-06 | 1,13164182 | 0,67780973 | 0,49789237 |
| VC0574 | 1,41547404 | 52,8138528 | 1,38799019 | 5,1536E-05 | 1,38799019 | 0,67770624 | 0,49795801 |
| VC0790 | 2,00857801 | 54,9110584 | 2,01978031 | 5,3583E-05 | 2,01978031 | 0,67633751 | 0,49882643 |
| VC1751 | 1,12379552 | 4,25864977 | 1,0925925 | 4,1556E-06 | 1,0925925 | 0,67630048 | 0,49884991 |
| tRNA-Leu-5 | -1,1518402 | -3,670646 | -1,1887896 | -3,582E-06 | -1,1887896 | -0,6758121 | 0,49915995 |

| | | | | | | | |
|---|---|---|---|---|---|---|---|
| VC1155 | -1,1723386 | -9,7792667 | -1,2532495 | -9,543E-06 | -1,2532495 | -0,6749325 | 0,49971869 |
| VC0112 | 1,3085287 | 79,2884497 | 1,20996809 | 7,737E-05 | 1,20996809 | 0,67479745 | 0,49980449 |
| rfaH | -1,0669262 | -5,3914058 | -1,1163893 | -5,261E-06 | -1,1163893 | -0,6746545 | 0,49989531 |
| VC0837 | 1,1250743 | 12,5502522 | 1,13971193 | 1,2247E-05 | 1,13971193 | 0,67392935 | 0,50035626 |
| VCA0580 | 1,15190941 | 3,27841018 | 1,09674278 | 3,1991E-06 | 1,09674278 | 0,67365343 | 0,5005317 |
| VC2183 | -1,181827 | -11,531575 | -1,2147373 | -1,125E-05 | -1,2147373 | -0,6735008 | 0,50062874 |
| VC0260 | 1,07779086 | 6,42855261 | 1,08999155 | 6,273E-06 | 1,08999155 | 0,67318262 | 0,50083115 |
| VCA0829 | 1,38748183 | 1,40807351 | 1,23983313 | 1,374E-06 | 1,23983313 | 0,67254516 | 0,50123672 |
| minC | 1,25625397 | 3,50917638 | 1,16928039 | 3,4243E-06 | 1,16928039 | 0,67250175 | 0,50126436 |
| VC1282 | 1,37677426 | 1,52552612 | 1,21929705 | 1,4886E-06 | 1,21929705 | 0,67248217 | 0,50127681 |
| VC0200 | 1,63707443 | 12,3973802 | 1,59315562 | 1,2097E-05 | 1,59315562 | 0,67122394 | 0,50207791 |
| VC1745 | 1,2272361 | 2,10666209 | 1,15343511 | 2,0557E-06 | 1,15343511 | 0,67105435 | 0,50218593 |
| VC2645 | 1,32380189 | 2,91894427 | 1,20062383 | 2,8483E-06 | 1,20062383 | 0,67093128 | 0,50226434 |
| ulaA | -1,1197994 | -1,5386792 | -1,3555654 | -1,501E-06 | -1,3555654 | -0,6708763 | 0,50229936 |
| VCA0264 | 1,41272223 | 1,43820692 | 1,23292292 | 1,4034E-06 | 1,23292292 | 0,67087566 | 0,50229977 |
| VC0194 | 1,5890762 | 0,90115372 | 1,3984245 | 8,7935E-07 | 1,3984245 | 0,67014788 | 0,50276355 |
| VC0381 | -1,1245257 | -9,0289266 | -1,13371 | -8,811E-06 | -1,13371 | -0,6692956 | 0,50330699 |
| flgC | -1,1121559 | -26,685921 | -1,121537 | -2,604E-05 | -1,121537 | -0,6692477 | 0,50333749 |
| VCA0595 | 1,21469629 | 2,46690583 | 1,12871431 | 2,4072E-06 | 1,12871431 | 0,66895357 | 0,50352512 |
| VC0618 | -1,8361624 | -5,7469774 | -1,9518426 | -5,608E-06 | -1,9518426 | -0,6676129 | 0,50438076 |
| VCA0558 | -1,149255 | -3,2652401 | -1,2301299 | -3,186E-06 | -1,2301299 | -0,6673696 | 0,50453609 |
| VC2128 | -1,0911765 | -7,9815701 | -1,1514385 | -7,788E-06 | -1,1514385 | -0,6672991 | 0,5045811 |
| VC2511 | 1,19863762 | 2,17739388 | 1,14574115 | 2,1247E-06 | 1,14574115 | 0,66609315 | 0,50535158 |
| VC2185 | -1,1520283 | -8,9827025 | -1,1678047 | -8,765E-06 | -1,1678047 | -0,6659239 | 0,5054598 |
| VC0391 | 1,26718723 | 4,17378139 | 1,20360985 | 4,0728E-06 | 1,20360985 | 0,6658535 | 0,50550478 |
| VCA0402 | 1,28210845 | 2,13395342 | 1,22492706 | 2,0823E-06 | 1,22492706 | 0,6655843 | 0,50567687 |
| VC2229 | 1,09203785 | 4,68871446 | 1,06629894 | 4,5753E-06 | 1,06629894 | 0,66540189 | 0,5057935 |
| VC1978 | 1,46294137 | 1,25316983 | 1,26683034 | 1,2229E-06 | 1,26683034 | 0,66523461 | 0,50590048 |
| VC1794 | 1,27730611 | 1,43762323 | 1,22859985 | 1,4028E-06 | 1,22859985 | 0,66513008 | 0,50596732 |
| astD | -1,3595405 | -3,6076254 | -1,5534121 | -3,52E-06 | -1,5534121 | -0,6640112 | 0,5066832 |
| VCA0905 | 1,56562045 | 5,33898831 | 1,37487244 | 5,2098E-06 | 1,37487244 | 0,66302904 | 0,507312 |
| fliJ | -1,0385333 | -2,8965764 | -1,1191853 | -2,827E-06 | -1,1191853 | -0,6624709 | 0,50766953 |
| VC1917 | -1,1193195 | -1,1684693 | -1,2860808 | -1,14E-06 | -1,2860808 | -0,6623215 | 0,50776523 |
| VC1363 | 1,12335741 | 3,50603456 | 1,08681298 | 3,4212E-06 | 1,08681298 | 0,66149754 | 0,50829332 |
| VC1614 | 1,76911618 | 11,7243843 | 1,56363513 | 1,1441E-05 | 1,56363513 | 0,66011124 | 0,50918249 |
| VCA1066 | -1,2128898 | -2,4391582 | -1,2970486 | -2,38E-06 | -1,2970486 | -0,6598831 | 0,50932888 |
| VC0212 | -1,1113544 | -7,4789494 | -1,1247838 | -7,298E-06 | -1,1247838 | -0,6595203 | 0,50956174 |
| VCA0100 | 1,18233423 | 2,83246014 | 1,10779438 | 2,7639E-06 | 1,10779438 | 0,659223 | 0,5097526 |
| glpE | -1,0375241 | -3,2500719 | -1,0977877 | -3,171E-06 | -1,0977877 | -0,6584551 | 0,51024579 |
| flgF | -1,1112728 | -19,566544 | -1,1422198 | -1,909E-05 | -1,1422198 | -0,6583061 | 0,51034146 |
| VC2095 | -1,1674537 | -18,651409 | -1,2801143 | -1,82E-05 | -1,2801143 | -0,6582827 | 0,51035651 |
| VC1744 | 1,18715109 | 2,64611917 | 1,11536827 | 2,5821E-06 | 1,11536827 | 0,65799209 | 0,51054324 |
| VCA0892 | 1,40404071 | 27,3985918 | 1,33832338 | 2,6736E-05 | 1,33832338 | 0,65777391 | 0,51068346 |
| VC1644 | 1,35229402 | 29,7586867 | 1,33510738 | 2,9039E-05 | 1,33510738 | 0,656561 | 0,51146327 |
| VC0581 | 1,14008819 | 4,6630428 | 1,09217775 | 4,5502E-06 | 1,09217775 | 0,65653794 | 0,51147809 |

| | | | | | | | |
|---|---|---|---|---|---|---|---|
| fre | 1,14349998 | 3,423351 | 1,10070436 | 3,3405E-06 | 1,10070436 | 0,65636688 | 0,51158812 |
| VCA0877 | 1,21255828 | 2,34451934 | 1,14328398 | 2,2878E-06 | 1,14328398 | 0,6563311 | 0,51161113 |
| VC1515 | -1,2206084 | -12,008041 | -1,2665291 | -1,172E-05 | -1,2665291 | -0,6563154 | 0,51162125 |
| VC0676 | -1,1164383 | -1,3981524 | -1,2286591 | -1,364E-06 | -1,2286591 | -0,6560119 | 0,51181646 |
| VC1622 | 1,29550946 | 12,2576396 | 1,24250456 | 1,1961E-05 | 1,24250456 | 0,65555855 | 0,51210823 |
| flgB | -1,120987 | -157,81392 | -1,1965164 | -0,000154 | -1,1965164 | -0,6551028 | 0,51240161 |
| VC0377 | -1,0763848 | -28,260411 | -1,1054178 | -2,758E-05 | -1,1054178 | -0,6540901 | 0,51305382 |
| VC2421 | 1,1943995 | 2,43622832 | 1,12433746 | 2,3773E-06 | 1,12433746 | 0,65405351 | 0,51307736 |
| VC0962 | 1,33243285 | 5,0950789 | 1,31097441 | 4,9718E-06 | 1,31097441 | 0,65329485 | 0,51356625 |
| VCA0417 | 1,19927358 | 2,28993341 | 1,14672079 | 2,2345E-06 | 1,14672079 | 0,65268142 | 0,51396172 |
| ribH | -1,1377554 | -50,627182 | -1,1801191 | -4,94E-05 | -1,1801191 | -0,6525566 | 0,51404222 |
| VC0115 | 1,2207999 | 4,36744032 | 1,18857068 | 4,2618E-06 | 1,18857068 | 0,65112485 | 0,51496593 |
| VC0535 | 1,29631784 | 1,64933869 | 1,1870183 | 1,6094E-06 | 1,1870183 | 0,65047926 | 0,51538272 |
| VC0352 | 1,18453773 | 2,72419998 | 1,10919652 | 2,6583E-06 | 1,10919652 | 0,65047647 | 0,51538452 |
| VC1303 | 1,22295322 | 2,18394355 | 1,13790426 | 2,1311E-06 | 1,13790426 | 0,65009922 | 0,51562815 |
| VC1375 | 1,33265213 | 2,86596547 | 1,25523736 | 2,7966E-06 | 1,25523736 | 0,64977599 | 0,51583697 |
| VC1898 | -1,1698191 | -23,175429 | -1,2569865 | -2,261E-05 | -1,2569865 | -0,6482578 | 0,51681828 |
| secF | -1,0647825 | -10,680751 | -1,1004053 | -1,042E-05 | -1,1004053 | -0,6482243 | 0,5168399 |
| VC2334 | 1,20549445 | 2,40088867 | 1,12366259 | 2,3428E-06 | 1,12366259 | 0,6476307 | 0,51722386 |
| tRNA-Thr-4 | -1,3082735 | -36,258521 | -1,2644844 | -3,538E-05 | -1,2644844 | -0,6474377 | 0,51734877 |
| VC1522 | -1,0748166 | -2,5845129 | -1,1619175 | -2,522E-06 | -1,1619175 | -0,6473603 | 0,51739881 |
| tRNA-Thr-5 | -1,3763333 | -48,299178 | -1,3439699 | -4,713E-05 | -1,3439699 | -0,6470737 | 0,5175843 |
| VC1928 | 1,90352852 | 1,96275694 | 1,78141142 | 1,9153E-06 | 1,78141142 | 0,64681296 | 0,51775305 |
| ispD | 1,20158915 | 1,96549765 | 1,1526279 | 1,9179E-06 | 1,1526279 | 0,64659497 | 0,51789413 |
| modC | -1,0946089 | -2,0325542 | -1,1479066 | -1,983E-06 | -1,1479066 | -0,646287 | 0,51809353 |
| VC0198 | 1,49907656 | 1,18091773 | 1,29699759 | 1,1523E-06 | 1,29699759 | 0,64544286 | 0,51864026 |
| VC1508 | 1,21430881 | 7,5239194 | 1,17571346 | 7,3419E-06 | 1,17571346 | 0,64476055 | 0,5190824 |
| trpD | -1,0943899 | -1,6355872 | -1,185063 | -1,596E-06 | -1,185063 | -0,644655 | 0,5191508 |
| VC1567 | 1,28491144 | 1,26934365 | 1,24466001 | 1,2386E-06 | 1,24466001 | 0,64425484 | 0,51941022 |
| VCA0939 | -1,0774598 | -2,0419801 | -1,1453142 | -1,993E-06 | -1,1453142 | -0,6441666 | 0,51946742 |
| VCA0077 | 1,18572584 | 2,20846299 | 1,13319973 | 2,155E-06 | 1,13319973 | 0,64319901 | 0,520095 |
| VC0234 | 1,12034664 | 6,4229833 | 1,10883012 | 6,2676E-06 | 1,10883012 | 0,64249967 | 0,52054883 |
| ipk | -1,1383016 | -8,0135649 | -1,1602449 | -7,82E-06 | -1,1602449 | -0,6417416 | 0,52104099 |
| VC2622 | -1,1354119 | -1,0723788 | -1,293346 | -1,046E-06 | -1,293346 | -0,6414915 | 0,52120343 |
| VCA0923 | -1,0762362 | -4,2600926 | -1,1991956 | -4,157E-06 | -1,1991956 | -0,6408935 | 0,52159189 |
| argC | 1,43557514 | 1,05712358 | 1,29753519 | 1,0316E-06 | 1,29753519 | 0,64085898 | 0,52161434 |
| VC0250 | 1,28087957 | 112,782364 | 1,21834086 | 0,00011005 | 1,21834086 | 0,6404859 | 0,52185679 |
| mscL | 1,20952149 | 2,64686921 | 1,11620031 | 2,5828E-06 | 1,11620031 | 0,64028627 | 0,52198653 |
| VC2671 | -1,123077 | -1,3886741 | -1,2182678 | -1,355E-06 | -1,2182678 | -0,6402509 | 0,52200955 |
| VC1833 | 1,12418048 | 5,57007846 | 1,11586734 | 5,4353E-06 | 1,11586734 | 0,63853711 | 0,52312414 |
| VCA0019 | 1,45017673 | 1,08485164 | 1,28609837 | 1,0586E-06 | 1,28609837 | 0,63820025 | 0,52334336 |
| VCA0645 | 1,43163533 | 0,98851248 | 1,31818325 | 9,646E-07 | 1,31818325 | 0,63799498 | 0,52347697 |
| VC2052 | -1,2842486 | -20,922818 | -1,2590266 | -2,042E-05 | -1,2590266 | -0,6375283 | 0,52378083 |
| VC1712 | 1,45112095 | 1,35312268 | 1,29101588 | 1,3204E-06 | 1,29101588 | 0,63719426 | 0,52399834 |
| VC2635 | 1,17600085 | 3,44225436 | 1,1087926 | 3,359E-06 | 1,1087926 | 0,63699231 | 0,52412988 |

| | | | | | | | |
|---|---|---|---|---|---|---|---|
| VC1701 | 1,53257596 | 2,35473272 | 1,39843542 | 2,2978E-06 | 1,39843542 | 0,63676086 | 0,52428066 |
| ompR | 1,24309248 | 6,13999723 | 1,1614405 | 5,9915E-06 | 1,1614405 | 0,63588948 | 0,52484849 |
| VC2278 | 1,22739342 | 5,80190161 | 1,20449902 | 5,6615E-06 | 1,20449902 | 0,63530007 | 0,52523276 |
| VCA0474 | 1,16681581 | 2,25498182 | 1,12688528 | 2,2004E-06 | 1,12688528 | 0,6352867 | 0,52524147 |
| VC2012 | -1,1091443 | -1,3577642 | -1,2190872 | -1,325E-06 | -1,2190872 | -0,6341553 | 0,52597951 |
| VCA0095 | 1,76812608 | 0,79902636 | 1,43445669 | 7,797E-07 | 1,43445669 | 0,63385029 | 0,52617856 |
| dnaG | 1,13256167 | 5,26068678 | 1,11801377 | 5,1334E-06 | 1,11801377 | 0,63382115 | 0,52619758 |
| VC1491 | -1,2017158 | -18,688689 | -1,1980879 | -1,824E-05 | -1,1980879 | -0,6326764 | 0,526945 |
| VCA0066 | 1,6362912 | 0,67964446 | 1,48848913 | 6,632E-07 | 1,48848913 | 0,63264704 | 0,52696418 |
| VC0008 | 1,36212912 | 3,34517929 | 1,3168812 | 3,2643E-06 | 1,3168812 | 0,63190968 | 0,52744593 |
| VCA0166 | -1,4754827 | -833,58124 | -1,1754851 | -0,0008134 | -1,1754851 | -0,6303903 | 0,5284393 |
| VC1497 | 1,16585946 | 42,3413808 | 1,15577035 | 4,1317E-05 | 1,15577035 | 0,63019537 | 0,52856681 |
| VC1853 | 1,08645712 | 15,143414 | 1,05338582 | 1,4777E-05 | 1,05338582 | 0,62988027 | 0,52877296 |
| VC0660 | 1,26395752 | 5,31513752 | 1,23090828 | 5,1866E-06 | 1,23090828 | 0,62966898 | 0,52891122 |
| VC2277 | 1,17132934 | 11,0827039 | 1,15307871 | 1,0815E-05 | 1,15307871 | 0,62902243 | 0,5293344 |
| VCA0193 | -1,1045427 | -1,4635398 | -1,197891 | -1,428E-06 | -1,197891 | -0,628745 | 0,52951601 |
| VC0603 | 1,13450625 | 3,25259421 | 1,08435471 | 3,1739E-06 | 1,08435471 | 0,62842428 | 0,52972605 |
| VC0257 | -1,3567611 | -8,7691603 | -1,344446 | -8,557E-06 | -1,344446 | -0,6282017 | 0,52987185 |
| VCA0356a | 1,21236694 | 7,71048324 | 1,17486122 | 7,524E-06 | 1,17486122 | 0,6274534 | 0,53036209 |
| VC1754 | 1,12467236 | 3,13830006 | 1,08727315 | 3,0624E-06 | 1,08727315 | 0,62743177 | 0,53037628 |
| glgA | -1,1168146 | -17,398195 | -1,1619555 | -1,698E-05 | -1,1619555 | -0,626273 | 0,53113592 |
| VC2533 | -1,1197754 | -1,4102385 | -1,2041785 | -1,376E-06 | -1,2041785 | -0,6260228 | 0,5313 |
| VC1957 | 1,10059123 | 10,6307111 | 1,08099856 | 1,0374E-05 | 1,08099856 | 0,62592096 | 0,5313668 |
| VC1711 | 1,34515567 | 1,18586852 | 1,25949052 | 1,1572E-06 | 1,25949052 | 0,62576887 | 0,53146658 |
| VC2382 | -1,1087917 | -1,5580687 | -1,1825397 | -1,52E-06 | -1,1825397 | -0,6252498 | 0,53180716 |
| VCA1105 | -1,1809489 | -3,8408278 | -1,2659882 | -3,748E-06 | -1,2659882 | -0,6251527 | 0,53187085 |
| VCA0815 | 1,21031894 | 5,54784589 | 1,15610097 | 5,4136E-06 | 1,15610097 | 0,62503305 | 0,5319494 |
| rimO | 1,27012766 | 1,36948454 | 1,21665936 | 1,3364E-06 | 1,21665936 | 0,62447605 | 0,53231502 |
| VC0229 | 1,07088103 | 8,41927785 | 1,07579742 | 8,2156E-06 | 1,07579742 | 0,62441043 | 0,53235811 |
| VC0132 | -1,0527188 | -2,2381083 | -1,122903 | -2,184E-06 | -1,122903 | -0,6234784 | 0,53297023 |
| VC0036 | 1,35967754 | 20,7637919 | 1,34824632 | 2,0261E-05 | 1,34824632 | 0,62315373 | 0,53318355 |
| VCA0896 | 1,27681171 | 2,60065291 | 1,21607811 | 2,5377E-06 | 1,21607811 | 0,62307631 | 0,5332344 |
| VCA0284 | -1,017805 | -2,9115089 | -1,0929509 | -2,841E-06 | -1,0929509 | -0,6228363 | 0,53339212 |
| VC2768 | 1,25318955 | 94,9075245 | 1,13525476 | 9,2612E-05 | 1,13525476 | 0,62279598 | 0,53341864 |
| tRNA-Gly-4 | 1,29331502 | 1,34613026 | 1,21225408 | 1,3136E-06 | 1,21225408 | 0,6224668 | 0,53363499 |
| VCA0018 | -1,4533131 | -0,5721785 | -1,5819313 | -5,583E-07 | -1,5819313 | -0,6219997 | 0,53394206 |
| VC2032 | -1,1887589 | -1,5637643 | -1,2441195 | -1,526E-06 | -1,2441195 | -0,6216794 | 0,53415273 |
| phhA | 1,5598716 | 1,45378677 | 1,31511896 | 1,4186E-06 | 1,31511896 | 0,6208533 | 0,53469618 |
| VC0896 | 1,32484631 | 1,35147065 | 1,210105 | 1,3188E-06 | 1,210105 | 0,62083643 | 0,53470727 |
| thiE | 1,42979668 | 0,99143401 | 1,29671683 | 9,6745E-07 | 1,29671683 | 0,61988443 | 0,5353339 |
| VC1580 | 1,36206566 | 1,30926425 | 1,21685736 | 1,2776E-06 | 1,21685736 | 0,61986044 | 0,53534969 |
| VC1517 | -1,1407004 | -12,717296 | -1,2003224 | -1,241E-05 | -1,2003224 | -0,6190048 | 0,53591327 |
| VC1927 | 1,63661171 | 0,99894342 | 1,52484074 | 9,7478E-07 | 1,52484074 | 0,61784992 | 0,5366743 |
| VCA0110 | 1,42519633 | 1,08129251 | 1,26672002 | 1,0551E-06 | 1,26672002 | 0,61782016 | 0,53669391 |
| VC1628 | 1,2422471 | 4,57581214 | 1,23527355 | 4,4651E-06 | 1,23527355 | 0,61779054 | 0,53671346 |

| | | | | | | | |
|---|---|---|---|---|---|---|---|
| VC0568 | 1,24815924 | 4,73350063 | 1,14464648 | 4,619E-06 | 1,14464648 | 0,61713781 | 0,53714386 |
| VC0393 | -1,0464866 | -2,2003124 | -1,1218508 | -2,147E-06 | -1,1218508 | -0,6156923 | 0,53809765 |
| VC0330 | -1,0966935 | -14,433266 | -1,1379046 | -1,408E-05 | -1,1379046 | -0,6152411 | 0,53839553 |
| VCA0092 | 1,20750168 | 2,16191101 | 1,12356165 | 2,1096E-06 | 1,12356165 | 0,61431751 | 0,53900555 |
| VC1147 | -1,1102467 | -8,1666361 | -1,1919002 | -7,969E-06 | -1,1919002 | -0,6140454 | 0,53918539 |
| VC1993 | 1,79610672 | 1,46339317 | 1,47494247 | 1,428E-06 | 1,47494247 | 0,61395759 | 0,53924339 |
| VC2211 | 1,33265354 | 1,29541715 | 1,21481897 | 1,2641E-06 | 1,21481897 | 0,61395151 | 0,53924739 |
| VC0426 | -1,0535681 | -2,6084213 | -1,1129196 | -2,545E-06 | -1,1129196 | -0,6133229 | 0,53966285 |
| VCA0561 | 1,1574507 | 5,52330301 | 1,10895898 | 5,3897E-06 | 1,10895898 | 0,61318759 | 0,53975233 |
| tRNA-Gln-5 | -1,362288 | -2,0578499 | -1,367443 | -2,008E-06 | -1,367443 | -0,613032 | 0,53985524 |
| VC2540 | 1,16806155 | 7,44710238 | 1,15276391 | 7,2669E-06 | 1,15276391 | 0,61156156 | 0,54082791 |
| VC0615 | -1,8054796 | -7,8582915 | -1,8681353 | -7,668E-06 | -1,8681353 | -0,6114542 | 0,54089893 |
| VC2168 | 1,20395461 | 2,19365158 | 1,12042078 | 2,1406E-06 | 1,12042078 | 0,61134765 | 0,54096948 |
| fliQ | -1,1039415 | -5,946923 | -1,138148 | -5,803E-06 | -1,138148 | -0,6113332 | 0,54097902 |
| VC0237 | 1,07109307 | 15,7638149 | 1,12989948 | 1,5382E-05 | 1,12989948 | 0,60961646 | 0,54211593 |
| VC1789 | -1,3675974 | -9,0009295 | -1,3484981 | -8,783E-06 | -1,3484981 | -0,6094918 | 0,54219855 |
| VC1884 | 1,21743558 | 1,81453914 | 1,14641199 | 1,7706E-06 | 1,14641199 | 0,60936627 | 0,54228172 |
| VCA0809 | -1,111839 | -1,7766769 | -1,2036005 | -1,734E-06 | -1,2036005 | -0,6086883 | 0,54273112 |
| VC0397 | -1,0340803 | -73,743903 | -1,2094584 | -7,196E-05 | -1,2094584 | -0,6084234 | 0,54290671 |
| VC2142 | -1,2127956 | -37,826052 | -1,23335 | -3,691E-05 | -1,23335 | -0,608081 | 0,54313381 |
| VC1840 | 1,13223889 | 6,68754376 | 1,10597788 | 6,5258E-06 | 1,10597788 | 0,60796826 | 0,54320855 |
| VC1411 | 1,25255436 | 1,54073911 | 1,17340103 | 1,5035E-06 | 1,17340103 | 0,60727184 | 0,54367055 |
| tRNA-Glu-4 | -1,5665082 | -0,4679144 | -1,7106368 | -4,566E-07 | -1,7106368 | -0,6066416 | 0,54408879 |
| VCA0628 | 1,45189794 | 42,7485136 | 1,40438421 | 4,1714E-05 | 1,40438421 | 0,6064974 | 0,54418454 |
| VC0269 | 1,15581029 | 13,9943424 | 1,16725522 | 1,3656E-05 | 1,16725522 | 0,60571444 | 0,54470442 |
| VC2202 | 1,01366272 | 6,52188176 | 1,0381431 | 6,3641E-06 | 1,0381431 | 0,60516623 | 0,54506857 |
| VC1204 | 1,88248807 | 2,45094356 | 1,56706886 | 2,3917E-06 | 1,56706886 | 0,60498788 | 0,54518708 |
| VC0266 | -1,0607982 | -1,615978 | -1,1735741 | -1,577E-06 | -1,1735741 | -0,6036552 | 0,54607294 |
| VC2153 | 1,17610292 | 2,99268064 | 1,11052984 | 2,9203E-06 | 1,11052984 | 0,60341019 | 0,54623587 |
| VCA1088 | -1,1325197 | -1,4118358 | -1,1880727 | -1,378E-06 | -1,1880727 | -0,6033738 | 0,54626005 |
| VCA1057 | 1,52872411 | 0,69571303 | 1,42229826 | 6,7888E-07 | 1,42229826 | 0,60321478 | 0,54636585 |
| mltA | 1,16179333 | 2,99414879 | 1,10067938 | 2,9217E-06 | 1,10067938 | 0,60277086 | 0,54666116 |
| VC2682 | -1,1196691 | -7,7428327 | -1,1427969 | -7,556E-06 | -1,1427969 | -0,6025787 | 0,54678902 |
| VC2686 | 1,16284617 | 39,3153959 | 1,18585162 | 3,8364E-05 | 1,18585162 | 0,60229951 | 0,54697482 |
| VC2397 | -1,0796693 | -33,21547 | -1,119285 | -3,241E-05 | -1,119285 | -0,6013377 | 0,54761511 |
| avtA | 1,15807884 | 2,81192436 | 1,08947535 | 2,7439E-06 | 1,08947535 | 0,60103878 | 0,54781417 |
| argS | 1,22902475 | 6,4055314 | 1,20388643 | 6,2506E-06 | 1,20388643 | 0,60087582 | 0,54792273 |
| VC2666 | 1,32855956 | 1,28267569 | 1,20685059 | 1,2516E-06 | 1,20685059 | 0,60056835 | 0,54812755 |
| rluD | -1,1203349 | -11,273976 | -1,1370433 | -1,1E-05 | -1,1370433 | -0,6000236 | 0,54849051 |
| VC1258 | -1,1170963 | -21,800976 | -1,1350196 | -2,127E-05 | -1,1350196 | -0,5998835 | 0,54858392 |
| VCA0329b | 1,21628176 | 1,59530201 | 1,16198084 | 1,5567E-06 | 1,16198084 | 0,59881166 | 0,54929851 |
| secE | 1,12703791 | 29,874403 | 1,11529366 | 2,9152E-05 | 1,11529366 | 0,59822493 | 0,54968988 |
| VC2320 | -1,1210684 | -1,5124885 | -1,1711934 | -1,476E-06 | -1,1711934 | -0,5981402 | 0,54974639 |
| VCA0041 | 1,16675116 | 3,16801861 | 1,10640274 | 3,0914E-06 | 1,10640274 | 0,59728032 | 0,55032026 |
| VC1209 | 1,13873732 | 34,1722521 | 1,13394244 | 3,3346E-05 | 1,13394244 | 0,59605508 | 0,55113845 |

| | | | | | | | |
|---|---|---|---|---|---|---|---|
| VC0948 | 1,09354056 | 9,17842532 | 1,08396543 | 8,9564E-06 | 1,08396543 | 0,59580454 | 0,55130584 |
| VC1671 | 1,1818037 | 2,49630014 | 1,09938282 | 2,4359E-06 | 1,09938282 | 0,59542143 | 0,55156182 |
| VC1080 | 1,388928 | 7,54656839 | 1,2968503 | 7,364E-06 | 1,2968503 | 0,59476158 | 0,55200289 |
| VC2338 | -1,2228592 | -36,674807 | -1,2998942 | -3,579E-05 | -1,2998942 | -0,5944338 | 0,55222204 |
| VC0579 | 1,07548814 | 6,73716889 | 1,06761115 | 6,5742E-06 | 1,06761115 | 0,59440918 | 0,55223849 |
| VCA0689 | 1,3612446 | 1,12691497 | 1,23314002 | 1,0997E-06 | 1,23314002 | 0,59409759 | 0,55244686 |
| VC1562 | 1,36765613 | 1,13372787 | 1,23134111 | 1,1063E-06 | 1,23134111 | 0,59382658 | 0,55262813 |
| VC0423 | 1,52058666 | 4,30292179 | 1,31528667 | 4,1988E-06 | 1,31528667 | 0,59357067 | 0,55279934 |
| VCA0991 | 1,38222571 | 1,12062943 | 1,2336246 | 1,0935E-06 | 1,2336246 | 0,59298947 | 0,55318822 |
| VC2600 | -1,0798467 | -1,8277404 | -1,1370096 | -1,784E-06 | -1,1370096 | -0,5929159 | 0,55323742 |
| VC2147 | -1,1002369 | -5,3700045 | -1,1459118 | -5,24E-06 | -1,1459118 | -0,5927913 | 0,55332086 |
| VC2434 | -1,3545492 | -24,94138 | -1,2620673 | -2,434E-05 | -1,2620673 | -0,5926008 | 0,55344835 |
| VC0658 | -1,025515 | -2,210223 | -1,1170244 | -2,157E-06 | -1,1170244 | -0,5918663 | 0,55394015 |
| VC0077 | -1,1242539 | -1,0058885 | -1,2624657 | -9,816E-07 | -1,2624657 | -0,591673 | 0,55406959 |
| VC0153 | 1,20508247 | 2,12747548 | 1,11595107 | 2,076E-06 | 1,11595107 | 0,59140039 | 0,5542522 |
| tRNA-Asp-4 | 1,40504182 | 34,2308637 | 1,47074696 | 3,3403E-05 | 1,47074696 | 0,5908691 | 0,55460818 |
| VC1759 | 1,1728359 | 2,47763041 | 1,09839033 | 2,4177E-06 | 1,09839033 | 0,59036087 | 0,55494876 |
| VC0213 | -1,0507523 | -4,0472848 | -1,103292 | -3,949E-06 | -1,103292 | -0,588409 | 0,55625785 |
| VC2305 | 1,16279936 | 14,3466534 | 1,17072105 | 1,4E-05 | 1,17072105 | 0,58800344 | 0,55653001 |
| VC1910 | -1,0793426 | -8,2102462 | -1,0632796 | -8,012E-06 | -1,0632796 | -0,5874978 | 0,55686947 |
| tRNA-Phe-1 | 1,74614453 | 0,51166179 | 1,6030737 | 4,9928E-07 | 1,6030737 | 0,58697313 | 0,55722178 |
| tRNA-Asn-3 | -1,3116152 | -27,061044 | -1,2757303 | -2,641E-05 | -1,2757303 | -0,5866951 | 0,55740855 |
| VC0627 | 1,24943137 | 15,9688741 | 1,21230695 | 1,5583E-05 | 1,21230695 | 0,58655982 | 0,55749941 |
| VC0936 | 1,38597254 | 0,98123395 | 1,26446578 | 9,575E-07 | 1,26446578 | 0,58634021 | 0,55764694 |
| VC1158 | -1,0672944 | -8,6564556 | -1,0828486 | -8,447E-06 | -1,0828486 | -0,5860595 | 0,55783557 |
| VCA0849 | 1,2819561 | 1,2712826 | 1,19776901 | 1,2405E-06 | 1,19776901 | 0,58582957 | 0,55799008 |
| fliN | -1,1083118 | -10,225457 | -1,1087793 | -9,978E-06 | -1,1087793 | -0,585522 | 0,55819684 |
| VC1633 | 1,19292507 | 7,97365557 | 1,17022575 | 7,7808E-06 | 1,17022575 | 0,58471359 | 0,55874034 |
| VC0118 | -1,0910413 | -3,7793964 | -1,1858058 | -3,688E-06 | -1,1858058 | -0,5846595 | 0,55877671 |
| VCA0217 | 1,3679301 | 1,21573303 | 1,2271159 | 1,1863E-06 | 1,2271159 | 0,58453936 | 0,55885751 |
| VCA0311 | -1,0953218 | -10,345928 | -1,1085414 | -1,01E-05 | -1,1085414 | -0,5838988 | 0,55928843 |
| 23Sg | -1,1201222 | -189,39501 | -1,1500032 | -0,0001848 | -1,1500032 | -0,5838328 | 0,55933285 |
| VC1484 | 1,5815919 | 544,744678 | 1,77915494 | 0,00053157 | 1,77915494 | 0,58372476 | 0,55940554 |
| VC1306 | -1,0518618 | -3,5713063 | -1,1284755 | -3,485E-06 | -1,1284755 | -0,5832435 | 0,55972945 |
| VCA0527 | 1,1620425 | 2,53069296 | 1,09367564 | 2,4695E-06 | 1,09367564 | 0,58283415 | 0,56000498 |
| VC2731 | 1,15301498 | 2,86412926 | 1,09242314 | 2,7948E-06 | 1,09242314 | 0,58255206 | 0,56019491 |
| VCA1046 | -1,1038787 | -1,9565941 | -1,1816052 | -1,909E-06 | -1,1816052 | -0,5825417 | 0,56020185 |
| VCA0011 | -1,3165827 | -31,661493 | -1,3741844 | -3,09E-05 | -1,3741844 | -0,5822617 | 0,56039043 |
| VCA0075 | 1,22001741 | 2,39176437 | 1,18142056 | 2,3339E-06 | 1,18142056 | 0,58224955 | 0,56039862 |
| VCA0067 | 1,39538315 | 1,11290038 | 1,2255772 | 1,086E-06 | 1,2255772 | 0,58172298 | 0,56075331 |
| VCA0590 | 1,00045884 | -2,5181219 | -1,0935902 | -2,457E-06 | -1,0935902 | -0,5811314 | 0,56115189 |
| VC0813 | 1,31790775 | 3,23447006 | 1,18472316 | 3,1562E-06 | 1,18472316 | 0,58106493 | 0,56119672 |
| VC2096 | -1,0767657 | -7,6486883 | -1,0724584 | -7,464E-06 | -1,0724584 | -0,579678 | 0,56213179 |
| VC1079 | 1,19645281 | 3,91162193 | 1,15510611 | 3,817E-06 | 1,15510611 | 0,57946751 | 0,56227378 |
| VC0227 | 1,13036061 | 11,030273 | 1,16754854 | 1,0763E-05 | 1,16754854 | 0,57921361 | 0,56244507 |

| | | | | | | | |
|---|---|---|---|---|---|---|---|
| VCA0279 | 1,58886479 | 46,5528817 | 1,46331075 | 4,5427E-05 | 1,46331075 | 0,57913639 | 0,56249719 |
| secA | 1,16113771 | 16,8656904 | 1,13900047 | 1,6458E-05 | 1,13900047 | 0,5791195 | 0,56250858 |
| VCA0494 | 1,33424937 | 1,59145089 | 1,23477612 | 1,553E-06 | 1,23477612 | 0,57894434 | 0,56262676 |
| VC1408 | 1,17178015 | 2,11423117 | 1,11145543 | 2,0631E-06 | 1,11145543 | 0,57862978 | 0,56283903 |
| VC2440 | 1,13178339 | 20,8661203 | 1,16115777 | 2,0361E-05 | 1,16115777 | 0,57850136 | 0,56292571 |
| VCA0382 | -1,1131552 | -8,4261441 | -1,1979569 | -8,222E-06 | -1,1979569 | -0,5777155 | 0,56345621 |
| thiP | 1,39886928 | 0,78051486 | 1,33241086 | 7,6163E-07 | 1,33241086 | 0,5776794 | 0,56348061 |
| VC2602 | 1,00356573 | -21,753159 | -1,0568684 | -2,123E-05 | -1,0568684 | -0,5774685 | 0,56362304 |
| VCA0156 | -1,2012126 | -0,8143317 | -1,315991 | -7,946E-07 | -1,315991 | -0,5773389 | 0,56371054 |
| VC2324 | 1,29413778 | 1,24753184 | 1,1953723 | 1,2174E-06 | 1,1953723 | 0,57711897 | 0,56385911 |
| VC2767 | 1,22769103 | 79,5756788 | 1,11009792 | 7,7651E-05 | 1,11009792 | 0,57684549 | 0,56404387 |
| ileS | 1,1337233 | 7,51538292 | 1,12081054 | 7,3336E-06 | 1,12081054 | 0,57677704 | 0,56409011 |
| VCA0286 | 1,14969228 | 3,16100107 | 1,08922692 | 3,0845E-06 | 1,08922692 | 0,57674175 | 0,56411395 |
| mglC | -1,2220067 | -51,628852 | -1,317428 | -5,038E-05 | -1,317428 | -0,5765053 | 0,56427373 |
| VC1257 | 1,13705113 | 3,67738088 | 1,09667678 | 3,5884E-06 | 1,09667678 | 0,57647475 | 0,56429436 |
| VC0538 | 1,52972657 | 13,9331687 | 1,58795482 | 1,3596E-05 | 1,58795482 | 0,57630271 | 0,56441062 |
| VC1048 | 1,18305226 | 6,9477762 | 1,16659476 | 6,7797E-06 | 1,16659476 | 0,57618488 | 0,56449025 |
| gltD (NC_002505 2535637..2537147) | 1,67957073 | 4,65450722 | 1,60576236 | 4,5419E-06 | 1,60576236 | 0,57508352 | 0,56523484 |
| VC2502 | 1,29170871 | 1,34522594 | 1,17815694 | 1,3127E-06 | 1,17815694 | 0,5745354 | 0,56560558 |
| VCA0919 | 1,06704941 | 18,6036376 | 1,09445649 | 1,8154E-05 | 1,09445649 | 0,57379443 | 0,56610695 |
| ligA | 1,19377432 | 2,13058882 | 1,11620547 | 2,079E-06 | 1,11620547 | 0,57352819 | 0,56628714 |
| VCA0878 | -1,0584873 | -1,9324224 | -1,1201976 | -1,886E-06 | -1,1201976 | -0,5732909 | 0,56644777 |
| VC2009 | 1,3218797 | 34,696495 | 1,25697533 | 3,3857E-05 | 1,25697533 | 0,57298365 | 0,5666558 |
| VC2672 | 1,07799631 | 13,3205156 | 1,05003076 | 1,2998E-05 | 1,05003076 | 0,57276542 | 0,56680355 |
| VCA0609 | -1,1306723 | -18,609773 | -1,1712751 | -1,816E-05 | -1,1712751 | -0,5727475 | 0,56681573 |
| VC0002 | 1,05679392 | 6,81630451 | 1,06213445 | 6,6514E-06 | 1,06213445 | 0,57232137 | 0,5671043 |
| nudC | 1,24086601 | 1,52344753 | 1,15438522 | 1,4866E-06 | 1,15438522 | 0,57229219 | 0,56712406 |
| VCA0341 | 1,20050246 | 1,54709291 | 1,16365872 | 1,5097E-06 | 1,16365872 | 0,57153562 | 0,56763664 |
| VCA0003 | -1,2843816 | -12,173121 | -1,257285 | -1,188E-05 | -1,257285 | -0,5711845 | 0,56787465 |
| VC1652 | 1,17679687 | 1,78585737 | 1,1295183 | 1,7427E-06 | 1,1295183 | 0,57083686 | 0,56811025 |
| VC0466 | 1,20270445 | 1,69413129 | 1,13699734 | 1,6531E-06 | 1,13699734 | 0,57080932 | 0,56812893 |
| tRNA-Thr-1 | 1,51623312 | 1,15926166 | 1,45624891 | 1,1312E-06 | 1,45624891 | 0,5707943 | 0,56813911 |
| VC0767 | -1,2141375 | -12,913163 | -1,170568 | -1,26E-05 | -1,170568 | -0,5707722 | 0,56815408 |
| VCA1080 | 1,16476326 | 2,44914851 | 1,09275525 | 2,3899E-06 | 1,09275525 | 0,57066875 | 0,56822422 |
| VCA0936 | 1,27016949 | 1,33673449 | 1,17675307 | 1,3044E-06 | 1,17675307 | 0,57064216 | 0,56824226 |
| VC1579 | 1,24774191 | 116,193449 | 1,13311684 | 0,00011338 | 1,13311684 | 0,57026695 | 0,56849668 |
| VC0619 | -1,4535211 | -2,6814108 | -1,5387015 | -2,617E-06 | -1,5387015 | -0,5698126 | 0,56880484 |
| VCA0469 | 1,14855431 | 4,25180426 | 1,09041328 | 4,1489E-06 | 1,09041328 | 0,56894014 | 0,56939679 |
| VCA0624 | 1,14609956 | 9,58904211 | 1,14600925 | 9,3571E-06 | 1,14600925 | 0,56887585 | 0,56944042 |
| VCA0799 | 1,16346929 | 2,90347569 | 1,07708657 | 2,8332E-06 | 1,07708657 | 0,56857805 | 0,56964255 |
| VC2232 | -1,0155605 | -2,801071 | -1,0798959 | -2,733E-06 | -1,0798959 | -0,5681617 | 0,56992521 |
| VC1658 | 1,29898847 | 1,21182648 | 1,19469677 | 1,1825E-06 | 1,19469677 | 0,56790333 | 0,57010063 |
| tdh | -1,2264908 | -16,694185 | -1,1862497 | -1,629E-05 | -1,1862497 | -0,5675365 | 0,57034978 |
| VCA0651 | -1,237308 | -0,5107433 | -1,5320007 | -4,984E-07 | -1,5320007 | -0,5673962 | 0,57044503 |
| VC1662 | 1,18357458 | 1,95319431 | 1,11617277 | 1,9059E-06 | 1,11617277 | 0,56717064 | 0,57059828 |

| | | | | | | | |
|---|---|---|---|---|---|---|---|
| VC0605 | 1,13723056 | 3,3425363 | 1,09710166 | 3,2617E-06 | 1,09710166 | 0,56694345 | 0,57075262 |
| emrD | -1,0609434 | -1,7707587 | -1,1283868 | -1,728E-06 | -1,1283868 | -0,5660808 | 0,57133889 |
| VC0475 | 1,48818809 | 2,98705977 | 1,40048231 | 2,9148E-06 | 1,40048231 | 0,56569053 | 0,5716042 |
| VC2606 | 1,25174487 | 2,70709497 | 1,14310558 | 2,6416E-06 | 1,14310558 | 0,56485553 | 0,57217205 |
| VC0624 | 1,19660778 | 2,06461132 | 1,1083383 | 2,0147E-06 | 1,1083383 | 0,5641634 | 0,57264295 |
| VC0297 | 1,10709335 | 11,9260126 | 1,1173163 | 1,1638E-05 | 1,1173163 | 0,56369755 | 0,57296002 |
| VCA0441 | 1,21179941 | 8,15679558 | 1,16868751 | 7,9595E-06 | 1,16868751 | 0,56350738 | 0,57308947 |
| VC0113 | 1,17705726 | 2,44508507 | 1,14433072 | 2,3859E-06 | 1,14433072 | 0,56239725 | 0,57384542 |
| VC2541 | -1,0752451 | -1,4674614 | -1,1539683 | -1,432E-06 | -1,1539683 | -0,5609732 | 0,57481582 |
| VC1942 | 1,18263987 | 1,6918734 | 1,13166767 | 1,6509E-06 | 1,13166767 | 0,5599217 | 0,57553286 |
| VCA0522 | -1,1404246 | -8,9942266 | -1,1331519 | -8,777E-06 | -1,1331519 | -0,5588413 | 0,57627004 |
| VCA0436 | 1,23102499 | 3,34812624 | 1,14208856 | 3,2671E-06 | 1,14208856 | 0,55881491 | 0,57628807 |
| VCA0695 | 1,34943753 | 1,51433237 | 1,24318003 | 1,4777E-06 | 1,24318003 | 0,55876739 | 0,57632051 |
| VC0061 | 1,34673976 | 1,06366879 | 1,21621489 | 1,0379E-06 | 1,21621489 | 0,55795846 | 0,57687277 |
| tRNA-Val-1 | -1,1327306 | -3,9666141 | -1,1772475 | -3,871E-06 | -1,1772475 | -0,5578127 | 0,5769723 |
| VC2390 | -1,1704343 | -5,8494846 | -1,170391 | -5,708E-06 | -1,170391 | -0,5575081 | 0,57718034 |
| VCA0831 | 1,18235822 | 2,07821192 | 1,10461911 | 2,0279E-06 | 1,10461911 | 0,55670965 | 0,57772585 |
| VCA0787 | -1,1346154 | -1,4674874 | -1,2762437 | -1,432E-06 | -1,2762437 | -0,5561833 | 0,57808556 |
| VC2519 | 1,15451581 | 2,0615414 | 1,11427834 | 2,0117E-06 | 1,11427834 | 0,55617951 | 0,57808817 |
| nhaB | -1,0776107 | -8,8113497 | -1,1171034 | -8,598E-06 | -1,1171034 | -0,5560474 | 0,57817849 |
| VC0540 | 1,4074404 | 4,41171047 | 1,38586224 | 4,305E-06 | 1,38586224 | 0,55578942 | 0,57835485 |
| VC0376 | -1,0907593 | -1,3695006 | -1,1694012 | -1,336E-06 | -1,1694012 | -0,5555739 | 0,57850221 |
| VC1673 | -1,0763716 | -1,2224255 | -1,1835738 | -1,193E-06 | -1,1835738 | -0,555258 | 0,57871826 |
| VC1786 | -1,1265029 | -0,9613958 | -1,2393389 | -9,381E-07 | -1,2393389 | -0,5552127 | 0,57874917 |
| VC2417 | 1,1635408 | 2,60056246 | 1,08890071 | 2,5377E-06 | 1,08890071 | 0,55465874 | 0,57912812 |
| VC0050 | 1,33084666 | 10,9283287 | 1,25246064 | 1,0664E-05 | 1,25246064 | 0,55431018 | 0,57936661 |
| VCA0348 | 1,16037743 | 2,55379802 | 1,11438959 | 2,492E-06 | 1,11438959 | 0,55416096 | 0,57946872 |
| VC2717 | -1,2950657 | -48,142576 | -1,3287756 | -4,698E-05 | -1,3287756 | -0,5538174 | 0,57970381 |
| VCA0005 | 1,16514006 | 29,692961 | 1,19687276 | 2,8975E-05 | 1,19687276 | 0,55291887 | 0,580319 |
| VC2669 | 1,36165998 | 2,58317356 | 1,23569859 | 2,5207E-06 | 1,23569859 | 0,5523793 | 0,58068854 |
| VCA0570 | -1,0098888 | -38,522112 | -1,1058416 | -3,759E-05 | -1,1058416 | -0,5522742 | 0,58076057 |
| VCA1070 | 1,36424029 | 40,1145983 | 1,30677563 | 3,9144E-05 | 1,30677563 | 0,55203979 | 0,58092112 |
| VCA0793 | 1,21525327 | 1,52717459 | 1,14162271 | 1,4902E-06 | 1,14162271 | 0,55043039 | 0,58202423 |
| VCA0904 | 1,54832664 | 2,44645499 | 1,29598585 | 2,3873E-06 | 1,29598585 | 0,54987994 | 0,58240175 |
| VC2642 | 1,32352934 | 1,03014057 | 1,21656235 | 1,0052E-06 | 1,21656235 | 0,54949216 | 0,58266776 |
| VC0278 | -1,2179802 | -14,54602 | -1,2804241 | -1,419E-05 | -1,2804241 | -0,5491789 | 0,5828827 |
| tRNA-Pro-1 | -1,3251637 | -40,103168 | -1,2926093 | -3,913E-05 | -1,2926093 | -0,5488311 | 0,58312137 |
| VCA0646 | 1,41997534 | 0,70714125 | 1,33092174 | 6,9003E-07 | 1,33092174 | 0,54879867 | 0,58314364 |
| 23Sa | -1,4494092 | -27,810457 | -1,5082188 | -2,714E-05 | -1,5082188 | -0,5482038 | 0,58355198 |
| VC2226 | -1,1871329 | -5,0587875 | -1,2064249 | -4,936E-06 | -1,2064249 | -0,5477202 | 0,58388403 |
| VC1348 | -1,0920604 | -1,1349928 | -1,1930062 | -1,108E-06 | -1,1930062 | -0,5474246 | 0,58408711 |
| VCA0370 | 1,20889023 | 1,50403436 | 1,14222641 | 1,4676E-06 | 1,14222641 | 0,54733013 | 0,58415197 |
| VC2753 | 1,79631176 | 146,398115 | 1,5651558 | 0,00014286 | 1,5651558 | 0,54711117 | 0,5843024 |
| trmB | 1,2301016 | 5,04402027 | 1,19386647 | 4,922E-06 | 1,19386647 | 0,54695701 | 0,5844083 |
| VCA0128 | -1,1701954 | -4,589578 | -1,4332214 | -4,479E-06 | -1,4332214 | -0,5468619 | 0,58447363 |

| | | | | | | | |
|---|---|---|---|---|---|---|---|
| VC0364 | 1,26903579 | 4,72617609 | 1,20019369 | 4,6118E-06 | 1,20019369 | 0,54650805 | 0,58471678 |
| nusA | 1,1300124 | 23,8427666 | 1,1579937 | 2,3266E-05 | 1,1579937 | 0,54562588 | 0,58532317 |
| VC1977 | 1,20453095 | 1,79383738 | 1,11687648 | 1,7504E-06 | 1,11687648 | 0,54509402 | 0,58568887 |
| rbgA | 1,05366296 | 4,74217128 | 1,05848619 | 4,6275E-06 | 1,05848619 | 0,54493394 | 0,58579897 |
| VC0705 | 1,1701738 | 10,9461961 | 1,15611726 | 1,0681E-05 | 1,15611726 | 0,54445583 | 0,58612786 |
| VC0273 | 1,09497718 | 370,174688 | 1,20890868 | 0,00036122 | 1,20890868 | 0,54380178 | 0,58657791 |
| VC0774 | 2,05822384 | 0,40286242 | 1,64226763 | 3,9312E-07 | 1,64226763 | 0,54201189 | 0,58781034 |
| VC0080 | -1,0852563 | -1,4197542 | -1,1478613 | -1,385E-06 | -1,1478613 | -0,5414939 | 0,58816722 |
| VC2535 | 1,09856399 | 5,10976003 | 1,06579364 | 4,9861E-06 | 1,06579364 | 0,54110514 | 0,58843514 |
| VCA0981 | 1,34369173 | 1,07223127 | 1,19859808 | 1,0463E-06 | 1,19859808 | 0,53903968 | 0,5898595 |
| VCA0439 | -1,1270911 | -10,142853 | -1,1824361 | -9,897E-06 | -1,1824361 | -0,5387233 | 0,59007782 |
| VC0384 | 1,56558273 | 7,04029261 | 1,63958937 | 6,87E-06 | 1,63958937 | 0,5380933 | 0,59051267 |
| VCA0614 | 1,24328001 | 1,18976407 | 1,19150295 | 1,161E-06 | 1,19150295 | 0,53808012 | 0,59052175 |
| panB | 1,15062132 | 5,99147138 | 1,12817539 | 5,8465E-06 | 1,12817539 | 0,53805597 | 0,59053843 |
| VCA0587 | -1,0849243 | -1,6875311 | -1,1429538 | -1,647E-06 | -1,1429538 | -0,5376862 | 0,59079376 |
| VCA0140 | -1,0719926 | -1,3906039 | -1,1483294 | -1,357E-06 | -1,1483294 | -0,5366952 | 0,59147817 |
| VC0379a | -1,1414477 | -1,4887753 | -1,1823341 | -1,453E-06 | -1,1823341 | -0,5361067 | 0,5918848 |
| VCA0371 | -1,3228743 | -7,8554522 | -1,2999369 | -7,665E-06 | -1,2999369 | -0,5357134 | 0,5921567 |
| VC2014 | 1,2065325 | 1,52134525 | 1,1341064 | 1,4845E-06 | 1,1341064 | 0,53554234 | 0,59227491 |
| VC0580 | 1,08569297 | 6,07699399 | 1,07289096 | 5,93E-06 | 1,07289096 | 0,53506302 | 0,59260631 |
| VC1984 | 1,28552619 | 3,68039889 | 1,16061365 | 3,5914E-06 | 1,16061365 | 0,53492848 | 0,59269934 |
| rnhB | 1,21256039 | 1,27383027 | 1,16172665 | 1,243E-06 | 1,16172665 | 0,53469809 | 0,59285866 |
| VC2739 | -1,0117701 | -1,8992795 | -1,1055821 | -1,853E-06 | -1,1055821 | -0,5345251 | 0,5929783 |
| VC2757 | -1,0170002 | -1,9539643 | -1,1018956 | -1,907E-06 | -1,1018956 | -0,5330837 | 0,59397568 |
| VCA0848 | -1,0908338 | -1,496053 | -1,1349293 | -1,46E-06 | -1,1349293 | -0,5325962 | 0,59431316 |
| VCA0519 | -1,5102621 | -34,639835 | -1,463899 | -3,38E-05 | -1,463899 | -0,531771 | 0,59488466 |
| VC2141 | -1,0847912 | -65,502033 | -1,1966816 | -6,392E-05 | -1,1966816 | -0,5312105 | 0,59527295 |
| VC1601 | -1,0745072 | -6,849042 | -1,1332898 | -6,683E-06 | -1,1332898 | -0,5306053 | 0,59569234 |
| VC1164 | 1,34555378 | 0,99484797 | 1,22457351 | 9,7078E-07 | 1,22457351 | 0,53006479 | 0,59606703 |
| VC0716 | 1,16306066 | 4,33733638 | 1,13380104 | 4,2324E-06 | 1,13380104 | 0,52993992 | 0,59615361 |
| VC1533 | -1,0788778 | -1,134761 | -1,1795801 | -1,107E-06 | -1,1795801 | -0,5296109 | 0,59638172 |
| VC1066 | -1,0876522 | -7,2756583 | -1,1652283 | -7,1E-06 | -1,1652283 | -0,529405 | 0,59652457 |
| VC0455 | 1,14731495 | 1,97170833 | 1,09933795 | 1,924E-06 | 1,09933795 | 0,52905741 | 0,59676564 |
| VC0769 | -1,2011315 | -2,3209836 | -1,3131253 | -2,265E-06 | -1,3131253 | -0,5289705 | 0,59682596 |
| VCA0970 | 1,18288973 | 8,26621735 | 1,11527206 | 8,0662E-06 | 1,11527206 | 0,52871579 | 0,59700264 |
| VCA0805 | 1,15092149 | 2,52572391 | 1,08576661 | 2,4646E-06 | 1,08576661 | 0,52798825 | 0,59750751 |
| VC2112 | 1,19010198 | 2,86640448 | 1,12859132 | 2,7971E-06 | 1,12859132 | 0,52737301 | 0,59793461 |
| VC2002 | -1,0158596 | -2,3423883 | -1,0821862 | -2,286E-06 | -1,0821862 | -0,5266669 | 0,59842496 |
| VC0251 | 1,20753357 | 76,5079864 | 1,15680813 | 7,4657E-05 | 1,15680813 | 0,52664421 | 0,59844072 |
| hutH | -1,1232138 | -1,7162431 | -1,2681243 | -1,675E-06 | -1,2681243 | -0,5263943 | 0,59861431 |
| VCA0144 | -1,0705729 | -1,7198521 | -1,169803 | -1,678E-06 | -1,169803 | -0,5253858 | 0,59931505 |
| VC1402 | 1,36594278 | 0,87118914 | 1,23599181 | 8,5011E-07 | 1,23599181 | 0,52520772 | 0,59943882 |
| VCA0837 | -1,021629 | -2,2805429 | -1,083639 | -2,225E-06 | -1,083639 | -0,5240574 | 0,60023865 |
| VC1949 | -1,2044104 | -0,7854702 | -1,2631093 | -7,665E-07 | -1,2631093 | -0,5234094 | 0,60068941 |
| gltX | 1,11424893 | 9,23377745 | 1,14255564 | 9,0104E-06 | 1,14255564 | 0,52330218 | 0,60076402 |

| | | | | | | | |
|---|---|---|---|---|---|---|---|
| VC0503 | -1,0617672 | -3,6501827 | -1,0943 | -3,562E-06 | -1,0943 | -0,5228937 | 0,60104824 |
| VC2438 | -1,026409 | -1,9987909 | -1,1078891 | -1,95E-06 | -1,1078891 | -0,5228562 | 0,60107432 |
| VC2335 | 1,0080738 | -2,3570886 | -1,0803359 | -2,3E-06 | -1,0803359 | -0,5225685 | 0,60127459 |
| VC1631 | -1,1850151 | -4,5621294 | -1,2639016 | -4,452E-06 | -1,2639016 | -0,5220297 | 0,60164968 |
| VCA0473 | 1,20194344 | 2,06207888 | 1,09354519 | 2,0122E-06 | 1,09354519 | 0,52193318 | 0,60171688 |
| rlmL | 1,14052016 | 3,14771739 | 1,09668259 | 3,0716E-06 | 1,09668259 | 0,5219224 | 0,60172438 |
| VC1956 | 1,09971207 | 4,83550867 | 1,06099299 | 4,7185E-06 | 1,06099299 | 0,52163533 | 0,60192428 |
| VC2683 | 1,32427236 | 1,02176044 | 1,19467989 | 9,9704E-07 | 1,19467989 | 0,52144842 | 0,60205445 |
| VCA0822 | -1,1411483 | -1,1795563 | -1,1990913 | -1,151E-06 | -1,1990913 | -0,5209172 | 0,6024245 |
| VC0371 | 1,12029557 | 3,00071321 | 1,06210971 | 2,9281E-06 | 1,06210971 | 0,52072366 | 0,60255931 |
| VC1973 | -1,1997116 | -56,804449 | -1,2185553 | -5,543E-05 | -1,2185553 | -0,5204018 | 0,60278356 |
| VCA0738 | -1,0775975 | -1,5486753 | -1,1235015 | -1,511E-06 | -1,1235015 | -0,5198205 | 0,60318869 |
| VC0934 | 1,75763975 | 0,40683695 | 1,56811638 | 3,97E-07 | 1,56811638 | 0,51961578 | 0,60333141 |
| VC2301 | -1,2372465 | -18,193329 | -1,2386661 | -1,775E-05 | -1,2386661 | -0,5193096 | 0,60354487 |
| VCA0931 | 1,28813167 | 1,08407829 | 1,18048993 | 1,0579E-06 | 1,18048993 | 0,51884996 | 0,60386539 |
| VC1106 | 1,1626645 | 1,56997958 | 1,12086777 | 1,532E-06 | 1,12086777 | 0,51809454 | 0,60439233 |
| VCA0339 | 1,1928937 | 1,63551102 | 1,11561455 | 1,5959E-06 | 1,11561455 | 0,51781964 | 0,60458413 |
| VC1108 | 1,1667224 | 2,64547629 | 1,10600658 | 2,5815E-06 | 1,10600658 | 0,51758769 | 0,604746 |
| VC0163 | 1,14751758 | 9,42643721 | 1,17162828 | 9,1984E-06 | 1,17162828 | 0,51677375 | 0,60531414 |
| VCA0768 | 1,24958741 | 6,8932312 | 1,18072934 | 6,7265E-06 | 1,18072934 | 0,5166425 | 0,60540577 |
| VC1890 | 1,25882079 | 9,10614131 | 1,26802908 | 8,8859E-06 | 1,26802908 | 0,51660215 | 0,60543395 |
| VC0609 | 1,26144055 | 1,54850742 | 1,17657216 | 1,511E-06 | 1,17657216 | 0,51639645 | 0,60557757 |
| VC1320 | 1,36567341 | 162,078762 | 1,29688027 | 0,00015816 | 1,29688027 | 0,51631458 | 0,60563474 |
| mazG | 1,09027279 | 3,80499191 | 1,07294656 | 3,7129E-06 | 1,07294656 | 0,51626885 | 0,60566667 |
| VC1998 | 1,08429895 | 4,18733752 | 1,08185109 | 4,086E-06 | 1,08185109 | 0,51592958 | 0,60590362 |
| VC1224 | 1,22377162 | 11,0100574 | 1,15988255 | 1,0744E-05 | 1,15988255 | 0,51579582 | 0,60599705 |
| VC2101 | -1,5375376 | -4,6877914 | -1,4807288 | -4,574E-06 | -1,4807288 | -0,5153319 | 0,60632114 |
| VCA0952 | -1,1027853 | -5,3547005 | -1,129132 | -5,225E-06 | -1,129132 | -0,5153159 | 0,60633232 |
| VCA0671 | 1,23927459 | 1,26484944 | 1,15031334 | 1,2343E-06 | 1,15031334 | 0,51502684 | 0,60653428 |
| VC2523 | 1,16957652 | 2,4745798 | 1,10786588 | 2,4147E-06 | 1,10786588 | 0,51497013 | 0,60657392 |
| VC0662 | 1,2827213 | 3,17317085 | 1,23296572 | 3,0964E-06 | 1,23296572 | 0,51418159 | 0,60712507 |
| VCA0591 | -1,0017911 | -3,4729749 | -1,0847347 | -3,389E-06 | -1,0847347 | -0,5141502 | 0,60714699 |
| VC0517 | -1,0516966 | -8,2059159 | -1,0365453 | -8,007E-06 | -1,0365453 | -0,5134004 | 0,60767127 |
| VCA0978 | 1,23233897 | 1,3228566 | 1,14220545 | 1,2909E-06 | 1,14220545 | 0,51327095 | 0,60776182 |
| VC1280 | 1,32633508 | 1,61040319 | 1,17294117 | 1,5714E-06 | 1,17294117 | 0,51284447 | 0,60806014 |
| VC0597 | 1,38199735 | 0,90121916 | 1,21549946 | 8,7942E-07 | 1,21549946 | 0,51281898 | 0,60807797 |
| thiH | 1,27437258 | 1,09964056 | 1,17297753 | 1,073E-06 | 1,17297753 | 0,51245378 | 0,60833348 |
| VC0690 | -1,0388205 | -3,8158669 | -1,1282096 | -3,724E-06 | -1,1282096 | -0,5123579 | 0,60840061 |
| VC0904 | 1,22890546 | 1,29739813 | 1,14407508 | 1,266E-06 | 1,14407508 | 0,51141535 | 0,60906027 |
| VCA0846 | 1,17513479 | 2,01491315 | 1,09041763 | 1,9662E-06 | 1,09041763 | 0,51133294 | 0,60911796 |
| VC2311 | 1,27132791 | 2,1243374 | 1,19793871 | 2,0729E-06 | 1,19793871 | 0,5111834 | 0,60922267 |
| VC0626 | 1,15742164 | 4,8997408 | 1,11803161 | 4,7812E-06 | 1,11803161 | 0,51107866 | 0,609296 |
| VC1022 | 1,14868634 | 2,3777124 | 1,07594391 | 2,3202E-06 | 1,07594391 | 0,51084111 | 0,60946234 |
| VC1530 | -1,0786059 | -1,5539095 | -1,1615777 | -1,516E-06 | -1,1615777 | -0,5100558 | 0,61001239 |
| VC1398 | 1,36558798 | 0,63312767 | 1,31729641 | 6,1781E-07 | 1,31729641 | 0,50997461 | 0,61006927 |

| | | | | | | | |
|---|---|---|---|---|---|---|---|
| VCA0424 | 1,16256855 | 2,50259013 | 1,09929144 | 2,442E-06 | 1,09929144 | 0,50986173 | 0,61014835 |
| VCA0291 | 1,17724094 | 2,25683308 | 1,11534723 | 2,2022E-06 | 1,11534723 | 0,50985798 | 0,61015098 |
| VCA0176 | -1,1250795 | -7,4781873 | -1,1921762 | -7,297E-06 | -1,1921762 | -0,5098353 | 0,61016691 |
| lldD | -1,2046369 | -2,3590684 | -1,3566461 | -2,302E-06 | -1,3566461 | -0,5096195 | 0,61031806 |
| VC1240 | -1,1124016 | -1,9182758 | -1,1833171 | -1,872E-06 | -1,1833171 | -0,5090385 | 0,61072524 |
| VC0405 | -1,0360651 | -3,2700889 | -1,0542326 | -3,191E-06 | -1,0542326 | -0,5089311 | 0,61080055 |
| VC0923 | 1,20768794 | 1,49980624 | 1,1220962 | 1,4635E-06 | 1,1220962 | 0,50880283 | 0,61089046 |
| VCA0925 | 1,16453773 | 1,8716331 | 1,0965926 | 1,8264E-06 | 1,0965926 | 0,5086162 | 0,61102128 |
| VC1193 | -1,2691569 | -11,724329 | -1,3385392 | -1,144E-05 | -1,3385392 | -0,5085468 | 0,61106997 |
| VC2350 | -1,2317285 | -24,460645 | -1,2389021 | -2,387E-05 | -1,2389021 | -0,5084772 | 0,61111877 |
| VC1088 | -1,0567391 | -1,1178402 | -1,1665311 | -1,091E-06 | -1,1665311 | -0,5077118 | 0,61165547 |
| VCA0202 | -1,2741536 | -0,5069238 | -1,4062658 | -4,947E-07 | -1,4062658 | -0,5067172 | 0,61235329 |
| VC0830 | 1,30224159 | 3,15543055 | 1,26219033 | 3,0791E-06 | 1,26219033 | 0,5065738 | 0,61245392 |
| VCA0035 | 1,25301892 | 1,15576745 | 1,1598328 | 1,1278E-06 | 1,1598328 | 0,50654777 | 0,61247218 |
| VCA1053 | 1,22213212 | 1,38486297 | 1,13164682 | 1,3514E-06 | 1,13164682 | 0,50654027 | 0,61247744 |
| VC1061 | 1,42358877 | 0,69967773 | 1,27807962 | 6,8275E-07 | 1,27807962 | 0,50618632 | 0,61272588 |
| argD | 1,77439135 | 7,95439589 | 1,52528739 | 7,762E-06 | 1,52528739 | 0,50609658 | 0,61278887 |
| VCA0115 | 1,42842272 | 0,7269339 | 1,26590217 | 7,0935E-07 | 1,26590217 | 0,50588187 | 0,6129396 |
| VC0135 | 1,29736466 | 2,64579829 | 1,15511342 | 2,5818E-06 | 1,15511342 | 0,50489057 | 0,61363572 |
| VC1495 | -1,0888104 | -0,9119341 | -1,2052801 | -8,899E-07 | -1,2052801 | -0,5046439 | 0,61380901 |
| ihfB | 1,03231446 | -82,35587 | -1,0897609 | -8,036E-05 | -1,0897609 | -0,5043741 | 0,61399853 |
| VC1762 | -1,1855989 | -9,5657936 | -1,1558064 | -9,334E-06 | -1,1558064 | -0,5035357 | 0,61458772 |
| VC1929 | 1,72201663 | 2,64210564 | 1,663657 | 2,5782E-06 | 1,663657 | 0,50334723 | 0,6147202 |
| sdhA | 1,53790308 | 33,1196621 | 1,45974544 | 3,2318E-05 | 1,45974544 | 0,50315308 | 0,61485668 |
| treR | 1,1391213 | 2,86891596 | 1,08428318 | 2,7995E-06 | 1,08428318 | 0,50240625 | 0,6153818 |
| VC0075 | -1,1072995 | -8,9976665 | -1,1525175 | -8,78E-06 | -1,1525175 | -0,5022296 | 0,61550607 |
| VC0361 | 1,10013284 | 21,8852483 | 1,13721912 | 2,1356E-05 | 1,13721912 | 0,50218122 | 0,61554007 |
| VC1893 | -1,0375064 | -3,1795216 | -1,0542315 | -3,103E-06 | -1,0542315 | -0,5018288 | 0,61578797 |
| VC2240 | 1,37408557 | 14,184791 | 1,25188798 | 1,3842E-05 | 1,25188798 | 0,50139114 | 0,61609591 |
| VC2761 | 1,45668041 | 7,31760938 | 1,47663518 | 7,1406E-06 | 1,47663518 | 0,50119005 | 0,61623741 |
| VC2612 | 1,2328617 | 1,14308289 | 1,15753156 | 1,1154E-06 | 1,15753156 | 0,50038741 | 0,61680234 |
| murG | -1,0354416 | -2,2730776 | -1,0916347 | -2,218E-06 | -1,0916347 | -0,4991357 | 0,61768385 |
| VCA0318 | 1,17452428 | 3,10058128 | 1,12109147 | 3,0256E-06 | 1,12109147 | 0,49905137 | 0,61774322 |
| VC2108 | 1,19849529 | 3,985866 | 1,15720561 | 3,8894E-06 | 1,15720561 | 0,49895183 | 0,61781334 |
| VC0071 | 1,12043466 | 8,64737184 | 1,10388584 | 8,4382E-06 | 1,10388584 | 0,49855263 | 0,61809461 |
| VC0010 | 1,37409237 | 20,5382804 | 1,42702482 | 2,0041E-05 | 1,42702482 | 0,49835538 | 0,61823361 |
| VC0747 | 1,12105023 | 5,28430606 | 1,10659989 | 5,1565E-06 | 1,10659989 | 0,49825264 | 0,61830601 |
| VCA0083 | 1,25074688 | 1,08171894 | 1,16550745 | 1,0556E-06 | 1,16550745 | 0,49802172 | 0,61846875 |
| VC1091 | 1,11928667 | 21,4104845 | 1,06816301 | 2,0893E-05 | 1,06816301 | 0,49798177 | 0,61849692 |
| VCA0116 | 1,32876987 | 0,76608196 | 1,24141169 | 7,4755E-07 | 1,24141169 | 0,49752779 | 0,61881693 |
| VC1127 | 1,20768092 | 2,63035845 | 1,15441942 | 2,5667E-06 | 1,15441942 | 0,49693363 | 0,61923588 |
| VC2480 | 1,11708227 | 2,65650814 | 1,06393902 | 2,5922E-06 | 1,06393902 | 0,49688956 | 0,61926696 |
| VC0004 | 1,10174688 | 10,6195105 | 1,08904069 | 1,0363E-05 | 1,08904069 | 0,49688301 | 0,61927157 |
| VC0090 | -1,0692174 | -1,1193279 | -1,1586149 | -1,092E-06 | -1,1586149 | -0,4967356 | 0,61937553 |
| VCA0881 | 1,42991135 | 0,71674245 | 1,25905808 | 6,994E-07 | 1,25905808 | 0,49656686 | 0,61949454 |

| | | | | | | | |
|---|---|---|---|---|---|---|---|
| VCA0853 | 1,48191396 | 0,48249043 | 1,40965218 | 4,7082E-07 | 1,40965218 | 0,49606175 | 0,61985086 |
| VCA0111 | 1,45731859 | 0,59357915 | 1,31947802 | 5,7922E-07 | 1,31947802 | 0,49525147 | 0,62042264 |
| VC1232 | 1,1682379 | 1,50573214 | 1,11482681 | 1,4693E-06 | 1,11482681 | 0,49524679 | 0,62042594 |
| rseB | 1,0861122 | 16,8491395 | 1,06954948 | 1,6442E-05 | 1,06954948 | 0,49489356 | 0,62067529 |
| VC1286 | -1,0312093 | -5,3655988 | -1,1257055 | -5,236E-06 | -1,1257055 | -0,4948719 | 0,62069061 |
| VC1697 | -1,0530242 | -6,325184 | -1,1860217 | -6,172E-06 | -1,1860217 | -0,4946982 | 0,62081317 |
| VCA0379 | 1,24641725 | 1,17924741 | 1,14856293 | 1,1507E-06 | 1,14856293 | 0,49459088 | 0,62088896 |
| VC1008 | -1,0558742 | -8,1217489 | -1,1012321 | -7,925E-06 | -1,1012321 | -0,4939169 | 0,62136491 |
| VCA0157 | -1,0728208 | -0,9515399 | -1,1868378 | -9,285E-07 | -1,1868378 | -0,493855 | 0,62140861 |
| VC0076 | -1,2102105 | -74,924226 | -1,2563315 | -7,311E-05 | -1,2563315 | -0,493749 | 0,62148351 |
| VCA0862 | -1,0314268 | -1,1444885 | -1,1932749 | -1,117E-06 | -1,1932749 | -0,4937361 | 0,62149257 |
| VCA0454 | 1,3800653 | 1,72216792 | 1,37501486 | 1,6805E-06 | 1,37501486 | 0,493447 | 0,6216968 |
| VCA0064 | 1,51313331 | 0,4819646 | 1,40485642 | 4,7031E-07 | 1,40485642 | 0,49337194 | 0,62174982 |
| VCA0132 | -1,065846 | -2,4100069 | -1,1494497 | -2,352E-06 | -1,1494497 | -0,4933101 | 0,62179349 |
| rph | -1,108632 | -10,134042 | -1,0982148 | -9,889E-06 | -1,0982148 | -0,4933023 | 0,62179907 |
| VCA0661 | 1,00610793 | -2,4762664 | -1,0675823 | -2,416E-06 | -1,0675823 | -0,4927792 | 0,62216863 |
| prfC | 1,11843414 | 3,10413342 | 1,08854723 | 3,029E-06 | 1,08854723 | 0,49206023 | 0,62267678 |
| tRNA-Glu-2 | 1,1771338 | 23,4658176 | 1,23670935 | 2,2898E-05 | 1,23670935 | 0,49180304 | 0,62285862 |
| VC1447 | 1,38124071 | 0,61218761 | 1,30320483 | 5,9738E-07 | 1,30320483 | 0,49170562 | 0,62292748 |
| VC1892 | 1,03324029 | 5,62156297 | 1,03771896 | 5,4856E-06 | 1,03771896 | 0,49145758 | 0,62310287 |
| VC1950 | 1,21008879 | 15,9105588 | 1,23409564 | 1,5526E-05 | 1,23409564 | 0,49135698 | 0,62317401 |
| VC1283 | 1,39933818 | 0,91789429 | 1,19588208 | 8,9569E-07 | 1,19588208 | 0,49049529 | 0,62378348 |
| VC2264 | -1,0732304 | -6,6639259 | -1,1044243 | -6,503E-06 | -1,1044243 | -0,490442 | 0,62382121 |
| VC0158 | 1,07052643 | 4,07726566 | 1,05673264 | 3,9786E-06 | 1,05673264 | 0,49011488 | 0,62405263 |
| VC0073 | -1,0457803 | -1,9037631 | -1,0915949 | -1,858E-06 | -1,0915949 | -0,4899594 | 0,62416265 |
| VC0152 | 1,14739936 | 2,27300725 | 1,07360522 | 2,218E-06 | 1,07360522 | 0,48981906 | 0,62426196 |
| VCA0838 | -1,043064 | -3,4524946 | -1,0785524 | -3,369E-06 | -1,0785524 | -0,48923 | 0,62467887 |
| VC1482 | 1,11765284 | 2,40365485 | 1,06839861 | 2,3455E-06 | 1,06839861 | 0,4883272 | 0,62531812 |
| VC0063 | 1,31261756 | 0,71760383 | 1,24828624 | 7,0024E-07 | 1,24828624 | 0,48758936 | 0,62584075 |
| VC1888 | 1,18626685 | 1,4939318 | 1,1119526 | 1,4578E-06 | 1,1119526 | 0,48742076 | 0,6259602 |
| VC2239 | 1,11098036 | 5,35037889 | 1,0617512 | 5,2209E-06 | 1,0617512 | 0,48677721 | 0,62641625 |
| VC1386 | 1,3678315 | 0,76756908 | 1,22931881 | 7,49E-07 | 1,22931881 | 0,48669166 | 0,62647688 |
| VC1945 | 1,37010619 | 0,80033083 | 1,21829855 | 7,8097E-07 | 1,21829855 | 0,48608441 | 0,62690733 |
| VCA0022 | 1,18479303 | 1,29823998 | 1,1426018 | 1,2668E-06 | 1,1426018 | 0,48606442 | 0,62692151 |
| ihfA | 1,02087108 | -60,590614 | -1,1257412 | -5,912E-05 | -1,1257412 | -0,4856731 | 0,62719896 |
| mobA | -1,0620104 | -3,5290045 | -1,1463366 | -3,444E-06 | -1,1463366 | -0,4854995 | 0,62732213 |
| VC0866 | 1,15479471 | 2,11823741 | 1,07703267 | 2,067E-06 | 1,07703267 | 0,48547829 | 0,62733713 |
| VC2065 | 1,11771258 | 28,9559141 | 1,05882445 | 2,8255E-05 | 1,05882445 | 0,48505627 | 0,62763646 |
| VC2431 | -1,0090475 | -2,8210589 | -1,0767523 | -2,753E-06 | -1,0767523 | -0,4844488 | 0,6280674 |
| nusB | -1,1646506 | -31,610923 | -1,1516081 | -3,085E-05 | -1,1516081 | -0,4840675 | 0,628338 |
| VCA0644 | -1,0351641 | -0,8070621 | -1,2136635 | -7,875E-07 | -1,2136635 | -0,4834197 | 0,62879776 |
| VC1242 | 1,32633768 | 11,6445588 | 1,22295494 | 1,1363E-05 | 1,22295494 | 0,4833257 | 0,62886453 |
| moaA | 1,14667264 | 2,20381048 | 1,09628272 | 2,1505E-06 | 1,09628272 | 0,48206383 | 0,62976062 |
| VC1763 | -1,217015 | -12,110657 | -1,1748442 | -1,182E-05 | -1,1748442 | -0,4820293 | 0,62978515 |
| VC2528 | -1,0824164 | -7,6991261 | -1,1063014 | -7,513E-06 | -1,1063014 | -0,4819629 | 0,62983232 |

| | | | | | | | |
|---|---|---|---|---|---|---|---|
| holA | 1,14511935 | 2,37844982 | 1,06720107 | 2,3209E-06 | 1,06720107 | 0,48162842 | 0,63006995 |
| VC0173 | 1,2231402 | 1,99293694 | 1,1617035 | 1,9447E-06 | 1,1617035 | 0,48155138 | 0,63012469 |
| VCA0533 | 1,18407749 | 6,55086887 | 1,1514961 | 6,3924E-06 | 1,1514961 | 0,48136219 | 0,63025913 |
| VC1208 | 1,12717608 | 26,2853164 | 1,12409068 | 2,5649E-05 | 1,12409068 | 0,48086907 | 0,63060957 |
| VCA0568 | -1,0578275 | -7,3940691 | -1,0555538 | -7,215E-06 | -1,0555538 | -0,4806148 | 0,63079032 |
| tRNA-Gln-4 | -1,2435659 | -3,3099263 | -1,2965371 | -3,23E-06 | -1,2965371 | -0,4805882 | 0,63080925 |
| VCA0503 | 1,14267654 | 1,91750001 | 1,08360791 | 1,8711E-06 | 1,08360791 | 0,48045096 | 0,63090678 |
| VC1000 | 1,08232429 | 9,09445189 | 1,08772923 | 8,8744E-06 | 1,08772923 | 0,48020548 | 0,63108132 |
| VC2640 | 1,08896908 | 5,35874195 | 1,06398447 | 5,2291E-06 | 1,06398447 | 0,47970999 | 0,63143364 |
| rpmH | 1,11194778 | 23,4743196 | 1,09482132 | 2,2906E-05 | 1,09482132 | 0,47938517 | 0,63166467 |
| VC1346 | 1,26684568 | 1,04823405 | 1,15735214 | 1,0229E-06 | 1,15735214 | 0,47892455 | 0,63199232 |
| VCA0270 | 1,00103855 | -2,6766905 | -1,1218253 | -2,612E-06 | -1,1218253 | -0,478884 | 0,63202114 |
| VC2513 | 1,15485666 | 1,95209489 | 1,08137607 | 1,9049E-06 | 1,08137607 | 0,47850828 | 0,6322885 |
| VC1785 | 1,45150325 | 0,62201502 | 1,2788142 | 6,0697E-07 | 1,2788142 | 0,4778203 | 0,63277813 |
| VCA1078 | -1,0594311 | -4,0730728 | -1,0781192 | -3,975E-06 | -1,0781192 | -0,4778111 | 0,63278469 |
| pdxA | 1,14214404 | 3,53129313 | 1,08346982 | 3,4459E-06 | 1,08346982 | 0,47674866 | 0,63354113 |
| VC1805 | 1,29283708 | 0,95161659 | 1,17226635 | 9,286E-07 | 1,17226635 | 0,47581332 | 0,63420739 |
| VC1974 | -1,1323819 | -9,1319445 | -1,1448404 | -8,911E-06 | -1,1448404 | -0,4746364 | 0,63504617 |
| VCA0006 | 1,16227684 | 34,1153262 | 1,15695105 | 3,329E-05 | 1,15695105 | 0,4733252 | 0,63598121 |
| VC0183 | 1,20552467 | 2,71997797 | 1,1405921 | 2,6542E-06 | 1,1405921 | 0,47314944 | 0,63610659 |
| VC1121 | -1,0153382 | -1,540004 | -1,122388 | -1,503E-06 | -1,122388 | -0,4729453 | 0,6362522 |
| VC0900 | 1,13483597 | 9,18065029 | 1,09212398 | 8,9586E-06 | 1,09212398 | 0,47294493 | 0,6362525 |
| VC2554 | 1,27075037 | 1,04435826 | 1,15345242 | 1,0191E-06 | 1,15345242 | 0,47250474 | 0,63656658 |
| VC2368 | -1,0096791 | -157,58987 | -1,2105013 | -0,0001538 | -1,2105013 | -0,471666 | 0,63716523 |
| VC2187 | -1,4074653 | -198,89521 | -1,3555544 | -0,0001941 | -1,3555544 | -0,4715825 | 0,63722486 |
| 16Sb | 1,26242606 | 5,97313521 | 1,20751806 | 5,8286E-06 | 1,20751806 | 0,47095496 | 0,63767292 |
| fliE | -1,0804483 | -17,158557 | -1,0841568 | -1,674E-05 | -1,0841568 | -0,4707357 | 0,63782951 |
| VC1506 | -1,0209033 | -0,8922967 | -1,1802824 | -8,707E-07 | -1,1802824 | -0,4704752 | 0,63801558 |
| VC0491 | 1,27860791 | 50,0320812 | 1,26759751 | 4,8822E-05 | 1,26759751 | 0,47029941 | 0,63814113 |
| VCA0344 | 1,3966742 | 123,291547 | 1,31248344 | 0,00012031 | 1,31248344 | 0,47017893 | 0,63822721 |
| VC1089 | 1,34865334 | 0,70369895 | 1,23360089 | 6,8668E-07 | 1,23360089 | 0,46988249 | 0,63843899 |
| VC1733 | 1,62302553 | 0,43833587 | 1,40325043 | 4,2773E-07 | 1,40325043 | 0,46973435 | 0,63854484 |
| VCA1041 | -1,1618386 | -10,673974 | -1,2221435 | -1,042E-05 | -1,2221435 | -0,4694514 | 0,63874702 |
| VC1939 | 1,20446498 | 1,7769085 | 1,08991543 | 1,7339E-06 | 1,08991543 | 0,46801191 | 0,63977609 |
| VC2610 | 1,17095052 | 1,6071785 | 1,09514698 | 1,5683E-06 | 1,09514698 | 0,46793625 | 0,63983019 |
| rpsT | 1,11460057 | 207,699508 | 1,16060898 | 0,00020267 | 1,16060898 | 0,46672347 | 0,64069776 |
| VC1492 | -1,2926123 | -20,661779 | -1,3106929 | -2,016E-05 | -1,3106929 | -0,4666266 | 0,64076709 |
| VC1708 | 1,28954578 | 0,83583798 | 1,1900582 | 8,1562E-07 | 1,1900582 | 0,46648554 | 0,64086801 |
| VC2265 | 1,01003464 | -2,1889658 | -1,0683932 | -2,136E-06 | -1,0683932 | -0,4659906 | 0,64122226 |
| VCA0893 | 1,09771915 | 2,78491562 | 1,05303059 | 2,7175E-06 | 1,05303059 | 0,46456192 | 0,64224522 |
| VC0415 | -1,0625478 | -10,555119 | -1,0876126 | -1,03E-05 | -1,0876126 | -0,4643594 | 0,64239032 |
| VC2028 | 1,23069516 | 5,58847727 | 1,19655731 | 5,4533E-06 | 1,19655731 | 0,46426853 | 0,6424554 |
| uvrD | 1,14412757 | 2,01605449 | 1,073831 | 1,9673E-06 | 1,073831 | 0,46403471 | 0,64262289 |
| VC1345 | 1,28443601 | 0,86682761 | 1,18041307 | 8,4586E-07 | 1,18041307 | 0,4638662 | 0,64274362 |
| VC1702 | 1,28256816 | 3,33132513 | 1,1379815 | 3,2507E-06 | 1,1379815 | 0,46328969 | 0,64315676 |

| | | | | | | | |
|---|---|---|---|---|---|---|---|
| VCA1002 | -1,135609 | -1,557575 | -1,2209691 | -1,52E-06 | -1,2209691 | -0,4630723 | 0,64331253 |
| VCA0407 | -1,2097606 | -0,5664882 | -1,2885679 | -5,528E-07 | -1,2885679 | -0,4629123 | 0,64342726 |
| VC1611 | 1,3938181 | 0,72670677 | 1,21731046 | 7,0913E-07 | 1,21731046 | 0,4622406 | 0,64390881 |
| VCA0496 | 1,15213342 | 5,25559577 | 1,13578883 | 5,1285E-06 | 1,13578883 | 0,46169681 | 0,64429878 |
| VC0130 | -1,0561446 | -3,8336709 | -1,1550476 | -3,741E-06 | -1,1550476 | -0,4616116 | 0,64435992 |
| VC2424 | -1,0811767 | -1,0379315 | -1,1467946 | -1,013E-06 | -1,1467946 | -0,4614306 | 0,64448973 |
| VCA0105 | 1,23054206 | 2,23375781 | 1,1198467 | 2,1797E-06 | 1,1198467 | 0,46139211 | 0,64451733 |
| VC1986 | 1,15568491 | 1,49596817 | 1,09949302 | 1,4598E-06 | 1,09949302 | 0,46117339 | 0,64467423 |
| tppB | 1,25759946 | 7,55553077 | 1,26517437 | 7,3728E-06 | 1,26517437 | 0,46094898 | 0,64483524 |
| VCA0154 | -1,0697177 | -0,8045033 | -1,1919777 | -7,85E-07 | -1,1919777 | -0,4597618 | 0,64568724 |
| pyrB | 1,16739715 | 2,75310055 | 1,13009572 | 2,6865E-06 | 1,13009572 | 0,4596076 | 0,64579792 |
| VC1861 | 1,3664517 | 0,72422955 | 1,21499933 | 7,0671E-07 | 1,21499933 | 0,45923111 | 0,64606823 |
| VC1900 | -1,08331 | -5,9197653 | -1,0569694 | -5,777E-06 | -1,0569694 | -0,4590401 | 0,64620536 |
| VC0108 | 1,14559921 | 1,79726182 | 1,08123739 | 1,7538E-06 | 1,08123739 | 0,45876297 | 0,64640441 |
| VC1796 | 1,29037282 | 1,33926092 | 1,21930901 | 1,3069E-06 | 1,21930901 | 0,45869232 | 0,64645515 |
| VC2346 | 1,17371126 | 11,1984745 | 1,16774183 | 1,0928E-05 | 1,16774183 | 0,45838263 | 0,6466776 |
| VC0647 | 1,06090561 | 14,2293467 | 1,07489612 | 1,3885E-05 | 1,07489612 | 0,4581263 | 0,64686172 |
| VC1423 | -1,0288591 | -4,736064 | -1,0476844 | -4,621E-06 | -1,0476844 | -0,4574663 | 0,64733593 |
| VC1400 | -1,0811922 | -1,0349879 | -1,1439945 | -1,01E-06 | -1,1439945 | -0,456658 | 0,64791692 |
| VCA0834 | -1,0421837 | -1,3939249 | -1,1048733 | -1,36E-06 | -1,1048733 | -0,4564602 | 0,64805914 |
| VC0635 | -1,0013546 | -60,811542 | -1,1316949 | -5,934E-05 | -1,1316949 | -0,4564119 | 0,64809383 |
| VCA1052 | -1,0128884 | -0,931375 | -1,1609901 | -9,088E-07 | -1,1609901 | -0,4562442 | 0,64821437 |
| VCA0759 | 1,58919304 | 5,83826401 | 1,47078592 | 5,697E-06 | 1,47078592 | 0,45545754 | 0,64878013 |
| VC0600 | 1,46249387 | 0,50183005 | 1,31949076 | 4,8969E-07 | 1,31949076 | 0,45537814 | 0,64883723 |
| VC0022 | 1,11835771 | 5,12705919 | 1,09855893 | 5,003E-06 | 1,09855893 | 0,4543707 | 0,64956206 |
| VC1296 | -1,0372641 | -1,6271095 | -1,0982283 | -1,588E-06 | -1,0982283 | -0,4541599 | 0,64971374 |
| VC1397 | 1,2501137 | 0,88541429 | 1,1679616 | 8,6399E-07 | 1,1679616 | 0,45364311 | 0,65008574 |
| VC1748 | 1,31901491 | 0,6050863 | 1,25559534 | 5,9045E-07 | 1,25559534 | 0,45354014 | 0,65015986 |
| tRNA-Asn-1 | 1,05935363 | 9,69132706 | 1,09410468 | 9,4569E-06 | 1,09410468 | 0,45320786 | 0,65039909 |
| VC2451 | -1,028409 | -3,0541156 | -1,0457058 | -2,98E-06 | -1,0457058 | -0,4524616 | 0,65093652 |
| VC2426 | 1,25862708 | 0,96077532 | 1,15282107 | 9,3753E-07 | 1,15282107 | 0,45233547 | 0,65102735 |
| tRNA-Pro-3 | -1,1212066 | -5,0369084 | -1,1316 | -4,915E-06 | -1,1316 | -0,4521173 | 0,65118449 |
| VCA0247 | 1,27755797 | 3,43858986 | 1,15526585 | 3,3554E-06 | 1,15526585 | 0,4514704 | 0,65165057 |
| VC0917 | 1,44567651 | 0,46902031 | 1,33794773 | 4,5767E-07 | 1,33794773 | 0,45098715 | 0,65199883 |
| VCA0357 | 1,20236039 | 1,89837274 | 1,13290336 | 1,8524E-06 | 1,13290336 | 0,45043687 | 0,65239548 |
| VC1905 | 1,1120672 | 9,12301397 | 1,08871043 | 8,9023E-06 | 1,08871043 | 0,45032688 | 0,65247478 |
| VCA0888 | 1,20599293 | 1,16212586 | 1,12302167 | 1,134E-06 | 1,12302167 | 0,44947231 | 0,65309099 |
| VCA0453 | 1,19279469 | 1,15810114 | 1,1314655 | 1,1301E-06 | 1,1314655 | 0,44931289 | 0,65320598 |
| VC1738 | 1,0860125 | 14,6531917 | 1,07397319 | 1,4299E-05 | 1,07397319 | 0,44905934 | 0,65338887 |
| VC1558 | -1,2078007 | -17,65307 | -1,2388349 | -1,723E-05 | -1,2388349 | -0,4490061 | 0,65342727 |
| VC0632 | 1,005515 | -1,8967141 | -1,0734374 | -1,851E-06 | -1,0734374 | -0,4489317 | 0,65348094 |
| VCA0681 | 1,21056139 | 1,1410708 | 1,12506825 | 1,1135E-06 | 1,12506825 | 0,44885501 | 0,65353627 |
| recX | -1,0513138 | -0,9384481 | -1,1540727 | -9,157E-07 | -1,1540727 | -0,4487452 | 0,65361549 |
| VCA0114 | 1,39430523 | 0,64008784 | 1,23271167 | 6,246E-07 | 1,23271167 | 0,44737719 | 0,65460276 |
| VC0964 | 1,09490235 | -35,585156 | -1,0713926 | -3,472E-05 | -1,0713926 | -0,445922 | 0,65565363 |

| | | | | | | | |
|---|---|---|---|---|---|---|---|
| VCA0965 | 1,18325184 | 1,52987558 | 1,09050716 | 1,4929E-06 | 1,09050716 | 0,44576717 | 0,65576546 |
| VC0852 | 1,16464137 | 4,92319653 | 1,10266012 | 4,8041E-06 | 1,10266012 | 0,44551898 | 0,65594477 |
| VCA0874 | -1,0772742 | -11,007375 | -1,1002177 | -1,074E-05 | -1,1002177 | -0,4452032 | 0,65617289 |
| VC1757 | 1,14515134 | 2,22323342 | 1,11223059 | 2,1695E-06 | 1,11223059 | 0,44511951 | 0,65623341 |
| VC1576 | 1,37863826 | 93,4237426 | 1,18264694 | 9,1164E-05 | 1,18264694 | 0,44474025 | 0,6565075 |
| VC2567 | 1,01002955 | -1,9141331 | -1,0713438 | -1,868E-06 | -1,0713438 | -0,4447384 | 0,65650885 |
| VC1527 | -1,0518454 | -3,3463716 | -1,138094 | -3,265E-06 | -1,138094 | -0,4444962 | 0,65668392 |
| VC2289 | 1,07555444 | 8,46209054 | 1,08876074 | 8,2574E-06 | 1,08876074 | 0,44409818 | 0,65697163 |
| VC2405 | 1,08226249 | 2,84019586 | 1,04737531 | 2,7715E-06 | 1,04737531 | 0,44404304 | 0,65701147 |
| VCA1106 | -1,1136667 | -6,1840643 | -1,199318 | -6,034E-06 | -1,199318 | -0,4435574 | 0,65736262 |
| VCA0009 | -1,1312869 | -0,9031222 | -1,1739337 | -8,813E-07 | -1,1739337 | -0,4427756 | 0,65792809 |
| VCA1018 | 1,36716918 | 0,71091806 | 1,20222676 | 6,9372E-07 | 1,20222676 | 0,44254713 | 0,65809335 |
| 5Sg | 1,01155115 | 1019,63477 | 1,21234534 | 0,00099497 | 1,21234534 | 0,44211433 | 0,65840651 |
| VC0081 | 1,18718847 | 1,26874467 | 1,10823654 | 1,2381E-06 | 1,10823654 | 0,44205637 | 0,65844843 |
| nagA | 1,21118078 | 11,5853751 | 1,14688426 | 1,1305E-05 | 1,14688426 | 0,44187933 | 0,65857656 |
| fliG | 1,04762745 | 3,66106783 | 1,03595147 | 3,5725E-06 | 1,03595147 | 0,44041396 | 0,65963733 |
| VCA0133 | -1,0460687 | -1,231233 | -1,115344 | -1,201E-06 | -1,115344 | -0,4402343 | 0,65976745 |
| VCA0472 | -1,0070913 | -1,9690599 | -1,076005 | -1,921E-06 | -1,076005 | -0,4401641 | 0,65981828 |
| VCA0423 | 1,06862002 | 5,34943638 | 1,05665673 | 5,22E-06 | 1,05665673 | 0,43967438 | 0,66017297 |
| VC1369 | 1,2895419 | 0,85827661 | 1,16230946 | 8,3751E-07 | 1,16230946 | 0,43963117 | 0,66020428 |
| VC1660 | -1,0221677 | -0,9538769 | -1,1439862 | -9,308E-07 | -1,1439862 | -0,438387 | 0,66110576 |
| VCA0065 | 1,41399378 | 0,5167857 | 1,2824618 | 5,0428E-07 | 1,2824618 | 0,43802049 | 0,66137145 |
| VCA0694 | -1,0851485 | -12,113695 | -1,1032024 | -1,182E-05 | -1,1032024 | -0,4369035 | 0,66218134 |
| tRNA-Lys-2 | 1,14431239 | 2,04028065 | 1,1265173 | 1,9909E-06 | 1,1265173 | 0,43672512 | 0,66231073 |
| fadA | -1,0767244 | -1,3694239 | -1,1901007 | -1,336E-06 | -1,1901007 | -0,4366674 | 0,66235257 |
| VC1597 | 1,2850413 | 0,86523732 | 1,15832837 | 8,4431E-07 | 1,15832837 | 0,43636521 | 0,66257179 |
| VCA0331 | -1,006292 | -1,6616822 | -1,0818264 | -1,621E-06 | -1,0818264 | -0,4359535 | 0,66287049 |
| VC0911 | -1,256557 | -75,633788 | -1,4290635 | -7,38E-05 | -1,4290635 | -0,4350765 | 0,66350691 |
| VCA0812 | 1,33871076 | 0,56925911 | 1,24882211 | 5,5549E-07 | 1,24882211 | 0,43469346 | 0,66378496 |
| VC1980 | 1,19063227 | 1,18168615 | 1,11235895 | 1,1531E-06 | 1,11235895 | 0,43424422 | 0,66411112 |
| VCA0989 | 1,3153943 | 0,79832404 | 1,17087384 | 7,7901E-07 | 1,17087384 | 0,43418199 | 0,66415631 |
| VCA0004 | -1,1253063 | -18,186583 | -1,1097243 | -1,775E-05 | -1,1097243 | -0,4341126 | 0,66420668 |
| VC0001 | -1,1011621 | -1,0057648 | -1,1397411 | -9,814E-07 | -1,1397411 | -0,4339725 | 0,66430845 |
| VC1333 | -1,0630621 | -0,5777138 | -1,2436448 | -5,637E-07 | -1,2436448 | -0,4338296 | 0,66441224 |
| mnmC | 1,21092442 | 1,02666564 | 1,12999477 | 1,0018E-06 | 1,12999477 | 0,43356139 | 0,66460699 |
| VC2006 | -1,1084575 | -17,686196 | -1,1130861 | -1,726E-05 | -1,1130861 | -0,4333417 | 0,66476659 |
| tRNA-Met-8 | -1,4267969 | -0,5257375 | -1,4416513 | -5,13E-07 | -1,4416513 | -0,4327948 | 0,66516384 |
| VC1812 | -1,0536999 | -0,8904746 | -1,1549734 | -8,689E-07 | -1,1549734 | -0,4321947 | 0,66559991 |
| VC2384 | 1,16391519 | 1,55030224 | 1,10716591 | 1,5128E-06 | 1,10716591 | 0,43180539 | 0,66588287 |
| VC2770 | 1,15072129 | 45,033808 | 1,1062278 | 4,3944E-05 | 1,1062278 | 0,43151013 | 0,66609751 |
| VC1377 | -1,0516022 | -8,2915118 | -1,1074324 | -8,091E-06 | -1,1074324 | -0,4313913 | 0,66618392 |
| deoA | -1,1057183 | -6,5564149 | -1,2248265 | -6,398E-06 | -1,2248265 | -0,4313342 | 0,66622539 |
| VC0778 | 1,54378819 | 0,35998339 | 1,41541362 | 3,5127E-07 | 1,41541362 | 0,43096966 | 0,66649045 |
| VC1096 | 1,22561521 | 1,01143442 | 1,12997041 | 9,8697E-07 | 1,12997041 | 0,4302954 | 0,66698079 |
| VCA0395 | -1,0427538 | -6,897041 | -1,0992459 | -6,73E-06 | -1,0992459 | -0,4302201 | 0,66703553 |

| | | | | | | | |
|---|---|---|---|---|---|---|---|
| VC0492 | 1,23014643 | 55,8033399 | 1,18100607 | 5,4453E-05 | 1,18100607 | 0,42975434 | 0,66737437 |
| VC0506 | -1,011453 | -2,2009669 | -1,0574016 | -2,148E-06 | -1,0574016 | -0,4292182 | 0,66776443 |
| VCA0737 | 1,13334594 | 2,19150666 | 1,06277433 | 2,1385E-06 | 1,06277433 | 0,42843312 | 0,66833583 |
| VCA1112 | 1,19944803 | 1,47520371 | 1,1539081 | 1,4395E-06 | 1,1539081 | 0,42772821 | 0,66884903 |
| tRNA-Ala-1 | 1,27119184 | 1,58226766 | 1,18778153 | 1,544E-06 | 1,18778153 | 0,42752775 | 0,668995 |
| VC1685 | -1,0066097 | -1,8065502 | -1,0697517 | -1,763E-06 | -1,0697517 | -0,4273755 | 0,66910587 |
| VC1361 | -1,0571317 | -0,7196788 | -1,1847439 | -7,023E-07 | -1,1847439 | -0,427283 | 0,6691732 |
| VC0267 | -1,0125015 | -1,7557535 | -1,1277069 | -1,713E-06 | -1,1277069 | -0,4255998 | 0,67039951 |
| VC2652 | 1,00476579 | -0,8522067 | -1,1524518 | -8,316E-07 | -1,1524518 | -0,4255336 | 0,67044773 |
| cpxP | 1,2057601 | 0,96181856 | 1,13372124 | 9,3855E-07 | 1,13372124 | 0,42524596 | 0,67065739 |
| VC0950 | 1,13716801 | 3,26484251 | 1,09330768 | 3,1859E-06 | 1,09330768 | 0,4250671 | 0,67078778 |
| VCA0425 | 1,15608691 | 4,75971372 | 1,11263176 | 4,6446E-06 | 1,11263176 | 0,42506126 | 0,67079203 |
| rpsR | 1,31582874 | 65,9747795 | 1,09973891 | 6,4379E-05 | 1,09973891 | 0,42503219 | 0,67081323 |
| VCA0553 | 1,17586244 | 1,12557784 | 1,11280373 | 1,0983E-06 | 1,11280373 | 0,42460273 | 0,67112632 |
| VC0476 | -1,2080123 | -15,949058 | -1,1504528 | -1,556E-05 | -1,1504528 | -0,4239128 | 0,67162942 |
| VC0894 | 1,21497608 | 1,94796232 | 1,17583508 | 1,9008E-06 | 1,17583508 | 0,42299622 | 0,67229804 |
| VCA0702 | 1,33981267 | 10,5163387 | 1,28192296 | 1,0262E-05 | 1,28192296 | 0,42236162 | 0,67276112 |
| VC0586 | -1,1429939 | -10,069396 | -1,1397026 | -9,826E-06 | -1,1397026 | -0,4217125 | 0,67323493 |
| VC2487 | 1,27305036 | 0,81976975 | 1,15582303 | 7,9994E-07 | 1,15582303 | 0,42161597 | 0,67330536 |
| VCA0937 | 1,21708494 | 0,95232329 | 1,13256014 | 9,2929E-07 | 1,13256014 | 0,42141529 | 0,67345187 |
| VCA0716 | 1,22108964 | 0,90532177 | 1,13983128 | 8,8342E-07 | 1,13983128 | 0,42128499 | 0,673547 |
| VC2716 | 1,1687497 | 1,56197519 | 1,09841647 | 1,5242E-06 | 1,09841647 | 0,4212067 | 0,67360416 |
| VCA0034 | -1,1390597 | -0,5661905 | -1,2329683 | -5,525E-07 | -1,2329683 | -0,4209685 | 0,67377808 |
| VC2478 | 1,10687837 | -71,85564 | -1,0818944 | -7,012E-05 | -1,0818944 | -0,4209442 | 0,67379586 |
| VC0148 | -1,0107001 | -2,5143424 | -1,0608164 | -2,454E-06 | -1,0608164 | -0,4205093 | 0,67411345 |
| VC1281 | 1,37926094 | 1,9546325 | 1,19336191 | 1,9073E-06 | 1,19336191 | 0,42035075 | 0,67422926 |
| VCA0403 | 1,14615574 | 1,6062381 | 1,08314297 | 1,5674E-06 | 1,08314297 | 0,42018623 | 0,67434943 |
| VCA0078 | 1,2783086 | 5,93382156 | 1,21186329 | 5,7903E-06 | 1,21186329 | 0,42007532 | 0,67443046 |
| VCA0385 | -1,0703693 | -8,2758115 | -1,087555 | -8,076E-06 | -1,087555 | -0,4199445 | 0,67452602 |
| VCA0638 | -1,0761895 | -0,6635144 | -1,1941378 | -6,475E-07 | -1,1941378 | -0,4196717 | 0,67472533 |
| VC1955 | 1,30766062 | 0,67723639 | 1,18893674 | 6,6085E-07 | 1,18893674 | 0,41876773 | 0,6753859 |
| VC1868 | -1,0902548 | -0,6367195 | -1,2015753 | -6,213E-07 | -1,2015753 | -0,4182032 | 0,67579859 |
| VCA0700 | -1,1167169 | -0,5855276 | -1,2206173 | -5,714E-07 | -1,2206173 | -0,4177514 | 0,67612893 |
| tRNA-Glu-3 | 1,12593096 | 19,8928192 | 1,18550774 | 1,9412E-05 | 1,18550774 | 0,41693449 | 0,67672635 |
| VC2026 | 1,26505998 | 56,0684716 | 1,07098823 | 5,4712E-05 | 1,07098823 | 0,41669827 | 0,67689913 |
| VC1194 | 1,17441791 | 1,26023477 | 1,10518701 | 1,2297E-06 | 1,10518701 | 0,41597344 | 0,67742945 |
| VCA0723 | -1,043416 | -1,1760622 | -1,103108 | -1,148E-06 | -1,103108 | -0,4159047 | 0,67747978 |
| VCA1058 | 1,36415603 | 0,66838396 | 1,18864025 | 6,5221E-07 | 1,18864025 | 0,41572339 | 0,67761244 |
| VCA0250 | -1,5174405 | -31,801041 | -1,688713 | -3,103E-05 | -1,688713 | -0,4151796 | 0,67801044 |
| VCA0654 | -1,0347924 | -1,8340283 | -1,1364139 | -1,79E-06 | -1,1364139 | -0,4146025 | 0,67843291 |
| hemD | -1,0319107 | -1,8451893 | -1,1146922 | -1,801E-06 | -1,1146922 | -0,4144482 | 0,67854594 |
| VC2428 | 1,10937955 | 3,72944314 | 1,07801521 | 3,6392E-06 | 1,07801521 | 0,41440375 | 0,67857846 |
| tRNA-Ala-4 | 1,28088431 | 1,1967189 | 1,27432152 | 1,1678E-06 | 1,27432152 | 0,41400115 | 0,67887328 |
| VC1170 | -1,029988 | -2,2700528 | -1,0701921 | -2,215E-06 | -1,0701921 | -0,413953 | 0,67890857 |
| VCA0769 | 1,193115 | 23,8664796 | 1,138442 | 2,3289E-05 | 1,138442 | 0,41377414 | 0,67903955 |

| | | | | | | | |
|---|---|---|---|---|---|---|---|
| VC0233 | 1,07205114 | 2,36781744 | 1,05165233 | 2,3105E-06 | 1,05165233 | 0,41330395 | 0,67938395 |
| VC0583 | 1,09734395 | 23,4778861 | 1,08637959 | 2,291E-05 | 1,08637959 | 0,41318261 | 0,67947284 |
| VC1925 | -1,0258598 | -1,189067 | -1,1384851 | -1,16E-06 | -1,1384851 | -0,4124514 | 0,68000858 |
| VC2212 | -1,0184661 | -2,4302711 | -1,0593054 | -2,371E-06 | -1,0593054 | -0,4122117 | 0,68018426 |
| VC1168 | 1,18281159 | 3,21571404 | 1,13028436 | 3,1379E-06 | 1,13028436 | 0,41205771 | 0,68029714 |
| VC2057 | 1,01180866 | -2,2852169 | -1,050744 | -2,23E-06 | -1,050744 | -0,4118802 | 0,68042724 |
| VC2560 | 1,44622165 | 1,96596631 | 1,48276257 | 1,9184E-06 | 1,48276257 | 0,41178013 | 0,68050061 |
| VCA0158 | -1,0301643 | -0,9373186 | -1,1274936 | -9,146E-07 | -1,1274936 | -0,4105026 | 0,68143731 |
| VC0913 | -1,0226984 | -1,158233 | -1,101907 | -1,13E-06 | -1,101907 | -0,4104463 | 0,68147863 |
| VCA0873 | -1,0097146 | -1,8516023 | -1,0624689 | -1,807E-06 | -1,0624689 | -0,4101844 | 0,68167072 |
| VC0802 | 1,17896421 | 1,88948642 | 1,12360348 | 1,8438E-06 | 1,12360348 | 0,40968573 | 0,68203652 |
| VCA0706 | -1,0262784 | -0,8964094 | -1,1330523 | -8,747E-07 | -1,1330523 | -0,4095678 | 0,68212306 |
| VC0299 | 1,30825775 | 0,57727383 | 1,21434587 | 5,6331E-07 | 1,21434587 | 0,40943694 | 0,68221905 |
| flhA | -1,0296665 | -4,162909 | -1,0758965 | -4,062E-06 | -1,0758965 | -0,409242 | 0,68236212 |
| VCA0868 | 1,18845253 | 3,85315333 | 1,12039268 | 3,7599E-06 | 1,12039268 | 0,40904525 | 0,68250647 |
| VC1700 | 1,13300785 | 2,19488613 | 1,08913157 | 2,1418E-06 | 1,08913157 | 0,408774 | 0,68270554 |
| VCA0890 | -1,017314 | -0,8580787 | -1,1384562 | -8,373E-07 | -1,1384562 | -0,4082552 | 0,68308637 |
| hmuV | -1,0314703 | -3,0234104 | -1,1380065 | -2,95E-06 | -1,1380065 | -0,4073282 | 0,683767 |
| VC0438 | -1,1498306 | -6,1864881 | -1,1479632 | -6,037E-06 | -1,1479632 | -0,4071259 | 0,68391554 |
| VC2692 | 1,22177312 | 1,42568994 | 1,10967002 | 1,3912E-06 | 1,10967002 | 0,40619797 | 0,68459716 |
| VCA0024 | 1,2227677 | 0,92340417 | 1,12616655 | 9,0107E-07 | 1,12616655 | 0,40544474 | 0,68515064 |
| VC0875 | 1,10229376 | 5,86922598 | 1,08395036 | 5,7272E-06 | 1,08395036 | 0,40512778 | 0,6853836 |
| VC1223 | 1,24983203 | 8,54909601 | 1,17500779 | 8,3423E-06 | 1,17500779 | 0,40436164 | 0,68594683 |
| VCA0593 | 1,1432135 | 1,47424961 | 1,07645886 | 1,4386E-06 | 1,07645886 | 0,40355491 | 0,68654007 |
| tRNA-Met-9 | -1,0373721 | -2,8069462 | -1,050024 | -2,739E-06 | -1,050024 | -0,4035204 | 0,68656545 |
| rpsF | 1,24118658 | 80,3992112 | 1,08143523 | 7,8454E-05 | 1,08143523 | 0,40213281 | 0,68758631 |
| VCA1005 | 1,22117498 | 0,5864503 | 1,20235063 | 5,7226E-07 | 1,20235063 | 0,40205546 | 0,68764322 |
| VC2299 | 1,15954574 | 6,61250064 | 1,1019104 | 6,4525E-06 | 1,1019104 | 0,40193657 | 0,68773073 |
| upp | 1,09314222 | 5,84586945 | 1,09332713 | 5,7044E-06 | 1,09332713 | 0,4015647 | 0,68800443 |
| VC2542 | 1,15923191 | 1,42916629 | 1,07801768 | 1,3946E-06 | 1,07801768 | 0,40121571 | 0,68826133 |
| VC0131 | -1,0428765 | -2,9213196 | -1,1281515 | -2,851E-06 | -1,1281515 | -0,4003536 | 0,68889611 |
| VCA0139 | -1,009166 | -1,4667092 | -1,0755336 | -1,431E-06 | -1,0755336 | -0,4001678 | 0,68903291 |
| csrC | 1,00606362 | 1012,1188 | 1,06398605 | 0,00098763 | 1,06398605 | 0,39988633 | 0,68924027 |
| VC2464 | -1,0139618 | -3,1987287 | -1,0337891 | -3,121E-06 | -1,0337891 | -0,3993051 | 0,68966844 |
| VC1583 | 1,26373919 | 0,79853063 | 1,14245099 | 7,7921E-07 | 1,14245099 | 0,39910299 | 0,68981734 |
| VC2751 | 1,12220507 | 4,77509898 | 1,07726153 | 4,6596E-06 | 1,07726153 | 0,39869356 | 0,69011903 |
| VC1290 | -1,0234031 | -1,0551787 | -1,1057278 | -1,03E-06 | -1,1057278 | -0,3986754 | 0,69013241 |
| VC1679 | 1,2620135 | 0,89040963 | 1,12651877 | 8,6887E-07 | 1,12651877 | 0,39865756 | 0,69014555 |
| VC1078 | -1,0516779 | -0,9124443 | -1,123002 | -8,904E-07 | -1,123002 | -0,3982414 | 0,69045228 |
| VC2729 | 1,09010121 | 4,30112191 | 1,0732213 | 4,1971E-06 | 1,0732213 | 0,3981442 | 0,69052391 |
| VC2309 | 1,19890462 | 0,9105673 | 1,12312769 | 8,8854E-07 | 1,12312769 | 0,39802296 | 0,69061327 |
| tRNA-Leu-8 | -1,169629 | -39,991096 | -1,1792408 | -3,902E-05 | -1,1792408 | -0,3977443 | 0,69081872 |
| VC1724 | 1,00111004 | -1,0212329 | -1,1085999 | -9,965E-07 | -1,1085999 | -0,3972307 | 0,69119737 |
| VC1832 | 1,07258219 | 2,35428352 | 1,04569962 | 2,2973E-06 | 1,04569962 | 0,39722496 | 0,69120159 |
| VCA1050 | 1,35239705 | 1,89412333 | 1,18076523 | 1,8483E-06 | 1,18076523 | 0,39704898 | 0,69133136 |

| | | | | | | | |
|---|---|---|---|---|---|---|---|
| VCA1001 | -1,0916616 | -0,5572798 | -1,2077647 | -5,438E-07 | -1,2077647 | -0,3966499 | 0,69162566 |
| VC0870 | -1,2436853 | -1,3121016 | -1,2058556 | -1,28E-06 | -1,2058556 | -0,3964096 | 0,69180292 |
| VCA0381 | -1,0018863 | -1,0463093 | -1,1053666 | -1,021E-06 | -1,1053666 | -0,3963516 | 0,69184571 |
| VCA0059 | -1,2659122 | -403,79591 | -1,1303153 | -0,000394 | -1,1303153 | -0,3962231 | 0,69194047 |
| coaD | 1,14845634 | 1,45466778 | 1,07424492 | 1,4195E-06 | 1,07424492 | 0,39523027 | 0,69267297 |
| VC0620 | -1,1738385 | -7,9490119 | -1,2985553 | -7,757E-06 | -1,2985553 | -0,394931 | 0,69289384 |
| VC1016 | 1,13107743 | 1,68027278 | 1,06384135 | 1,6396E-06 | 1,06384135 | 0,39488365 | 0,69292877 |
| VC2730 | 1,08616435 | 8,04333576 | 1,08681384 | 7,8488E-06 | 1,08681384 | 0,39475981 | 0,69302017 |
| VCA0856 | -1,1668144 | -0,3037131 | -1,4097399 | -2,964E-07 | -1,4097399 | -0,3936059 | 0,69387202 |
| VCA0280 | 1,33322151 | 30,2606007 | 1,27109103 | 2,9529E-05 | 1,27109103 | 0,39342619 | 0,69400475 |
| VC0857 | 1,19957605 | 1,71282118 | 1,13504664 | 1,6714E-06 | 1,13504664 | 0,39294502 | 0,6943601 |
| VCA0418 | 1,21746607 | 0,94325055 | 1,13664721 | 9,2043E-07 | 1,13664721 | 0,39283688 | 0,69443998 |
| rplU | -1,0194823 | 274,775076 | 1,17018809 | 0,00026813 | 1,17018809 | 0,39277613 | 0,69448486 |
| VCA0431 | 1,24134012 | 5,9741423 | 1,1734908 | 5,8296E-06 | 1,1734908 | 0,39255319 | 0,69464954 |
| VC2175 | -1,0561203 | -7,5277305 | -1,1161671 | -7,346E-06 | -1,1161671 | -0,3921811 | 0,69492443 |
| VCA0530 | -1,158571 | -0,5329873 | -1,2124723 | -5,201E-07 | -1,2124723 | -0,3918609 | 0,69516104 |
| rho | 1,07420129 | 17,9524348 | 1,06384007 | 1,7518E-05 | 1,06384007 | 0,39148495 | 0,69543882 |
| VCA0031 | 1,1678038 | 0,94904736 | 1,11378185 | 9,2609E-07 | 1,11378185 | 0,39148296 | 0,6954403 |
| VC0151 | 1,38241494 | 6,67248523 | 1,23159558 | 6,5111E-06 | 1,23159558 | 0,39118242 | 0,69566243 |
| mutL | 1,11044692 | 2,0807223 | 1,05021873 | 2,0304E-06 | 1,05021873 | 0,39102945 | 0,69577548 |
| VC0508 | -1,031898 | -1,2773055 | -1,1246139 | -1,246E-06 | -1,1246139 | -0,3908836 | 0,69588327 |
| VC0575 | 1,14634727 | 24,2424485 | 1,14405084 | 2,3656E-05 | 1,14405084 | 0,39044079 | 0,69621065 |
| VCA0639 | -1,0463419 | -0,5959199 | -1,1863475 | -5,815E-07 | -1,1863475 | -0,390353 | 0,69627553 |
| VC1600 | -1,0291686 | -4,243301 | -1,0837569 | -4,141E-06 | -1,0837569 | -0,3899147 | 0,69659966 |
| artP | 1,37849693 | 3,25597866 | 1,29551476 | 3,1772E-06 | 1,29551476 | 0,38989204 | 0,69661641 |
| VC1531 | 1,07431385 | 4,86635138 | 1,07480637 | 4,7486E-06 | 1,07480637 | 0,38981493 | 0,69667343 |
| VCA0691 | 1,51978345 | 0,35311588 | 1,33343478 | 3,4457E-07 | 1,33343478 | 0,38906945 | 0,69722478 |
| VC0809 | 1,21708654 | 1,01494121 | 1,10460658 | 9,9039E-07 | 1,10460658 | 0,3890248 | 0,69725782 |
| VC1405 | 1,16163994 | 1,0702067 | 1,09879989 | 1,0443E-06 | 1,09879989 | 0,38876718 | 0,69744839 |
| VC0205 | 2,09847039 | 0,18885985 | 1,72731932 | 1,8429E-07 | 1,72731932 | 0,38870944 | 0,69749111 |
| VCA0894 | 1,11651814 | 1,83111115 | 1,05656023 | 1,7868E-06 | 1,05656023 | 0,38869585 | 0,69750116 |
| VC2018 | 1,11016473 | 1,86976838 | 1,05533284 | 1,8245E-06 | 1,05533284 | 0,38860851 | 0,69756578 |
| VC1709 | 1,16664812 | 1,97873407 | 1,10387098 | 1,9309E-06 | 1,10387098 | 0,38858578 | 0,6975826 |
| VC1555 | -1,0581134 | -5,2203713 | -1,1442901 | -5,094E-06 | -1,1442901 | -0,3883827 | 0,69773285 |
| VC0398 | 1,01737668 | -1,6857173 | -1,0614742 | -1,645E-06 | -1,0614742 | -0,388344 | 0,69776148 |
| VC1076 | -1,0076023 | -1,0736739 | -1,0980518 | -1,048E-06 | -1,0980518 | -0,3879885 | 0,69802454 |
| rumB | 1,18133488 | 1,01237305 | 1,10364719 | 9,8788E-07 | 1,10364719 | 0,38683469 | 0,69887861 |
| VC0807 | 1,15588429 | 1,35597321 | 1,09060268 | 1,3232E-06 | 1,09060268 | 0,38666368 | 0,69900522 |
| VCA0641 | 1,41638826 | 0,38864652 | 1,29382495 | 3,7924E-07 | 1,29382495 | 0,38645822 | 0,69915736 |
| ftsA | -1,0521551 | -6,8287532 | -1,0449079 | -6,664E-06 | -1,0449079 | -0,3864548 | 0,69915989 |
| VC2163 | -1,0229421 | -1,8103188 | -1,0564193 | -1,767E-06 | -1,0564193 | -0,3860141 | 0,69948624 |
| VC1165 | 1,28387994 | 0,70485874 | 1,15093831 | 6,8781E-07 | 1,15093831 | 0,38521084 | 0,70008122 |
| VC1783 | -1,0702701 | -4,8887115 | -1,2063938 | -4,77E-06 | -1,2063938 | -0,3850556 | 0,70019627 |
| csrB | -1,0291799 | -5010,3194 | -1,0931925 | -0,0048891 | -1,0931925 | -0,3849075 | 0,70030594 |
| VC1596 | -1,1225415 | -31,585191 | -1,1910024 | -3,082E-05 | -1,1910024 | -0,3845985 | 0,70053491 |

| | | | | | | | |
|---|---|---|---|---|---|---|---|
| VC1012 | 1,14304453 | 1,1639193 | 1,08840034 | 1,1358E-06 | 1,08840034 | 0,38445462 | 0,70064154 |
| VC0127 | -1,0139191 | -2,0219042 | -1,0499275 | -1,973E-06 | -1,0499275 | -0,3843708 | 0,70070363 |
| VC2617 | 1,47706318 | 2,40246762 | 1,27181645 | 2,3443E-06 | 1,27181645 | 0,38418274 | 0,70084303 |
| VC0262 | 1,08782473 | 2,64936778 | 1,05554035 | 2,5853E-06 | 1,05554035 | 0,38409109 | 0,70091095 |
| VC1206 | 1,4109616 | 2,44933481 | 1,22892074 | 2,3901E-06 | 1,22892074 | 0,38405354 | 0,70093878 |
| VCA0260 | 1,28104838 | 0,97784968 | 1,18117548 | 9,5419E-07 | 1,18117548 | 0,38335015 | 0,70146018 |
| cls | 1,00190185 | -1,4660358 | -1,0691091 | -1,431E-06 | -1,0691091 | -0,3832778 | 0,70151384 |
| VCA0420 | 1,13755052 | 2,80924013 | 1,07518023 | 2,7413E-06 | 1,07518023 | 0,38297037 | 0,70174175 |
| VC1199 | 1,20244593 | 0,86132482 | 1,13245566 | 8,4049E-07 | 1,13245566 | 0,3826259 | 0,70199718 |
| VCA0880 | -1,2097864 | -0,2634281 | -1,45385 | -2,571E-07 | -1,45385 | -0,3823174 | 0,70222599 |
| tRNA-Met-3 | -1,1589537 | -25,669377 | -1,1581797 | -2,505E-05 | -1,1581797 | -0,3820272 | 0,70244118 |
| VCA0708 | -1,1135095 | -0,5005223 | -1,2151349 | -4,884E-07 | -1,2151349 | -0,3818812 | 0,70254951 |
| VC2023 | 1,12269975 | 6,69557793 | 1,10433673 | 6,5336E-06 | 1,10433673 | 0,38174351 | 0,70265164 |
| VCA0103 | 1,13913151 | 1,37636181 | 1,07316464 | 1,3431E-06 | 1,07316464 | 0,38173746 | 0,70265612 |
| VCA0758 | 1,43411703 | 3,05336296 | 1,35513967 | 2,9795E-06 | 1,35513967 | 0,38165213 | 0,70271943 |
| VCA1063 | 1,56292313 | 4,04579459 | 1,53601994 | 3,9479E-06 | 1,53601994 | 0,38158043 | 0,70277263 |
| VC1736 | 1,26103832 | 0,71090352 | 1,14634367 | 6,9371E-07 | 1,14634367 | 0,38133302 | 0,70295617 |
| VCA0358 | -1,0089047 | -2,1919918 | -1,1098044 | -2,139E-06 | -1,1098044 | -0,3809223 | 0,70326091 |
| VC0308 | -1,0224493 | -1,5446082 | -1,0787741 | -1,507E-06 | -1,0787741 | -0,3806363 | 0,70347312 |
| VCA0179 | 1,35727424 | 1,16314451 | 1,31038032 | 1,135E-06 | 1,31038032 | 0,38031732 | 0,7037099 |
| VC1416 | 1,27869547 | 0,52146325 | 1,20371002 | 5,0885E-07 | 1,20371002 | 0,38027875 | 0,70373852 |
| VCA0267 | 1,26958289 | 0,5080987 | 1,20835636 | 4,9581E-07 | 1,20835636 | 0,37923119 | 0,70451621 |
| VC1613 | -1,0323695 | -4,0905706 | -1,1443518 | -3,992E-06 | -1,1443518 | -0,3789351 | 0,70473605 |
| VCA0884 | 1,13389309 | 1,27769404 | 1,0775572 | 1,2468E-06 | 1,0775572 | 0,37827901 | 0,70522335 |
| VCA0549 | 1,1456155 | 9,38484335 | 1,11418661 | 9,1578E-06 | 1,11418661 | 0,37818447 | 0,70529358 |
| VC0972 | -1,0951784 | -45,566242 | -1,1986198 | -4,446E-05 | -1,1986198 | -0,3766707 | 0,70641834 |
| lipB | 1,11138858 | 2,19739343 | 1,08358236 | 2,1442E-06 | 1,08358236 | 0,37654205 | 0,70651398 |
| VC2711 | 1,16879615 | 1,38234776 | 1,07073373 | 1,3489E-06 | 1,07073373 | 0,37637843 | 0,70663559 |
| VC1212 | 1,2159754 | 0,82114358 | 1,12176602 | 8,0128E-07 | 1,12176602 | 0,37599841 | 0,70691809 |
| VC1390 | 1,41006243 | 0,44376531 | 1,23760454 | 4,3303E-07 | 1,23760454 | 0,37598754 | 0,70692617 |
| VCA0160 | 1,2916912 | 0,99463041 | 1,16773434 | 9,7057E-07 | 1,16773434 | 0,37594015 | 0,70696141 |
| VC1967 | -1,0199657 | -1,0750823 | -1,0914661 | -1,049E-06 | -1,0914661 | -0,3755682 | 0,70723798 |
| VC1681 | -1,042165 | -1,0453942 | -1,0969567 | -1,02E-06 | -1,0969567 | -0,3754653 | 0,70731449 |
| VC1430 | 1,05547933 | 2,57897458 | 1,0502686 | 2,5166E-06 | 1,0502686 | 0,37519491 | 0,70751553 |
| 23Sc | -1,042861 | -0,8428757 | -1,1178038 | -8,225E-07 | -1,1178038 | -0,3750428 | 0,70762863 |
| queF | 1,12948446 | 1,37734104 | 1,07029331 | 1,344E-06 | 1,07029331 | 0,37456459 | 0,70798432 |
| VC1994 | 1,14117124 | 1,68529331 | 1,06751113 | 1,6445E-06 | 1,06751113 | 0,37429141 | 0,70818753 |
| VCA0557 | -1,0041633 | -1,4040998 | -1,0736756 | -1,37E-06 | -1,0736756 | -0,3730664 | 0,70909901 |
| potE | 1,45636425 | 2,45937608 | 1,45137226 | 2,3999E-06 | 1,45137226 | 0,3728389 | 0,70926837 |
| VCA0285 | 1,12320517 | 1,43496865 | 1,06671694 | 1,4003E-06 | 1,06671694 | 0,37278986 | 0,70930486 |
| tRNA-Met-2 | -1,2166501 | -16,900796 | -1,1610901 | -1,649E-05 | -1,1610901 | -0,3719924 | 0,70989855 |
| VC2660 | 1,10973246 | 25,1372396 | 1,11387672 | 2,4529E-05 | 1,11387672 | 0,37134066 | 0,71038383 |
| VCA0709 | -1,0366166 | -0,8773456 | -1,1105281 | -8,561E-07 | -1,1105281 | -0,3712689 | 0,71043726 |
| deoD | -1,0871409 | -10,922126 | -1,0625943 | -1,066E-05 | -1,0625943 | -0,3698316 | 0,71150797 |
| VCA0620 | 1,1834416 | 2,61238523 | 1,09309004 | 2,5492E-06 | 1,09309004 | 0,36977714 | 0,71154856 |

| | | | | | | | |
|---|---|---|---|---|---|---|---|
| VC1981 | -1,0004135 | -1,1224803 | -1,0843521 | -1,095E-06 | -1,0843521 | -0,3691606 | 0,71200806 |
| tRNA-Tyr-1 | -1,2181037 | -16,794377 | -1,1535596 | -1,639E-05 | -1,1535596 | -0,3683652 | 0,71260099 |
| VC1519 | -1,0079626 | -3,0989848 | -1,1082386 | -3,024E-06 | -1,1082386 | -0,3681407 | 0,71276834 |
| VC0542 | 1,16502628 | 1,29070556 | 1,07239786 | 1,2595E-06 | 1,07239786 | 0,36779382 | 0,71302699 |
| tRNA-Leu-9 | -1,2239382 | -11,495984 | -1,205957 | -1,122E-05 | -1,205957 | -0,3667722 | 0,71378895 |
| VCA1084 | -1,0079505 | -1,4155672 | -1,0651168 | -1,381E-06 | -1,0651168 | -0,3659358 | 0,71441298 |
| VC1582 | 1,3502087 | 0,40141652 | 1,24934976 | 3,9171E-07 | 1,24934976 | 0,36537146 | 0,71483415 |
| VC2109 | -1,0551013 | -10,054605 | -1,0795186 | -9,811E-06 | -1,0795186 | -0,3652583 | 0,71491859 |
| VC0304 | -1,0442753 | -2,8323409 | -1,0545055 | -2,764E-06 | -1,0545055 | -0,3652339 | 0,71493681 |
| VC2383 | 1,17474957 | 1,01199305 | 1,09167574 | 9,8751E-07 | 1,09167574 | 0,36478071 | 0,71527512 |
| VC0534 | 1,20858791 | 2,94389505 | 1,10812861 | 2,8727E-06 | 1,10812861 | 0,36471356 | 0,71532526 |
| VC0587 | 1,1842757 | 0,8389925 | 1,11155343 | 8,187E-07 | 1,11155343 | 0,36465467 | 0,71536921 |
| VC0829 | -1,1642579 | -3,0161417 | -1,1703382 | -2,943E-06 | -1,1703382 | -0,3644286 | 0,71553799 |
| VC1023 | 1,154175 | 1,53124047 | 1,059433 | 1,4942E-06 | 1,059433 | 0,36410611 | 0,71577879 |
| tRNA-Val-4 | 1,13470698 | 7,62557352 | 1,167456 | 7,4411E-06 | 1,167456 | 0,36286873 | 0,71670296 |
| VC0396 | -1,0647114 | -0,5375785 | -1,177806 | -5,246E-07 | -1,177806 | -0,3628657 | 0,71670524 |
| VC2011 | 1,17551769 | 1,81179423 | 1,08609714 | 1,768E-06 | 1,08609714 | 0,36243035 | 0,71703048 |
| VC0825 | 1,16651341 | 1,63571967 | 1,11435549 | 1,5961E-06 | 1,11435549 | 0,36223666 | 0,7171752 |
| VC1896 | -1,0403963 | -3,1633266 | -1,047487 | -3,087E-06 | -1,047487 | -0,3618598 | 0,71745678 |
| VC0648 | -1,0117708 | -4,4499669 | -1,0742606 | -4,342E-06 | -1,0742606 | -0,3616867 | 0,71758615 |
| VC0965 | 1,32142005 | 71,6875851 | 1,23362498 | 6,9953E-05 | 1,23362498 | 0,36133712 | 0,71784746 |
| VC2462 | 1,08691785 | 3,37559213 | 1,06541879 | 3,2939E-06 | 1,06541879 | 0,36111498 | 0,71801351 |
| hemG | 1,16649616 | 1,49716014 | 1,05977771 | 1,4609E-06 | 1,05977771 | 0,36104363 | 0,71806684 |
| VC1542 | 1,26482201 | 0,50338446 | 1,1888406 | 4,9121E-07 | 1,1888406 | 0,36095393 | 0,7181339 |
| purN | -1,0886786 | -3,2398176 | -1,1086592 | -3,161E-06 | -1,1086592 | -0,3605738 | 0,71841807 |
| VC0452 | 1,28207807 | 0,66565153 | 1,13920148 | 6,4955E-07 | 1,13920148 | 0,36048008 | 0,71848816 |
| VC1656 | 1,20614759 | 0,81200227 | 1,11269825 | 7,9236E-07 | 1,11269825 | 0,36047962 | 0,71848851 |
| VCA1095 | -1,1174392 | -0,4917287 | -1,1931564 | -4,798E-07 | -1,1931564 | -0,3604489 | 0,71851146 |
| VC1645 | 1,36405241 | 15,6320924 | 1,28475418 | 1,5254E-05 | 1,28475418 | 0,36032107 | 0,71860706 |
| VCA0063 | 1,35776671 | 0,41653134 | 1,23168607 | 4,0645E-07 | 1,23168607 | 0,36017893 | 0,71871334 |
| vrrA | 1,28466237 | 48,840604 | 1,13222636 | 4,7659E-05 | 1,13222636 | 0,36002158 | 0,71883101 |
| VC1725 | 1,00667327 | -1,2927323 | -1,0691304 | -1,261E-06 | -1,0691304 | -0,3599645 | 0,71887372 |
| hemA | -1,0124341 | -3,7214577 | -1,0389997 | -3,631E-06 | -1,0389997 | -0,359351 | 0,71933253 |
| VCA0510 | 1,17327732 | 1,51166105 | 1,09451508 | 1,4751E-06 | 1,09451508 | 0,35882538 | 0,71972574 |
| VC2388 | 1,13197691 | 2,43063231 | 1,11924829 | 2,3718E-06 | 1,11924829 | 0,3585088 | 0,7199626 |
| VCA0567 | 1,02969706 | 4,91112817 | 1,05862751 | 4,7923E-06 | 1,05862751 | 0,35716519 | 0,72096816 |
| VC2435 | -1,2385166 | -34,880064 | -1,180279 | -3,404E-05 | -1,180279 | -0,3562233 | 0,72167339 |
| VCA0734 | 1,03826092 | -1,4213462 | -1,0612429 | -1,387E-06 | -1,0612429 | -0,3559418 | 0,72188416 |
| VC1042 | -1,0208053 | -0,8279192 | -1,1073807 | -8,079E-07 | -1,1073807 | -0,3557525 | 0,72202594 |
| VC0761 | 1,1029966 | 18,2409406 | 1,07210042 | 1,78E-05 | 1,07210042 | 0,35560156 | 0,72213899 |
| VC1682 | -1,0544515 | -1,2830494 | -1,1044998 | -1,252E-06 | -1,1044998 | -0,3555791 | 0,72215584 |
| VC0897 | 1,14351144 | 1,16119044 | 1,07509911 | 1,1331E-06 | 1,07509911 | 0,35506981 | 0,72253731 |
| VC2178 | -1,0378584 | -5,8730654 | -1,0798042 | -5,731E-06 | -1,0798042 | -0,3548964 | 0,72266724 |
| VCA0667 | 1,00019855 | -0,6512837 | -1,1377407 | -6,355E-07 | -1,1377407 | -0,3548138 | 0,72272911 |
| ispF | 1,13019137 | 1,36985711 | 1,06320098 | 1,3367E-06 | 1,06320098 | 0,3548086 | 0,722733 |

| | | | | | | | |
|---|---|---|---|---|---|---|---|
| VC2756 | 1,0162384 | -1,2435959 | -1,0697454 | -1,214E-06 | -1,0697454 | -0,3545712 | 0,72291087 |
| VC2637 | 1,27012798 | 0,58918225 | 1,15307459 | 5,7493E-07 | 1,15307459 | 0,35449357 | 0,72296904 |
| VC2743 | 1,22832932 | 0,62604079 | 1,17346907 | 6,109E-07 | 1,17346907 | 0,35440785 | 0,72303326 |
| VC1084 | -1,0415708 | -2,7266314 | -1,1245561 | -2,661E-06 | -1,1245561 | -0,3543498 | 0,72307675 |
| VC2262 | 1,16944306 | 0,94354594 | 1,09279177 | 9,2072E-07 | 1,09279177 | 0,35427143 | 0,72313549 |
| VC1935 | 1,39720621 | 0,60769347 | 1,1563563 | 5,9299E-07 | 1,1563563 | 0,35308612 | 0,72402389 |
| VC1410 | 1,19358086 | 1,01258362 | 1,11256635 | 9,8809E-07 | 1,11256635 | 0,3528399 | 0,72420848 |
| VC2534 | 1,01863252 | -1,5694906 | -1,0542029 | -1,532E-06 | -1,0542029 | -0,352481 | 0,72447755 |
| mnmA | 1,1312659 | 8,52380111 | 1,14611713 | 8,3176E-06 | 1,14611713 | 0,35190419 | 0,72491014 |
| VC2422 | 1,03063893 | -1,5379323 | -1,0642335 | -1,501E-06 | -1,0642335 | -0,3516451 | 0,72510444 |
| VCA0317 | 1,11945904 | 1,58968565 | 1,07681928 | 1,5512E-06 | 1,07681928 | 0,35142471 | 0,72526976 |
| VCA0039 | 1,18057691 | 3,4702373 | 1,08653858 | 3,3863E-06 | 1,08653858 | 0,35136881 | 0,72531169 |
| VC1059 | 1,22189757 | 10,2186775 | 1,12956166 | 9,9715E-06 | 1,12956166 | 0,35127538 | 0,72538179 |
| VC1804 | -1,0553849 | -0,6821098 | -1,1277074 | -6,656E-07 | -1,1277074 | -0,3504621 | 0,72599194 |
| VC0899 | -1,1152061 | -12,909571 | -1,1871409 | -1,26E-05 | -1,1871409 | -0,3502184 | 0,72617484 |
| VC1937 | 1,31390881 | 0,59008299 | 1,14888371 | 5,7581E-07 | 1,14888371 | 0,35021538 | 0,72617708 |
| VC1760 | 1,06302052 | 5,46274467 | 1,08210176 | 5,3306E-06 | 1,08210176 | 0,34987271 | 0,72643424 |
| VCA0026 | -1,0390347 | -7,3147221 | -1,0672986 | -7,138E-06 | -1,0672986 | -0,3494441 | 0,72675597 |
| VC1795 | 1,18138576 | 0,78205533 | 1,10978763 | 7,6314E-07 | 1,10978763 | 0,34941239 | 0,72677974 |
| VC2233 | 1,09189726 | 1,96549555 | 1,04227579 | 1,9179E-06 | 1,04227579 | 0,34937789 | 0,72680564 |
| VC1180 | 1,01842065 | -1,6478429 | -1,0542814 | -1,608E-06 | -1,0542814 | -0,3492615 | 0,72689302 |
| VC1344 | 1,26910479 | 0,48792228 | 1,18169899 | 4,7612E-07 | 1,18169899 | 0,34915271 | 0,72697467 |
| VC1116 | 1,17662206 | 1,54067436 | 1,12373067 | 1,5034E-06 | 1,12373067 | 0,34899945 | 0,72708973 |
| VCA0382a | -1,0359114 | -2,8188916 | -1,1317167 | -2,751E-06 | -1,1317167 | -0,3482378 | 0,72766163 |
| adk | -1,0094349 | -55,333467 | -1,1797627 | -5,399E-05 | -1,1797627 | -0,3481792 | 0,72770565 |
| VC0220 | -1,0139274 | -1,2595211 | -1,0838592 | -1,229E-06 | -1,0838592 | -0,347906 | 0,72791081 |
| VCA0033 | 1,25336687 | 0,61033731 | 1,14152044 | 5,9557E-07 | 1,14152044 | 0,34785243 | 0,72795102 |
| VCA1033 | -1,0288738 | -0,6335939 | -1,1358093 | -6,183E-07 | -1,1358093 | -0,3476568 | 0,72809795 |
| VC0143 | 1,32704079 | 0,56847473 | 1,16284353 | 5,5472E-07 | 1,16284353 | 0,34731936 | 0,72835142 |
| VCA0008 | -1,0776274 | -0,7170825 | -1,1283791 | -6,997E-07 | -1,1283791 | -0,3471858 | 0,72845174 |
| VC2156 | -1,0159202 | -5,5447964 | -1,0538627 | -5,411E-06 | -1,0538627 | -0,3469537 | 0,72862615 |
| VC2437 | -1,0519068 | -5,3068798 | -1,0938178 | -5,178E-06 | -1,0938178 | -0,3467987 | 0,72874258 |
| VC2466 | 1,05644533 | -23,148388 | -1,0549143 | -2,259E-05 | -1,0549143 | -0,3467644 | 0,72876834 |
| VCA0090 | 1,01112868 | -0,8027424 | -1,1050477 | -7,833E-07 | -1,1050477 | -0,3466672 | 0,72884134 |
| VC1118 | 1,00132525 | -1,7493481 | -1,1142566 | -1,707E-06 | -1,1142566 | -0,3466324 | 0,7288675 |
| ispG | 1,08061367 | 2,85199746 | 1,03753142 | 2,783E-06 | 1,03753142 | 0,3464198 | 0,72902726 |
| VCA0419 | 1,02522128 | -1,4908408 | -1,065368 | -1,455E-06 | -1,065368 | -0,3457938 | 0,7294977 |
| VC0033 | 1,23346029 | 1,0096006 | 1,15846799 | 9,8518E-07 | 1,15846799 | 0,34578508 | 0,72950425 |
| VC1350 | 1,09987532 | 6,51073649 | 1,11065638 | 6,3532E-06 | 1,11065638 | 0,34560803 | 0,72963733 |
| VC0403 | 1,04749739 | 2,37873696 | 1,03624805 | 2,3212E-06 | 1,03624805 | 0,34528414 | 0,72988078 |
| VCA0740 | 1,11187869 | 1,54208091 | 1,05282129 | 1,5048E-06 | 1,05282129 | 0,34502416 | 0,73007622 |
| VC0550 | -1,0937199 | -13,36561 | -1,1546665 | -1,304E-05 | -1,1546665 | -0,3444675 | 0,73049477 |
| VC0612 | -1,0350704 | -0,8441032 | -1,115063 | -8,237E-07 | -1,115063 | -0,344299 | 0,73062146 |
| 23Sd | -1,0350452 | -0,7676316 | -1,1085275 | -7,491E-07 | -1,1085275 | -0,3442857 | 0,73063148 |
| VC1253 | 1,02354378 | -1,7943216 | -1,0542019 | -1,751E-06 | -1,0542019 | -0,343378 | 0,73131412 |

| | | | | | | | |
|---|---|---|---|---|---|---|---|
| VC0788 | 1,24833585 | 1,47768786 | 1,15179008 | 1,4419E-06 | 1,15179008 | 0,34337684 | 0,73131499 |
| VC2337 | -1,0823388 | -11,392977 | -1,1465712 | -1,112E-05 | -1,1465712 | -0,3431584 | 0,73147934 |
| VC0044 | 1,11769431 | 1,52515872 | 1,05278749 | 1,4883E-06 | 1,05278749 | 0,34301883 | 0,73158431 |
| VC0672 | 1,00067078 | -2,3718216 | -1,0367666 | -2,314E-06 | -1,0367666 | -0,3425215 | 0,73195844 |
| VCA1004 | -1,0682973 | -5,0903291 | -1,1741195 | -4,967E-06 | -1,1741195 | -0,3424647 | 0,73200124 |
| purH | 1,20681847 | 1,42929164 | 1,16390642 | 1,3947E-06 | 1,16390642 | 0,34224736 | 0,73216476 |
| VCA0249 | 1,23360857 | 5,06349925 | 1,18475718 | 4,941E-06 | 1,18475718 | 0,34181077 | 0,73249332 |
| VC1723 | -1,0278251 | -0,9675548 | -1,0837274 | -9,441E-07 | -1,0837274 | -0,3415185 | 0,7327133 |
| VCA0683 | -1,0848014 | -1,7255621 | -1,1657177 | -1,684E-06 | -1,1657177 | -0,3412778 | 0,73289448 |
| VC0859 | 1,20887815 | 0,52059968 | 1,15934811 | 5,0801E-07 | 1,15934811 | 0,3394888 | 0,73424154 |
| VCA0975 | 1,13066464 | 7,44079436 | 1,09857167 | 7,2608E-06 | 1,09857167 | 0,33913585 | 0,73450741 |
| VCA0949 | -1,012644 | -0,8773929 | -1,0913713 | -8,562E-07 | -1,0913713 | -0,3391168 | 0,73452178 |
| VCA1081 | 1,11992322 | 1,50966089 | 1,05945533 | 1,4731E-06 | 1,05945533 | 0,33818347 | 0,73522494 |
| VC1731 | -1,1607809 | -6,5932103 | -1,1447708 | -6,434E-06 | -1,1447708 | -0,3381206 | 0,7352723 |
| VC1525 | -1,0471097 | -3,2737282 | -1,1396841 | -3,195E-06 | -1,1396841 | -0,3379153 | 0,73542706 |
| VCA0618 | 1,16283414 | 0,93226297 | 1,08505709 | 9,0971E-07 | 1,08505709 | 0,33777582 | 0,73553215 |
| VC1587 | 1,32470695 | 0,41689671 | 1,20033521 | 4,0681E-07 | 1,20033521 | 0,33745011 | 0,73577763 |
| VC2113 | 1,25063915 | 0,7341323 | 1,13969531 | 7,1637E-07 | 1,13969531 | 0,3373137 | 0,73588045 |
| 5Sc | 1,01150901 | 742,907109 | 1,16035841 | 0,00072494 | 1,16035841 | 0,3372799 | 0,73590593 |
| VC0047 | 1,25718199 | 0,66023019 | 1,12178004 | 6,4426E-07 | 1,12178004 | 0,33716883 | 0,73598965 |
| VC0633 | -1,0289859 | 248,66608 | 1,05634374 | 0,00024265 | 1,05634374 | 0,33684961 | 0,73623029 |
| VC2475 | 1,08655455 | 1,73557694 | 1,04453786 | 1,6936E-06 | 1,04453786 | 0,33678916 | 0,73627585 |
| VC0344 | 1,10173491 | 1,85467488 | 1,04146366 | 1,8098E-06 | 1,04146366 | 0,33617637 | 0,73673788 |
| VC1246 | 1,11098232 | 3,66332543 | 1,09895701 | 3,5747E-06 | 1,09895701 | 0,33579463 | 0,73702576 |
| VC1302 | 1,19654216 | 3,9977387 | 1,1172252 | 3,901E-06 | 1,1172252 | 0,33531727 | 0,73738578 |
| VC1129 | 1,13035594 | 2,78501842 | 1,12828281 | 2,7176E-06 | 1,12828281 | 0,33515198 | 0,73751046 |
| VC0140 | 1,03059233 | -1,5910576 | -1,0780398 | -1,553E-06 | -1,0780398 | -0,3351492 | 0,73751254 |
| VC1743 | 1,00884574 | -3,0437891 | -1,0763675 | -2,97E-06 | -1,0763675 | -0,3349035 | 0,73769789 |
| VC2332 | 1,01068788 | -2,3466951 | -1,0715616 | -2,29E-06 | -1,0715616 | -0,3336844 | 0,73861777 |
| VC1797 | -1,0421905 | -0,7980883 | -1,0971556 | -7,788E-07 | -1,0971556 | -0,3330478 | 0,73909822 |
| VCA0603 | 1,25305436 | 0,6273613 | 1,12523767 | 6,1218E-07 | 1,12523767 | 0,33303076 | 0,73911107 |
| VC0287 | -1,0912835 | -1,9205655 | -1,259809 | -1,874E-06 | -1,259809 | -0,3324306 | 0,73956414 |
| VC0641 | -1,0860124 | -12,765556 | -1,066728 | -1,246E-05 | -1,066728 | -0,3324043 | 0,73958402 |
| VC0064 | 1,25651966 | 0,64864966 | 1,12020317 | 6,3296E-07 | 1,12020317 | 0,33215146 | 0,7397749 |
| VC1473 | -1,0482183 | -0,8292026 | -1,0927894 | -8,091E-07 | -1,0927894 | -0,332108 | 0,73980772 |
| VCA0164 | 1,00892311 | -1,2578058 | -1,060149 | -1,227E-06 | -1,060149 | -0,3319228 | 0,73994758 |
| VC2704 | -1,0853914 | -0,3159176 | -1,2624708 | -3,083E-07 | -1,2624708 | -0,3315864 | 0,74020156 |
| VC0593 | 1,095932 | 1,99121578 | 1,06318353 | 1,943E-06 | 1,06318353 | 0,3310273 | 0,74062388 |
| VCA0833 | 1,0026618 | -0,9290845 | -1,0816768 | -9,066E-07 | -1,0816768 | -0,3306994 | 0,74087156 |
| VC0177 | 1,18875 | 1,35643298 | 1,10855699 | 1,3236E-06 | 1,10855699 | 0,33013609 | 0,74129715 |
| rnpA | -1,0928629 | -23,727385 | -1,0996071 | -2,315E-05 | -1,0996071 | -0,3300976 | 0,74132624 |
| VC1335 | 1,36466943 | 0,5606246 | 1,15048236 | 5,4706E-07 | 1,15048236 | 0,32988207 | 0,74148908 |
| VCA0789 | -1,0423336 | -0,689966 | -1,1106531 | -6,733E-07 | -1,1106531 | -0,3294194 | 0,7418387 |
| VCA1017 | 1,15892684 | 1,17143383 | 1,10391809 | 1,1431E-06 | 1,10391809 | 0,32894663 | 0,74219604 |
| VCA0198 | -1,0290826 | -9,9373912 | -1,0404109 | -9,697E-06 | -1,0404109 | -0,3278237 | 0,74304498 |

| | | | | | | | |
|---|---|---|---|---|---|---|---|
| VC1710 | 1,23672159 | 1,66472962 | 1,14206128 | 1,6245E-06 | 1,14206128 | 0,32780282 | 0,74306077 |
| VC1693 | 1,23880813 | 0,51084116 | 1,15055819 | 4,9848E-07 | 1,15055819 | 0,32755251 | 0,74325004 |
| VC0601 | 1,15801083 | 0,78592855 | 1,09510141 | 7,6692E-07 | 1,09510141 | 0,32714852 | 0,74355557 |
| VC1295 | 1,26376117 | 0,55198524 | 1,13774084 | 5,3863E-07 | 1,13774084 | 0,32664732 | 0,74393466 |
| VC0687 | -1,0982748 | -41,747908 | -1,3606916 | -4,074E-05 | -1,3606916 | -0,3266311 | 0,74394696 |
| VC1755 | 1,09517261 | 2,12566867 | 1,06048712 | 2,0742E-06 | 1,06048712 | 0,32631787 | 0,74418388 |
| VC0338 | -1,0529937 | -4,6761948 | -1,1508767 | -4,563E-06 | -1,1508767 | -0,3261565 | 0,74430598 |
| VC2719 | -1,0794662 | -0,4313917 | -1,1789668 | -4,21E-07 | -1,1789668 | -0,3260303 | 0,74440144 |
| vibF | 1,31238159 | 0,40971709 | 1,18865665 | 3,9981E-07 | 1,18865665 | 0,32549957 | 0,74480302 |
| pbpG | 1,15717498 | 0,9683307 | 1,07557812 | 9,4491E-07 | 1,07557812 | 0,32523988 | 0,74499954 |
| VC0717 | 1,12642528 | 2,43564065 | 1,08950017 | 2,3767E-06 | 1,08950017 | 0,32506697 | 0,7451304 |
| VC1013 | 1,12294048 | 1,26455165 | 1,0602579 | 1,234E-06 | 1,0602579 | 0,32454147 | 0,74552815 |
| VCA0559 | 1,01839238 | -1,4892885 | -1,0480803 | -1,453E-06 | -1,0480803 | -0,323867 | 0,74603875 |
| VCA0836 | 1,07595717 | 1,74556106 | 1,04087425 | 1,7033E-06 | 1,04087425 | 0,32385764 | 0,74604583 |
| VC1680 | -1,0126195 | -1,1361654 | -1,0673157 | -1,109E-06 | -1,0673157 | -0,3237367 | 0,7461374 |
| VC2254 | -1,0123264 | -2,5788335 | -1,0629994 | -2,516E-06 | -1,0629994 | -0,3235885 | 0,74624961 |
| VC0560 | -1,0243075 | -2,755545 | -1,0256308 | -2,689E-06 | -1,0256308 | -0,3234362 | 0,7463649 |
| VCA0647 | -1,0906546 | -0,4089076 | -1,1862629 | -3,99E-07 | -1,1862629 | -0,3232851 | 0,74647933 |
| VC1650 | 1,25761952 | 0,4953307 | 1,15106424 | 4,8335E-07 | 1,15106424 | 0,32304509 | 0,7466611 |
| VC2287 | 1,21521939 | 0,65675124 | 1,11162955 | 6,4086E-07 | 1,11162955 | 0,32273282 | 0,74689761 |
| VCA0927 | 1,18198193 | 0,58904294 | 1,12524076 | 5,7479E-07 | 1,12524076 | 0,3227037 | 0,74691966 |
| thiG | 1,29217187 | 0,49600047 | 1,15027959 | 4,84E-07 | 1,15027959 | 0,32248162 | 0,74708787 |
| VC0622 | -1,0069052 | -3,4891522 | -1,1310813 | -3,405E-06 | -1,1310813 | -0,3224201 | 0,74713448 |
| VCA0405 | 1,13111416 | 8,89661529 | 1,10252833 | 8,6814E-06 | 1,10252833 | 0,32193585 | 0,74750131 |
| VC1553 | 1,23269931 | 0,49520129 | 1,14947 | 4,8322E-07 | 1,14947 | 0,32141311 | 0,74789736 |
| VC1851 | 1,02163415 | -1,405345 | -1,0501955 | -1,371E-06 | -1,0501955 | -0,3212867 | 0,74799311 |
| VCA0002 | 1,09083462 | 3,53516366 | 1,04962349 | 3,4496E-06 | 1,04962349 | 0,32113063 | 0,74811141 |
| VCA0818 | 1,22607652 | 0,80792395 | 1,10003204 | 7,8838E-07 | 1,10003204 | 0,32106895 | 0,74815815 |
| truA | 1,13591085 | 1,0308024 | 1,06894044 | 1,0059E-06 | 1,06894044 | 0,32100711 | 0,74820502 |
| VCA0811 | 1,28211424 | 0,50941579 | 1,14452832 | 4,9709E-07 | 1,14452832 | 0,32092845 | 0,74826463 |
| VCA0369 | 1,1235791 | 0,98262944 | 1,07233314 | 9,5886E-07 | 1,07233314 | 0,32077269 | 0,74838267 |
| VC2238 | 1,14119298 | 0,97233095 | 1,07283843 | 9,4881E-07 | 1,07283843 | 0,32016081 | 0,74884644 |
| VC2551 | 1,19305728 | 1,49195973 | 1,08866973 | 1,4559E-06 | 1,08866973 | 0,32004899 | 0,7489312 |
| VC1422 | 1,23007369 | 4,47701578 | 1,23511488 | 4,3687E-06 | 1,23511488 | 0,31968909 | 0,74920405 |
| VC2626 | 1,08829303 | 2,75197387 | 1,06396022 | 2,6854E-06 | 1,06396022 | 0,31816267 | 0,75036156 |
| VCA0177 | -1,0369663 | -5,2794288 | -1,0459474 | -5,152E-06 | -1,0459474 | -0,3179053 | 0,75055677 |
| VC1311 | -1,0288761 | -0,6512601 | -1,1088896 | -6,355E-07 | -1,1088896 | -0,3176183 | 0,75077453 |
| VC2373 | 1,28985826 | 5,05667514 | 1,33541854 | 4,9343E-06 | 1,33541854 | 0,31679212 | 0,75140136 |
| VCA0325 | 1,15301315 | 1,07514688 | 1,11568186 | 1,0491E-06 | 1,11568186 | 0,31646555 | 0,75164919 |
| VCA0481 | 1,13758021 | 1,88164567 | 1,07479705 | 1,8361E-06 | 1,07479705 | 0,31621855 | 0,75183664 |
| VC1528 | 1,071272 | 2,06383726 | 1,04084579 | 2,0139E-06 | 1,04084579 | 0,31614262 | 0,75189427 |
| smpB | 1,05862597 | 3,27270749 | 1,05071762 | 3,1935E-06 | 1,05071762 | 0,31496272 | 0,75278997 |
| VC1459 | -1,1606245 | -0,2961925 | -1,2513003 | -2,89E-07 | -1,2513003 | -0,3149399 | 0,75280728 |
| VC1811 | -1,0099696 | -1,5313651 | -1,0440651 | -1,494E-06 | -1,0440651 | -0,3147078 | 0,75298352 |
| trmE | -1,0057321 | -1,2199763 | -1,0554573 | -1,19E-06 | -1,0554573 | -0,3142435 | 0,75333615 |

| | | | | | | | |
|---|---|---|---|---|---|---|---|
| VC1605 | 1,31130528 | 1,75697028 | 1,14649654 | 1,7145E-06 | 1,14649654 | 0,31403347 | 0,75349563 |
| VCA0330 | 1,14875979 | 1,08951587 | 1,0622082 | 1,0632E-06 | 1,0622082 | 0,31400663 | 0,75351601 |
| VC1394 | -1,0172465 | -1,0641423 | -1,0637235 | -1,038E-06 | -1,0637235 | -0,31397 | 0,75354381 |
| VC0706 | 1,07665498 | 1062,35326 | 1,22165954 | 0,00103665 | 1,22165954 | 0,31391569 | 0,75358509 |
| VC1776 | -1,177441 | -17,034693 | -1,2698811 | -1,662E-05 | -1,2698811 | -0,313208 | 0,75412263 |
| VC0305 | 1,04250634 | 2,55680954 | 1,02589301 | 2,495E-06 | 1,02589301 | 0,31312234 | 0,75418773 |
| VC0471 | 1,28538882 | 0,56136369 | 1,12303297 | 5,4778E-07 | 1,12303297 | 0,31240372 | 0,75473373 |
| VC2353 | 1,01084555 | -1,0407776 | -1,0644571 | -1,016E-06 | -1,0644571 | -0,3122306 | 0,75486528 |
| VC2176 | -1,0437592 | -6,6374737 | -1,0739607 | -6,477E-06 | -1,0739607 | -0,3121873 | 0,7548982 |
| VCA0523 | -1,0941609 | -4,4169994 | -1,1297249 | -4,31E-06 | -1,1297249 | -0,3117818 | 0,75520638 |
| VC1887 | 1,08743075 | 3,90536224 | 1,0565011 | 3,8109E-06 | 1,0565011 | 0,31153573 | 0,7553934 |
| VCA1085 | 1,001122 | -4,0076576 | -1,1213149 | -3,911E-06 | -1,1213149 | -0,3114395 | 0,75546655 |
| tRNA-Glu-1 | 1,32641813 | 0,23237512 | 1,32289156 | 2,2675E-07 | 1,32289156 | 0,31129311 | 0,75557782 |
| VC1807 | 1,36208059 | 0,39622606 | 1,17618941 | 3,8664E-07 | 1,17618941 | 0,31022316 | 0,75639127 |
| VCA0631 | 1,01150161 | -0,771008 | -1,0867269 | -7,524E-07 | -1,0867269 | -0,3100532 | 0,7565205 |
| VCA1109 | 1,3716651 | 0,32208374 | 1,22088005 | 3,1429E-07 | 1,22088005 | 0,30999955 | 0,75656131 |
| VC2634 | 1,18197818 | 0,66166551 | 1,10168521 | 6,4566E-07 | 1,10168521 | 0,30990344 | 0,7566344 |
| VCA0537 | 1,22310623 | 3,23431775 | 1,10398122 | 3,1561E-06 | 1,10398122 | 0,30982386 | 0,75669492 |
| VC0052 | 1,12792201 | 1,43590248 | 1,10487669 | 1,4012E-06 | 1,10487669 | 0,3098061 | 0,75670843 |
| VCA0182 | 1,2341383 | 0,64056005 | 1,10486767 | 6,2506E-07 | 1,10486767 | 0,30942138 | 0,75700102 |
| helD | 1,00831776 | -1,1730831 | -1,055868 | -1,145E-06 | -1,055868 | -0,3092528 | 0,75712927 |
| VC1960 | 1,15089877 | 1,3930712 | 1,05548108 | 1,3594E-06 | 1,05548108 | 0,30909762 | 0,75724728 |
| VC2399 | 1,03485574 | 7,60053065 | 1,03910267 | 7,4167E-06 | 1,03910267 | 0,30901868 | 0,75730733 |
| VCA0777 | 1,00186477 | -0,8670923 | -1,0761432 | -8,461E-07 | -1,0761432 | -0,3088749 | 0,75741667 |
| VC1990 | 1,12117979 | 0,9165536 | 1,07165933 | 8,9438E-07 | 1,07165933 | 0,3084037 | 0,75777518 |
| VC0757 | -1,0235912 | -1,4239122 | -1,0720819 | -1,389E-06 | -1,0720819 | -0,3082895 | 0,75786204 |
| VC1196 | 1,09791367 | 3,04323578 | 1,06980369 | 2,9696E-06 | 1,06980369 | 0,30815058 | 0,75796777 |
| VC0777 | 1,42156544 | 0,25361448 | 1,28423531 | 2,4748E-07 | 1,28423531 | 0,30769218 | 0,75831658 |
| VC2063 | 1,01394722 | 5,82142494 | 1,033128 | 5,6806E-06 | 1,033128 | 0,30701566 | 0,75883146 |
| VC1674 | 1,0297692 | -0,5108115 | -1,1310034 | -4,985E-07 | -1,1310034 | -0,3069312 | 0,75889572 |
| VC0689 | 1,14571325 | 1,00950693 | 1,06400693 | 9,8509E-07 | 1,06400693 | 0,30646202 | 0,7592529 |
| metF | 1,01610302 | -1,4967685 | -1,0873344 | -1,461E-06 | -1,0873344 | -0,3055141 | 0,75997463 |
| VC1439 | -1,0515867 | -7,0084356 | -1,0599091 | -6,839E-06 | -1,0599091 | -0,3048042 | 0,76051528 |
| VC1606 | 1,20532887 | 1,50280033 | 1,12653212 | 1,4664E-06 | 1,12653212 | 0,3032744 | 0,76168077 |
| VCA0485 | 1,15017037 | 2,86918212 | 1,06909949 | 2,7998E-06 | 1,06909949 | 0,30298592 | 0,7619006 |
| VC2676 | -1,0000885 | -3,6476927 | -1,0614165 | -3,559E-06 | -1,0614165 | -0,3029313 | 0,76194225 |
| VC1780 | 1,3788869 | 6,34834778 | 1,19142598 | 6,1948E-06 | 1,19142598 | 0,30292168 | 0,76194956 |
| VC1876 | 1,11668116 | 5,54289704 | 1,08306535 | 5,4088E-06 | 1,08306535 | 0,3028398 | 0,76201196 |
| VCA0354 | -1,0254611 | -1,3836423 | -1,0898016 | -1,35E-06 | -1,0898016 | -0,3026141 | 0,76218395 |
| VC1466 | 1,12018008 | 1,14440536 | 1,10256385 | 1,1167E-06 | 1,10256385 | 0,30256086 | 0,76222456 |
| VCA0104 | 1,14710367 | 1,53311595 | 1,0614244 | 1,496E-06 | 1,0614244 | 0,30244686 | 0,76231145 |
| VC2432 | -1,1576216 | -10,056226 | -1,1147757 | -9,813E-06 | -1,1147757 | -0,3019149 | 0,76271694 |
| VC0473 | -1,0348275 | -4,6421861 | -1,1069418 | -4,53E-06 | -1,1069418 | -0,3018216 | 0,7627881 |
| thyA | 1,02272208 | 3,79853613 | 1,02956811 | 3,7066E-06 | 1,02956811 | 0,30145975 | 0,76306395 |
| VC1469 | -1,103109 | -0,2763485 | -1,2461903 | -2,697E-07 | -1,2461903 | -0,3014405 | 0,76307859 |

| | | | | | | | |
|---|---|---|---|---|---|---|---|
| engB | -1,0416467 | -3,0851803 | -1,031474 | -3,011E-06 | -1,031474 | -0,300732 | 0,7636189 |
| VC0454 | 1,14548326 | 0,90493979 | 1,0747665 | 8,8305E-07 | 1,0747665 | 0,30034127 | 0,76391687 |
| VC2075 | -1,0846922 | -3,6094154 | -1,1081528 | -3,522E-06 | -1,1081528 | -0,300269 | 0,76397201 |
| VC2358 | 1,12063085 | 6,20610805 | 1,08947206 | 6,056E-06 | 1,08947206 | 0,30008095 | 0,76411543 |
| VC0230 | -1,0168368 | -1,9926803 | -1,0402053 | -1,944E-06 | -1,0402053 | -0,2995923 | 0,76448815 |
| VCA0061 | -1,1400242 | -6,9828316 | -1,098617 | -6,814E-06 | -1,098617 | -0,2995701 | 0,76450509 |
| VC2604 | -1,0143951 | -5,4010962 | -1,0224453 | -5,27E-06 | -1,0224453 | -0,299478 | 0,76457536 |
| VCA0871 | 1,12772367 | 1,07175814 | 1,05732914 | 1,0458E-06 | 1,05732914 | 0,2993291 | 0,76468897 |
| rrmA | 1,12132562 | 1,05996633 | 1,05798511 | 1,0343E-06 | 1,05798511 | 0,29932831 | 0,76468957 |
| VC1722 | 1,00823852 | -23,059358 | -1,068557 | -2,25E-05 | -1,068557 | -0,2987127 | 0,76515924 |
| VCA0889 | 1,19511941 | 0,69939814 | 1,08868677 | 6,8248E-07 | 1,08868677 | 0,29848146 | 0,76533573 |
| mraW | 1,16476363 | 1,11619721 | 1,05830805 | 1,0892E-06 | 1,05830805 | 0,298463 | 0,76534982 |
| VC0701 | 1,08860271 | 1,32458894 | 1,04826478 | 1,2925E-06 | 1,04826478 | 0,29820768 | 0,76554467 |
| VC1772 | 1,17571197 | 1,60150529 | 1,10424785 | 1,5628E-06 | 1,10424785 | 0,29812221 | 0,7656099 |
| VCA0962 | 1,17397764 | 0,8990111 | 1,06814479 | 8,7726E-07 | 1,06814479 | 0,29810681 | 0,76562166 |
| VC1471 | 1,11727764 | 1,041197 | 1,07646861 | 1,016E-06 | 1,07646861 | 0,29798141 | 0,76571736 |
| VC1391 | 1,13727333 | 0,59877148 | 1,10639851 | 5,8429E-07 | 1,10639851 | 0,29791659 | 0,76576683 |
| VCA1077 | -1,0021343 | -2,4087372 | -1,0565854 | -2,35E-06 | -1,0565854 | -0,2974162 | 0,76614882 |
| VC2621 | 1,00979226 | -1,4717725 | -1,1095792 | -1,436E-06 | -1,1095792 | -0,2963431 | 0,76696812 |
| VCA0208 | 1,01580299 | -1,0318559 | -1,0582962 | -1,007E-06 | -1,0582962 | -0,2961011 | 0,76715286 |
| VC1028 | 1,10286411 | 1,3479502 | 1,0666913 | 1,3153E-06 | 1,0666913 | 0,29605477 | 0,76718827 |
| VC1675 | 1,02452305 | -0,5927823 | -1,1034997 | -5,784E-07 | -1,1034997 | -0,2958066 | 0,76737777 |
| VC2710 | 1,08080157 | 2,46991123 | 1,05193811 | 2,4102E-06 | 1,05193811 | 0,29529265 | 0,76777034 |
| VC0096 | 1,15204694 | 0,43232345 | 1,14399738 | 4,2186E-07 | 1,14399738 | 0,29514191 | 0,76788548 |
| VC0069 | 1,1213823 | 0,88430197 | 1,06788902 | 8,6291E-07 | 1,06788902 | 0,29512087 | 0,76790156 |
| pyrH | 1,08546616 | 3,69049455 | 1,07345523 | 3,6012E-06 | 1,07345523 | 0,29418035 | 0,76862011 |
| VC1395 | 1,11354104 | 0,69298482 | 1,08642218 | 6,7622E-07 | 1,08642218 | 0,29345112 | 0,76917737 |
| VCA0773 | -1,0357928 | -4,9968851 | -1,1039317 | -4,876E-06 | -1,1039317 | -0,2930841 | 0,76945788 |
| VC2720 | 1,24234697 | 5,48143485 | 1,11160612 | 5,3488E-06 | 1,11160612 | 0,29238863 | 0,76998952 |
| secD (NC_002506 631616..633495) | 1,19352659 | 2,19004653 | 1,12947287 | 2,1371E-06 | 1,12947287 | 0,29228596 | 0,770068 |
| VCA0416 | 1,15015405 | 3,60007352 | 1,08705235 | 3,513E-06 | 1,08705235 | 0,29215585 | 0,77016748 |
| VCA0178 | 1,14450468 | 1,1243227 | 1,05764374 | 1,0971E-06 | 1,05764374 | 0,29207233 | 0,77023133 |
| VCA0070 | 1,22769229 | 0,5131167 | 1,1172793 | 5,007E-07 | 1,1172793 | 0,29200567 | 0,77028229 |
| VCA0832 | 1,02888141 | 4,38753059 | 1,0366141 | 4,2814E-06 | 1,0366141 | 0,29192484 | 0,77034411 |
| VC0817 | 1,14431078 | 1,00038347 | 1,05844381 | 9,7618E-07 | 1,05844381 | 0,29190892 | 0,77035627 |
| VC1380 | 1,02571849 | -1,0660287 | -1,0547438 | -1,04E-06 | -1,0547438 | -0,2919026 | 0,77036107 |
| VCA0731 | 1,31319833 | 0,27423584 | 1,23079559 | 2,676E-07 | 1,23079559 | 0,29174732 | 0,77047983 |
| VC1366 | 1,09090449 | 1,65216956 | 1,05317614 | 1,6122E-06 | 1,05317614 | 0,2917341 | 0,77048995 |
| VCA0212 | 1,10706735 | 0,81076869 | 1,07365683 | 7,9116E-07 | 1,07365683 | 0,29169776 | 0,77051773 |
| VC1858 | 1,0211532 | -1,1945147 | -1,0486055 | -1,166E-06 | -1,0486055 | -0,2915918 | 0,77059878 |
| VC2070 | 1,1657236 | 10,3367495 | 1,12962638 | 1,0087E-05 | 1,12962638 | 0,29158464 | 0,77060424 |
| VCA0976 | 1,30415079 | 0,36537946 | 1,16787476 | 3,5654E-07 | 1,16787476 | 0,29134583 | 0,77078685 |
| VC1607 | 1,23597918 | 1,12941318 | 1,16300982 | 1,1021E-06 | 1,16300982 | 0,29107002 | 0,77099778 |
| murC | 1,07164346 | 1,63124589 | 1,03520213 | 1,5918E-06 | 1,03520213 | 0,29094507 | 0,77109334 |
| VC1636 | 1,18596741 | 0,61255646 | 1,09638129 | 5,9774E-07 | 1,09638129 | 0,2906674 | 0,77130571 |

| | | | | | | | |
|---|---|---|---|---|---|---|---|
| VCA1006 | 1,00306002 | -0,4805991 | -1,1243905 | -4,69E-07 | -1,1243905 | -0,2905557 | 0,77139114 |
| VC1356 | 1,15411369 | 0,80008165 | 1,07263552 | 7,8073E-07 | 1,07263552 | 0,29003022 | 0,77179313 |
| VC1944 | 1,22087944 | 0,58595223 | 1,10048604 | 5,7178E-07 | 1,10048604 | 0,28999199 | 0,77182237 |
| VC1541 | 1,20713234 | 0,49208447 | 1,12049814 | 4,8018E-07 | 1,12049814 | 0,28963603 | 0,77209471 |
| VC1632 | 1,24977115 | 0,53350618 | 1,1105368 | 5,206E-07 | 1,1105368 | 0,28952657 | 0,77217846 |
| VC1031 | 1,21492301 | 0,52536976 | 1,11232682 | 5,1266E-07 | 1,11232682 | 0,28950454 | 0,77219532 |
| VCA0712 | 1,00553932 | -0,7210616 | -1,0803751 | -7,036E-07 | -1,0803751 | -0,2890938 | 0,77250959 |
| VC0892 | 1,03170124 | -1,6548528 | -1,0506772 | -1,615E-06 | -1,0506772 | -0,2889083 | 0,77265157 |
| nrdR | 1,13458965 | 1,81678231 | 1,04739332 | 1,7728E-06 | 1,04739332 | 0,28869068 | 0,7728181 |
| VC0433 | 1,14983259 | 0,8428206 | 1,0788773 | 8,2243E-07 | 1,0788773 | 0,28857134 | 0,77290944 |
| VC1556 | -1,0081994 | -2,7025592 | -1,1138701 | -2,637E-06 | -1,1138701 | -0,2884622 | 0,77299294 |
| VCA0159 | 1,26171875 | 0,44184563 | 1,13345121 | 4,3116E-07 | 1,13345121 | 0,28794966 | 0,77338529 |
| VC1407 | 1,10675112 | 1,70431773 | 1,04928919 | 1,6631E-06 | 1,04928919 | 0,28755303 | 0,77368892 |
| phhB | -1,0147868 | -2,4293708 | -1,0864599 | -2,371E-06 | -1,0864599 | -0,2872499 | 0,77392099 |
| VC1277 | 1,03124527 | -1,1133347 | -1,0773186 | -1,086E-06 | -1,0773186 | -0,2868913 | 0,77419558 |
| VCA0219 | -1,1514772 | -162,93983 | -1,1532034 | -0,000159 | -1,1532034 | -0,2868292 | 0,77424314 |
| xni | 1,01252231 | -1,1843202 | -1,0473618 | -1,156E-06 | -1,0473618 | -0,2866932 | 0,77434723 |
| VC0724 | -1,0093286 | -0,8884753 | -1,0634965 | -8,67E-07 | -1,0634965 | -0,286391 | 0,77457866 |
| VC1571 | -1,0697501 | -0,6424025 | -1,1538329 | -6,269E-07 | -1,1538329 | -0,2856525 | 0,77514427 |
| VC0342 | 1,11192233 | 2,22362428 | 1,08935892 | 2,1698E-06 | 1,08935892 | 0,28560596 | 0,77517995 |
| VCA0987 | 1,25288907 | 9,2335568 | 1,29101414 | 9,0102E-06 | 1,29101414 | 0,28554516 | 0,77522653 |
| VCA0239 | 1,11486491 | 0,81442219 | 1,06897432 | 7,9472E-07 | 1,06897432 | 0,28540013 | 0,77533761 |
| VC1816 | 1,176457 | 0,59448687 | 1,09566885 | 5,8011E-07 | 1,09566885 | 0,28533635 | 0,77538648 |
| VC1211 | 1,15202131 | 0,71966584 | 1,07765409 | 7,0226E-07 | 1,07765409 | 0,28406887 | 0,77635761 |
| VCA0028 | -1,0458632 | -0,3064889 | -1,1920509 | -2,991E-07 | -1,1920509 | -0,2838254 | 0,77654421 |
| VC2772 | -1,0470004 | -6,9586749 | -1,0429531 | -6,79E-06 | -1,0429531 | -0,2835959 | 0,77672013 |
| VCA0579 | 1,12791682 | 1,13957338 | 1,04811508 | 1,112E-06 | 1,04811508 | 0,28340033 | 0,77686998 |
| VCA0165 | -1,0137045 | -1,5221225 | -1,1019241 | -1,485E-06 | -1,1019241 | -0,2830905 | 0,77710751 |
| VC1342 | -1,015198 | -0,8375221 | -1,0763742 | -8,173E-07 | -1,0763742 | -0,2826108 | 0,77747518 |
| VC2081 | 1,16934248 | 1,25595552 | 1,10425132 | 1,2256E-06 | 1,10425132 | 0,28232591 | 0,77769363 |
| VC0725 | -1,0037923 | -1,0450916 | -1,0521545 | -1,02E-06 | -1,0521545 | -0,2822819 | 0,77772737 |
| VCA0355 | -1,0095449 | -0,8496381 | -1,0644726 | -8,291E-07 | -1,0644726 | -0,2821392 | 0,77783678 |
| tRNA-Gly-8 | -1,1173887 | -5,255273 | -1,1181518 | -5,128E-06 | -1,1181518 | -0,2818277 | 0,77807563 |
| VCA0253 | 1,14829425 | 1,27827804 | 1,1037473 | 1,2474E-06 | 1,1037473 | 0,28175788 | 0,77812918 |
| VC0940 | 1,11237622 | 2,41868715 | 1,08768313 | 2,3602E-06 | 1,08768313 | 0,28108449 | 0,77864561 |
| VCA0999 | 1,10123665 | 0,91960937 | 1,05894821 | 8,9736E-07 | 1,05894821 | 0,28104661 | 0,77867466 |
| VC0673 | 1,0137283 | -1,5238357 | -1,0350561 | -1,487E-06 | -1,0350561 | -0,280629 | 0,77899498 |
| VCA0412 | 1,28421565 | 0,50220316 | 1,11134299 | 4,9005E-07 | 1,11134299 | 0,27971601 | 0,77969541 |
| VC1441 | 1,10079082 | 16,9621582 | 1,08515386 | 1,6552E-05 | 1,08515386 | 0,27971215 | 0,77969837 |
| VCA0191 | 1,12370686 | 1,25303565 | 1,04232938 | 1,2227E-06 | 1,04232938 | 0,27913018 | 0,78014493 |
| fliM | -1,003309 | -2,6257088 | -1,0411981 | -2,562E-06 | -1,0411981 | -0,2787633 | 0,7804265 |
| tRNA-Tyr-4 | -1,0961427 | -0,9506586 | -1,1797913 | -9,277E-07 | -1,1797913 | -0,2785478 | 0,78059187 |
| VCA0511 | 1,21556433 | 17,991614 | 1,18447124 | 1,7556E-05 | 1,18447124 | 0,27830725 | 0,78077652 |
| VCA0203 | 1,02031994 | 13,0637911 | 1,05021572 | 1,2748E-05 | 1,05021572 | 0,27818494 | 0,78087041 |
| VC0474 | 1,25857497 | 0,35545258 | 1,15581448 | 3,4685E-07 | 1,15581448 | 0,27761965 | 0,78130436 |

| | | | | | | | |
|---|---|---|---|---|---|---|---|
| VCA0073 | 1,16160208 | 0,62600831 | 1,08545302 | 6,1086E-07 | 1,08545302 | 0,27740597 | 0,78146841 |
| VC2024 | 1,06426934 | 3,8230044 | 1,06603803 | 3,7305E-06 | 1,06603803 | 0,2765206 | 0,78214826 |
| VC0758 | 1,06833527 | 1,65721608 | 1,03115456 | 1,6171E-06 | 1,03115456 | 0,27615377 | 0,78242997 |
| VCA0630 | 1,27083648 | 0,51892029 | 1,10292133 | 5,0637E-07 | 1,10292133 | 0,27602807 | 0,78252651 |
| VCA0554 | -1,0472223 | -1,0114318 | -1,0825906 | -9,87E-07 | -1,0825906 | -0,2759633 | 0,78257628 |
| VC2479 | -1,0510758 | -4,1064946 | -1,0739718 | -4,007E-06 | -1,0739718 | -0,275421 | 0,7829928 |
| VC1718 | 1,02890535 | -2,3671826 | -1,0633643 | -2,31E-06 | -1,0633643 | -0,2753474 | 0,78304937 |
| VCA0971 | 1,00577219 | -3,6248869 | -1,0642054 | -3,537E-06 | -1,0642054 | -0,2752897 | 0,78309372 |
| fliA | 1,00799786 | 4,45121647 | 1,02560841 | 4,3435E-06 | 1,02560841 | 0,27528887 | 0,78309433 |
| VC0500 | 1,14803082 | 0,75740577 | 1,06892823 | 7,3908E-07 | 1,06892823 | 0,27513967 | 0,78320895 |
| VC1591 | 1,18628185 | 1,92213313 | 1,14380372 | 1,8756E-06 | 1,14380372 | 0,27471021 | 0,7835389 |
| VCA0386 | 1,00794691 | -1,1352637 | -1,0494766 | -1,108E-06 | -1,0494766 | -0,2745837 | 0,78363614 |
| ansA | 1,21314347 | 3,74806552 | 1,1008437 | 3,6574E-06 | 1,1008437 | 0,27408366 | 0,78402035 |
| VC1965 | 1,0331311 | 2,23766451 | 1,03759284 | 2,1835E-06 | 1,03759284 | 0,27403849 | 0,78405505 |
| VCA0282 | 1,01790111 | -1,2304377 | -1,0461504 | -1,201E-06 | -1,0461504 | -0,2733258 | 0,78460281 |
| VC2378 | 1,30563909 | 2,18400421 | 1,1326553 | 2,1312E-06 | 1,1326553 | 0,27298533 | 0,78486451 |
| VC1409 | -1,0136106 | -0,8286424 | -1,1001965 | -8,086E-07 | -1,1001965 | -0,2727308 | 0,78506013 |
| tRNA-Leu-6 | -1,0491856 | -2,3579081 | -1,0910276 | -2,301E-06 | -1,0910276 | -0,2724244 | 0,78529576 |
| VC0174 | 1,04582628 | -1,3702192 | -1,0492264 | -1,337E-06 | -1,0492264 | -0,2723888 | 0,78532313 |
| tRNA-Ile-1 | 1,12988457 | 28,7879745 | 1,12918647 | 2,8092E-05 | 1,12918647 | 0,27215869 | 0,78550002 |
| trkA | 1,02324809 | -1,2775849 | -1,0392749 | -1,247E-06 | -1,0392749 | -0,2716955 | 0,7858562 |
| VC0378 | 1,03210653 | -8,5652651 | -1,0247287 | -8,358E-06 | -1,0247287 | -0,2716295 | 0,78590695 |
| VC1605a | 1,20636021 | 17,4508406 | 1,17529522 | 1,7029E-05 | 1,17529522 | 0,27146753 | 0,78603148 |
| VC0973 | -1,0627984 | -17,546347 | -1,1219745 | -1,712E-05 | -1,1219745 | -0,2713599 | 0,78611425 |
| VCA0313 | 1,09138917 | 1,60935076 | 1,04637261 | 1,5704E-06 | 1,04637261 | 0,27126909 | 0,78618409 |
| VC2733 | 1,1313786 | 1,19112717 | 1,04192926 | 1,1623E-06 | 1,04192926 | 0,27088446 | 0,7864799 |
| tRNA-Trp-1 | 1,1405099 | 12,1874487 | 1,19354317 | 1,1893E-05 | 1,19354317 | 0,27079253 | 0,78655062 |
| VCA0346 | 1,11642889 | 3,20215794 | 1,07102223 | 3,1247E-06 | 1,07102223 | 0,2707771 | 0,78656249 |
| VC2015 | 1,12216863 | 0,915743 | 1,05477665 | 8,9359E-07 | 1,05477665 | 0,27062413 | 0,78668015 |
| ruvC | 1,12621133 | 0,71723567 | 1,07039531 | 6,9988E-07 | 1,07039531 | 0,2704824 | 0,78678916 |
| VCA1008 | 1,21899582 | 0,37644348 | 1,13754494 | 3,6734E-07 | 1,13754494 | 0,26957264 | 0,78748906 |
| VC1119 | 1,25197688 | 0,8872305 | 1,12179564 | 8,6577E-07 | 1,12179564 | 0,26827184 | 0,78849009 |
| murB | 1,00555679 | -1,5034797 | -1,0323592 | -1,467E-06 | -1,0323592 | -0,2679896 | 0,78870734 |
| VC2242 | -1,0145193 | -0,67249 | -1,0733867 | -6,562E-07 | -1,0733867 | -0,2672232 | 0,78929731 |
| dipZ | 1,12865508 | 0,84406627 | 1,05791929 | 8,2365E-07 | 1,05791929 | 0,26696205 | 0,78949839 |
| pflA | 1,04001984 | 4,67022389 | 1,06004389 | 4,5572E-06 | 1,06004389 | 0,2667742 | 0,78964303 |
| VC1604 | 1,23472915 | 1,6966353 | 1,09865049 | 1,6556E-06 | 1,09865049 | 0,26539792 | 0,79070296 |
| VC1694 | 1,41835459 | 0,22023576 | 1,23787002 | 2,1491E-07 | 1,23787002 | 0,2650069 | 0,79100415 |
| VC0791 | 1,20627735 | 4,10480727 | 1,18360524 | 4,0055E-06 | 1,18360524 | 0,26492208 | 0,7910695 |
| ispH | 1,08051343 | 1,43620825 | 1,03817738 | 1,4015E-06 | 1,03817738 | 0,26458006 | 0,79133299 |
| VC2536 | -1,0435811 | -4,6013408 | -1,0365726 | -4,49E-06 | -1,0365726 | -0,2644589 | 0,79142636 |
| ftsB | 1,08806139 | 1,41983272 | 1,04557571 | 1,3855E-06 | 1,04557571 | 0,26422548 | 0,79160619 |
| VC0596 | 1,12691659 | 17,3073767 | 1,07136114 | 1,6889E-05 | 1,07136114 | 0,26421204 | 0,79161654 |
| VCA0197 | 1,11200904 | 0,95277661 | 1,05513127 | 9,2973E-07 | 1,05513127 | 0,26397457 | 0,79179952 |
| VC2564 | 1,19871932 | 0,55340224 | 1,09820963 | 5,4001E-07 | 1,09820963 | 0,26345347 | 0,79220109 |

| | | | | | | | |
|---|---|---|---|---|---|---|---|
| coaE | 1,13655165 | 0,84164059 | 1,05650935 | 8,2128E-07 | 1,05650935 | 0,26340379 | 0,79223938 |
| VC0970 | 1,06328692 | 6,1615862 | 1,03995065 | 6,0125E-06 | 1,03995065 | 0,2630395 | 0,79252015 |
| VC0103 | 1,0118055 | -0,9734727 | -1,0484868 | -9,499E-07 | -1,0484868 | -0,2629189 | 0,79261308 |
| VC1379 | 1,15629111 | 0,73073766 | 1,06500893 | 7,1306E-07 | 1,06500893 | 0,26270562 | 0,79277749 |
| VC0764 | 1,14170136 | 1,63191067 | 1,08660418 | 1,5924E-06 | 1,08660418 | 0,26203827 | 0,79329195 |
| VC0221 | 1,11632112 | 0,86606122 | 1,05427363 | 8,4511E-07 | 1,05427363 | 0,26200144 | 0,79332034 |
| VC2553 | 1,21884538 | 0,50386472 | 1,09467118 | 4,9168E-07 | 1,09467118 | 0,26137844 | 0,79380069 |
| VC1347 | 1,00948966 | -0,7225606 | -1,064522 | -7,051E-07 | -1,064522 | -0,2602822 | 0,79464612 |
| VCA0254 | 1,24841891 | 0,28738616 | 1,16829808 | 2,8043E-07 | 1,16829808 | 0,25868666 | 0,79587702 |
| uvrC | 1,04479867 | -1,6667615 | -1,0399389 | -1,626E-06 | -1,0399389 | -0,2583976 | 0,79610004 |
| VCA0316 | 1,10886518 | 0,88309016 | 1,0539862 | 8,6173E-07 | 1,0539862 | 0,25753729 | 0,79676404 |
| leuO | 1,10161019 | 0,85217483 | 1,05322305 | 8,3156E-07 | 1,05322305 | 0,25743067 | 0,79684633 |
| VC1524 | -1,0443865 | -1,8647856 | -1,1071816 | -1,82E-06 | -1,1071816 | -0,2573645 | 0,79689745 |
| VCA0466 | 1,20013592 | 1,04852297 | 1,07626332 | 1,0232E-06 | 1,07626332 | 0,25727408 | 0,79696721 |
| VC0968 | 1,07333901 | 29,4383201 | 1,14051447 | 2,8726E-05 | 1,14051447 | 0,25721407 | 0,79701353 |
| VC2343 | 1,10541667 | 1,16014709 | 1,03862151 | 1,1321E-06 | 1,03862151 | 0,25678519 | 0,79734461 |
| tRNA-Val-3 | -1,0933161 | -14,383877 | -1,0694286 | -1,404E-05 | -1,0694286 | -0,2565652 | 0,79751445 |
| VCA0606 | 1,06905744 | -0,3780153 | -1,1221943 | -3,689E-07 | -1,1221943 | -0,2555341 | 0,79831064 |
| VCA0779 | 1,14182156 | 0,7765096 | 1,08105037 | 7,5772E-07 | 1,08105037 | 0,255498 | 0,7983385 |
| VC1538 | 1,04195173 | 8,04233045 | 1,03746421 | 7,8478E-06 | 1,03746421 | 0,2550799 | 0,7986614 |
| purU | 1,10362501 | 3,67332152 | 1,07807146 | 3,5845E-06 | 1,07807146 | 0,25463222 | 0,79900719 |
| VC0919 | 1,02560618 | -0,488926 | -1,0924402 | -4,771E-07 | -1,0924402 | -0,2545583 | 0,79906425 |
| VC1440 | 1,06034394 | 8,05827122 | 1,05981859 | 7,8633E-06 | 1,05981859 | 0,25445059 | 0,79914748 |
| VCA0804 | -1,2597365 | -17,630205 | -1,1818113 | -1,72E-05 | -1,1818113 | -0,2538789 | 0,79958913 |
| VC1732 | 1,1006744 | 3,06075106 | 1,07032861 | 2,9867E-06 | 1,07032861 | 0,25356033 | 0,79983526 |
| VC2341 | 1,24590959 | 0,65774109 | 1,08997929 | 6,4183E-07 | 1,08997929 | 0,25276847 | 0,80044714 |
| VC1775 | -1,0330137 | -6,3726286 | -1,1344398 | -6,218E-06 | -1,1344398 | -0,2521292 | 0,80094124 |
| VC2326 | 1,00503697 | -22,265067 | -1,0926784 | -2,173E-05 | -1,0926784 | -0,2520539 | 0,8009994 |
| VCA0589 | 1,03617326 | -0,9619371 | -1,0449963 | -9,387E-07 | -1,0449963 | -0,2519884 | 0,80105002 |
| VCA1034 | 1,03114466 | -0,4830686 | -1,0914723 | -4,714E-07 | -1,0914723 | -0,251759 | 0,80122736 |
| VC1934 | -1,0137106 | -0,5089271 | -1,0866018 | -4,966E-07 | -1,0866018 | -0,2517291 | 0,80125046 |
| pcm | 1,03873242 | -1,2424336 | -1,0417458 | -1,212E-06 | -1,0417458 | -0,2514121 | 0,80149549 |
| VCA0442 | 1,13136508 | 0,97715801 | 1,08012684 | 9,5352E-07 | 1,08012684 | 0,25136008 | 0,80153574 |
| VC1040 | 1,11382365 | 1,42699742 | 1,07907929 | 1,3925E-06 | 1,07907929 | 0,25047292 | 0,80222165 |
| VC2630 | 1,14564252 | 0,60006051 | 1,07221172 | 5,8554E-07 | 1,07221172 | 0,25046484 | 0,8022279 |
| VC2638 | 1,22762861 | 0,46569395 | 1,0938409 | 4,5443E-07 | 1,0938409 | 0,25022822 | 0,80241087 |
| VCA0106 | 1,14978683 | 0,62948993 | 1,06848735 | 6,1426E-07 | 1,06848735 | 0,25005513 | 0,80254472 |
| VC1947 | 1,19981833 | 0,4764919 | 1,09134406 | 4,6496E-07 | 1,09134406 | 0,24987163 | 0,80268663 |
| metL | 1,15944138 | 0,6078132 | 1,0707345 | 5,9311E-07 | 1,0707345 | 0,24957496 | 0,80291607 |
| VC2076 | 1,13015019 | 6,37031974 | 1,14621329 | 6,2162E-06 | 1,14621329 | 0,24917214 | 0,80322764 |
| VC2407 | 1,12058007 | 0,8776027 | 1,04820501 | 8,5637E-07 | 1,04820501 | 0,24892752 | 0,80341685 |
| VC1399 | 1,187817 | 0,42241162 | 1,1025708 | 4,1219E-07 | 1,1025708 | 0,24863716 | 0,80364147 |
| VC2609 | 1,14813455 | 2,54478745 | 1,06978048 | 2,4832E-06 | 1,06978048 | 0,2486369 | 0,80364167 |
| VCA0119 | 1,13016396 | 0,46261508 | 1,09308167 | 4,5142E-07 | 1,09308167 | 0,24843377 | 0,80379881 |
| VC1004 | 1,01520105 | -1,1102287 | -1,0377143 | -1,083E-06 | -1,0377143 | -0,2482872 | 0,80391217 |

| | | | | | | | |
|---|---|---|---|---|---|---|---|
| VCA0992 | 1,33087154 | 0,22595005 | 1,19993929 | 2,2048E-07 | 1,19993929 | 0,24820517 | 0,80397567 |
| VC0864 | 1,00080755 | -1,230053 | -1,052846 | -1,2E-06 | -1,052846 | -0,2481153 | 0,80404518 |
| VC0920 | 1,226384 | 0,44126912 | 1,09751797 | 4,3059E-07 | 1,09751797 | 0,24808636 | 0,8040676 |
| VC2005 | -1,0769773 | -13,820806 | -1,1156524 | -1,349E-05 | -1,1156524 | -0,2479095 | 0,80420447 |
| VC1038 | 1,07256654 | 10,0873404 | 1,13080634 | 9,8433E-06 | 1,13080634 | 0,24776322 | 0,80431762 |
| VC0539 | 1,18264635 | 1,52449114 | 1,16984125 | 1,4876E-06 | 1,16984125 | 0,24735662 | 0,80463226 |
| yebU | -1,0122053 | -0,6637118 | -1,0687135 | -6,477E-07 | -1,0687135 | -0,2470564 | 0,80486458 |
| VC1643 | 1,0335502 | -0,4515931 | -1,0943007 | -4,407E-07 | -1,0943007 | -0,2469865 | 0,80491864 |
| murD | 1,01508818 | -1,1606281 | -1,0442072 | -1,133E-06 | -1,0442072 | -0,2469162 | 0,80497306 |
| VC2508 | 1,04996113 | -0,3952245 | -1,1083659 | -3,857E-07 | -1,1083659 | -0,2468634 | 0,80501393 |
| VCA0210 | 1,00230506 | -0,6005317 | -1,0699036 | -5,86E-07 | -1,0699036 | -0,2466637 | 0,80516855 |
| VC0949 | 1,09282931 | 1,62657777 | 1,04592085 | 1,5872E-06 | 1,04592085 | 0,24660545 | 0,8052136 |
| VC0231 | 1,067028 | 1,38252036 | 1,02969955 | 1,3491E-06 | 1,02969955 | 0,24635733 | 0,80540564 |
| VC2497 | 1,09611439 | 4,18828094 | 1,05461161 | 4,087E-06 | 1,05461161 | 0,24619739 | 0,80552945 |
| VC1421 | 1,01433377 | -0,7617695 | -1,0543698 | -7,433E-07 | -1,0543698 | -0,2459319 | 0,80573495 |
| VCA0550 | 1,0239274 | -1,2298976 | -1,0333051 | -1,2E-06 | -1,0333051 | -0,2458427 | 0,80580399 |
| VC0966 | 1,26319949 | 86,4921118 | 1,12515229 | 8,44E-05 | 1,12515229 | 0,24570583 | 0,80590997 |
| VC0144 | 1,04553845 | -2,500881 | -1,0627704 | -2,44E-06 | -1,0627704 | -0,2453068 | 0,80621892 |
| VC0169 | 1,21341479 | 0,41043517 | 1,10274555 | 4,0051E-07 | 1,10274555 | 0,24528555 | 0,80623535 |
| VCA1068 | -1,0079258 | -6,0646251 | -1,1077982 | -5,918E-06 | -1,1077982 | -0,2452718 | 0,80624598 |
| VC2473 | -1,07369 | -14,627691 | -1,0973864 | -1,427E-05 | -1,0973864 | -0,2447593 | 0,8066428 |
| VC0963 | 1,10516206 | 0,81395051 | 1,05011886 | 7,9426E-07 | 1,05011886 | 0,24432856 | 0,80697637 |
| VCA0262 | -1,0045852 | -0,6253336 | -1,0657214 | -6,102E-07 | -1,0657214 | -0,2443071 | 0,80699296 |
| fbpC | 1,37287321 | 3,98082175 | 1,29286245 | 3,8845E-06 | 1,29286245 | 0,24364384 | 0,80750668 |
| VCA0542 | 1,14595196 | 10,8878504 | 1,12428335 | 1,0624E-05 | 1,12428335 | 0,24294391 | 0,80804886 |
| VC1585 | 1,01981142 | -0,8251314 | -1,0507239 | -8,052E-07 | -1,0507239 | -0,2426454 | 0,8082801 |
| VCA1003 | 1,01748057 | -2,4183351 | -1,095569 | -2,36E-06 | -1,095569 | -0,2416484 | 0,80905263 |
| VC0146 | -1,0194602 | -3,3259063 | -1,0356092 | -3,245E-06 | -1,0356092 | -0,2411564 | 0,80943392 |
| VC2067 | -1,0287408 | -4,3084888 | -1,0287797 | -4,204E-06 | -1,0287797 | -0,2409428 | 0,80959942 |
| VCA1059 | 1,10819704 | 0,81270472 | 1,05775866 | 7,9304E-07 | 1,05775866 | 0,24073099 | 0,80976363 |
| artM | 1,25857355 | 1,26229186 | 1,20250943 | 1,2318E-06 | 1,20250943 | 0,24063443 | 0,80983847 |
| VCA0194 | 1,05254751 | -0,9828722 | -1,0430869 | -9,591E-07 | -1,0430869 | -0,2402971 | 0,81009993 |
| asnB | 1,09105402 | -43,908497 | -1,0855186 | -4,285E-05 | -1,0855186 | -0,2400216 | 0,81031353 |
| VC2439 | -1,009952 | -4,7761959 | -1,0737032 | -4,661E-06 | -1,0737032 | -0,2398755 | 0,81042679 |
| VCA0852 | 1,01540193 | -0,4650836 | -1,0859936 | -4,538E-07 | -1,0859936 | -0,2398302 | 0,81046191 |
| VC1806 | 1,23761652 | 0,36673465 | 1,11005099 | 3,5786E-07 | 1,11005099 | 0,23954541 | 0,8106827 |
| VC2546 | -1,0238453 | -2,184204 | -1,0236591 | -2,131E-06 | -1,0236591 | -0,2393696 | 0,81081904 |
| VCA0415 | 1,12130484 | 0,75324624 | 1,06999538 | 7,3502E-07 | 1,06999538 | 0,239339 | 0,81084274 |
| VCA0600 | 1,12843756 | 0,63474442 | 1,06133029 | 6,1939E-07 | 1,06133029 | 0,23802691 | 0,81186023 |
| VCA0529 | -1,0406862 | -0,2502185 | -1,1630676 | -2,442E-07 | -1,1630676 | -0,2378863 | 0,8119693 |
| VC0585 | -1,0699207 | -5,5361819 | -1,0440982 | -5,402E-06 | -1,0440982 | -0,2377394 | 0,81208322 |
| VC0783 | 1,20054948 | 0,40420887 | 1,09740816 | 3,9443E-07 | 1,09740816 | 0,23731253 | 0,81241435 |
| VC0665 | 1,04770366 | -1,1273041 | -1,0524768 | -1,1E-06 | -1,0524768 | -0,2369063 | 0,8127295 |
| VC1130 | 1,01508246 | -7,2019368 | -1,0223402 | -7,028E-06 | -1,0223402 | -0,2367925 | 0,81281779 |
| VC0041 | 1,07335011 | 1,36117511 | 1,02775925 | 1,3282E-06 | 1,02775925 | 0,23644159 | 0,81309003 |

| | | | | | | | |
|---|---|---|---|---|---|---|---|
| VC0921 | 1,18404728 | 0,32701925 | 1,1206968 | 3,1911E-07 | 1,1206968 | 0,23629606 | 0,81320295 |
| tarB | -1,0232695 | -1,624687 | -1,0583282 | -1,585E-06 | -1,0583282 | -0,2361024 | 0,81335324 |
| VCA0688 | 1,20086541 | 0,35571832 | 1,1096554 | 3,4711E-07 | 1,1096554 | 0,23551779 | 0,81380688 |
| VCA0574 | 1,00836639 | -1,4996238 | -1,0844227 | -1,463E-06 | -1,0844227 | -0,2354308 | 0,8138744 |
| VC2115 | 1,00642416 | -0,8488037 | -1,0444816 | -8,283E-07 | -1,0444816 | -0,2353787 | 0,81391482 |
| VCA1090 | -1,0685602 | -0,3440159 | -1,1133775 | -3,357E-07 | -1,1133775 | -0,235302 | 0,81397438 |
| VC1551 | 1,25586405 | 0,26959046 | 1,14666722 | 2,6307E-07 | 1,14666722 | 0,23507014 | 0,8141543 |
| VC1933 | 1,13212056 | 0,54206505 | 1,07026845 | 5,2895E-07 | 1,07026845 | 0,23493888 | 0,81425618 |
| VC1581 | 1,4072428 | 0,17790145 | 1,23040396 | 1,736E-07 | 1,23040396 | 0,23480306 | 0,8143616 |
| VCA0960 | 1,07702958 | 1,26644394 | 1,02938396 | 1,2358E-06 | 1,02938396 | 0,23455027 | 0,81455782 |
| VCA0361a | -1,8E+308 | -0,0567154 | -1,9542881 | -5,534E-08 | -1,9542881 | -0,2344361 | 0,81464648 |
| torD | 1,21683981 | 0,38048596 | 1,10116797 | 3,7128E-07 | 1,10116797 | 0,23443471 | 0,81464752 |
| VC0093 | -1,0355345 | -0,847796 | -1,0609413 | -8,273E-07 | -1,0609413 | -0,2339326 | 0,81503733 |
| VCA0257 | 1,23029049 | 0,40531754 | 1,09393559 | 3,9551E-07 | 1,09393559 | 0,2335569 | 0,815329 |
| VCA0601 | 1,26389848 | 0,26754367 | 1,14576311 | 2,6107E-07 | 1,14576311 | 0,23350238 | 0,81537133 |
| VC0265 | -1,0046306 | -0,7306934 | -1,0508663 | -7,13E-07 | -1,0508663 | -0,2331726 | 0,81562737 |
| VC2771 | 1,19450931 | 31,3899879 | 1,06008584 | 3,0631E-05 | 1,06008584 | 0,23293719 | 0,81581019 |
| VC1608 | 1,14930886 | 0,80173682 | 1,09228764 | 7,8234E-07 | 1,09228764 | 0,23272266 | 0,81597678 |
| VC2031 | -1,0310441 | -0,4473501 | -1,1015669 | -4,365E-07 | -1,1015669 | -0,2323538 | 0,81626321 |
| VC1968 | 1,032215 | -1,4347506 | -1,0305206 | -1,4E-06 | -1,0305206 | -0,2323521 | 0,81626455 |
| VC1894 | -1,0073813 | -7,264565 | -1,0242456 | -7,089E-06 | -1,0242456 | -0,232281 | 0,81631975 |
| tRNA-Met-6 | -1,0178958 | -0,5989225 | -1,0617199 | -5,844E-07 | -1,0617199 | -0,2319241 | 0,81659701 |
| VC0693 | -1,0009058 | -0,7019778 | -1,0524171 | -6,85E-07 | -1,0524171 | -0,2319151 | 0,81660399 |
| VCA0042 | 1,08061617 | 1,47452383 | 1,03208812 | 1,4389E-06 | 1,03208812 | 0,23146102 | 0,81695667 |
| VC0147 | -1,010547 | -2,8244652 | -1,0369536 | -2,756E-06 | -1,0369536 | -0,2309847 | 0,81732667 |
| fliL | -1,0046041 | -1,7103141 | -1,0305864 | -1,669E-06 | -1,0305864 | -0,2309326 | 0,81736715 |
| tRNA-Asp-1 | 1,11722477 | 2,27864798 | 1,11777805 | 2,2235E-06 | 1,11777805 | 0,23022019 | 0,81792069 |
| VC1005 | -1,0324995 | -1,4211915 | -1,077903 | -1,387E-06 | -1,077903 | -0,2300825 | 0,81802771 |
| VC0196 | 1,11081947 | 0,71991545 | 1,05131837 | 7,025E-07 | 1,05131837 | 0,22941626 | 0,81854541 |
| VC1015 | 1,09671344 | 1,01625328 | 1,03475879 | 9,9167E-07 | 1,03475879 | 0,22821479 | 0,81947927 |
| VC0496 | 1,16612561 | 0,52658641 | 1,06799637 | 5,1385E-07 | 1,06799637 | 0,22791092 | 0,8197155 |
| VC0922 | 1,13386925 | 0,53824302 | 1,06636839 | 5,2522E-07 | 1,06636839 | 0,22773426 | 0,81985284 |
| VC0072 | 1,09101415 | 1,06443158 | 1,05913016 | 1,0387E-06 | 1,05913016 | 0,22727545 | 0,82020957 |
| VC1389 | -1,0338191 | -0,3549052 | -1,1019362 | -3,463E-07 | -1,1019362 | -0,2272333 | 0,82024233 |
| VC2270 | 1,01417907 | -2,0004789 | -1,0359718 | -1,952E-06 | -1,0359718 | -0,2272054 | 0,82026406 |
| VCA0343 | 1,0396062 | -0,8996372 | -1,0389792 | -8,779E-07 | -1,0389792 | -0,2271484 | 0,82030839 |
| VCA0786 | 1,00935329 | -0,7531798 | -1,0465793 | -7,35E-07 | -1,0465793 | -0,2267751 | 0,8205986 |
| VC1911 | 1,00644181 | 2,89871986 | 1,03135818 | 2,8286E-06 | 1,03135818 | 0,22661989 | 0,82071933 |
| VC1247 | 1,03998407 | 11,1432549 | 1,04369593 | 1,0874E-05 | 1,04369593 | 0,22590738 | 0,82127346 |
| VCA0380 | 1,2194315 | 0,3943673 | 1,08951329 | 3,8483E-07 | 1,08951329 | 0,22512993 | 0,8218782 |
| VC0903 | 1,12260641 | 0,73418867 | 1,04685433 | 7,1643E-07 | 1,04685433 | 0,22454286 | 0,82233493 |
| VCA0096 | 1,15476471 | 0,56092491 | 1,06172293 | 5,4736E-07 | 1,06172293 | 0,2244518 | 0,82240577 |
| VC0808 | 1,13484503 | 0,55818211 | 1,06166633 | 5,4468E-07 | 1,06166633 | 0,22380275 | 0,8229108 |
| fadE | 1,05499359 | -0,3815691 | -1,0913448 | -3,723E-07 | -1,0913448 | -0,2236028 | 0,82306637 |
| VCA0958 | 1,2396971 | 0,38173121 | 1,09124618 | 3,725E-07 | 1,09124618 | 0,22353481 | 0,8231193 |

| | | | | | | | |
|---|---|---|---|---|---|---|---|
| VCA0754 | 1,16841643 | 3,29751124 | 1,17246082 | 3,2177E-06 | 1,17246082 | 0,22297469 | 0,82355521 |
| VC1746 | 1,17350468 | 0,45532238 | 1,07549844 | 4,4431E-07 | 1,07549844 | 0,22290984 | 0,82360569 |
| VC0938 | 1,18534056 | 0,37541039 | 1,09184779 | 3,6633E-07 | 1,09184779 | 0,222374 | 0,82402276 |
| VC1931 | 1,20628424 | 0,35108275 | 1,09817381 | 3,4259E-07 | 1,09817381 | 0,22199504 | 0,82431775 |
| VCA0720 | 1,00367114 | -0,3521 | -1,0972085 | -3,436E-07 | -1,0972085 | -0,2212716 | 0,82488096 |
| VC2500 | 1,00362704 | -1,1222601 | -1,0307784 | -1,095E-06 | -1,0307784 | -0,2211216 | 0,82499777 |
| VCA0141 | 1,00317596 | -1,0104397 | -1,0888171 | -9,86E-07 | -1,0888171 | -0,2206696 | 0,82534968 |
| fadJ | 1,18502651 | 0,95489538 | 1,07956728 | 9,318E-07 | 1,07956728 | 0,22013761 | 0,82576399 |
| VC1766 | 1,0632076 | 2,8193129 | 1,05434644 | 2,7511E-06 | 1,05434644 | 0,219886 | 0,82595995 |
| ppnK | 1,11668523 | 1,10224772 | 1,03399836 | 1,0756E-06 | 1,03399836 | 0,21908686 | 0,8265824 |
| VC1162 | -1,0055598 | -0,6018212 | -1,0544268 | -5,873E-07 | -1,0544268 | -0,2187045 | 0,82688027 |
| VC0722 | -1,018313 | -3,8965532 | -1,0768928 | -3,802E-06 | -1,0768928 | -0,2186649 | 0,82691109 |
| VC1213 | 1,07789692 | 8,05246632 | 1,04475248 | 7,8577E-06 | 1,04475248 | 0,21801227 | 0,82741956 |
| VC1730 | 1,08857738 | 4,6950586 | 1,04753763 | 4,5815E-06 | 1,04753763 | 0,21798179 | 0,82744331 |
| VCA0146 | -1,0307536 | -0,2155755 | -1,1583052 | -2,104E-07 | -1,1583052 | -0,2177969 | 0,82758735 |
| VC1179 | -1,0015161 | -1,5530665 | -1,038111 | -1,515E-06 | -1,038111 | -0,2176981 | 0,82766433 |
| VC1465 | 1,00537445 | -0,7198673 | -1,0447091 | -7,025E-07 | -1,0447091 | -0,2173064 | 0,82796961 |
| VC1429 | 1,09006038 | 0,81761558 | 1,03924047 | 7,9784E-07 | 1,03924047 | 0,21725641 | 0,82800853 |
| VCA0776 | 1,15758218 | 0,59986109 | 1,05385222 | 5,8535E-07 | 1,05385222 | 0,2172229 | 0,82803465 |
| VC1856 | 1,0089186 | -1,3941233 | -1,0358931 | -1,36E-06 | -1,0358931 | -0,2170618 | 0,82816022 |
| VC1647 | 1,2346427 | 0,35449237 | 1,09263392 | 3,4592E-07 | 1,09263392 | 0,21697181 | 0,82823032 |
| VC0595 | 1,20394775 | 0,55190653 | 1,06864241 | 5,3856E-07 | 1,06864241 | 0,21689661 | 0,82828892 |
| VCA0926 | 1,14793086 | 0,4752444 | 1,06775047 | 4,6375E-07 | 1,06775047 | 0,21613632 | 0,8288815 |
| VC0317 | 1,12898072 | 0,62635729 | 1,05098189 | 6,112E-07 | 1,05098189 | 0,21612321 | 0,82889172 |
| cpdB | 1,01482976 | -3,1434856 | -1,0922714 | -3,067E-06 | -1,0922714 | -0,2160086 | 0,82898107 |
| VCA0085 | 1,23198142 | 0,35963099 | 1,08995515 | 3,5093E-07 | 1,08995515 | 0,21549371 | 0,82938242 |
| tRNA-Met-7 | -1,0802881 | -3,385983 | -1,063794 | -3,304E-06 | -1,063794 | -0,2153871 | 0,82946554 |
| VCA0935 | 1,1759352 | 1,20991162 | 1,12132208 | 1,1806E-06 | 1,12132208 | 0,21532986 | 0,82951016 |
| VCA0632 | 1,16425532 | 0,67634746 | 1,05519531 | 6,5999E-07 | 1,05519531 | 0,21529509 | 0,82953727 |
| VC0995 | 1,14381856 | 10,9429088 | 1,08920616 | 1,0678E-05 | 1,08920616 | 0,21516578 | 0,82963808 |
| tRNA-Ala-3 | -1,056794 | -1,2102147 | -1,1207169 | -1,181E-06 | -1,1207169 | -0,2146841 | 0,83001365 |
| VC1269 | 1,1112723 | 5,35789245 | 1,07711858 | 5,2283E-06 | 1,07711858 | 0,21453482 | 0,83013003 |
| VC0847 | 1,15452883 | 1,42985607 | 1,06734239 | 1,3953E-06 | 1,06734239 | 0,21449641 | 0,83015997 |
| VC1777 | -1,0015674 | -6,923622 | -1,1729238 | -6,756E-06 | -1,1729238 | -0,2143339 | 0,83028666 |
| VC1788 | 1,10931833 | 0,58089397 | 1,05395622 | 5,6684E-07 | 1,05395622 | 0,21396196 | 0,83057673 |
| dinG | 1,09096826 | 1,52229603 | 1,04945034 | 1,4855E-06 | 1,04945034 | 0,2137992 | 0,83070366 |
| VC0498 | -1,0202586 | -0,3388434 | -1,0941451 | -3,306E-07 | -1,0941451 | -0,2137748 | 0,83072267 |
| VC1909 | -1,0236843 | -2,691216 | -1,0191079 | -2,626E-06 | -1,0191079 | -0,213146 | 0,83121308 |
| VCA0895 | 1,1932165 | 0,44937618 | 1,06963975 | 4,3851E-07 | 1,06963975 | 0,21298465 | 0,83133895 |
| VC1081 | 1,17985261 | 3,48202873 | 1,09662253 | 3,3978E-06 | 1,09662253 | 0,21209733 | 0,83203112 |
| VCA0074 | 1,09880283 | 0,79882007 | 1,0601886 | 7,795E-07 | 1,0601886 | 0,21201355 | 0,83209647 |
| VCA1015 | 1,12578758 | 0,7253053 | 1,05795327 | 7,0776E-07 | 1,05795327 | 0,21163808 | 0,8323894 |
| VCA0727 | 1,00394598 | -0,5556954 | -1,0546757 | -5,423E-07 | -1,0546757 | -0,2106235 | 0,83318106 |
| VCA1054 | 1,13240257 | 0,3468403 | 1,08904229 | 3,3845E-07 | 1,08904229 | 0,21059634 | 0,83320227 |
| VC1077 | 1,09135328 | 13,7850064 | 1,06200594 | 1,3452E-05 | 1,06200594 | 0,21021116 | 0,83350288 |

| | | | | | | | |
|---|---|---|---|---|---|---|---|
| VC0910 | -1,0681526 | -63,315412 | -1,2281453 | -6,178E-05 | -1,2281453 | -0,2101304 | 0,83356588 |
| VCA0690 | 1,26635338 | 0,19827384 | 1,1602095 | 1,9348E-07 | 1,1602095 | 0,2100341 | 0,83364106 |
| VC2303 | -1,0792471 | -3,1653667 | -1,0932535 | -3,089E-06 | -1,0932535 | -0,2100288 | 0,83364524 |
| prmA | 1,09104513 | 0,82690604 | 1,05834588 | 8,069E-07 | 1,05834588 | 0,2098449 | 0,83378873 |
| VCA0961 | 1,16688835 | 0,45395177 | 1,06595936 | 4,4297E-07 | 1,06595936 | 0,20851837 | 0,83482425 |
| apaG | 1,08847964 | 1,34770694 | 1,03368837 | 1,3151E-06 | 1,03368837 | 0,20743639 | 0,83566907 |
| VC2208 | 1,04809045 | -1,1826647 | -1,0438756 | -1,154E-06 | -1,0438756 | -0,2071452 | 0,83589647 |
| VCA0080 | 1,10801545 | 0,72991078 | 1,03993233 | 7,1225E-07 | 1,03993233 | 0,20703983 | 0,83597876 |
| VC0084 | 1,02807955 | -1,0260791 | -1,0281628 | -1,001E-06 | -1,0281628 | -0,2067499 | 0,8362052 |
| qrr4 | 1,02916476 | -1,6394345 | -1,054573 | -1,6E-06 | -1,054573 | -0,2066327 | 0,83629676 |
| VCA0863 | 1,03485873 | -0,5739468 | -1,0671228 | -5,601E-07 | -1,0671228 | -0,2066216 | 0,83630539 |
| VC2504 | 1,24021682 | 2,09201871 | 1,09768276 | 2,0414E-06 | 1,09768276 | 0,20655638 | 0,83635634 |
| VC2094 | -1,0105683 | -4,0953702 | -1,0702937 | -3,996E-06 | -1,0702937 | -0,2064998 | 0,83640055 |
| VC1325 | 1,23855287 | 62,039306 | 1,1437537 | 6,0538E-05 | 1,1437537 | 0,20642595 | 0,83645822 |
| VC2557 | 1,18713901 | 1,35537525 | 1,05773015 | 1,3226E-06 | 1,05773015 | 0,20638662 | 0,83648893 |
| VC0819 | 1,1894073 | 0,35560847 | 1,08295933 | 3,4701E-07 | 1,08295933 | 0,20583642 | 0,8369187 |
| VC0463 | 1,0368637 | -0,5887202 | -1,048451 | -5,745E-07 | -1,048451 | -0,2043884 | 0,83805003 |
| VC2082 | 1,12540511 | 0,57834431 | 1,04932655 | 5,6435E-07 | 1,04932655 | 0,20435779 | 0,83807392 |
| VC1197 | -1,0331247 | -7,4385864 | -1,0697832 | -7,259E-06 | -1,0697832 | -0,2043233 | 0,83810086 |
| VC0129 | -1,0031534 | -1,1429507 | -1,0246107 | -1,115E-06 | -1,0246107 | -0,2041624 | 0,83822662 |
| VC1017 | 1,11512968 | 0,75594642 | 1,03723378 | 7,3766E-07 | 1,03723378 | 0,20359104 | 0,83867311 |
| VC1305 | 1,00911874 | -1,7325801 | -1,0476074 | -1,691E-06 | -1,0476074 | -0,2031396 | 0,83902592 |
| VC0678 | 1,25669663 | 17,6372009 | 1,04102428 | 1,7211E-05 | 1,04102428 | 0,20310646 | 0,83905184 |
| VCA0813 | 1,15212218 | 0,41846304 | 1,06779648 | 4,0834E-07 | 1,06779648 | 0,20288045 | 0,83922848 |
| tRNA-Ile-3 | 1,09188816 | 19,4596677 | 1,08294893 | 1,8989E-05 | 1,08294893 | 0,20258675 | 0,83945806 |
| VC2489 | 1,22246351 | 0,40865117 | 1,06921535 | 3,9877E-07 | 1,06921535 | 0,20250542 | 0,83952162 |
| VCA0426 | 1,06342329 | 2,3311096 | 1,04659751 | 2,2747E-06 | 1,04659751 | 0,20217949 | 0,83977641 |
| VC1442 | 1,12790525 | 7,26855894 | 1,07011846 | 7,0927E-06 | 1,07011846 | 0,20200342 | 0,83991407 |
| VC1153 | 1,13381558 | 14,8357791 | 1,10025135 | 1,4477E-05 | 1,10025135 | 0,20144817 | 0,84034816 |
| VC1083 | 1,01303534 | -1,9744801 | -1,0769273 | -1,927E-06 | -1,0769273 | -0,2010238 | 0,84067998 |
| VC1507 | 1,12673001 | 1,98044326 | 1,05308368 | 1,9325E-06 | 1,05308368 | 0,20057805 | 0,84102853 |
| VCA0788 | 1,01583771 | -2,8890559 | -1,0538095 | -2,819E-06 | -1,0538095 | -0,2005569 | 0,84104504 |
| VC2599 | -1,0226596 | -2,5771654 | -1,0288867 | -2,515E-06 | -1,0288867 | -0,1998977 | 0,84156062 |
| VC1120 | 1,0495648 | -0,9271377 | -1,0643969 | -9,047E-07 | -1,0643969 | -0,1998344 | 0,84161012 |
| ugpC | 1,37766536 | 0,16563027 | 1,17476926 | 1,6162E-07 | 1,17476926 | 0,19982813 | 0,841615 |
| VC0957 | -1,0535892 | -22,084593 | -1,1037333 | -2,155E-05 | -1,1037333 | -0,1992363 | 0,84207788 |
| lplA | 1,11062453 | 0,49977741 | 1,06071638 | 4,8769E-07 | 1,06071638 | 0,19895213 | 0,8423002 |
| VCA0222 | 1,21630232 | 0,26319854 | 1,10520408 | 2,5683E-07 | 1,10520408 | 0,19864247 | 0,84254243 |
| VC0887 | 1,0211596 | -0,4961911 | -1,0542521 | -4,842E-07 | -1,0542521 | -0,1982753 | 0,84282968 |
| VC1664 | 1,00882379 | -0,5966456 | -1,0448252 | -5,822E-07 | -1,0448252 | -0,1980865 | 0,84297738 |
| VCA1020 | 1,02611929 | -1,175428 | -1,0223736 | -1,147E-06 | -1,0223736 | -0,1975183 | 0,84342195 |
| VC1560 | -1,0043598 | -0,9595732 | -1,0560516 | -9,364E-07 | -1,0560516 | -0,197509 | 0,84342924 |
| VC1782 | 1,03296903 | -2,769009 | -1,1182384 | -2,702E-06 | -1,1182384 | -0,1973998 | 0,84351473 |
| VC1660a | 1,03305997 | -0,5288 | -1,0527624 | -5,16E-07 | -1,0527624 | -0,1972064 | 0,84366606 |
| VC1897 | -1,0366493 | -3,1695158 | -1,0467875 | -3,093E-06 | -1,0467875 | -0,197021 | 0,84381111 |

| | | | | | | | |
|---|---|---|---|---|---|---|---|
| envZ | 1,04882864 | -0,7064887 | -1,0372421 | -6,894E-07 | -1,0372421 | -0,1968399 | 0,84395284 |
| cyaA | 1,04377338 | 13,3769431 | 1,12432929 | 1,3053E-05 | 1,12432929 | 0,19656562 | 0,84416749 |
| VC1870 | 1,02348255 | -0,4069124 | -1,0651245 | -3,971E-07 | -1,0651245 | -0,1962057 | 0,84444916 |
| VCA0007 | 1,09351381 | 1,82945043 | 1,05648274 | 1,7852E-06 | 1,05648274 | 0,19603369 | 0,8445838 |
| VC1156 | 1,02621276 | -3,0740796 | -1,0868656 | -3E-06 | -1,0868656 | -0,1957603 | 0,8447978 |
| VCA0988 | -1,0609283 | -0,2352499 | -1,1146457 | -2,296E-07 | -1,1146457 | -0,1956077 | 0,84491725 |
| VC1422a | 1,07256426 | 1,07469648 | 1,02389496 | 1,0487E-06 | 1,02389496 | 0,19510678 | 0,84530936 |
| VCA1100 | 1,22291482 | 0,20645342 | 1,13084228 | 2,0146E-07 | 1,13084228 | 0,19501608 | 0,84538037 |
| VC0099 | 1,09105429 | 0,92902157 | 1,02762212 | 9,0655E-07 | 1,02762212 | 0,19485689 | 0,84550499 |
| VCA0544 | 1,19963729 | 1,27325762 | 1,06773314 | 1,2425E-06 | 1,06773314 | 0,1947541 | 0,84558546 |
| VCA0297 | 1,10553894 | 0,48713746 | 1,0537879 | 4,7535E-07 | 1,0537879 | 0,19424462 | 0,84598435 |
| VCA0649 | 1,36282582 | 0,12794008 | 1,21800303 | 1,2485E-07 | 1,21800303 | 0,19422909 | 0,84599651 |
| VCA0854 | 1,03871532 | -0,1503692 | -1,181659 | -1,467E-07 | -1,181659 | -0,1938096 | 0,84632497 |
| VC0095 | 1,12232068 | 0,78469677 | 1,0321284 | 7,6571E-07 | 1,0321284 | 0,19292379 | 0,84701866 |
| VC1229 | 1,25228176 | 0,23233452 | 1,11246679 | 2,2671E-07 | 1,11246679 | 0,19263499 | 0,84724485 |
| VC1598 | 1,1347298 | 0,64395101 | 1,03916711 | 6,2837E-07 | 1,03916711 | 0,19263045 | 0,8472484 |
| purT | -1,0251359 | -0,9393093 | -1,0698711 | -9,166E-07 | -1,0698711 | -0,1926121 | 0,8472628 |
| VCA1092 | -1,0519843 | -0,2723947 | -1,0951129 | -2,658E-07 | -1,0951129 | -0,192609 | 0,84726517 |
| lspA | -1,0099776 | -2,040003 | -1,0432503 | -1,991E-06 | -1,0432503 | -0,1925313 | 0,84732606 |
| cobS | 1,00922463 | -1,1930502 | -1,0767243 | -1,164E-06 | -1,0767243 | -0,1922264 | 0,84756487 |
| VC1006 | 1,09980941 | 2,14552994 | 1,06768391 | 2,0936E-06 | 1,06768391 | 0,19107196 | 0,84846923 |
| VC2318 | -1,0234868 | -1,0074072 | -1,047328 | -9,83E-07 | -1,047328 | -0,1906995 | 0,84876101 |
| alr (NC_002505 387951..389119) | 1,02870496 | -0,9652087 | -1,0254223 | -9,419E-07 | -1,0254223 | -0,1906466 | 0,84880245 |
| VCA0102 | 1,00207836 | -0,9421495 | -1,0260729 | -9,194E-07 | -1,0260729 | -0,1906351 | 0,84881149 |
| VC2162 | 1,01363094 | -0,9603057 | -1,0254535 | -9,371E-07 | -1,0254535 | -0,1902771 | 0,84909203 |
| VC0728 | 1,04863811 | -0,9734418 | -1,0391553 | -9,499E-07 | -1,0391553 | -0,190188 | 0,84916184 |
| VCA0505 | 1,02088302 | -1,6642658 | -1,0387852 | -1,624E-06 | -1,0387852 | -0,1899018 | 0,84938608 |
| VC2111 | -1,0059936 | -1,2293466 | -1,0314795 | -1,2E-06 | -1,0314795 | -0,1898012 | 0,84946491 |
| VCA1010 | 1,05290833 | -0,3617054 | -1,0683557 | -3,53E-07 | -1,0683557 | -0,1893713 | 0,84980186 |
| VC2310 | 1,16886065 | 0,57800914 | 1,0893421 | 5,6403E-07 | 1,0893421 | 0,18934438 | 0,84982292 |
| VC0851 | 1,07574878 | 4,29258671 | 1,03718095 | 4,1887E-06 | 1,03718095 | 0,18928457 | 0,84986979 |
| VC1997 | 1,05084059 | -1,8901315 | -1,048532 | -1,844E-06 | -1,048532 | -0,188757 | 0,85028332 |
| VCA0464 | 1,0738725 | 0,84706203 | 1,02833877 | 8,2657E-07 | 1,02833877 | 0,18842776 | 0,85054135 |
| VC1310 | 1,07018249 | 1,01293564 | 1,02778923 | 9,8843E-07 | 1,02778923 | 0,18824721 | 0,85068287 |
| VC0992 | 1,02117934 | -0,6185694 | -1,0389216 | -6,036E-07 | -1,0389216 | -0,1882145 | 0,8507085 |
| VC0868 | -1,0548642 | -0,2215879 | -1,1122188 | -2,162E-07 | -1,1122188 | -0,1879306 | 0,85093106 |
| VC1953 | 1,20334631 | 0,22232959 | 1,11178542 | 2,1695E-07 | 1,11178542 | 0,18790027 | 0,85095484 |
| VC2550 | 1,0830616 | 3,23277131 | 1,05983096 | 3,1546E-06 | 1,05983096 | 0,18787878 | 0,85097169 |
| VCA1019 | 1,23549765 | 0,27139686 | 1,09043831 | 2,6483E-07 | 1,09043831 | 0,1876814 | 0,85112642 |
| VC1918 | 1,04068207 | 5,17278133 | 1,03539048 | 5,0476E-06 | 1,03539048 | 0,18755388 | 0,85122639 |
| VC1122 | -1,0411371 | -2,8609923 | -1,0646644 | -2,792E-06 | -1,0646644 | -0,1874878 | 0,8512782 |
| VC2455 | 1,02387995 | -0,435227 | -1,0552641 | -4,247E-07 | -1,0552641 | -0,1873735 | 0,85136782 |
| mraY | 1,01437587 | -1,0207317 | -1,0231254 | -9,96E-07 | -1,0231254 | -0,1870935 | 0,85158731 |
| VC2677 | 1,02557275 | -6,3536084 | -1,0740335 | -6,2E-06 | -1,0740335 | -0,1869374 | 0,85170971 |
| VC0303 | -1,0055273 | -0,8646634 | -1,0730391 | -8,437E-07 | -1,0730391 | -0,1867281 | 0,85187381 |

| | | | | | | | |
|---|---|---|---|---|---|---|---|
| VCA0766 | 1,01517471 | -0,4955817 | -1,0479864 | -4,836E-07 | -1,0479864 | -0,1866449 | 0,85193905 |
| VC0779 | 1,01751053 | -0,3482103 | -1,0687867 | -3,398E-07 | -1,0687867 | -0,1863704 | 0,85215435 |
| VC1003 | 1,06945462 | 1,29199684 | 1,03289917 | 1,2607E-06 | 1,03289917 | 0,18602533 | 0,8524249 |
| ectC | -1,0063159 | -0,4245026 | -1,0637236 | -4,142E-07 | -1,0637236 | -0,1855295 | 0,85281372 |
| VCA0990 | 1,22175677 | 0,33956107 | 1,06987877 | 3,3135E-07 | 1,06987877 | 0,18544743 | 0,85287812 |
| VC0038 | 1,07381666 | 7,24900306 | 1,05712222 | 7,0736E-06 | 1,05712222 | 0,18486237 | 0,85333699 |
| VC0481 | 1,07557407 | 2,17913371 | 1,06089594 | 2,1264E-06 | 1,06089594 | 0,18453229 | 0,8535959 |
| VCA0930 | 1,09133625 | 0,67628901 | 1,03409136 | 6,5993E-07 | 1,03409136 | 0,18440301 | 0,85369732 |
| VC0235 | 1,07541464 | 1,79080595 | 1,03735536 | 1,7475E-06 | 1,03735536 | 0,18433707 | 0,85374904 |
| VCA0025 | 1,15554684 | 5,86854593 | 1,1073906 | 5,7266E-06 | 1,1073906 | 0,18421645 | 0,85384366 |
| zntA | 1,06807989 | 1,87693097 | 1,04391745 | 1,8315E-06 | 1,04391745 | 0,18408509 | 0,85394671 |
| VC2695 | 1,02423289 | -2,0224743 | -1,0430848 | -1,974E-06 | -1,0430848 | -0,1838676 | 0,85411733 |
| VCA0615 | 1,04919509 | -0,3505139 | -1,0662691 | -3,42E-07 | -1,0662691 | -0,183644 | 0,85429278 |
| glgC (NC_002505 1865312..1866570) | -1,0060319 | -5,876606 | -1,0587399 | -5,734E-06 | -1,0587399 | -0,183461 | 0,85443635 |
| VCA0484 | 1,03989538 | -1,3490184 | -1,0567694 | -1,316E-06 | -1,0567694 | -0,1832844 | 0,85457488 |
| VC1475 | -1,027444 | -0,1681044 | -1,1421207 | -1,64E-07 | -1,1421207 | -0,1829183 | 0,85486215 |
| VC1433 | -1,027226 | -4,653088 | -1,0576601 | -4,541E-06 | -1,0576601 | -0,1827978 | 0,85495665 |
| VC2300 | 1,05234003 | -1,0727664 | -1,0275147 | -1,047E-06 | -1,0275147 | -0,1825207 | 0,85517412 |
| VC2463 | -1,0226144 | -3,2928043 | -1,0248548 | -3,213E-06 | -1,0248548 | -0,1824459 | 0,85523281 |
| VC1867 | 1,0724246 | -1,1399126 | -1,0493553 | -1,112E-06 | -1,0493553 | -0,18222 | 0,85541009 |
| VC1875 | 1,09668024 | 2,69640918 | 1,05745567 | 2,6312E-06 | 1,05745567 | 0,18154755 | 0,85593782 |
| VC0755 | 1,02917193 | -3,8758841 | -1,0474337 | -3,782E-06 | -1,0474337 | -0,1814906 | 0,85598256 |
| tRNA-Gly-3 | -1,0767309 | -1,8883733 | -1,0632858 | -1,843E-06 | -1,0632858 | -0,1809837 | 0,85638035 |
| VC2631 | -1,007335 | -0,2821796 | -1,0803526 | -2,754E-07 | -1,0803526 | -0,1808239 | 0,85650583 |
| obgE | 1,06157936 | 3,86742867 | 1,06510241 | 3,7739E-06 | 1,06510241 | 0,1806522 | 0,85664059 |
| VC1057 | -1,0012345 | -0,472146 | -1,0470958 | -4,607E-07 | -1,0470958 | -0,1805191 | 0,85674508 |
| ubiB | 1,06254761 | 0,99231233 | 1,02464166 | 9,6831E-07 | 1,02464166 | 0,18033086 | 0,85689283 |
| VCA0770 | 1,15375277 | 16,4725941 | 1,05467187 | 1,6074E-05 | 1,05467187 | 0,18017261 | 0,85701707 |
| tRNA-Leu-2 | -1,1657615 | -4,243009 | -1,1242803 | -4,14E-06 | -1,1242803 | -0,179908 | 0,85722484 |
| VC2017 | 1,08050997 | 0,69061164 | 1,03169919 | 6,739E-07 | 1,03169919 | 0,17979461 | 0,85731382 |
| VCA0594 | 1,06227964 | 22,7262581 | 1,08986263 | 2,2176E-05 | 1,08986263 | 0,17976435 | 0,85733759 |
| VCA0422 | 1,07403966 | 0,69978055 | 1,03280547 | 6,8285E-07 | 1,03280547 | 0,17940938 | 0,85761628 |
| VC0582 | 1,00825927 | -1,1437446 | -1,0235089 | -1,116E-06 | -1,0235089 | -0,1790098 | 0,85792997 |
| VCA0685 | 1,3106434 | 9,80748366 | 1,24162143 | 9,5702E-06 | 1,24162143 | 0,1786536 | 0,85820971 |
| VCA0604 | 1,02629685 | -0,143705 | -1,1595102 | -1,402E-07 | -1,1595102 | -0,1784487 | 0,85837062 |
| VCA0027 | 1,16955182 | 0,43227064 | 1,07102187 | 4,2181E-07 | 1,07102187 | 0,1778552 | 0,8588367 |
| VC2355 | 1,06748912 | 1,53001038 | 1,02351539 | 1,493E-06 | 1,02351539 | 0,17728958 | 0,85928094 |
| VCA0265 | 1,08229177 | 4,99372025 | 1,04741255 | 4,8729E-06 | 1,04741255 | 0,17682146 | 0,85964864 |
| VCA0703 | 1,02253484 | -0,3644725 | -1,058359 | -3,557E-07 | -1,058359 | -0,176071 | 0,86023817 |
| VC0930 | 1,11717174 | 0,40938328 | 1,05120124 | 3,9948E-07 | 1,05120124 | 0,17509088 | 0,86100823 |
| VC1188 | 1,13814715 | 2,53870899 | 1,06106737 | 2,4773E-06 | 1,06106737 | 0,17398715 | 0,86187557 |
| VCA0733 | 1,2101321 | 0,25985577 | 1,08002966 | 2,5357E-07 | 1,08002966 | 0,17318819 | 0,86250352 |
| VCA0275 | 1,28074482 | 0,16387834 | 1,12977191 | 1,5991E-07 | 1,12977191 | 0,17307944 | 0,86258899 |
| VCA0886 | -1,0713401 | -5,0421087 | -1,0591282 | -4,92E-06 | -1,0591282 | -0,1730619 | 0,8626028 |
| VC0772 | -1,0133384 | -0,2288404 | -1,0906933 | -2,233E-07 | -1,0906933 | -0,1725718 | 0,862988 |

| | | | | | | | |
|---|---|---|---|---|---|---|---|
| VCA1016 | 1,12651199 | 0,52460951 | 1,05578997 | 5,1192E-07 | 1,05578997 | 0,17242078 | 0,86310674 |
| VC2357 | 1,02364906 | -5,8853496 | -1,0676444 | -5,743E-06 | -1,0676444 | -0,1724099 | 0,86311533 |
| VCA0455 | 1,10383573 | 1,17207 | 1,04212496 | 1,1437E-06 | 1,04212496 | 0,17228949 | 0,86320996 |
| VC1750 | 1,03479674 | -0,6997767 | -1,0286377 | -6,828E-07 | -1,0286377 | -0,1721522 | 0,86331789 |
| 16Sc | 1,0175049 | 7,34154833 | 1,07211283 | 7,1639E-06 | 1,07211283 | 0,17210345 | 0,86335621 |
| VC2527 | -1,0434552 | -6,5981085 | -1,0399703 | -6,438E-06 | -1,0399703 | -0,1720411 | 0,8634052 |
| VC1341 | -1,0121232 | -2,3348716 | -1,0437905 | -2,278E-06 | -1,0437905 | -0,1714943 | 0,86383512 |
| VC0404 | 1,05357505 | 1,24400693 | 1,02501246 | 1,2139E-06 | 1,02501246 | 0,17128465 | 0,86399996 |
| VC1665 | 1,04753105 | -0,3217486 | -1,0626438 | -3,14E-07 | -1,0626438 | -0,1712171 | 0,86405306 |
| VCA0684 | -1,1454543 | -4,3493307 | -1,2023684 | -4,244E-06 | -1,2023684 | -0,170786 | 0,86439206 |
| VCA1099 | 1,41013427 | 0,12084917 | 1,17431074 | 1,1793E-07 | 1,17431074 | 0,17048411 | 0,86462944 |
| VC1438 | -1,0232228 | -2,9492457 | -1,0576467 | -2,878E-06 | -1,0576467 | -0,170271 | 0,86479699 |
| VCA0977 | 1,16650629 | 0,33270724 | 1,0598267 | 3,2466E-07 | 1,0598267 | 0,17026491 | 0,86480181 |
| VCA0774 | -1,0123428 | -10,570813 | -1,0799308 | -1,032E-05 | -1,0799308 | -0,1702368 | 0,86482391 |
| VC0349 | 1,04560037 | -6,2886728 | -1,0169195 | -6,137E-06 | -1,0169195 | -0,1697253 | 0,86522618 |
| VC0490 | 1,03677005 | 15,571902 | 1,06065995 | 1,5195E-05 | 1,06065995 | 0,16927549 | 0,86557996 |
| VCA1094 | -1,0577141 | -0,2899554 | -1,0859538 | -2,829E-07 | -1,0859538 | -0,1685964 | 0,86611413 |
| VC0937 | 1,03429551 | -0,3655651 | -1,0530438 | -3,567E-07 | -1,0530438 | -0,1683305 | 0,86632329 |
| VC1289 | 1,08115828 | 0,97404044 | 1,01951587 | 9,5048E-07 | 1,01951587 | 0,16804674 | 0,86654651 |
| tRNA-Ile-2 | 1,09600204 | 18,1635081 | 1,0768361 | 1,7724E-05 | 1,0768361 | 0,16800759 | 0,86657732 |
| VC2563 | 1,12583967 | 0,43328886 | 1,04415541 | 4,2281E-07 | 1,04415541 | 0,16756624 | 0,86692454 |
| VC2288 | 1,0701276 | 1,67927099 | 1,03511494 | 1,6386E-06 | 1,03511494 | 0,16717043 | 0,86723596 |
| VC2613 | 1,08126982 | -0,8253198 | -1,0302967 | -8,054E-07 | -1,0302967 | -0,1668847 | 0,86746077 |
| VCA0714 | 1,10458439 | 0,43940376 | 1,04311976 | 4,2877E-07 | 1,04311976 | 0,16679613 | 0,86753047 |
| VC0032 | 1,23555132 | 0,15387651 | 1,12680044 | 1,5015E-07 | 1,12680044 | 0,16589912 | 0,86823635 |
| hemC | 1,05571655 | -0,631877 | -1,0293961 | -6,166E-07 | -1,0293961 | -0,1657078 | 0,86838692 |
| VC0704 | 1,14103314 | 0,41334158 | 1,04528049 | 4,0334E-07 | 1,04528049 | 0,16569 | 0,86840092 |
| VCA0353 | 1,08613652 | 0,41749393 | 1,04478191 | 4,0739E-07 | 1,04478191 | 0,16562106 | 0,86845518 |
| VC0979 | 1,1567512 | 0,65759182 | 1,04531564 | 6,4168E-07 | 1,04531564 | 0,16546367 | 0,86857905 |
| VC1097 | -1,0013439 | 11,7935135 | 1,04888627 | 1,1508E-05 | 1,04888627 | 0,16479087 | 0,86910859 |
| VC1991 | 1,16405913 | 3,47310233 | 1,08674021 | 3,3891E-06 | 1,08674021 | 0,16438255 | 0,86943001 |
| VC0070 | 1,05034831 | 2,98421526 | 1,03599481 | 2,912E-06 | 1,03599481 | 0,16433813 | 0,86946497 |
| hemH (NC_002505 1051232..1052247) | 1,11892174 | 0,75148768 | 1,02418901 | 7,3331E-07 | 1,02418901 | 0,16413899 | 0,86962173 |
| VC1612 | 1,05400338 | -1,1416215 | -1,0588921 | -1,114E-06 | -1,0588921 | -0,1641207 | 0,8696361 |
| VCA0255 | -1,0012279 | -0,1798231 | -1,1051035 | -1,755E-07 | -1,1051035 | -0,1641178 | 0,86963845 |
| VCA0830 | 1,1572868 | 0,36276354 | 1,05065516 | 3,5399E-07 | 1,05065516 | 0,16396062 | 0,86976215 |
| gpsA | 1,12597679 | 0,60201249 | 1,02992459 | 5,8745E-07 | 1,02992459 | 0,16317057 | 0,87038414 |
| gcp | 1,04182948 | 1,46547119 | 1,02389365 | 1,43E-06 | 1,02389365 | 0,16283516 | 0,87064823 |
| VC1337 | 1,05713255 | -0,3636325 | -1,0918124 | -3,548E-07 | -1,0918124 | -0,1627571 | 0,87070968 |
| VCA0375 | 1,15304463 | 0,2389874 | 1,07669709 | 2,3321E-07 | 1,07669709 | 0,16272403 | 0,87073573 |
| VC1171 | 1,0392876 | -0,4896229 | -1,0365391 | -4,778E-07 | -1,0365391 | -0,1623406 | 0,87103768 |
| VC2467 | 1,13603183 | -21,352275 | -1,0268176 | -2,084E-05 | -1,0268176 | -0,1620292 | 0,87128284 |
| recF | -1,0127494 | -3,5597431 | -1,0346244 | -3,474E-06 | -1,0346244 | -0,1619939 | 0,87131065 |
| VCA0271 | -1,0124286 | -2,1183029 | -1,0239346 | -2,067E-06 | -1,0239346 | -0,1617651 | 0,87149084 |
| VC0268 | 1,05021174 | 2,19375945 | 1,03872944 | 2,1407E-06 | 1,03872944 | 0,16134402 | 0,87182247 |

| | | | | | | | |
|---|---|---|---|---|---|---|---|
| VC0947 | 1,04607591 | 4,22588243 | 1,01419815 | 4,1237E-06 | 1,01419815 | 0,16055835 | 0,87244127 |
| VCA0211 | 1,04374134 | -0,3089261 | -1,0570497 | -3,015E-07 | -1,0570497 | -0,1603221 | 0,87262733 |
| VC1550 | 1,04409096 | -0,1637574 | -1,1101694 | -1,598E-07 | -1,1101694 | -0,1601524 | 0,87276106 |
| VC1603 | 1,00482824 | -3,6655205 | -1,043947 | -3,577E-06 | -1,043947 | -0,1601027 | 0,8728002 |
| VCA0751 | 1,14769808 | 1,79608924 | 1,06474321 | 1,7526E-06 | 1,06474321 | 0,15972842 | 0,87309502 |
| VC0379 | 1,13352289 | 0,47839525 | 1,04229779 | 4,6682E-07 | 1,04229779 | 0,15971289 | 0,87310725 |
| VCA0153 | 1,2056028 | 0,2758877 | 1,0635987 | 2,6921E-07 | 1,0635987 | 0,15971263 | 0,87310745 |
| VC2531 | 1,04776417 | 5,61381941 | 1,03645359 | 5,478E-06 | 1,03645359 | 0,15862764 | 0,87396226 |
| VC1976 | 1,12457655 | 0,49609352 | 1,0342524 | 4,8409E-07 | 1,0342524 | 0,15830281 | 0,8742182 |
| VCA0847 | 1,15144283 | 0,26396614 | 1,06522201 | 2,5758E-07 | 1,06522201 | 0,15814277 | 0,8743443 |
| VCA0865 | 1,19815597 | 0,23906542 | 1,07223109 | 2,3328E-07 | 1,07223109 | 0,15811124 | 0,87436915 |
| ribB | -1,0166907 | -1,0879535 | -1,0463672 | -1,062E-06 | -1,0463672 | -0,1579476 | 0,87449811 |
| VC0142 | 1,02801574 | -3,8237036 | -1,0489192 | -3,731E-06 | -1,0489192 | -0,1579184 | 0,87452113 |
| VCA0772 | 1,16726687 | 0,24720704 | 1,0695144 | 2,4123E-07 | 1,0695144 | 0,15783197 | 0,87458921 |
| VC0392 | 1,13369757 | 0,64547179 | 1,03985427 | 6,2986E-07 | 1,03985427 | 0,15775021 | 0,87465364 |
| VC1360 | 1,02750506 | -0,2306844 | -1,0742511 | -2,251E-07 | -1,0742511 | -0,1573951 | 0,87493349 |
| VC2339 | 1,21176953 | 0,22445807 | 1,07634814 | 2,1903E-07 | 1,07634814 | 0,15735409 | 0,8749658 |
| VC0861 | 1,05315494 | -0,272505 | -1,0623526 | -2,659E-07 | -1,0623526 | -0,1572153 | 0,8750752 |
| VC2395 | 1,03052909 | -0,4668481 | -1,035826 | -4,556E-07 | -1,035826 | -0,156993 | 0,87525036 |
| VC0770 | -1,1613843 | -13,66746 | -1,0890284 | -1,334E-05 | -1,0890284 | -0,1565309 | 0,87561461 |
| VCA0228 | 1,15132865 | 0,33306191 | 1,0498169 | 3,25E-07 | 1,0498169 | 0,15583155 | 0,87616581 |
| recC | 1,11726624 | 0,45665346 | 1,03564983 | 4,4561E-07 | 1,03564983 | 0,15489377 | 0,87690507 |
| VC1784 | -1,0328942 | -2,6859476 | -1,1038285 | -2,621E-06 | -1,1038285 | -0,154614 | 0,87712567 |
| folE | 1,07001789 | 0,62842742 | 1,02679271 | 6,1322E-07 | 1,02679271 | 0,15451907 | 0,87720048 |
| tRNA-Leu-12 | 1,05593562 | 5,71983358 | 1,08064543 | 5,5815E-06 | 1,08064543 | 0,15390868 | 0,87768175 |
| VC2064 | 1,08199927 | 7,68181197 | 1,0157057 | 7,496E-06 | 1,0157057 | 0,15370236 | 0,87784443 |
| VC0301 | 1,08063482 | -0,1318567 | -1,1268078 | -1,287E-07 | -1,1268078 | -0,1535751 | 0,87794477 |
| VC0943 | 1,06358871 | 0,75926605 | 1,03830312 | 7,409E-07 | 1,03830312 | 0,15351268 | 0,877994 |
| VC1779 | 1,02091306 | -8,1276844 | -1,1065145 | -7,931E-06 | -1,1065145 | -0,1534826 | 0,87801773 |
| VC0628 | 1,10745388 | 0,74081735 | 1,03788183 | 7,229E-07 | 1,03788183 | 0,15252275 | 0,87877465 |
| VC0721 | 1,09486676 | 0,39739005 | 1,03976902 | 3,8778E-07 | 1,03976902 | 0,15245906 | 0,87882488 |
| VC1359 | 1,02005914 | -0,312944 | -1,0505704 | -3,054E-07 | -1,0505704 | -0,1521619 | 0,87905923 |
| tRNA-Leu-3 | -1,0879455 | -2,2756567 | -1,0793344 | -2,221E-06 | -1,0793344 | -0,1518779 | 0,87928325 |
| VC0723 | 1,05011014 | -0,9339006 | -1,0352452 | -9,113E-07 | -1,0352452 | -0,1517467 | 0,87938675 |
| rpmA | -1,0622017 | -51,077168 | -1,0359105 | -4,984E-05 | -1,0359105 | -0,1516713 | 0,87944621 |
| VCA0640 | 1,51374775 | 0,10867267 | 1,14952694 | 1,0604E-07 | 1,14952694 | 0,15059454 | 0,88029557 |
| VC1284 | 1,18006653 | 0,32046994 | 1,04795877 | 3,1272E-07 | 1,04795877 | 0,15004762 | 0,88072705 |
| VCA0739 | 1,01306465 | -0,4759267 | -1,0356984 | -4,644E-07 | -1,0356984 | -0,1497911 | 0,88092944 |
| VC0051 | 1,09097435 | 0,444017 | 1,03816926 | 4,3328E-07 | 1,03816926 | 0,14896492 | 0,88158132 |
| VCA0993 | 1,04773995 | -0,1619686 | -1,0956417 | -1,581E-07 | -1,0956417 | -0,1489161 | 0,88161986 |
| VC1161 | 1,01875292 | -0,4800816 | -1,0312328 | -4,685E-07 | -1,0312328 | -0,148815 | 0,88169959 |
| VC1663 | 1,01179698 | -5,2017281 | -1,0171392 | -5,076E-06 | -1,0171392 | -0,1483394 | 0,88207492 |
| VCA0619 | 1,05414896 | -0,8387715 | -1,034098 | -8,185E-07 | -1,034098 | -0,148121 | 0,88224725 |
| VC2223 | 1,06839539 | 0,57719821 | 1,02558064 | 5,6323E-07 | 1,02558064 | 0,14787982 | 0,88243762 |
| VC0680 | 1,00261342 | -0,6506599 | -1,0334093 | -6,349E-07 | -1,0334093 | -0,1478094 | 0,88249317 |

| | | | | | | | |
|---|---|---|---|---|---|---|---|
| VCA0274 | -1,0362596 | -2,8107697 | -1,0814662 | -2,743E-06 | -1,0814662 | -0,1477068 | 0,88257415 |
| VCA0562 | 1,05370363 | 4,29946809 | 1,02574086 | 4,1955E-06 | 1,02574086 | 0,14758839 | 0,88266762 |
| recA | 1,04885668 | 5,23117596 | 1,01435581 | 5,1046E-06 | 1,01435581 | 0,14754094 | 0,88270507 |
| VC2646 | 1,03646949 | 7,60203649 | 1,11755093 | 7,4181E-06 | 1,11755093 | 0,14710409 | 0,88304987 |
| VCA0791 | -1,1561883 | -2,8887399 | -1,0757242 | -2,819E-06 | -1,0757242 | -0,1466078 | 0,8834416 |
| VC1203 | 1,05430062 | -0,7902801 | -1,0907981 | -7,712E-07 | -1,0907981 | -0,1465773 | 0,88346567 |
| VC0501 | 1,15023955 | 0,58211302 | 1,04569767 | 5,6803E-07 | 1,04569767 | 0,14615268 | 0,88380087 |
| VC2603 | 1,13080321 | 2,86809553 | 1,06344734 | 2,7987E-06 | 1,06344734 | 0,14553786 | 0,88428623 |
| VC0901 | 1,07369157 | 0,56011623 | 1,0254504 | 5,4657E-07 | 1,0254504 | 0,14530847 | 0,88446734 |
| VC0858 | 1,13306495 | 0,2234098 | 1,06470237 | 2,1801E-07 | 1,06470237 | 0,14492519 | 0,88476995 |
| VCA0663 | 1,09968338 | 1,07889285 | 1,03393481 | 1,0528E-06 | 1,03393481 | 0,1446733 | 0,88496883 |
| VC2702 | 1,02278999 | -2,8639806 | -1,0499577 | -2,795E-06 | -1,0499577 | -0,1446703 | 0,88497121 |
| VCA0037 | 1,06789761 | 3,17707684 | 1,04860895 | 3,1002E-06 | 1,04860895 | 0,14417081 | 0,8853656 |
| VC2256 | -1,0081543 | -1,4224219 | -1,0269227 | -1,388E-06 | -1,0269227 | -0,144166 | 0,88536942 |
| VC1590 | 1,06442722 | 0,83172067 | 1,08863684 | 8,116E-07 | 1,08863684 | 0,14406559 | 0,88544869 |
| VC0522 | 1,04630868 | 2,89243531 | 1,02854533 | 2,8225E-06 | 1,02854533 | 0,14373101 | 0,88571289 |
| VCA1067 | 1,11808805 | 0,63512538 | 1,04342207 | 6,1976E-07 | 1,04342207 | 0,14345802 | 0,88592847 |
| VC2525 | -1,0386655 | -6,5276626 | -1,034981 | -6,37E-06 | -1,034981 | -0,1431189 | 0,8861963 |
| VC0023 | 1,02336458 | -2,5298171 | -1,048692 | -2,469E-06 | -1,048692 | -0,1423862 | 0,88677499 |
| VCA0526 | -1,0182952 | -0,403894 | -1,0512407 | -3,941E-07 | -1,0512407 | -0,1422497 | 0,88688274 |
| VCA0903 | 1,15737392 | 0,74206305 | 1,05653893 | 7,2411E-07 | 1,05653893 | 0,14214135 | 0,88696836 |
| VC0929 | 1,1093423 | 0,28338432 | 1,05165866 | 2,7653E-07 | 1,05165866 | 0,14202888 | 0,8870572 |
| VCA0825 | 1,22311317 | 0,20158769 | 1,06899309 | 1,9671E-07 | 1,06899309 | 0,14200922 | 0,88707273 |
| VC0510 | 1,01637235 | -0,3247331 | -1,0419529 | -3,169E-07 | -1,0419529 | -0,1414764 | 0,88749364 |
| VCA1098 | 1,2585972 | 0,13705271 | 1,10197589 | 1,3374E-07 | 1,10197589 | 0,14123412 | 0,887685 |
| VC2444 | 1,02530736 | -0,4642717 | -1,0333241 | -4,53E-07 | -1,0333241 | -0,1406779 | 0,88812443 |
| VC1074 | 1,14242194 | 0,3197514 | 1,05501685 | 3,1202E-07 | 1,05501685 | 0,1406032 | 0,88818343 |
| tRNA-Lys-1 | 1,01560439 | -0,5016561 | -1,0330113 | -4,895E-07 | -1,0330113 | -0,1403307 | 0,88839875 |
| nudE | 1,08781796 | 0,75126439 | 1,02454557 | 7,3309E-07 | 1,02454557 | 0,14000175 | 0,88865861 |
| VCA0233 | 1,05394473 | -0,2098073 | -1,0636186 | -2,047E-07 | -1,0636186 | -0,1392993 | 0,88921363 |
| VCA0322 | 1,01089999 | -2,1399363 | -1,0388969 | -2,088E-06 | -1,0388969 | -0,1392778 | 0,88923061 |
| VCA0276 | -1,0272639 | -3,680673 | -1,0479087 | -3,592E-06 | -1,0479087 | -0,1384174 | 0,88991055 |
| wecC | 1,16043668 | 0,17296676 | 1,07586036 | 1,6878E-07 | 1,07586036 | 0,13770535 | 0,89047329 |
| VC1191 | 1,11567801 | 0,72774724 | 1,04453217 | 7,1014E-07 | 1,04453217 | 0,13747673 | 0,890654 |
| VC0525 | 1,08470564 | 0,45264731 | 1,02822171 | 4,417E-07 | 1,02822171 | 0,13746053 | 0,89066679 |
| queA | 1,10492771 | 0,62097571 | 1,04196239 | 6,0595E-07 | 1,04196239 | 0,13715885 | 0,89090524 |
| VCA0406 | 1,12120025 | 1,424669 | 1,04437383 | 1,3902E-06 | 1,04437383 | 0,13709965 | 0,89095204 |
| VC2045 | 1,17089133 | 20,8494206 | 1,03562723 | 2,0345E-05 | 1,03562723 | 0,1368782 | 0,89112707 |
| VCA0704 | 1,13558748 | 0,25637596 | 1,04983373 | 2,5017E-07 | 1,04983373 | 0,13674227 | 0,89123452 |
| VC1778 | 1,05267791 | -3,23578 | -1,0706025 | -3,158E-06 | -1,0706025 | -0,1367163 | 0,89125504 |
| VC2712 | -1,0020152 | -0,4558764 | -1,0502541 | -4,448E-07 | -1,0502541 | -0,1365573 | 0,89138073 |
| VC1392 | 1,07306208 | 0,3713014 | 1,03366869 | 3,6232E-07 | 1,03366869 | 0,13579986 | 0,89197951 |
| VC0024 | 1,02490453 | -0,4531606 | -1,027368 | -4,422E-07 | -1,027368 | -0,1354707 | 0,89223971 |
| fieF | 1,09748371 | 0,58412514 | 1,02089069 | 5,6999E-07 | 1,02089069 | 0,13459363 | 0,89293319 |
| VC0224 | 1,08715916 | 0,54273767 | 1,02230358 | 5,2961E-07 | 1,02230358 | 0,13400621 | 0,89339766 |

| | | | | | | | |
|---|---|---|---|---|---|---|---|
| VCA0345 | 1,05827959 | 4,68171772 | 1,04918649 | 4,5685E-06 | 1,04918649 | 0,13387151 | 0,89350418 |
| tRNA-Thr-3 | -1,0924213 | -8,1531847 | -1,0467388 | -7,956E-06 | -1,0467388 | -0,1338491 | 0,8935219 |
| VC2043 | 1,11093132 | 0,32623116 | 1,03701923 | 3,1834E-07 | 1,03701923 | 0,13336485 | 0,89390485 |
| VCA0209 | 1,10011863 | 0,34152823 | 1,03519042 | 3,3327E-07 | 1,03519042 | 0,13310236 | 0,89411243 |
| hisS | 1,00906446 | -1,7467667 | -1,0209191 | -1,705E-06 | -1,0209191 | -0,1326541 | 0,89446694 |
| VCA0707 | 1,04686685 | -0,0768725 | -1,1646395 | -7,501E-08 | -1,1646395 | -0,1324404 | 0,89463599 |
| VC0182 | 1,03622389 | 1,70655422 | 1,0306538 | 1,6653E-06 | 1,0306538 | 0,13235777 | 0,89470132 |
| VC2647 | 1,1597242 | 28,1049565 | 1,05128395 | 2,7425E-05 | 1,05128395 | 0,13226625 | 0,89477371 |
| VC0088 | 1,07508108 | 0,58769592 | 1,01977805 | 5,7348E-07 | 1,01977805 | 0,13139634 | 0,89546179 |
| tRNA-Gln-1 | -1,1248187 | -12,12429 | -1,0840453 | -1,183E-05 | -1,0840453 | -0,1308885 | 0,89586351 |
| VCA0350 | 1,14948373 | 1,21103483 | 1,05165025 | 1,1817E-06 | 1,05165025 | 0,13085537 | 0,89588972 |
| VCA0408 | 1,1393611 | 0,72576203 | 1,04069524 | 7,082E-07 | 1,04069524 | 0,13065007 | 0,89605213 |
| tRNA-Ala-2 | -1,0503112 | -2,1392915 | -1,0436371 | -2,088E-06 | -1,0436371 | -0,1304501 | 0,89621038 |
| VCA0942 | 1,08650146 | 0,78585883 | 1,02756231 | 7,6685E-07 | 1,02756231 | 0,12964156 | 0,89685003 |
| VC1687 | 1,10971457 | 1,5841644 | 1,05666937 | 1,5458E-06 | 1,05666937 | 0,12875557 | 0,89755107 |
| VC2548 | 1,06187671 | 0,67074802 | 1,01644183 | 6,5452E-07 | 1,01644183 | 0,12809474 | 0,89807401 |
| VCA0534 | 1,07543678 | 0,32582276 | 1,03412814 | 3,1794E-07 | 1,03412814 | 0,12806208 | 0,89809985 |
| VC2344 | 1,03036095 | -0,8774127 | -1,0192405 | -8,562E-07 | -1,0192405 | -0,1278615 | 0,89825862 |
| VC0780 | 1,03423623 | -0,1723702 | -1,0651582 | -1,682E-07 | -1,0651582 | -0,1277321 | 0,89836101 |
| VCA0188 | 1,3056707 | 0,17801518 | 1,06252673 | 1,7371E-07 | 1,06252673 | 0,12723976 | 0,89875064 |
| VC0954 | 1,02336883 | -0,8430202 | -1,0184668 | -8,226E-07 | -1,0184668 | -0,1271561 | 0,89881683 |
| VC1554 | 1,02891792 | 1,00394793 | 1,01740608 | 9,7966E-07 | 1,01740608 | 0,12713531 | 0,89883331 |
| VC1602 | -1,0280696 | -4,5907765 | -1,0386979 | -4,48E-06 | -1,0386979 | -0,1271027 | 0,89885908 |
| VC0400 | 1,04454491 | -0,6509559 | -1,0166597 | -6,352E-07 | -1,0166597 | -0,1270171 | 0,8989269 |
| VCA1111 | 1,17752187 | 0,17770991 | 1,0615992 | 1,7341E-07 | 1,0615992 | 0,12621254 | 0,89956369 |
| VC1340 | 1,00286419 | -0,2156381 | -1,0503759 | -2,104E-07 | -1,0503759 | -0,1260721 | 0,89967488 |
| VC0726 | 1,02153205 | -0,6997428 | -1,0199388 | -6,828E-07 | -1,0199388 | -0,1259965 | 0,89973469 |
| VC1332 | 1,0340954 | -0,1691584 | -1,0640313 | -1,651E-07 | -1,0640313 | -0,1254717 | 0,9001501 |
| VCA0876 | 1,11122192 | 0,41526844 | 1,02547508 | 4,0522E-07 | 1,02547508 | 0,12517652 | 0,90038381 |
| VCA1056 | 1,01186322 | -0,1822682 | -1,0585215 | -1,779E-07 | -1,0585215 | -0,12468 | 0,9007769 |
| VC1989 | 1,0406486 | -0,9749483 | -1,0295641 | -9,514E-07 | -1,0295641 | -0,1245036 | 0,90091654 |
| tRNA-Val-2 | -1,0540913 | -2,2767754 | -1,0541646 | -2,222E-06 | -1,0541646 | -0,1243278 | 0,90105577 |
| VC1105 | 1,09428089 | 0,29770829 | 1,03511331 | 2,9051E-07 | 1,03511331 | 0,12413651 | 0,9012072 |
| tpx | 1,03475627 | 4,14299 | 1,01382196 | 4,0428E-06 | 1,01382196 | 0,12393471 | 0,90136697 |
| VCA0543 | 1,06873097 | 1,39773176 | 1,02404162 | 1,3639E-06 | 1,02404162 | 0,12390326 | 0,90139188 |
| zntR | -1,0343852 | -1,4617591 | -1,0314075 | -1,426E-06 | -1,0314075 | -0,1232964 | 0,90187238 |
| VC2342 | 1,16156245 | -8,272616 | -1,0156158 | -8,072E-06 | -1,0156158 | -0,1227424 | 0,9023111 |
| VC1261 | 1,14900996 | 0,22413776 | 1,04562818 | 2,1872E-07 | 1,04562818 | 0,12246795 | 0,90252843 |
| VC0009 | -1,0023975 | -0,7161772 | -1,0382581 | -6,989E-07 | -1,0382581 | -0,1221913 | 0,90274755 |
| VCA0802 | 1,04779063 | -0,8685744 | -1,0179446 | -8,476E-07 | -1,0179446 | -0,1218163 | 0,90304453 |
| rpmB | 1,11324149 | -37,480701 | -1,0384027 | -3,657E-05 | -1,0384027 | -0,1216783 | 0,90315381 |
| VCA0023 | 1,04778102 | -0,2415295 | -1,0414666 | -2,357E-07 | -1,0414666 | -0,1213179 | 0,90343926 |
| VC0005 | -1,0592577 | -4,0605791 | -1,0385253 | -3,962E-06 | -1,0385253 | -0,1210524 | 0,90364958 |
| VC2030 | 1,05056488 | 2,70077742 | 1,02932627 | 2,6354E-06 | 1,02932627 | 0,12076937 | 0,90387372 |
| VCA0040 | 1,03930253 | -0,5030127 | -1,0194843 | -4,908E-07 | -1,0194843 | -0,120664 | 0,90395717 |

| | | | | | | | |
|---|---|---|---|---|---|---|---|
| VC0258 | 1,01215204 | 13,753381 | 1,0373832 | 1,3421E-05 | 1,0373832 | 0,12060749 | 0,90400194 |
| VC1895 | -1,0154302 | -1,132255 | -1,0105619 | -1,105E-06 | -1,0105619 | -0,1203639 | 0,90419487 |
| 23Sf | 1,03275144 | -0,2732135 | -1,0359064 | -2,666E-07 | -1,0359064 | -0,120232 | 0,90429938 |
| VCA0256 | 1,14023672 | 0,21612332 | 1,0454859 | 2,1089E-07 | 1,0454859 | 0,12007501 | 0,90442373 |
| VC1493 | 1,06307248 | -0,466618 | -1,020769 | -4,553E-07 | -1,020769 | -0,1199488 | 0,90452373 |
| VC2678 | 1,05455414 | -0,4141968 | -1,0229242 | -4,042E-07 | -1,0229242 | -0,1186657 | 0,90554025 |
| VCA0778 | 1,01242886 | -0,2633483 | -1,0361292 | -2,57E-07 | -1,0361292 | -0,1184006 | 0,90575028 |
| VCA0440 | 1,10348531 | 0,61413576 | 1,03180462 | 5,9928E-07 | 1,03180462 | 0,11831296 | 0,90581969 |
| VC2280 | 1,02231923 | -0,4503723 | -1,0208335 | -4,395E-07 | -1,0208335 | -0,1180232 | 0,9060493 |
| VC1467 | 1,01250073 | -0,2846333 | -1,0329817 | -2,777E-07 | -1,0329817 | -0,1176996 | 0,90630571 |
| cca | 1,03567718 | -0,2896809 | -1,0322183 | -2,827E-07 | -1,0322183 | -0,1173785 | 0,90656016 |
| VCA0956 | 1,07653038 | 0,46310924 | 1,01989225 | 4,5191E-07 | 1,01989225 | 0,11697288 | 0,90688154 |
| xerC | 1,01272324 | 0,84016276 | 1,01090399 | 8,1984E-07 | 1,01090399 | 0,11691113 | 0,90693048 |
| tRNA-Met-1 | -1,1141261 | -2,5683658 | -1,0724586 | -2,506E-06 | -1,0724586 | -0,1168273 | 0,9069969 |
| rpsI | 1,19078876 | 8,05564296 | 1,01512474 | 7,8608E-06 | 1,01512474 | 0,11681893 | 0,90700355 |
| VC1707 | 1,05131493 | -0,940854 | -1,0374645 | -9,181E-07 | -1,0374645 | -0,1166896 | 0,90710604 |
| VC0298 | 1,08577105 | -0,1634089 | -1,0568139 | -1,595E-07 | -1,0568139 | -0,1163668 | 0,90736187 |
| VC0465 | 1,2088853 | 0,20250607 | 1,04555125 | 1,9761E-07 | 1,04555125 | 0,11631228 | 0,90740506 |
| VC1355 | 1,03616882 | -0,3483347 | -1,0260126 | -3,399E-07 | -1,0260126 | -0,1158332 | 0,90778478 |
| VC2053 | -1,0630359 | -9,3281716 | -1,0444171 | -9,103E-06 | -1,0444171 | -0,1155916 | 0,90797623 |
| VC0254 | -1,0230405 | 8,30171878 | 1,03169543 | 8,1009E-06 | 1,03169543 | 0,11553362 | 0,90802218 |
| VC2718 | 1,12699069 | 0,24737892 | 1,03599857 | 2,4139E-07 | 1,03599857 | 0,11455059 | 0,90880135 |
| VC1245 | 1,03053708 | -0,3626353 | -1,024286 | -3,539E-07 | -1,024286 | -0,114246 | 0,90904276 |
| VC2445 | 1,07750011 | -0,2476522 | -1,0350703 | -2,417E-07 | -1,0350703 | -0,1131522 | 0,90990988 |
| gyrB | 1,0649752 | 2,57896624 | 1,02674184 | 2,5166E-06 | 1,02674184 | 0,11311608 | 0,90993853 |
| VC0736 | 1,05488223 | -0,5378393 | -1,0341688 | -5,248E-07 | -1,0341688 | -0,1125141 | 0,91041581 |
| VC2703 | 1,11040104 | 0,30176857 | 1,02822834 | 2,9447E-07 | 1,02822834 | 0,11224944 | 0,91062563 |
| 16Sa | 1,02317603 | -0,2545822 | -1,0334968 | -2,484E-07 | -1,0334968 | -0,1121646 | 0,91069286 |
| VCA0269 | 1,03733463 | -0,1666908 | -1,0515836 | -1,627E-07 | -1,0515836 | -0,1121317 | 0,91071899 |
| VC0931 | 1,13636448 | 0,16481405 | 1,05167704 | 1,6083E-07 | 1,05167704 | 0,11159706 | 0,91114291 |
| VCA0803 | 1,02436852 | -0,1742367 | -1,0486774 | -1,7E-07 | -1,0486774 | -0,1114444 | 0,91126398 |
| VC1815 | 1,19196957 | 0,18077023 | 1,04686797 | 1,764E-07 | 1,04686797 | 0,11143409 | 0,91127213 |
| VC2307 | 1,13265822 | 0,22781483 | 1,03672481 | 2,223E-07 | 1,03672481 | 0,11101114 | 0,91160751 |
| VCA0914 | 1,17350418 | 0,2366877 | 1,03492533 | 2,3096E-07 | 1,03492533 | 0,11039412 | 0,91209682 |
| VCA0240 | 1,04531011 | -0,3705911 | -1,0220871 | -3,616E-07 | -1,0220871 | -0,1101998 | 0,91225091 |
| VCA0068 | 1,09183417 | -0,2270607 | -1,0361693 | -2,216E-07 | -1,0361693 | -0,1100008 | 0,91240871 |
| VC0984 | -1,0095936 | 1,76183766 | 1,01309256 | 1,7192E-06 | 1,01309256 | 0,10991907 | 0,91247357 |
| VC1276 | 1,05333945 | -0,450888 | -1,0389618 | -4,4E-07 | -1,0389618 | -0,1099003 | 0,91248846 |
| VC0505 | 1,03766182 | -0,5672368 | -1,0270325 | -5,535E-07 | -1,0270325 | -0,1097432 | 0,91261307 |
| VCA0794 | 1,02918476 | -0,3230935 | -1,0305195 | -3,153E-07 | -1,0305195 | -0,1096611 | 0,91267812 |
| VC0450 | 1,12418129 | 0,83986571 | 1,02827131 | 8,1955E-07 | 1,02827131 | 0,10951245 | 0,91279605 |
| VC1774 | 1,13671391 | 1,74945731 | 1,06436989 | 1,7071E-06 | 1,06436989 | 0,10948426 | 0,91281841 |
| VC0762 | 1,00172256 | -2,1222227 | -1,0228243 | -2,071E-06 | -1,0228243 | -0,1093291 | 0,91294148 |
| VC0884 | 1,03263532 | 0,83564934 | 1,02377047 | 8,1543E-07 | 1,02377047 | 0,10859351 | 0,91352491 |
| VCA0974 | 1,04791665 | 1,37229544 | 1,05714995 | 1,3391E-06 | 1,05714995 | 0,10851604 | 0,91358636 |

| | | | | | | | |
|---|---|---|---|---|---|---|---|
| VC1659 | 1,18422369 | 0,21338438 | 1,03745663 | 2,0822E-07 | 1,03745663 | 0,10848341 | 0,91361224 |
| VCA0963 | 1,0429031 | -0,3052156 | -1,025959 | -2,978E-07 | -1,025959 | -0,1083166 | 0,91374457 |
| VC2705 | 1,02011423 | -0,0984943 | -1,0820496 | -9,611E-08 | -1,0820496 | -0,1079094 | 0,91406759 |
| VCA1091 | 1,01412276 | -0,1860781 | -1,0424964 | -1,816E-07 | -1,0424964 | -0,1077716 | 0,91417685 |
| VC2217 | 1,21120521 | 0,19446472 | 1,04037466 | 1,8976E-07 | 1,04037466 | 0,10744385 | 0,91443687 |
| VC0623 | 1,04420898 | -0,3083903 | -1,025267 | -3,009E-07 | -1,025267 | -0,1074359 | 0,91444317 |
| VCA0310 | 1,03501007 | 0,66488764 | 1,01464293 | 6,488E-07 | 1,01464293 | 0,10739699 | 0,91447404 |
| VC1913 | 1,01187596 | 0,90087038 | 1,01250873 | 8,7908E-07 | 1,01250873 | 0,10729197 | 0,91455736 |
| VCA0277 | 1,06458787 | 10,9708988 | 1,03437864 | 1,0705E-05 | 1,03437864 | 0,10712743 | 0,9146879 |
| VC0775 | 1,14778941 | 0,15974031 | 1,04906142 | 1,5588E-07 | 1,04906142 | 0,10711769 | 0,91469562 |
| 5Sd | 1,00322127 | 204,481457 | 1,04208871 | 0,00019953 | 1,04208871 | 0,10634982 | 0,91530481 |
| VC1432 | 1,05611117 | 1,00205836 | 1,04573562 | 9,7782E-07 | 1,04573562 | 0,10611868 | 0,91548819 |
| VCA0824 | 1,03153067 | -0,1493023 | -1,0514551 | -1,457E-07 | -1,0514551 | -0,1059932 | 0,91558778 |
| VCA1038 | 1,10813067 | 0,28845725 | 1,02620439 | 2,8148E-07 | 1,02620439 | 0,1057911 | 0,9157481 |
| VCA0495 | -1,0398914 | -3,9438449 | -1,0279457 | -3,848E-06 | -1,0279457 | -0,1054008 | 0,91605777 |
| VCA0882 | 1,11119353 | 0,28132789 | 1,02645029 | 2,7452E-07 | 1,02645029 | 0,10495825 | 0,91640894 |
| murE | 1,05485765 | 0,73257448 | 1,01458999 | 7,1485E-07 | 1,01458999 | 0,10495697 | 0,91640996 |
| VCA0634 | 1,16759913 | 0,2024371 | 1,03664523 | 1,9754E-07 | 1,03664523 | 0,10453408 | 0,91674553 |
| VC2697 | 1,0798158 | -0,6456876 | -1,0310204 | -6,301E-07 | -1,0310204 | -0,1043152 | 0,91691918 |
| VCA0917 | 1,0247655 | -10,265099 | -1,0416801 | -1,002E-05 | -1,0416801 | -0,1042428 | 0,91697666 |
| VC1187 | 1,04491623 | -0,3618565 | -1,0257064 | -3,531E-07 | -1,0257064 | -0,1026669 | 0,91822732 |
| dnaE | 1,0591382 | 0,5967596 | 1,01679317 | 5,8232E-07 | 1,01679317 | 0,10253323 | 0,91833344 |
| VCA0351 | 1,10890992 | 2,24708771 | 1,0374059 | 2,1927E-06 | 1,0374059 | 0,10226339 | 0,91854762 |
| VC0942 | 1,05379259 | 1,71051161 | 1,02122844 | 1,6691E-06 | 1,02122844 | 0,10192609 | 0,91881534 |
| VCA0072 | 1,20784805 | 0,10736571 | 1,06659747 | 1,0477E-07 | 1,06659747 | 0,10188135 | 0,91885086 |
| VCA0780 | 1,08079411 | -0,1869399 | -1,0372463 | -1,824E-07 | -1,0372463 | -0,1012587 | 0,91934511 |
| VC0860 | 1,13600688 | 0,1474313 | 1,0474245 | 1,4386E-07 | 1,0474245 | 0,10121702 | 0,91937819 |
| VC0300 | 1,02824353 | -0,1115749 | -1,0630296 | -1,089E-07 | -1,0630296 | -0,1011262 | 0,91945026 |
| VC2302 | -1,0550728 | -7,605795 | -1,0429183 | -7,422E-06 | -1,0429183 | -0,1008559 | 0,91966487 |
| VC1610 | 1,09779581 | 0,37864219 | 1,01785698 | 3,6948E-07 | 1,01785698 | 0,10026248 | 0,92013594 |
| VC0653 | 1,07475539 | 3,120252 | 1,03589045 | 3,0448E-06 | 1,03589045 | 0,10021094 | 0,92017687 |
| VC2248 | 1,02170184 | 0,63848672 | 1,01049853 | 6,2304E-07 | 1,01049853 | 0,10001424 | 0,92033303 |
| VC1472 | -1,0145385 | -0,360467 | -1,0348605 | -3,517E-07 | -1,0348605 | -0,0995968 | 0,92066443 |
| VC1834 | 1,1182599 | 10,8127992 | 1,04160735 | 1,0551E-05 | 1,04160735 | 0,0995642 | 0,92069032 |
| cobU | 1,09893352 | 0,39613859 | 1,03456169 | 3,8656E-07 | 1,03456169 | 0,09920714 | 0,92097381 |
| VC1539 | -1,0246322 | -4,1529465 | -1,039906 | -4,052E-06 | -1,039906 | -0,0987582 | 0,92133024 |
| VCA0617 | 1,10820378 | 0,28606928 | 1,02277254 | 2,7915E-07 | 1,02277254 | 0,09829493 | 0,9216981 |
| VC2474 | 1,08039255 | 0,45749603 | 1,01410824 | 4,4643E-07 | 1,01410824 | 0,09805195 | 0,92189105 |
| VCA1013 | -1,0106775 | -1,2210451 | -1,0403828 | -1,192E-06 | -1,0403828 | -0,0980262 | 0,92191149 |
| VC0209 | 1,02995016 | 0,90956419 | 1,01409439 | 8,8756E-07 | 1,01409439 | 0,09795159 | 0,92197073 |
| VC0383 | 1,03728742 | -0,247501 | -1,0261814 | -2,415E-07 | -1,0261814 | -0,0979508 | 0,92197133 |
| VC2275 | 1,11486414 | 1,30883556 | 1,04278313 | 1,2772E-06 | 1,04278313 | 0,09768622 | 0,92218147 |
| VCA0979 | 1,08626196 | -0,221985 | -1,028787 | -2,166E-07 | -1,028787 | -0,0972085 | 0,9225608 |
| VC0507 | 1,00456506 | -0,7849666 | -1,0184668 | -7,66E-07 | -1,0184668 | -0,0967966 | 0,92288791 |
| VC2253 | 1,05301042 | 1,13832787 | 1,01610912 | 1,1108E-06 | 1,01610912 | 0,09642358 | 0,92318417 |

| | | | | | | | |
|---|---|---|---|---|---|---|---|
| VCA1055 | 1,07475329 | 0,31429443 | 1,01950921 | 3,0669E-07 | 1,01950921 | 0,09543975 | 0,92396555 |
| VC1167 | 1,01448999 | -0,3706286 | -1,0190066 | -3,617E-07 | -1,0190066 | -0,095182 | 0,92417028 |
| VC1501 | 1,02717329 | -0,2872148 | -1,0210685 | -2,803E-07 | -1,0210685 | -0,0947749 | 0,92449363 |
| VC1540 | 1,0962688 | 0,29176482 | 1,03065314 | 2,8471E-07 | 1,03065314 | 0,09459395 | 0,92463735 |
| anmK | 1,06958267 | -0,6858465 | -1,0250017 | -6,693E-07 | -1,0250017 | -0,0945094 | 0,92470453 |
| VCA0801 | 1,03602205 | 1,40101901 | 1,05143943 | 1,3671E-06 | 1,05143943 | 0,09440635 | 0,92478637 |
| VC0402 | 1,04588904 | 0,80168014 | 1,01157684 | 7,8229E-07 | 1,01157684 | 0,09417734 | 0,92496829 |
| VCA0223 | 1,13530955 | 0,10770455 | 1,05628111 | 1,051E-07 | 1,05628111 | 0,0940412 | 0,92507643 |
| VC1058 | 1,04826099 | 0,56416087 | 1,02767933 | 5,5051E-07 | 1,02767933 | 0,09381496 | 0,92525615 |
| VC1458 | 1,06314153 | -0,1435232 | -1,0414294 | -1,401E-07 | -1,0414294 | -0,093478 | 0,92552384 |
| VC1307 | 1,07140068 | 0,92996789 | 1,03011052 | 9,0747E-07 | 1,03011052 | 0,09341683 | 0,92557242 |
| VCA0410 | 1,05582547 | -0,9281153 | -1,0294483 | -9,057E-07 | -1,0294483 | -0,0932084 | 0,92573798 |
| VC0425 | 1,04919112 | -0,3587419 | -1,0256412 | -3,501E-07 | -1,0256412 | -0,0930666 | 0,92585066 |
| VC0773 | 1,33178706 | 0,05673155 | 1,10705655 | 5,5359E-08 | 1,10705655 | 0,09299115 | 0,9259106 |
| VCA1110 | 1,31023794 | 0,04766905 | 1,12661058 | 4,6516E-08 | 1,12661058 | 0,092272 | 0,92648193 |
| VC1773 | 1,10384816 | 1,56112162 | 1,04864286 | 1,5234E-06 | 1,04864286 | 0,0922292 | 0,92651594 |
| VC0932 | 1,04188997 | -0,1234966 | -1,0469736 | -1,205E-07 | -1,0469736 | -0,092206 | 0,92653435 |
| VC2506 | 1,0984613 | 0,24026158 | 1,03832087 | 2,3445E-07 | 1,03832087 | 0,09203381 | 0,92667118 |
| VCA0306 | 1,06215996 | -0,2429679 | -1,023222 | -2,371E-07 | -1,023222 | -0,0914672 | 0,9271214 |
| VC1932 | 1,08488663 | 0,21920021 | 1,02572323 | 2,139E-07 | 1,02572323 | 0,09138094 | 0,92718991 |
| VC1630 | -1,0487968 | -2,1746281 | -1,0335625 | -2,122E-06 | -1,0335625 | -0,0913799 | 0,9271907 |
| VC1655 | 1,084318 | -0,2647582 | -1,0292653 | -2,584E-07 | -1,0292653 | -0,0910568 | 0,92744744 |
| nadE | 1,04512193 | -0,4447301 | -1,0124813 | -4,34E-07 | -1,0124813 | -0,0909663 | 0,92751938 |
| VC0509 | 1,06812281 | -0,2004437 | -1,0346465 | -1,956E-07 | -1,0346465 | -0,0908702 | 0,92759572 |
| VC0098 | 1,08771791 | -0,1902203 | -1,0290764 | -1,856E-07 | -1,0290764 | -0,0904299 | 0,92794563 |
| VC0674 | -1,0037638 | -1,3920584 | -1,010523 | -1,358E-06 | -1,010523 | -0,0901963 | 0,92813123 |
| VCA0361 | 1,03310226 | -0,32633 | -1,0227565 | -3,184E-07 | -1,0227565 | -0,0898788 | 0,9283835 |
| VCA0147 | 1,1356051 | 0,14287211 | 1,03821915 | 1,3942E-07 | 1,03821915 | 0,08964999 | 0,92856536 |
| VC1814 | 1,08616012 | -0,1034826 | -1,0528948 | -1,01E-07 | -1,0528948 | -0,0894371 | 0,92873453 |
| tbpA | 1,04359831 | -0,1648198 | -1,0328054 | -1,608E-07 | -1,0328054 | -0,0893287 | 0,92882071 |
| VCA0387 | 1,09440149 | 0,36404295 | 1,01811764 | 3,5524E-07 | 1,01811764 | 0,08867846 | 0,92933746 |
| VCA1012 | -1,0486923 | -2,1388426 | -1,0342985 | -2,087E-06 | -1,0342985 | -0,088366 | 0,92958577 |
| VCA0722 | 1,07441014 | -0,1756436 | -1,0299627 | -1,714E-07 | -1,0299627 | -0,0881911 | 0,92972481 |
| VCA0625 | 1,0618767 | -0,0908402 | -1,0585725 | -8,864E-08 | -1,0585725 | -0,088057 | 0,92983141 |
| VC1683 | 1,10338788 | 0,54931054 | 1,02061704 | 5,3602E-07 | 1,02061704 | 0,08769318 | 0,93012054 |
| VCA0855 | 1,0843474 | -0,0531499 | -1,100676 | -5,186E-08 | -1,100676 | -0,0874169 | 0,93034012 |
| VCA0216 | 1,12562343 | 0,24477624 | 1,02223615 | 2,3885E-07 | 1,02223615 | 0,08741582 | 0,930341 |
| VC1131 | 1,07079068 | 0,66477771 | 1,02901105 | 6,487E-07 | 1,02901105 | 0,08728425 | 0,93044558 |
| VCA0304 | 1,00647493 | -0,2096736 | -1,0325401 | -2,046E-07 | -1,0325401 | -0,0868903 | 0,93075872 |
| VC1629 | -1,0171603 | -0,7031199 | -1,0289745 | -6,861E-07 | -1,0289745 | -0,0864875 | 0,9310789 |
| greA | 1,12103713 | -7,3537537 | -1,018346 | -7,176E-06 | -1,018346 | -0,0862423 | 0,93127382 |
| VC1154 | -1,0075561 | -4,2266684 | -1,0233429 | -4,124E-06 | -1,0233429 | -0,086115 | 0,931375 |
| VC2008 | 1,07118202 | -0,7892337 | -1,0283161 | -7,701E-07 | -1,0283161 | -0,0857104 | 0,93169662 |
| VC0159 | 1,02306874 | -7,3273784 | -1,0266411 | -7,15E-06 | -1,0266411 | -0,0856857 | 0,9317163 |
| VCA0800 | 1,0257253 | -0,5452505 | -1,0119696 | -5,321E-07 | -1,0119696 | -0,0854473 | 0,93190577 |

| | | | | | | | |
|---|---|---|---|---|---|---|---|
| VCA0414 | 1,04013334 | -0,5780664 | -1,0275046 | -5,641E-07 | -1,0275046 | -0,0853739 | 0,93196413 |
| VC0661 | 1,07629459 | -0,1925516 | -1,0255314 | -1,879E-07 | -1,0255314 | -0,0853303 | 0,93199879 |
| 5Sb | -1,1594224 | 992,224048 | 1,02311193 | 0,00096822 | 1,02311193 | 0,08484466 | 0,93238489 |
| VC2443 | 1,07950897 | 0,52134864 | 1,01355234 | 5,0874E-07 | 1,01355234 | 0,08429738 | 0,93282 |
| VC1151 | 1,02949877 | -0,5893999 | -1,022377 | -5,751E-07 | -1,022377 | -0,083771 | 0,93323851 |
| VCA1032 | 1,0657537 | 0,33826026 | 1,01743941 | 3,3008E-07 | 1,01743941 | 0,08338062 | 0,9335489 |
| VCA0032 | 1,09856879 | 0,18062972 | 1,0255681 | 1,7626E-07 | 1,0255681 | 0,08270515 | 0,93408599 |
| VC1589 | 1,02964933 | 0,15849711 | 1,03965682 | 1,5466E-07 | 1,03965682 | 0,08225162 | 0,93444663 |
| VCA0782 | 1,08289761 | -0,1892505 | -1,0240196 | -1,847E-07 | -1,0240196 | -0,0820835 | 0,9345803 |
| VCA1082 | 1,02687784 | 0,72488481 | 1,01217018 | 7,0735E-07 | 1,01217018 | 0,08195896 | 0,93467935 |
| VC2007 | 1,01887899 | -3,1311803 | -1,0435065 | -3,055E-06 | -1,0435065 | -0,0819137 | 0,93471536 |
| VC1291 | 1,11582036 | 0,35787202 | 1,01255542 | 3,4921E-07 | 1,01255542 | 0,08184125 | 0,93477296 |
| fliH | 1,02461272 | 0,57938121 | 1,00769214 | 5,6537E-07 | 1,00769214 | 0,08160823 | 0,93495826 |
| VCA0864 | 1,09294628 | 0,26253319 | 1,01697209 | 2,5618E-07 | 1,01697209 | 0,08140925 | 0,93511649 |
| VC1349 | 1,08460404 | 0,21227498 | 1,02100449 | 2,0714E-07 | 1,02100449 | 0,08135507 | 0,93515958 |
| VC0252 | 1,0836063 | -10,272274 | -1,0163698 | -1,002E-05 | -1,0163698 | -0,0810775 | 0,93538034 |
| djlA | 1,07685147 | 0,40463143 | 1,01512699 | 3,9484E-07 | 1,01512699 | 0,08091724 | 0,93550778 |
| VC0015a | 1,07430904 | -0,1494256 | -1,0295768 | -1,458E-07 | -1,0295768 | -0,0808252 | 0,93558093 |
| VC0195 | 1,06907248 | -0,1543118 | -1,0285007 | -1,506E-07 | -1,0285007 | -0,0806495 | 0,93572069 |
| VC2529 | -1,0119505 | -3,1739442 | -1,0203384 | -3,097E-06 | -1,0203384 | -0,0805905 | 0,93576763 |
| VC1520 | -1,0149927 | -0,9312361 | -1,0274709 | -9,087E-07 | -1,0274709 | -0,0793197 | 0,93677838 |
| pssA | 1,07802459 | 0,36737474 | 1,0112319 | 3,5849E-07 | 1,0112319 | 0,07845449 | 0,93746653 |
| miaA | -1,0026296 | 0,81613612 | 1,00575685 | 7,9639E-07 | 1,00575685 | 0,07833389 | 0,93756246 |
| VC0236 | 1,04371855 | 0,63690985 | 1,0160141 | 6,215E-07 | 1,0160141 | 0,07787593 | 0,93792675 |
| VC0737 | 1,00722659 | -1,8960174 | -1,0368757 | -1,85E-06 | -1,0368757 | -0,0775098 | 0,93821801 |
| VCA0851 | 1,09404371 | 0,22629661 | 1,01783325 | 2,2082E-07 | 1,01783325 | 0,07745955 | 0,93825797 |
| VCA0605 | 1,26103915 | 0,04842225 | 1,08611943 | 4,7251E-08 | 1,08611943 | 0,0774398 | 0,93827368 |
| VCA0710 | 1,0726196 | -0,3262262 | -1,0162111 | -3,183E-07 | -1,0162111 | -0,0772722 | 0,93840698 |
| VC2135 | 1,06048199 | 0,41202453 | 1,0096745 | 4,0206E-07 | 1,0096745 | 0,07714052 | 0,93851176 |
| VCA0224 | 1,00736187 | -0,0496558 | -1,0829836 | -4,845E-08 | -1,0829836 | -0,0770369 | 0,93859419 |
| VCA0642 | 1,40116875 | 0,0474289 | 1,08608335 | 4,6282E-08 | 1,08608335 | 0,07662598 | 0,93892109 |
| VC0341 | 1,02611394 | -0,519233 | -1,0107007 | -5,067E-07 | -1,0107007 | -0,0757077 | 0,93965163 |
| VC1684 | 1,09154035 | 0,4790134 | 1,01723285 | 4,6743E-07 | 1,01723285 | 0,07569901 | 0,93965857 |
| VC1699 | -1,0061424 | -2,5359759 | -1,0191944 | -2,475E-06 | -1,0191944 | -0,0750509 | 0,94017422 |
| VC1817 | 1,25336969 | 0,07944014 | 1,04819596 | 7,7518E-08 | 1,04819596 | 0,07488598 | 0,94030543 |
| lnt | 1,02080126 | -0,606287 | -1,0131106 | -5,916E-07 | -1,0131106 | -0,0743136 | 0,94076085 |
| VC2740 | 1,07589172 | 0,38089402 | 1,0095408 | 3,7168E-07 | 1,0095408 | 0,07365719 | 0,94128317 |
| VC0888 | 1,08684883 | 0,23030362 | 1,01581345 | 2,2473E-07 | 1,01581345 | 0,0736211 | 0,94131188 |
| VCA1073 | 1,12093791 | -0,422986 | -1,050736 | -4,128E-07 | -1,050736 | -0,0735671 | 0,94135489 |
| VCA0546 | 1,06410297 | 0,43824933 | 1,01936787 | 4,2765E-07 | 1,01936787 | 0,07347926 | 0,94142475 |
| VC2764 | 1,15520895 | 5,71230022 | 1,00835471 | 5,5741E-06 | 1,00835471 | 0,07345174 | 0,94144665 |
| VCA0151 | 1,08734132 | -0,0926106 | -1,0395722 | -9,037E-08 | -1,0395722 | -0,0734204 | 0,94147156 |
| fabZ | 1,01333301 | -0,9496502 | -1,0085762 | -9,267E-07 | -1,0085762 | -0,0732234 | 0,94162832 |
| VCA0117 | 1,07095176 | -0,1200558 | -1,0299269 | -1,172E-07 | -1,0299269 | -0,0728692 | 0,9419102 |
| VC0283 | 1,23181985 | 0,05900806 | 1,06147477 | 5,7581E-08 | 1,06147477 | 0,07265675 | 0,94207927 |

| | | | | | | | |
|---|---|---|---|---|---|---|---|
| VC2073 | 1,02069172 | -0,1779978 | -1,0201722 | -1,737E-07 | -1,0201722 | -0,072444 | 0,94224857 |
| VC0311 | 1,07943438 | 0,26915372 | 1,01379657 | 2,6264E-07 | 1,01379657 | 0,07186333 | 0,94271068 |
| VC1376 | 1,05748958 | -1,6473748 | -1,0269661 | -1,608E-06 | -1,0269661 | -0,0717503 | 0,94280066 |
| VC2750 | 1,13867894 | 0,43094558 | 1,01959829 | 4,2052E-07 | 1,01959829 | 0,07134262 | 0,94312507 |
| tRNA-Cys-2 | 1,07616487 | 0,5759518 | 1,04315462 | 5,6202E-07 | 1,04315462 | 0,07129 | 0,94316695 |
| VC1809 | 1,05851455 | -0,2316419 | -1,0145405 | -2,26E-07 | -1,0145405 | -0,070823 | 0,94353862 |
| VC1321 | 1,03829381 | -0,445009 | -1,0094042 | -4,342E-07 | -1,0094042 | -0,0704585 | 0,94382873 |
| VCA0127 | 1,09774666 | -0,9908468 | -1,0512246 | -9,669E-07 | -1,0512246 | -0,0700065 | 0,94418851 |
| VCA0181 | 1,04192335 | -0,2698487 | -1,0119658 | -2,633E-07 | -1,0119658 | -0,0693884 | 0,94468047 |
| VC2330 | 1,06449742 | -0,1002241 | -1,0319583 | -9,78E-08 | -1,0319583 | -0,0687672 | 0,9451749 |
| VCA0781 | 1,09502923 | -0,13127 | -1,0242815 | -1,281E-07 | -1,0242815 | -0,0687301 | 0,94520443 |
| VC1273 | 1,07435338 | -0,0802606 | -1,0397036 | -7,832E-08 | -1,0397036 | -0,0684611 | 0,9454186 |
| VC2004 | 1,13360952 | 0,13303868 | 1,02324899 | 1,2982E-07 | 1,02324899 | 0,06772175 | 0,94600714 |
| VC2619 | 1,00550774 | -0,4119364 | -1,0260271 | -4,02E-07 | -1,0260271 | -0,0674009 | 0,94626259 |
| VC0340 | 1,04195035 | -0,3917483 | -1,0077542 | -3,823E-07 | -1,0077542 | -0,0673733 | 0,94628451 |
| VC1104 | 1,05913478 | 0,22550369 | 1,01350879 | 2,2005E-07 | 1,01350879 | 0,06737092 | 0,94628642 |
| VC1902 | 1,07857141 | 0,26086957 | 1,01155122 | 2,5456E-07 | 1,01155122 | 0,0670387 | 0,9465509 |
| VCA1101 | 1,16839806 | 0,07385687 | 1,04113142 | 7,207E-08 | 1,04113142 | 0,06682021 | 0,94672483 |
| VC1878 | 1,03241297 | 0,54297319 | 1,00958575 | 5,2984E-07 | 1,00958575 | 0,06673276 | 0,94679445 |
| VC1523 | 1,05441735 | 1,94432232 | 1,02739005 | 1,8973E-06 | 1,02739005 | 0,06647547 | 0,94699929 |
| VC1052 | 1,03303278 | -0,7466271 | -1,014532 | -7,286E-07 | -1,014532 | -0,0661616 | 0,94724919 |
| VCA0122 | 1,06312754 | 0,17861852 | 1,01642754 | 1,743E-07 | 1,01642754 | 0,06607257 | 0,94732005 |
| VC1050 | 1,06081369 | -0,2794243 | -1,0144733 | -2,727E-07 | -1,0144733 | -0,0660511 | 0,94733712 |
| VCA1069 | 1,0325778 | -5,4871855 | -1,0392565 | -5,354E-06 | -1,0392565 | -0,0660478 | 0,94733974 |
| VC1322 | 1,09278341 | -0,2472543 | -1,0117353 | -2,413E-07 | -1,0117353 | -0,0656926 | 0,94762253 |
| VCA0790 | 1,02505354 | -0,3949096 | -1,0202877 | -3,854E-07 | -1,0202877 | -0,0655204 | 0,94775967 |
| VC0935 | 1,13103647 | 0,10616523 | 1,02732413 | 1,036E-07 | 1,02732413 | 0,06551854 | 0,94776115 |
| VC0489 | 1,06597183 | -0,0971119 | -1,0298628 | -9,476E-08 | -1,0298628 | -0,0654681 | 0,94780132 |
| VCA0152 | 1,13032329 | 0,13154798 | 1,02188296 | 1,2837E-07 | 1,02188296 | 0,06535503 | 0,94789133 |
| VCA0583 | 1,07037478 | -0,1041529 | -1,027481 | -1,016E-07 | -1,027481 | -0,0650782 | 0,94811175 |
| VCA0220 | 1,10647272 | 2,65863098 | 1,02969609 | 2,5943E-06 | 1,02969609 | 0,06484721 | 0,94829566 |
| VC0497 | 1,07627495 | 0,13943778 | 1,02010037 | 1,3606E-07 | 1,02010037 | 0,06451607 | 0,94855932 |
| VC0804 | 1,07181453 | -0,1146197 | -1,0280652 | -1,118E-07 | -1,0280652 | -0,0644708 | 0,94859537 |
| VC2220 | 1,08536554 | -0,2116182 | -1,0152137 | -2,065E-07 | -1,0152137 | -0,0643762 | 0,94867065 |
| VCA0596 | 1,05116134 | -0,1440299 | -1,0192474 | -1,405E-07 | -1,0192474 | -0,064177 | 0,94882927 |
| VC2633 | 1,12364612 | 0,07221432 | 1,03854871 | 7,0467E-08 | 1,03854871 | 0,06400547 | 0,94896588 |
| VCA0983 | 1,123243 | -0,1543172 | -1,0317244 | -1,506E-07 | -1,0317244 | -0,0638914 | 0,94905672 |
| VC2623 | 1,01604855 | 0,68750995 | 1,00804178 | 6,7088E-07 | 1,00804178 | 0,06377262 | 0,94915128 |
| VC0116 | 1,08516413 | 0,3377053 | 1,01009702 | 3,2954E-07 | 1,01009702 | 0,06372856 | 0,94918637 |
| VCA0973 | 1,077471 | 0,12712213 | 1,04295571 | 1,2405E-07 | 1,04295571 | 0,06346863 | 0,94939334 |
| VC0282 | 1,14741078 | 0,38400392 | 1,0266634 | 3,7471E-07 | 1,0266634 | 0,06318583 | 0,94961853 |
| VC2285 | 1,05798617 | 0,31700289 | 1,00828946 | 3,0933E-07 | 1,00828946 | 0,06265424 | 0,95004184 |
| VC0844 | 1,03982879 | -0,1734717 | -1,0183837 | -1,693E-07 | -1,0183837 | -0,0625107 | 0,95015616 |
| VCA0148 | 1,17933858 | 0,09285384 | 1,0282667 | 9,0608E-08 | 1,0282667 | 0,06230692 | 0,95031842 |
| VC1940 | 1,0662919 | -0,1665613 | -1,0156443 | -1,625E-07 | -1,0156443 | -0,062276 | 0,95034306 |

| | | | | | | | |
|---|---|---|---|---|---|---|---|
| VC1549 | 1,16577464 | 0,05940139 | 1,04440402 | 5,7964E-08 | 1,04440402 | 0,06221384 | 0,95039254 |
| VC0684 | 1,06790978 | 0,32342258 | 1,00899814 | 3,156E-07 | 1,00899814 | 0,06164831 | 0,9508429 |
| VC0826 | -1,1036507 | -37,385952 | -1,0224364 | -3,648E-05 | -1,0224364 | -0,0607427 | 0,95156414 |
| VC1924 | 1,06747342 | -0,1355621 | -1,0181632 | -1,323E-07 | -1,0181632 | -0,0604992 | 0,95175808 |
| VCA0928 | 1,08401674 | -0,1105817 | -1,0222435 | -1,079E-07 | -1,0222435 | -0,0604071 | 0,95183144 |
| VCA0374 | 1,12420714 | 0,10664393 | 1,0230192 | 1,0406E-07 | 1,0230192 | 0,06033576 | 0,95188822 |
| aspA | 1,06819159 | 35,4308609 | 1,0336701 | 3,4574E-05 | 1,0336701 | 0,06029279 | 0,95192245 |
| VC1879 | 1,07475166 | 0,28577015 | 1,01513506 | 2,7886E-07 | 1,01513506 | 0,05989684 | 0,9522378 |
| VC0154 | 1,05251271 | -0,1719666 | -1,0139811 | -1,678E-07 | -1,0139811 | -0,0598451 | 0,95227904 |
| VCA1049 | 1,08960172 | -0,0956428 | -1,0252734 | -9,333E-08 | -1,0252734 | -0,0598381 | 0,95228461 |
| VC1382 | 1,02588306 | -0,2275595 | -1,0111002 | -2,221E-07 | -1,0111002 | -0,0595425 | 0,95252006 |
| VC1041 | 1,09103564 | 0,19036681 | 1,01230342 | 1,8576E-07 | 1,01230342 | 0,05909159 | 0,95287916 |
| VCA0957 | 1,08051025 | -0,1280676 | -1,0181055 | -1,25E-07 | -1,0181055 | -0,0587104 | 0,95318281 |
| lpxK | 1,04862908 | 0,59728811 | 1,01274837 | 5,8284E-07 | 1,01274837 | 0,05859157 | 0,95327742 |
| VC2430 | 1,09151377 | 0,43387684 | 1,01401322 | 4,2338E-07 | 1,01401322 | 0,05844755 | 0,95339214 |
| VCA0142 | 1,1009872 | 0,29938047 | 1,01654582 | 2,9214E-07 | 1,01654582 | 0,05842879 | 0,95340709 |
| VC2544 | 1,07216434 | 3,06552553 | 1,03141726 | 2,9914E-06 | 1,03141726 | 0,05830008 | 0,95350961 |
| VC2605 | -1,0348715 | -2,2900005 | -1,0215078 | -2,235E-06 | -1,0215078 | -0,0581679 | 0,9536149 |
| VCA0287 | 1,20398227 | 6,38250494 | 1,01385506 | 6,2281E-06 | 1,01385506 | 0,05805929 | 0,95370141 |
| VC1749 | 1,09906476 | -0,0985152 | -1,0228241 | -9,613E-08 | -1,0228241 | -0,0577472 | 0,95395003 |
| VCA0036 | 1,07478195 | 0,87938574 | 1,02161341 | 8,5811E-07 | 1,02161341 | 0,05768353 | 0,95400071 |
| VC0401 | 1,06877345 | 0,32476835 | 1,00887262 | 3,1691E-07 | 1,00887262 | 0,05756176 | 0,95409771 |
| VC2537 | 1,07353348 | 0,12302559 | 1,01810539 | 1,2005E-07 | 1,01810539 | 0,05754292 | 0,95411272 |
| VC1270 | 1,07149616 | 0,54327425 | 1,01057937 | 5,3013E-07 | 1,01057937 | 0,05740808 | 0,95422013 |
| VCA0409 | 1,10179925 | 0,60045903 | 1,01526299 | 5,8593E-07 | 1,01526299 | 0,05728585 | 0,95431749 |
| VC0703 | 1,05412438 | -0,1730368 | -1,0125583 | -1,689E-07 | -1,0125583 | -0,0569147 | 0,95461314 |
| VC0149 | 1,02269972 | -0,3975095 | -1,0065962 | -3,879E-07 | -1,0065962 | -0,056904 | 0,95462165 |
| tRNA-Ser-3 | 1,01001563 | 0,67883864 | 1,01499105 | 6,6242E-07 | 1,01499105 | 0,05667225 | 0,95480628 |
| VC1145 | 1,06385255 | 0,25978071 | 1,00944757 | 2,535E-07 | 1,00944757 | 0,05644255 | 0,95498927 |
| VC1941 | 1,02967355 | -0,2631618 | -1,008458 | -2,568E-07 | -1,008458 | -0,0563194 | 0,95508734 |
| VC1287 | 1,04215424 | -0,5506903 | -1,0144846 | -5,374E-07 | -1,0144846 | -0,0563051 | 0,95509873 |
| phoR | 1,0947671 | 0,12146559 | 1,01726355 | 1,1853E-07 | 1,01726355 | 0,05584348 | 0,9554665 |
| VC0354 | 1,11943505 | 13,2937373 | 1,01639791 | 1,2972E-05 | 1,01639791 | 0,05565889 | 0,95561356 |
| VC2561 | -1,0756216 | -0,0805179 | -1,044778 | -7,857E-08 | -1,044778 | -0,0555887 | 0,95566945 |
| tRNA-Cys-3 | 1,01527202 | -0,0418186 | -1,0501135 | -4,081E-08 | -1,0501135 | -0,0553776 | 0,95583764 |
| maf (NC_002505 2179182..2179804) | 1,03784176 | 0,58027668 | 1,01292527 | 5,6624E-07 | 1,01292527 | 0,05528437 | 0,95591192 |
| VC1704 | 1,16143511 | 0,0839856 | 1,02443 | 8,1954E-08 | 1,02443 | 0,05514093 | 0,95602619 |
| VC1334 | 1,22133794 | 0,05469292 | 1,03767305 | 5,337E-08 | 1,03767305 | 0,05507756 | 0,95607668 |
| VC2484 | -1,017089 | 2,60946757 | 1,01894212 | 2,5463E-06 | 1,01894212 | 0,05492255 | 0,95620017 |
| VC0176 | 1,07273408 | 0,18119729 | 1,01257422 | 1,7681E-07 | 1,01257422 | 0,0542707 | 0,9567195 |
| VCA0120 | 1,0346268 | -0,1091433 | -1,0179073 | -1,065E-07 | -1,0179073 | -0,0539045 | 0,95701123 |
| VCA0221 | 1,09229796 | -0,0495044 | -1,0398055 | -4,831E-08 | -1,0398055 | -0,0538344 | 0,95706708 |
| VC2054 | -1,0500796 | -2,3662559 | -1,0214667 | -2,309E-06 | -1,0214667 | -0,0537452 | 0,95713816 |
| VC1653 | 1,05520739 | 0,15956468 | 1,01442523 | 1,557E-07 | 1,01442523 | 0,05344488 | 0,95737745 |
| tRNA-Tyr-3 | 1,13737107 | 0,23782179 | 1,03630723 | 2,3207E-07 | 1,03630723 | 0,05343657 | 0,95738407 |

| | | | | | | | |
|---|---|---|---|---|---|---|---|
| VCA0413 | 1,12860748 | 0,52631846 | 1,01756177 | 5,1359E-07 | 1,01756177 | 0,05338786 | 0,95742288 |
| VCA0556 | 1,10076391 | -0,0717447 | -1,0267176 | -7,001E-08 | -1,0267176 | -0,053267 | 0,95751919 |
| VC2056 | 1,05905137 | -0,2256893 | -1,0083947 | -2,202E-07 | -1,0083947 | -0,0531987 | 0,95757362 |
| VC2505 | 1,00069583 | -0,4393753 | -1,0145671 | -4,287E-07 | -1,0145671 | -0,0530761 | 0,9576713 |
| VC1468 | 1,10219257 | 0,10174911 | 1,0183399 | 9,9288E-08 | 1,0183399 | 0,05266578 | 0,95799821 |
| VC0192 | 1,07837246 | -0,1185186 | -1,015546 | -1,157E-07 | -1,015546 | -0,0523682 | 0,95823529 |
| fxsA | 1,08368278 | 0,72077528 | 1,01302385 | 7,0334E-07 | 1,01302385 | 0,05217042 | 0,95839291 |
| VC0208 | 1,04714202 | -0,2509369 | -1,007218 | -2,449E-07 | -1,007218 | -0,052031 | 0,95850397 |
| VC2042 | 1,05234205 | -0,141219 | -1,0128403 | -1,378E-07 | -1,0128403 | -0,0519866 | 0,95853934 |
| VC2530 | 1,02258775 | 9,12878488 | 1,02122711 | 8,9079E-06 | 1,02122711 | 0,05125899 | 0,95911915 |
| tRNA-Pro-4 | 1,06377967 | 0,30228897 | 1,03088204 | 2,9498E-07 | 1,03088204 | 0,05061601 | 0,95963151 |
| VC1327 | 1,03404998 | -5,0679053 | -1,0263606 | -4,945E-06 | -1,0263606 | -0,0504667 | 0,95975046 |
| VC1988 | 1,08885526 | 0,20762833 | 1,00934848 | 2,0261E-07 | 1,00934848 | 0,05023629 | 0,95993409 |
| VCA0872 | 1,09648139 | 0,12003522 | 1,01401688 | 1,1713E-07 | 1,01401688 | 0,05006232 | 0,96007273 |
| VC1505 | 1,09751333 | 0,22576229 | 1,0145636 | 2,203E-07 | 1,0145636 | 0,04954647 | 0,96048381 |
| VC0806 | 1,07527928 | 0,14966787 | 1,01710861 | 1,4605E-07 | 1,01710861 | 0,04939422 | 0,96060514 |
| VC1275 | 1,15436606 | 0,0881021 | 1,0185891 | 8,5971E-08 | 1,0185891 | 0,04933556 | 0,96065188 |
| prfB | 1,01510657 | 1,18672809 | 1,01314361 | 1,158E-06 | 1,01314361 | 0,04900393 | 0,96091617 |
| 5Sa | 1,00370239 | 91,0096958 | 1,01848942 | 8,8808E-05 | 1,01848942 | 0,04873611 | 0,9611296 |
| 5Se | -1,3078267 | -135,73507 | -1,0157493 | -0,0001325 | -1,0157493 | -0,0486458 | 0,96120158 |
| VC2408 | 1,1051152 | 0,13674088 | 1,01159963 | 1,3343E-07 | 1,01159963 | 0,04863664 | 0,96120887 |
| VCA0483 | 1,09616228 | 0,6805988 | 1,01629835 | 6,6413E-07 | 1,01629835 | 0,04850279 | 0,96131554 |
| aat | 1,11304525 | 0,06493389 | 1,024433 | 6,3363E-08 | 1,024433 | 0,04848792 | 0,9613274 |
| VCA0307 | 1,07445642 | 0,38413428 | 1,01082756 | 3,7484E-07 | 1,01082756 | 0,04835522 | 0,96143315 |
| VCA0531 | 1,07534034 | 0,11741279 | 1,01231701 | 1,1457E-07 | 1,01231701 | 0,04643279 | 0,9629653 |
| VC0424 | -1,0080941 | -1,3425875 | -1,016528 | -1,31E-06 | -1,016528 | -0,0462022 | 0,96314909 |
| VCA0635 | 1,07329883 | -0,0851693 | -1,0166785 | -8,311E-08 | -1,0166785 | -0,0459688 | 0,9633351 |
| VC0264 | 1,0654796 | 0,21866214 | 1,008565 | 2,1337E-07 | 1,008565 | 0,04595822 | 0,96334355 |
| gmk | -1,0124041 | -0,4863556 | -1,0172191 | -4,746E-07 | -1,0172191 | -0,0447735 | 0,96428784 |
| VC0370 | 1,03368225 | -0,2694591 | -1,0151489 | -2,629E-07 | -1,0151489 | -0,0447697 | 0,96429085 |
| VC2732 | 1,0894656 | 0,18296796 | 1,00907446 | 1,7854E-07 | 1,00907446 | 0,04474896 | 0,9643074 |
| VCA0389 | 1,06315206 | 0,20207902 | 1,01096225 | 1,9719E-07 | 1,01096225 | 0,04463147 | 0,96440106 |
| tRNA-Arg-1 | -1,0426181 | -1,0985505 | -1,017597 | -1,072E-06 | -1,017597 | -0,0440748 | 0,96484475 |
| VCA0599 | 1,06063258 | -0,1256391 | -1,0102982 | -1,226E-07 | -1,0102982 | -0,0439416 | 0,96495092 |
| VC0745 | -1,0377956 | -1,6226039 | -1,0206698 | -1,583E-06 | -1,0206698 | -0,0438086 | 0,96505695 |
| VC1635 | 1,01628828 | 0,65681854 | 1,01008661 | 6,4093E-07 | 1,01008661 | 0,04324181 | 0,96550877 |
| VCA0808 | -1,009488 | 1,06977031 | 1,01215119 | 1,0439E-06 | 1,01215119 | 0,04287059 | 0,96580469 |
| 16Sg | 1,03865195 | -0,0959129 | -1,012533 | -9,359E-08 | -1,012533 | -0,0423308 | 0,96623503 |
| VC2146 | 1,02061305 | -0,4268974 | -1,0095417 | -4,166E-07 | -1,0095417 | -0,0421576 | 0,96637307 |
| tRNA-Asp-3 | -1,0058803 | -0,390787 | -1,0199671 | -3,813E-07 | -1,0199671 | -0,0408937 | 0,96738064 |
| VCA0783 | 1,10507291 | 0,14179597 | 1,00774406 | 1,3837E-07 | 1,00774406 | 0,04050669 | 0,96768917 |
| VC0602 | 1,06209277 | 0,14024754 | 1,0076623 | 1,3685E-07 | 1,0076623 | 0,0400725 | 0,96803532 |
| VC0969 | 1,02370326 | -0,2460306 | -1,0053114 | -2,401E-07 | -1,0053114 | -0,0399116 | 0,9681636 |
| VCA1108 | 1,0321321 | -0,0488457 | -1,0208183 | -4,766E-08 | -1,0208183 | -0,0388538 | 0,96900695 |
| VC1503 | -1,0248015 | -0,8517078 | -1,0100936 | -8,311E-07 | -1,0100936 | -0,0387541 | 0,9690864 |

| | | | | | | | |
|---|---|---|---|---|---|---|---|
| VCA1040 | 1,07363663 | 0,15028669 | 1,00658304 | 1,4665E-07 | 1,00658304 | 0,03846009 | 0,96932085 |
| VC0462 | 1,09239065 | 0,127292 | 1,00776529 | 1,2421E-07 | 1,00776529 | 0,03843147 | 0,96934367 |
| VCA0121 | 1,09292341 | 0,07760049 | 1,01260573 | 7,5723E-08 | 1,01260573 | 0,03818553 | 0,96953976 |
| VC2055 | -1,0058089 | 1,29357418 | 1,01490453 | 1,2623E-06 | 1,01490453 | 0,03743213 | 0,97014046 |
| VC0519 | 1,00639958 | -12,261116 | -1,0093206 | -1,196E-05 | -1,0093206 | -0,0373508 | 0,97020534 |
| sdhB | 1,04027723 | 1,76932101 | 1,02049146 | 1,7265E-06 | 1,02049146 | 0,03558919 | 0,97160993 |
| VC1372 | 1,04971462 | -0,1204645 | -1,0069585 | -1,176E-07 | -1,0069585 | -0,0353984 | 0,97176209 |
| VCA0735 | 1,04183664 | 0,25379346 | 1,00406012 | 2,4765E-07 | 1,00406012 | 0,03508665 | 0,97201065 |
| slmA | 1,07212239 | -0,2529516 | -1,0071398 | -2,468E-07 | -1,0071398 | -0,0348925 | 0,97216547 |
| VC0974 | 1,09370456 | 0,26840704 | 1,00758872 | 2,6191E-07 | 1,00758872 | 0,03487642 | 0,97217829 |
| VC1446 | 1,07043163 | -0,0887319 | -1,0088784 | -8,659E-08 | -1,0088784 | -0,0343 | 0,97263795 |
| VC2471 | 1,01308036 | 1,35704339 | 1,00733132 | 1,3242E-06 | 1,00733132 | 0,03397567 | 0,97289656 |
| VC1274 | 1,11545832 | -0,0593571 | -1,0129624 | -5,792E-08 | -1,0129624 | -0,0338628 | 0,97298656 |
| VCA0669 | 1,08314604 | -0,040036 | -1,0191448 | -3,907E-08 | -1,0191448 | -0,0337465 | 0,97307933 |
| tRNA-Tyr-5 | 1,03980194 | -0,1559443 | -1,0226811 | -1,522E-07 | -1,0226811 | -0,0335035 | 0,97327307 |
| VCA0328 | 1,03874613 | -0,0881048 | -1,008439 | -8,597E-08 | -1,008439 | -0,0333257 | 0,9734149 |
| VCA1089 | 1,01518684 | -0,0451259 | -1,0164711 | -4,403E-08 | -1,0164711 | -0,0332536 | 0,97347232 |
| VC1952 | 1,12186825 | -0,0578654 | -1,0127823 | -5,647E-08 | -1,0127823 | -0,033203 | 0,97351268 |
| VC0042 | 1,0632788 | -0,1586999 | -1,0046125 | -1,549E-07 | -1,0046125 | -0,0330985 | 0,97359601 |
| VC2379 | 1,04856393 | 0,17337162 | 1,00410465 | 1,6918E-07 | 1,00410465 | 0,03263897 | 0,97396249 |
| VC0993 | 1,04712078 | -1,1993925 | -1,0114344 | -1,17E-06 | -1,0114344 | -0,0325569 | 0,97402791 |
| VC1009 | 1,02781306 | -0,2463628 | -1,0049493 | -2,404E-07 | -1,0049493 | -0,0321803 | 0,97432824 |
| VCA0547 | 1,09759812 | -4,725167 | -1,0133858 | -4,611E-06 | -1,0133858 | -0,0321607 | 0,97434393 |
| def (NC_002506 167966..168513) | 1,03159468 | 0,20486183 | 1,01278152 | 1,9991E-07 | 1,01278152 | 0,03208194 | 0,97440671 |
| VC2488 | 1,13170755 | -0,0619188 | -1,0110436 | -6,042E-08 | -1,0110436 | -0,0319388 | 0,97452088 |
| VC0611 | 1,07854124 | 0,07067372 | 1,00955708 | 6,8964E-08 | 1,00955708 | 0,03175439 | 0,97466793 |
| VC0495 | 1,11520912 | -0,0624121 | -1,0106925 | -6,09E-08 | -1,0106925 | -0,0315547 | 0,97482717 |
| VCA0292 | 1,06307062 | 0,67317223 | 1,01159781 | 6,5689E-07 | 1,01159781 | 0,0314657 | 0,97489815 |
| VC1609 | 1,0623064 | 0,11286597 | 1,00691099 | 1,1014E-07 | 1,00691099 | 0,03136922 | 0,97497509 |
| VCA0428 | -1,0159993 | -0,5139312 | -1,022673 | -5,015E-07 | -1,022673 | -0,03114 | 0,97515788 |
| VCA0875 | 1,00167041 | -0,8939185 | -1,0058515 | -8,723E-07 | -1,0058515 | -0,0304798 | 0,97568444 |
| VC1304 | 1,05909045 | 2,55745174 | 1,01374305 | 2,4956E-06 | 1,01374305 | 0,03019979 | 0,97590772 |
| VC1396 | 1,04535209 | 0,11414468 | 1,00728071 | 1,1138E-07 | 1,00728071 | 0,0300449 | 0,97603124 |
| VC2775 | 1,01228749 | -0,3483524 | -1,0050142 | -3,399E-07 | -1,0050142 | -0,0290227 | 0,97684653 |
| secD (NC_002505 795818..797712) | 1,06763312 | 0,37965689 | 1,00557202 | 3,7047E-07 | 1,00557202 | 0,02891883 | 0,97692933 |
| VC1339 | 1,14995519 | 0,04145005 | 1,01548716 | 4,0447E-08 | 1,01548716 | 0,02865621 | 0,97713877 |
| VC2573 | 1,1249211 | 2,33532121 | 1,00216978 | 2,2788E-06 | 1,00216978 | 0,02859567 | 0,97718706 |
| VCA0891 | 1,08580864 | 0,18997045 | 1,01331727 | 1,8537E-07 | 1,01331727 | 0,0285383 | 0,97723282 |
| engA | -1,0113075 | -0,935935 | -1,0087575 | -9,133E-07 | -1,0087575 | -0,0284128 | 0,97733292 |
| VC2216 | -1,0185294 | 0,33313472 | 1,0045652 | 3,2508E-07 | 1,0045652 | 0,02841122 | 0,97733419 |
| VC0499 | 1,07162136 | 0,06780594 | 1,0078312 | 6,6166E-08 | 1,0078312 | 0,02816741 | 0,97752863 |
| VC2773 | 1,03649438 | 0,39689016 | 1,00417429 | 3,8729E-07 | 1,00417429 | 0,02804902 | 0,97762305 |
| VC1703 | 1,07107192 | 0,88007183 | 1,00866744 | 8,5878E-07 | 1,00866744 | 0,02773213 | 0,97787579 |
| tRNA-Tyr-2 | -1,0690288 | -0,6139224 | -1,0201142 | -5,991E-07 | -1,0201142 | -0,0277108 | 0,97789282 |
| VC0382 | 1,077094 | -0,0840985 | -1,0074935 | -8,206E-08 | -1,0074935 | -0,0274292 | 0,9781174 |

| | | | | | | | |
|---|---|---|---|---|---|---|---|
| VCA0686 | 1,0656401 | 0,36446009 | 1,03146764 | 3,5564E-07 | 1,03146764 | 0,0274202 | 0,97812458 |
| VC1065 | 1,02646548 | 0,17374747 | 1,00288317 | 1,6954E-07 | 1,00288317 | 0,02739301 | 0,97814628 |
| VC0434 | 1,03787152 | -0,2191992 | -1,0044812 | -2,139E-07 | -1,0044812 | -0,0272513 | 0,97825934 |
| VC1619 | 1,04426305 | -0,0942327 | -1,0053237 | -9,195E-08 | -1,0053237 | -0,0270081 | 0,97845328 |
| VC1160 | 1,04937827 | -0,1425387 | -1,0054053 | -1,391E-07 | -1,0054053 | -0,0264463 | 0,97890137 |
| VCA0682 | 1,09478868 | 0,16665175 | 1,01249172 | 1,6262E-07 | 1,01249172 | 0,02604354 | 0,97922261 |
| VCA0552 | 1,09549901 | 0,06232065 | 1,00722077 | 6,0813E-08 | 1,00722077 | 0,02593418 | 0,97930984 |
| VC1207 | 1,10702934 | 0,16976508 | 1,00713805 | 1,6566E-07 | 1,00713805 | 0,02584358 | 0,97938211 |
| VC0017 | 1,03738672 | 0,06839512 | 1,00623896 | 6,6741E-08 | 1,00623896 | 0,02526039 | 0,97984727 |
| VC1667 | 1,04670768 | -0,073387 | -1,0056883 | -7,161E-08 | -1,0056883 | -0,0249882 | 0,98006441 |
| VC0059 | 1,13251967 | -4,8212275 | -1,0109799 | -4,705E-06 | -1,0109799 | -0,0241194 | 0,98075739 |
| VC1323 | 1,10027107 | -0,1124776 | -1,0052014 | -1,098E-07 | -1,0052014 | -0,0239967 | 0,98085527 |
| VC0708 | 1,00036747 | -0,3031706 | -1,0019252 | -2,958E-07 | -1,0019252 | -0,0237489 | 0,98105292 |
| psd | 1,05231805 | -0,1154421 | -1,003671 | -1,126E-07 | -1,003671 | -0,0232746 | 0,98143123 |
| VCA0091 | 1,07235675 | -0,0666836 | -1,0053648 | -6,507E-08 | -1,0053648 | -0,023134 | 0,98154338 |
| VCA0118 | 1,01383887 | -0,029177 | -1,011921 | -2,847E-08 | -1,011921 | -0,0227736 | 0,98183084 |
| VCA0969 | 1,113191 | 0,10958511 | 1,00521996 | 1,0693E-07 | 1,00521996 | 0,02261173 | 0,98195999 |
| cobT | 1,07442552 | -0,1243305 | -1,0070939 | -1,213E-07 | -1,0070939 | -0,0225298 | 0,98202535 |
| VC1049 | -1,0018894 | -0,8526893 | -1,0080229 | -8,321E-07 | -1,0080229 | -0,0224376 | 0,98209886 |
| apaH | 1,05170228 | -0,1192041 | -1,0036522 | -1,163E-07 | -1,0036522 | -0,0223163 | 0,98219567 |
| VC2547 | 1,05070296 | 0,09148154 | 1,00360681 | 8,9268E-08 | 1,00360681 | 0,02222736 | 0,9822666 |
| VC2281 | 1,05759478 | -0,0787336 | -1,0040455 | -7,683E-08 | -1,0040455 | -0,0218363 | 0,98257855 |
| VC0306 | 1,12661482 | -3,8692292 | -1,005329 | -3,776E-06 | -1,005329 | -0,0218253 | 0,98258735 |
| VCA0232 | 1,07729531 | -0,0233141 | -1,0133591 | -2,275E-08 | -1,0133591 | -0,0215426 | 0,98281284 |
| VC1142 | 1,05135589 | -0,3090225 | -1,0044988 | -3,015E-07 | -1,0044988 | -0,0215363 | 0,98281786 |
| VC2359 | 1,06599675 | -0,0817492 | -1,003681 | -7,977E-08 | -1,003681 | -0,0212265 | 0,98306501 |
| VC1634 | 1,0429903 | -0,114825 | -1,0050145 | -1,12E-07 | -1,0050145 | -0,0210794 | 0,9831823 |
| VC1500 | 1,04836789 | -0,0526145 | -1,0055954 | -5,134E-08 | -1,0055954 | -0,020985 | 0,98325765 |
| VC2423 | 1,06910389 | 0,1467509 | 1,00623023 | 1,432E-07 | 1,00623023 | 0,02048962 | 0,9836528 |
| nrdG | 1,08085073 | 0,7167242 | 1,01314909 | 6,9939E-07 | 1,01314909 | 0,0203854 | 0,98373594 |
| VCA0281 | 1,06353005 | -0,5539978 | -1,0081761 | -5,406E-07 | -1,0081761 | -0,0201528 | 0,98392152 |
| VC0629 | 1,05167578 | 0,05663924 | 1,00467786 | 5,5269E-08 | 1,00467786 | 0,01991238 | 0,98411327 |
| VC0594 | 1,03733955 | 0,09992923 | 1,00260491 | 9,7512E-08 | 1,00260491 | 0,01974757 | 0,98424473 |
| VC1706 | 1,088906 | -0,0422328 | -1,006379 | -4,121E-08 | -1,006379 | -0,0196754 | 0,98430232 |
| VCA1048 | 1,11184858 | 0,08315356 | 1,00390779 | 8,1142E-08 | 1,00390779 | 0,01960195 | 0,98436091 |
| VCA0071 | 1,19671637 | -0,0172127 | -1,0149022 | -1,68E-08 | -1,0149022 | -0,0195426 | 0,98440822 |
| VC0549 | 1,08895298 | 0,52389395 | 1,00891599 | 5,1122E-07 | 1,00891599 | 0,01921302 | 0,98467117 |
| VC2235 | 1,06352841 | -0,1024033 | -1,0023911 | -9,993E-08 | -1,0023911 | -0,0191537 | 0,98471853 |
| VCA1000 | 1,0825431 | -0,0299811 | -1,0085458 | -2,926E-08 | -1,0085458 | -0,0185648 | 0,98518829 |
| VC0998 | 1,0338889 | -0,2747815 | -1,0046625 | -2,681E-07 | -1,0046625 | -0,0183762 | 0,98533877 |
| VC1557 | 1,11075329 | 0,09285187 | 1,0044849 | 9,0606E-08 | 1,0044849 | 0,01762234 | 0,98594013 |
| VCA0268 | 1,08520222 | -0,0348382 | -1,0059564 | -3,4E-08 | -1,0059564 | -0,0176166 | 0,9859447 |
| VCA0263 | 1,08023809 | 0,049067 | 1,00415052 | 4,788E-08 | 1,00415052 | 0,01745998 | 0,98606966 |
| VCA0650 | 1,06197724 | -0,0145404 | -1,0139448 | -1,419E-08 | -1,0139448 | -0,0173793 | 0,98613404 |
| VC0697 | 1,10206719 | 0,15625544 | 1,00433647 | 1,5248E-07 | 1,00433647 | 0,01663555 | 0,98672736 |

| | | | | | | | |
|---|---|---|---|---|---|---|---|
| VCA0274a | 1,0540241 | 0,08312559 | 1,00372877 | 8,1115E-08 | 1,00372877 | 0,01654285 | 0,98680131 |
| VC1401 | 1,06274666 | -0,0246896 | -1,0070547 | -2,409E-08 | -1,0070547 | -0,0161354 | 0,98712639 |
| VC0309 | 1,06509336 | -0,0463642 | -1,0037427 | -4,524E-08 | -1,0037427 | -0,0161185 | 0,98713987 |
| VC1570 | 1,08081429 | 0,03679611 | 1,00660604 | 3,5906E-08 | 1,00660604 | 0,01565237 | 0,98751174 |
| VC2425 | 1,02044512 | -0,0187072 | -1,0085304 | -1,825E-08 | -1,0085304 | -0,0154387 | 0,98768216 |
| VC1781 | 1,16543565 | 0,23972025 | 1,00907233 | 2,3392E-07 | 1,00907233 | 0,01514725 | 0,98791472 |
| VC0464 | 1,07504223 | 0,03864053 | 1,00343104 | 3,7706E-08 | 1,00343104 | 0,01408997 | 0,98875821 |
| VC0494 | 1,07292345 | -0,0449894 | -1,0040282 | -4,39E-08 | -1,0040282 | -0,0140248 | 0,98881016 |
| VCA0364 | 1,05087634 | 0,29237172 | 1,00496065 | 2,853E-07 | 1,00496065 | 0,0139752 | 0,98884977 |
| VC1034 | -1,0074381 | -0,4267944 | -1,0030641 | -4,165E-07 | -1,0030641 | -0,0137787 | 0,98900658 |
| VC1535 | 1,14336059 | 0,02117602 | 1,00567514 | 2,0664E-08 | 1,00567514 | 0,0134073 | 0,98930285 |
| VC2481 | -1,0016257 | 0,16153582 | 1,00562905 | 1,5763E-07 | 1,00562905 | 0,01297889 | 0,98964462 |
| VC0333 | 1,02723314 | -0,0755802 | -1,0014213 | -7,375E-08 | -1,0014213 | -0,0126899 | 0,98987521 |
| VC0872 | -1,016671 | -0,2357448 | -1,0041435 | -2,3E-07 | -1,0041435 | -0,0123883 | 0,9901158 |
| VC2636 | 1,05924268 | 0,08405859 | 1,00262598 | 8,2025E-08 | 1,00262598 | 0,012198 | 0,99026766 |
| VC1669 | 1,10283396 | -0,0308852 | -1,0031686 | -3,014E-08 | -1,0031686 | -0,0121063 | 0,99034078 |
| VC2237 | 1,08237273 | -0,0616151 | -1,0021611 | -6,012E-08 | -1,0021611 | -0,0118538 | 0,99054222 |
| VC1259 | 1,10948878 | -0,0216603 | -1,0040651 | -2,114E-08 | -1,0040651 | -0,0114809 | 0,99083976 |
| VCA0076 | 1,13192577 | 0,02097372 | 1,00411499 | 2,0466E-08 | 1,00411499 | 0,01136639 | 0,99093113 |
| VC1857 | 1,05894076 | -0,0483483 | -1,0017692 | -4,718E-08 | -1,0017692 | -0,0113225 | 0,99096614 |
| VC2456 | 1,06332245 | -0,0349813 | -1,0021749 | -3,414E-08 | -1,0021749 | -0,010677 | 0,99148114 |
| alaS | 1,00916397 | 0,13963311 | 1,00167643 | 1,3626E-07 | 1,00167643 | 0,01055451 | 0,9915789 |
| VCA0602 | 1,17484918 | -0,0062736 | -1,0118709 | -6,122E-09 | -1,0118709 | -0,010538 | 0,99159203 |
| VCA0293 | 1,04766239 | 0,34530534 | 1,00355661 | 3,3695E-07 | 1,00355661 | 0,01048529 | 0,99163409 |
| VC1338 | 1,12864464 | 0,0161254 | 1,00506288 | 1,5735E-08 | 1,00506288 | 0,00960331 | 0,99233779 |
| ddl | -1,0031321 | 0,16723312 | 1,0006124 | 1,6319E-07 | 1,0006124 | 0,00941797 | 0,99248566 |
| VC1618 | 1,08161032 | 0,01873386 | 1,00308408 | 1,8281E-08 | 1,00308408 | 0,00930226 | 0,99257797 |
| VC1082 | 1,12909673 | 0,07280767 | 1,00371108 | 7,1046E-08 | 1,00371108 | 0,0091252 | 0,99271924 |
| VC2734 | 1,09068964 | 0,03229319 | 1,00144664 | 3,1512E-08 | 1,00144664 | 0,00836815 | 0,99332326 |
| VCA0850 | 1,08497369 | 0,02122635 | 1,00217561 | 2,0713E-08 | 1,00217561 | 0,00831839 | 0,99336298 |
| VCA0365 | -1,0252846 | -0,015806 | -1,0038417 | -1,542E-08 | -1,0038417 | -0,0079499 | 0,993657 |
| VCA0578 | 1,11916931 | -0,0482281 | -1,0020153 | -4,706E-08 | -1,0020153 | -0,0075274 | 0,99399402 |
| VCA0155 | 1,11878709 | 0,01258093 | 1,00292633 | 1,2277E-08 | 1,00292633 | 0,00742586 | 0,99407507 |
| VC2498 | 1,10837675 | 0,03294449 | 1,00128694 | 3,2148E-08 | 1,00128694 | 0,00735573 | 0,99413101 |
| VCA0038 | 1,11334869 | 0,05175936 | 1,00184927 | 5,0507E-08 | 1,00184927 | 0,00718322 | 0,99426866 |
| VC0350 | 1,06238985 | -0,358565 | -1,0009693 | -3,499E-07 | -1,0009693 | -0,0070134 | 0,99440417 |
| VC2072 | 1,07696317 | 0,02031097 | 1,00159432 | 1,982E-08 | 1,00159432 | 0,00696671 | 0,99444143 |
| VC1073 | 1,09106312 | 0,00899955 | 1,00353396 | 8,7818E-09 | 1,00353396 | 0,00690088 | 0,99449394 |
| VCA0994 | 1,12587625 | 0,01887031 | 1,0022485 | 1,8414E-08 | 1,0022485 | 0,00674711 | 0,99461659 |
| VC2728 | 1,06462658 | -0,0441142 | -1,0015664 | -4,305E-08 | -1,0015664 | -0,0065733 | 0,99475535 |
| VCA0648 | 1,14356338 | -0,0041037 | -1,0063249 | -4,004E-09 | -1,0063249 | -0,0062298 | 0,99502939 |
| VC2499 | 1,04230423 | -0,0369666 | -1,0008418 | -3,607E-08 | -1,0008418 | -0,0062262 | 0,99503226 |
| ndk | 1,0330528 | -0,5690937 | -1,0019459 | -5,553E-07 | -1,0019459 | -0,0061911 | 0,99506021 |
| VC2512 | 1,04285733 | 0,4805176 | 1,00216046 | 4,6889E-07 | 1,00216046 | 0,00581426 | 0,99536093 |
| VC2522 | 1,03591125 | -0,0198943 | -1,0011547 | -1,941E-08 | -1,0011547 | -0,0057015 | 0,99545088 |

| | | | | | | | |
|---|---|---|---|---|---|---|---|
| VCA0633 | 1,09536494 | -0,0136965 | -1,0013832 | -1,337E-08 | -1,0013832 | -0,0053003 | 0,99577096 |
| VC0197 | 1,08855439 | 0,00955857 | 1,00195633 | 9,3273E-09 | 1,00195633 | 0,00529361 | 0,9957763 |
| VC2744 | -1,0135659 | 0,12598126 | 1,00175979 | 1,2293E-07 | 1,00175979 | 0,00487432 | 0,99611088 |
| VCA0573 | 1,03370817 | -0,251466 | -1,0014506 | -2,454E-07 | -1,0014506 | -0,0046357 | 0,99630128 |
| VC0803 | 1,06380095 | 0,01233957 | 1,00104267 | 1,2041E-08 | 1,00104267 | 0,00439197 | 0,99649571 |
| tRNA-Asp-2 | 1,00406231 | -0,0376715 | -1,0019447 | -3,676E-08 | -1,0019447 | -0,0043471 | 0,99653153 |
| VC2356 | 1,00906483 | 0,11077134 | 1,00164249 | 1,0809E-07 | 1,00164249 | 0,00431445 | 0,99655759 |
| VC0681 | 1,01554558 | -0,0458717 | -1,0007225 | -4,476E-08 | -1,0007225 | -0,0037763 | 0,99698696 |
| VC1646 | 1,21742543 | -0,004775 | -1,0015746 | -4,659E-09 | -1,0015746 | -0,003357 | 0,99732154 |
| VC1996 | 1,08251 | -0,0605844 | -1,0011192 | -5,912E-08 | -1,0011192 | -0,0033107 | 0,99735846 |
| VCA1093 | 1,06284939 | -0,0045035 | -1,0013321 | -4,395E-09 | -1,0013321 | -0,0029988 | 0,99760735 |
| VC1975 | 1,05228408 | 0,01037715 | 1,00057631 | 1,0126E-08 | 1,00057631 | 0,0029947 | 0,99761053 |
| VC0215 | 1,0340995 | 0,00863578 | 1,00058972 | 8,4269E-09 | 1,00058972 | 0,0027635 | 0,99779507 |
| VC2632 | 1,13316723 | -0,0040577 | -1,0011906 | -3,96E-09 | -1,0011906 | -0,0026911 | 0,99785279 |
| VC0449 | 1,12104497 | -0,004336 | -1,0009705 | -4,231E-09 | -1,0009705 | -0,0025118 | 0,99799589 |
| VCA0798 | 1,10786615 | 0,06529161 | 1,00089541 | 6,3712E-08 | 1,00089541 | 0,00225341 | 0,99820208 |
| VC1692 | 1,10506676 | 0,0034 | 1,00094679 | 3,3177E-09 | 1,00094679 | 0,0021969 | 0,99824714 |
| VC2494 | 1,05103315 | 0,00555103 | 1,00054206 | 5,4167E-09 | 1,00054206 | 0,00205637 | 0,99835928 |
| rnfD | 1,05942729 | -0,0063379 | -1,0002601 | -6,185E-09 | -1,0002601 | -0,0013159 | 0,99895008 |
| VCA0134 | 1,07922866 | -0,002973 | -1,0004006 | -2,901E-09 | -1,0004006 | -0,0012095 | 0,99903496 |
| VC0939 | 1,03498128 | 0,00758364 | 1,00026088 | 7,4002E-09 | 1,00026088 | 0,00099872 | 0,99920317 |
| VC1244 | 1,09405649 | 0,00169297 | 1,00023608 | 1,652E-09 | 1,00023608 | 0,00077424 | 0,99938194 |
| VC0702 | 1,0173625 | 0,00628814 | 1,00011556 | 6,136E-09 | 1,00011556 | 0,0006665 | 0,99946812 |
| VC2308 | 1,0715251 | -0,0011941 | -1,0001338 | -1,165E-09 | -1,0001338 | -0,0004895 | 0,99960944 |
| greB | 1,12643403 | 0,0001861 | 1,00002286 | 1,816E-10 | 1,00002286 | 7,8614E-05 | 0,99993411 |

**Table S2:** Function prediction of ncRNA386 using CopaRNA (1), Prodoric virtual footprint (2) and Bprom (3) tools.

(1)

| fdr | p-value | NC_002505 | NC_011744 | NC_009457 | Annotation | Additional ho | Amount samp |
|---|---|---|---|---|---|---|---|
| 0,67133365 | 0,00047039 | vca1077(N/A\| | vs_rs14880(N | vc0395_rs00830(N/A\|-11.88730\|0.05013 | | | 0 |
| 0,67133365 | 0,00061721 | vc1781(N/A\|-19.15150\|0.0 | | vc0395_rs122 | N-acetylmannosamine-6-pho | | 1 |
| 0,67133365 | 0,00100862 | vc1647(N/A\|- | vs_rs20450(N | vc0395_rs11650(N/A\|-18.14830\|0.00062 | | | 0 |
| 0,67133365 | 0,00119614 | vc1403(N/A\|-12.36740\|0.0 | | vc0395_rs104 | methyl-accepting chemotaxi | | 1 |
| 0,67133365 | 0,00125118 | vc1199(N/A\|- | vs_rs05685(N | vc0395_rs09455(N/A\|-15.36210\|0.0052 | | | 0 |
| 0,67133365 | 0,00151222 | vc2281(N/A\|- | vs_rs20455(N | vc0395_rs147 | ribosomal protein S6 modifi | | 0 |
| 0,67133365 | 0,00152885 | vc1052(N/A\|- | vs_rs04240(N | vc0395_rs08215(N/A\|-11.08940\|0.07915 | | | 0 |
| 0,67133365 | 0,00160577 | vc0032(N/A\|- | vs_rs14695(N | vc0395_rs177 | ComM-like protein | | 0 |
| 0,67133365 | 0,00178546 | vca0179(N/A\| | vs_rs16310(N | vc0395_rs051 | NupC family protein | | 0 |
| 0,72344254 | 0,00213783 | vc0190(uvrD\| | vs_rs14450(u | vc0395_rs181 | DNA-dependent helicase II | | 0 |
| 0,76310201 | 0,00248053 | vc1312(alr\|-13.84970\|0.01 | | vc0395_rs100 | alanine racemase | | 1 |
| 0,81833189 | 0,00290189 | vca0083(N/A\| | vs_rs16260(N | vc0395_rs002 | multidrug resistance protein | | 0 |
| 0,86989414 | 0,00348331 | vca1015(N/A\| | vs_rs21435(N | vc0395_rs011 | Na+/H+ antiporter | | 0 |
| 0,86989414 | 0,00374343 | vca0513(N/A\| | vs_rs18175(N | vc0395_rs021 | aromatic amino acid aminot | | 0 |
| 0,86989414 | 0,00400451 | vc2550(N/A\|- | vs_rs13055(N | vc0395_rs15990(N/A\|-19.62410\|0.00017 | | | 0 |
| 0,86989414 | 0,00411297 | vca0892(N/A\| | vs_rs16845(N | vc0395_rs01670(N/A\|-17.69280\|0.00090 | | | 0 |
| 0,96177228 | 0,00504086 | vc2552(N/A\|- | vs_rs13060(N | vc0395_rs16000(N/A\|-16.60970\|0.00211 | | | 0 |
| 0,96177228 | 0,00608944 | vc1684(N/A\|- | vs_rs09365(N | vc0395_rs118 | peptide ABC transporter ATP | | 0 |
| 0,96177228 | 0,00654767 | vca0727(N/A\| | vs_rs18205(N | vc0395_rs032 | cobyric acid synthase | | 0 |
| 0,96177228 | 0,00688622 | vc2003(rrmA | vs_rs04305(rr | vc0395_rs133 | 23S rRNA (guanine(745)-N(1 | | 0 |
| 0,96177228 | 0,00692035 | vca0867(N/A\| | vs_rs20750(N | vc0395_rs017 | outer membrane protein W | | 0 |
| 0,96177228 | 0,00733564 | vca0988(N/A\|-12.05430\|0. | | vc0395_rs012 | methyl-accepting chemotaxi | | 1 |
| 0,96177228 | 0,00784734 | vc1944(N/A\|-10.24010\|0.1 | | vc0395_rs130 | pyoverdine biosynthesis pro | | 1 |
| 0,96177228 | 0,00788481 | vca0281(N/A\| | vs_rs05730(N | vc0395_rs045 | integrase | | 0 |

| | | | | | | | |
|---|---|---|---|---|---|---|---|
| 0,96177228 | 0,00793299 | vc0652(N/A\|- | vs_rs11630(N | vc0395_rs062 | protease | | 0 |
| 0,96177228 | 0,00793777 | vc1044(N/A\|- | vs_rs03995(N | vc0395_rs08170(N/A\|-11.45410\|0.06442 | | | 0 |
| 0,96177228 | 0,0079538 | vc1125(N/A\|- | vs_rs21580(N | vc0395_rs08565(N/A\|-13.88420\|0.01450 | | | 0 |
| 0,96177228 | 0,00851087 | vc1228(purT\| | vs_rs08025(N | vc0395_rs096 | phosphoribosylglycinamide | | 0 |
| 0,96177228 | 0,00943368 | vc1570(N/A\|-15.50820\|0.0 | | vc0395_rs112 | quinol oxidase subunit II | | 1 |
| 0,96177228 | 0,01047807 | vca0557(N/A\|-14.30300\|0.( | | vc0395_rs023 | diguanylate cyclase | | 1 |
| 0,96177228 | 0,01051466 | vc1493(N/A\|-10.49660\|0.1 | | vc0395_rs10880(N/A\|-10.35260\|0.11816 | | | 1 |
| 0,96177228 | 0,01055318 | vca1028(lamE | vs_rs15560(N | vc0395_rs01( | maltoporin | | 0 |
| 0,96177228 | 0,01070212 | vc1246(N/A\|- | vs_rs07875(N | vc0395_rs09680(N/A\|-13.03100\|0.02505 | | | 0 |
| 0,96177228 | 0,01115173 | vca0753(N/A\| | vs_rs18110(N | vc0395_rs03320(N/A\|-9.34270\|0.197207 | | | 0 |
| 0,96177228 | 0,01124335 | vca1000(N/A\| | vs_rs16515(N | vc0395_rs01175(N/A\|-9.53263\|0.179732 | | | 0 |
| 0,96177228 | 0,01147732 | vca1057(N/A\| | vs_rs20745(N | vc0395_rs009 | short chain dehydrogenase/r | | 0 |
| 0,96177228 | 0,01165943 | vca0927(N/A\|-14.66740\|0.( | | vc0395_rs01505(N/A\|-14.67740\|0.00853 | | | 1 |
| 0,96177228 | 0,01189884 | vca0529(N/A\|-11.90870\|0.( | | vc0395_rs022 | potassium uptake protein K( | | 1 |
| 0,96177228 | 0,0126299 | vc2309(N/A\|- | vs_rs03130(N | vc0395_rs148 | class V aminotransferase | | 0 |
| 0,96177228 | 0,01388258 | vc0026(N/A\|- | vs_rs00190(N | vc0395_rs177 | zinc-binding alcohol dehydr( | | 0 |
| 0,96177228 | 0,01394383 | vc1941(N/A\|-12.24450\|0.0 | | vc0395_rs13035(N/A\|-12.24450\|0.04056 | | | 1 |
| 0,96177228 | 0,01425891 | vc1833(N/A\|- | vs_rs05235(N | vc0395_rs125 | quinolinate synthetase | | 0 |
| 0,96177228 | 0,01477909 | vc2544(N/A\|- | vs_rs13025(N | vc0395_rs159 | fructose-1 6-bisphosphatase | | 0 |
| 0,96177228 | 0,01508287 | vca0843(N/A\| | vs_rs16440(N | vc0395_rs019 | glyceraldehyde-3-phosphate | | 0 |
| 0,96177228 | 0,01533392 | vc2665(groES | vs_rs01095(gr | vc0395_rs165 | co-chaperonin GroES | | 0 |
| 0,96177228 | 0,01538447 | vc1821(N/A\|- | vs_rs20960(N | vc0395_rs124 | PTS system fructose-specific | | 0 |
| 0,96177228 | 0,01552815 | vc2706(N/A\|- | vs_rs14410(N | vc0395_rs16735(N/A\|-12.26410\|0.04009 | | | 0 |
| 0,96177228 | 0,01557809 | vc0632(N/A\|- | vs_rs11740(N | vc0395_rs061 | D-alanyl-D-alanine carboxyp | | 0 |
| 0,96177228 | 0,01566796 | vc0474(N/A\|-15.16530\|0.0 | | vc0395_rs055 | iron-regulated virulence regu | | 1 |

| | | | | | | |
|---|---|---|---|---|---|---|
| 0,96177228 | 0,01567577 | vca0774(N/A\| | vs_rs18335(N | vc0395_rs034 | UDP-glucose 4-epimerase | 0 |
| 0,96177228 | 0,01674201 | vc1183(N/A\|- | vs_rs05580(N | vc0395_rs09380(N/A\|-16.04250\|0.00323 | | 0 |
| 0,96177228 | 0,01675882 | vc0035(N/A\|- | vs_rs14680(N | vc0395_rs177 | serine/threonine protein kin | 0 |
| 0,96177228 | 0,01693918 | vc0233(N/A\|- | vs_rs00865(N | vc0395_rs183 | 3-deoxy-D-manno-octulosor | 0 |
| 0,96177228 | 0,01698483 | vc0300(N/A\|- | vs_rs13830(N | vc0395_rs18745(N/A\|-9.23014\|0.208182 | | 0 |
| 0,96177228 | 0,01726985 | vca1067(N/A\| | vs_rs05805(N | vc0395_rs008 | aldehyde dehydrogenase | 0 |
| 0,96177228 | 0,01745259 | vc0788(N/A\|- | vs_rs19715(N | vc0395_rs069 | DOPA-dioxygenase-like prote | 0 |
| 0,96177228 | 0,01750979 | vc2542(N/A\|- | vs_rs13020(N | vc0395_rs159 | UDP-N-acetylmuramate:L-ala | 0 |
| 0,96177228 | 0,01811381 | vc0790(N/A\|-16.00690\|0.0 | | vc0395_rs069 | transcriptional regulator Cit | 1 |
| 0,96177228 | 0,01844746 | vc1675(N/A\|- | vs_rs09320(N | vc0395_rs117 | multidrug resistance protein | 0 |
| 0,96177228 | 0,01850848 | vca0386(N/A\|-11.17030\|0.0 | | vc0395_rs04440(N/A\|-11.17030\|0.07565 | | 1 |
| 0,96177228 | 0,01888105 | vca0217(N/A\|-15.47000\|0.0 | | vc0395_rs047 | diguanylate cyclase | 1 |
| 0,96177228 | 0,01890993 | vc2209(vibF\|-13.52890\|0.0 | | vc0395_rs143 | nonribosomal peptide synth | 1 |
| 0,96177228 | 0,01982672 | vc1418(N/A\|-16.44920\|0.0 | | vc0395_rs10495(N/A\|-16.44920\|0.00239 | | 1 |
| 0,96177228 | 0,02027173 | vca0154(N/A\|-11.55830\|0.0 | | vc0395_rs05310(N/A\|-11.55830\|0.06069 | | 1 |
| 0,96177228 | 0,02027869 | vc1063(N/A\|- | vs_rs10145(N | vc0395_rs082 | acyl-CoA thioesterase II | 0 |
| 0,96177228 | 0,02065843 | vc1899(N/A\|- | vs_rs09760(N | vc0395_rs12835(N/A\|-10.06280\|0.13750 | | 0 |
| 0,96177228 | 0,02077914 | vc2625(N/A\|- | vs_rs13480(N | vc0395_rs163 | ribulose-phosphate 3-epime | 0 |
| 0,96177228 | 0,02111223 | vc1111(N/A\|- | vs_rs09565(N | vc0395_rs084 | adenosylmethionine--8-amir | 0 |
| 0,96177228 | 0,02285477 | vc2183(N/A\|- | vs_rs03440(N | vc0395_rs141 | ribose-phosphate pyrophosp | 0 |
| 0,96177228 | 0,02301722 | vc0117(N/A\|- | vs_rs14345(N | vc0395_rs173 | protoheme IX synthesis prot | 0 |
| 0,96177228 | 0,02323439 | vc1664(N/A\|- | vs_rs05460(N | vc0395_rs117 | ABC transporter substrate-bi | 0 |
| 0,96177228 | 0,02339939 | vca0837(N/A\| | vs_rs03025(N | vc0395_rs019 | hemolysin | 0 |
| 0,96177228 | 0,0234136 | vc0170(N/A\|- | vs_rs00325(N | vc0395_rs181 | peptide ABC transporter ATP | 0 |
| 0,96177228 | 0,02370758 | vca0677(N/A\| | vs_rs17635(N | vc0395_rs029 | nitrate reductase assembly p | 0 |
| 0,96177228 | 0,02379802 | vc2046(N/A\|- | vs_rs04455(N | vc0395_rs13540(N/A\|-9.72305\|0.163486 | | 0 |
| 0,96177228 | 0,02415393 | vc0081(N/A\|- | vs_rs00430(N | vc0395_rs17475(N/A\|-13.71210\|0.01622 | | 0 |
| 0,96177228 | 0,02444667 | vc1170(N/A\|- | vs_rs09230(N | vc0395_rs093 | tryptophan synthase subuni | 0 |
| 0,96177228 | 0,02453345 | vca0105(N/A\| | vs_rs17705(N | vc0395_rs00170(N/A\|-9.88 | vca0284 | 0 |
| 0,96177228 | 0,02455566 | vc0396(N/A\|- | vs_rs01470(N | vc0395_rs193 | LuxR family transcriptional r | 0 |
| 0,96177228 | 0,02475606 | vc0862(N/A\|-11.65160\|0.0 | | vc0395_rs07290(N/A\|-11.51890\|0.06208 | | 1 |
| 0,96177228 | 0,02481513 | vca0936(N/A\| | vs_rs16085(N | vc0395_rs01470(N/A\|-12.94740\|0.02640 | | 0 |
| 0,96177228 | 0,02590472 | vca0036(N/A\| | vs_rs15640(N | vc0395_rs004 | serine/threonine transporte | 0 |
| 0,96177228 | 0,02641594 | vc2111(N/A\|- | vs_rs04160(N | vc0395_rs13845(N/A\|-10.88680\|0.08856 | | 0 |
| 0,96177228 | 0,02686364 | vc0968(N/A\|- | vs_rs10740(N | vc0395_rs078 | cysteine synthase A | 0 |
| 0,96177228 | 0,02728307 | vc0674(N/A\|- | vs_rs02420(N | vc0395_rs063 | prolipoprotein diacylglycery | 0 |
| 0,96177228 | 0,0273354 | vc2113(N/A\|- | vs_rs04150(N | vc0395_rs13855(N/A\|-11.68960\|0.05626 | | 0 |
| 0,96177228 | 0,0274794 | vc0678(N/A\|- | vs_rs02450(N | vc0395_rs064 | transcriptional activator Hly | 0 |
| 0,96177228 | 0,02794084 | vca0851(N/A\| | vs_rs21700(N | vc0395_rs01870(N/A\|-12.16330\|0.04258 | | 0 |
| 0,96177228 | 0,02796304 | vc1275(N/A\|- | vs_rs08960(N | vc0395_rs09820(N/A\|-14.68360\|0.00849 | | 0 |
| 0,96177228 | 0,02871491 | vc0550(N/A\|- | vs_rs12110(N | vc0395_rs057 | oxaloacetate decarboxylase | 0 |
| 0,96177228 | 0,02907195 | vc0947(N/A\|- | vs_rs03260(N | vc0395_rs077 | D-alanyl-D-alanine carboxyp | 0 |
| 0,96177228 | 0,02943709 | vc1210(N/A\|- | vs_rs09170(N | vc0395_rs09510(N/A\|-10.74550\|0.09568 | | 0 |
| 0,96177228 | 0,03053746 | vc1502(yebU | vs_rs06625(ye | vc0395_rs109 | 16S rRNA (cytosine(1407)-C( | 0 |
| 0,96177228 | 0,03070528 | vc1780(N/A\|-16.16140\|0.0 | | vc0395_rs12270(N/A\|-16.16140\|0.00296 | | 1 |
| 0,96177228 | 0,03078269 | vc1712(N/A\|- | vs_rs15620(N | vc0395_rs11960(N/A\|-13.20910\|0.02240 | | 0 |
| 0,96177228 | 0,03095321 | vca0164(N/A\| | vs_rs20505(N | vc0395_rs05260(N/A\|-10.62300\|0.10225 | | 0 |
| 0,96177228 | 0,03127869 | vca0650(N/A\| | vs_rs13115(N | vc0395_rs02845(N/A\|-11.11840\|0.07788 | | 0 |
| 0,96177228 | 0,03134949 | vc1380(N/A\|- | vs_rs08300(N | vc0395_rs10330(N/A\|-16.22570\|0.00282 | | 0 |
| 0,96177228 | 0,03173774 | vc0142a(N/A\|-10.90540\|0.0 | | vc0395_rs17200(N/A\|-10.90540\|0.08765 | | 1 |
| 0,96177228 | 0,03187251 | vc0040(N/A\|- | vs_rs14660(N | vc0395_rs177 | hemolysin | 0 |

| fdr | p-value | NC_002505 | NC_011744 | NC_009457 | Annotation | Additional ho | Amount samp |
|---|---|---|---|---|---|---|---|
| 0,7047429 | 0,0002085 | vc0969(N/A\|- | vs_rs10735(N | vc0395_rs078 | sulfate transport protein Cys | | 0 |
| 0,72258242 | 0,00066371 | vca0535(N/A\| | -16.45890\|0.( | vc0395_rs02255(N/A\|-16.28460\|0.00113 | | | 1 |
| 0,72258242 | 0,00080215 | vca1028(lamE | vs_rs15560(N | vc0395_rs01( | maltoporin | | 0 |
| 0,72258242 | 0,00093523 | vc0734(N/A\|- | vs_rs16065(N | vc0395_rs06€ | malate synthase | | 0 |
| 0,72258242 | 0,00181617 | vc2269(N/A\|- | vs_rs11335(N | vc0395_rs14€ | 3 4-dihydroxy-2-butanone-4 | | 0 |
| 0,72258242 | 0,00203364 | vc0299(N/A\|- | vs_rs13835(N | vc0395_rs187 | DNA polymerase III subunit e | | 0 |
| 0,72258242 | 0,00220172 | vc2120(flhB\|- | vs_rs03870(N | vc0395_rs138 | flagellar biosynthesis proteir | | 0 |
| 0,72258242 | 0,00235208 | vc2551(N/A\|- | 18.02100\|0.0 | vc0395_rs15995(N/A\|-17.65710\|0.00028 | | | 1 |
| 0,72258242 | 0,00247155 | vc1571(N/A\|- | 15.54290\|0.0 | vc0395_rs112 | quinol oxidase subunit I | | 1 |
| 0,72258242 | 0,00276011 | vc2341(N/A\|- | vs_rs11525(N | vc0395_rs14¢ | long-chain-fatty-acid--CoA li | | 0 |
| 0,72258242 | 0,00288888 | vca1091(N/A\| | -12.39940\|0.( | vc0395_rs007 | chemotaxis protein CheR | | 1 |
| 0,72258242 | 0,00325645 | vc1045(N/A\|- | vs_rs04000(N | vc0395_rs081 | RNA polymerase sigma factoı | | 0 |
| 0,72258242 | 0,00340164 | vc2678(N/A\|- | vs_rs13625(N | vc0395_rs16€ | primosome assembly proteir | | 0 |
| 0,72258242 | 0,00351211 | vc1330(N/A\|- | 19.06540\|5.6 | vc0395_rs10090(N/A\|-19.06540\|5.61605 | | | 1 |
| 0,72258242 | 0,00376348 | vc2112(N/A\|- | vs_rs04155(N | vc0395_rs13850(N/A\|-13.90110\|0.00868 | | | 0 |
| 0,72258242 | 0,00384544 | vc1124(N/A\|- | vs_rs21575(N | vc0395_rs08560(N/A\|-13.88420\|0.0088C | | | 0 |
| 0,72258242 | 0,0039303 | vc0234(N/A\|- | 15.39800\|0.0 | vc0395_rs18370(N/A\|-15.39800\|0.00253 | | | 1 |
| 0,72258242 | 0,0039557 | vc1683(N/A\|- | vs_rs09360(N | vc0395_rs118 | peptide ABC transporter ATP | | 0 |
| 0,72258242 | 0,00406185 | vca1073(N/A\| | vs_rs14895(N | vc0395_rs008 | bifunctional proline dehydrc | | 0 |
| 0,76229193 | 0,00501556 | vc0795(N/A\|- | 15.37000\|0.0 | vc0395_rs06¢ | citrate/sodium symporter | | 1 |
| 0,76229193 | 0,00506414 | vc2714(ompF | vs_rs00700(o | vc0395_rs167 | osmolarity response regulatc | | 0 |
| 0,76229193 | 0,00511154 | vc1888(N/A\|- | 14.59300\|0.0 | vc0395_rs127 | hemolysin-like protein | | 1 |
| 0,76229193 | 0,00518719 | vc1417(N/A\|- | 16.33530\|0.0 | vc0395_rs10490(N/A\|-16.33530\|0.00108 | | | 1 |
| 0,77508134 | 0,00574572 | vc0475(N/A\|- | vs_rs07620(N | vc0395_rs055 | enterobactin receptor prote | | 0 |
| 0,77508134 | 0,00575849 | vc1328(mglC | \|-14.15800\|0. | vc0395_rs10( | beta-methylgalactoside tran | | 1 |
| 0,77508134 | 0,00638798 | vca0621(N/A\| | vs_rs17005(N | vc0395_rs027 | SorC family transcriptional r | | 0 |
| 0,77508134 | 0,00661863 | vc0653(N/A\|- | vs_rs11625(N | vc0395_rs062 | c-di-GMP phosphodiesterase | | 0 |
| 0,77508134 | 0,00682871 | vca0223(N/A\| | vs_rs19785(N | vc0395_rs047 | protease | | 0 |
| 0,77508134 | 0,00684278 | vc2454(N/A\|- | 11.63590\|0.0 | vc0395_rs155 | GGDEF family protein | | 1 |
| 0,77508134 | 0,00687942 | vc2308(N/A\|- | 13.40890\|0.0 | vc0395_rs148 | 4-methyl-5(B-hydroxyethyl)- | | 1 |
| 0,81772983 | 0,00749989 | vc0861(N/A\|- | vs_rs03015(N | vc0395_rs072 | type IV pilin | | 0 |
| 0,8353599 | 0,0087217 | vc1317(N/A\|- | 12.93670\|0.0 | vc0395_rs10025(N/A\|-12.93670\|0.01787 | | | 1 |
| 0,8353599 | 0,00884169 | vc2046(N/A\|- | vs_rs04455(N | vc0395_rs13540(N/A\|-13.56780\|0.01121 | | | 0 |
| 0,8353599 | 0,00895575 | vca0090(N/A\| | vs_rs15250(N | vc0395_rs00245(N/A\|-13.59600\|0.01097 | | | 0 |
| 0,8353599 | 0,00923794 | vca0104(N/A\| | vs_rs20410(N | vc0395_rs001 | ribosomal large subunit pseu | | 0 |
| 0,8353599 | 0,00966643 | vc2337(N/A\|- | vs_rs21270(N | vc0395_rs14¢ | LacI family transcription reg | | 0 |
| 0,8353599 | 0,0099177 | vc1820(N/A\|- | vs_rs20955(N | vc0395_rs124 | PTS system fructose-specific | | 0 |
| 0,8353599 | 0,00992829 | vc0605(N/A\|- | vs_rs11875(N | vc0395_rs06035(N/A\|-10.83440\|0.07263 | | | 0 |
| 0,8353599 | 0,01011842 | vca0676(N/A\| | vs_rs17640(N | vc0395_rs02¢ | (Fe-S)-binding protein | | 0 |
| 0,8353599 | 0,010179 | vc2489(N/A\|- | vs_rs01755(N | vc0395_rs156 | TetR family transcriptional r | | 0 |
| 0,8353599 | 0,0104376 | vc1612(N/A\|- | vs_rs07835(N | vc0395_rs114 | fimbrial biogenesis and twitc | | 0 |
| 0,8353599 | 0,01060633 | vc1557(N/A\|- | vs_rs15270(N | vc0395_rs111 | LacI family transcription reg | | 0 |
| 0,8353599 | 0,01062736 | vc2318(N/A\|- | 14.64220\|0.0 | vc0395_rs14865(N/A\|-14.49120\|0.00543 | | | 1 |
| 0,89523845 | 0,01165399 | vc0332(hemE | vs_rs13925(h | vc0395_rs18¢ | uroporphyrinogen decarbox | | 0 |
| 0,90699675 | 0,01237922 | vc1274(N/A\|- | vs_rs08965(N | vc0395_rs09815(N/A\|-14.89070\|0.00391 | | | 0 |
| 0,90699675 | 0,01283052 | vc1354(N/A\|- | vs_rs06545(N | vc0395_rs10200(N/A\|-12.12710\|0.0315C | | | 0 |
| 0,90699675 | 0,01310509 | vca0471(N/A\| | -13.00440\|0.( | vc0395_rs03845(N/A\|-13.00440\|0.01701 | | | 1 |

| | | | | | | | |
|---|---|---|---|---|---|---|---|
| 0,90699675 | 0,0133838 | vc2421(N/A\|- | vs_rs11990(N | vc0395_rs153 | N-acetyl-anhydromuranmyl- | | 0 |
| 0,90699675 | 0,01356139 | vc2103(N/A\|-12.78870\|0.0 | | vc0395_rs138 | LysR family transcriptional r | | 1 |
| 0,90699675 | 0,01365468 | vc0506(N/A\|- | vs_rs07755(N/A\|-7.67656\|0.2552526\|148\|159\|166\|1 | | | | 1 |
| 0,90699675 | 0,01368545 | vc1105(N/A\|- | vs_rs04655(N/A\|-13.13910\|0.007982914\|183\|212\|14 | | | | 1 |
| 0,91070222 | 0,0140108 | vc0118(N/A\|- | vs_rs14340(N | vc0395_rs173 | uroporphyrin-III C-methyltra | | 0 |
| 0,91806554 | 0,0143957 | vc0191(N/A\|- | vs_rs14435(N | vc0395_rs18165(N/A\|-12.77340\|0.0200 | | | 0 |
| 0,93732673 | 0,01508615 | vc1193(N/A\|- | vs_rs05655(N | vc0395_rs09425(N/A\|-11.13730\|0.06018 | | | 0 |
| 0,93732673 | 0,01525236 | vc2710(N/A\|- | vs_rs00715(N | vc0395_rs167 | bifunctional (p)ppGpp synth | | 0 |
| 0,94474656 | 0,01573798 | vca0747(glpA\|-12.92320\|0 | | vc0395_rs032 | sn-glycerol-3-phosphate deh | | 1 |
| 0,94474656 | 0,01624288 | vca0150(def\|-12.12540\|0.0 | | vc0395_rs053 | peptide deformylase | | 1 |
| 0,94474656 | 0,0163942 | vc0509(N/A\|- | vs_rs07735(N/A\|-17.04020\|0.0003319843\|6\|49\|149\| | | | | 1 |
| 0,94474656 | 0,01649114 | vc1165(N/A\|- | vs_rs05150(N | vc0395_rs09025(N/A\|-11.65510\|0.04316 | | | 0 |
| 0,94795314 | 0,01735956 | vca0832(N/A\| | vs_rs16950(N | vc0395_rs01 | methyltransferase | | 0 |
| 0,94795314 | 0,01736028 | vca0852(N/A\| | vs_rs21705(N | vc0395_rs01865(N/A\|-12.16330\|0.03073 | | | 0 |
| 0,94795314 | 0,01738849 | vc2690(fieF\|- | vs_rs13700(fi | vc0395_rs166 | ferrous iron efflux protein F | | 0 |
| 0,95877982 | 0,01816075 | vca0937(N/A\| | vs_rs15875(N | vc0395_rs014 | AraC family transcriptional r | | 0 |
| 0,95877982 | 0,01861071 | vc1435(N/A\|- | vs_rs06870(N | vc0395_rs10585(N/A\|-9.00220\|0.204151 | | | 0 |
| 0,95877982 | 0,01878804 | vc0549(N/A\|- | vs_rs12115(N | vc0395_rs057 | oxaloacetate decarboxylase | | 0 |
| 0,95877982 | 0,01895165 | vc2472(N/A\|- | vs_rs12390(N | vc0395_rs15590(N/A\|-11.64280\|0.04351 | | | 0 |
| 0,95877982 | 0,0190054 | vc0036(N/A\|- | vs_rs14675(N | vc0395_rs177 | FixG-like protein | | 0 |
| 0,96551865 | 0,01999628 | vc2182(ipk\|- | vs_rs03445(N | vc0395_rs141 | 4-diphosphocytidyl-2C-meth | | 0 |
| 0,96551865 | 0,02023693 | vc0188(N/A\|- | vs_rs00375(N | vc0395_rs181 | oligopeptidase A | | 0 |
| 0,96551865 | 0,02035321 | vc0628(N/A\|- | vs_rs11765(N | vc0395_rs06150(N/A\|-13.84430\|0.00907 | | | 0 |
| 0,96551865 | 0,02082078 | vc0452(N/A\|- | vs_rs12680(N | vc0395_rs053 | A/G-specific adenine glycosy | | 0 |
| 0,96551865 | 0,02101114 | vc1166(aspS\| | vs_rs05155(N | vc0395_rs090 | aspartyl-tRNA synthetase | | 0 |
| 0,96551865 | 0,02144085 | vc2001(N/A\|- | vs_rs04315(N | vc0395_rs13305(N/A\|-12.55370\|0.02346 | | | 0 |
| 0,96551865 | 0,02160732 | vca0651(N/A\| | vs_rs13110(N | vc0395_rs02850(N/A\|-11.08730\|0.06210 | | | 0 |
| 0,96551865 | 0,02226931 | vc0755(N/A\|- | vs_rs02815(N | vc0395_rs067 | aminopeptidase | | 0 |
| 0,96551865 | 0,02245986 | vca0933(N/A\| | vs_rs00795(N | vc0395_rs014 | cold shock do | vca0166 | 0 |
| 0,96551865 | 0,0228798 | vc0142a(N/A\|-11.37390\|0. | | vc0395_rs17200(N/A\|-11.37390\|0.05178 | | | 1 |
| 0,96551865 | 0,02334764 | vc2119(N/A\|- | vs_rs04025(N | vc0395_rs13880(N/A\|-11.52910\|0.04685 | | | 0 |
| 0,96551865 | 0,02336704 | vc0282(N/A\|-13.18480\|0.0 | | vc0395_rs185 | methyl-accepting chemotaxi | | 1 |
| 0,96551865 | 0,02354968 | vca0653(N/A\|-12.26510\|0. | | vc0395_rs028 | PTS system sucrose-specific t | | 1 |
| 0,96551865 | 0,02363142 | vc2094(N/A\|- | vs_rs10550(N | vc0395_rs13770(N/A\|-11.16730\|0.05905 | | | 0 |
| 0,96551865 | 0,02364395 | vc1521(N/A\|- | vs_rs07010(N | vc0395_rs110 | sensor histidine kinase | | 0 |
| 0,96551865 | 0,02399223 | vc1544(N/A\|- | vs_rs18015(N | vc0395_rs111 | TonB2 protein | | 0 |
| 0,96551865 | 0,02399514 | vc0673(N/A\|- | vs_rs02415(N | vc0395_rs06375(N/A\|-10.29960\|0.10009 | | | 0 |
| 0,96619257 | 0,02459458 | vca0245(N/A\| | vs_rs16290(N | vc0395_rs046 | PTS system transporter subu | | 0 |
| 0,96619257 | 0,02476053 | vc2552(N/A\|- | vs_rs13060(N | vc0395_rs16000(N/A\|-14.04000\|0.00779 | | | 0 |
| 0,96619257 | 0,02498759 | vca0742(N/A\|-12.41760\|0. | | vc0395_rs03270(N/A\|-13.96260\|0.00828 | | | 1 |
| 0,96619257 | 0,02578974 | vc2380(N/A\|- | vs_rs02190(N | vc0395_rs151 | adenosylcobinamide-phosp | | 0 |
| 0,96619257 | 0,02613617 | vc1180(N/A\|- | vs_rs05545(N | vc0395_rs093 | cysteine/glutathione ABC tra | | 0 |
| 0,96619257 | 0,02616266 | vca0971(N/A\| | vs_rs21370(N | vc0395_rs01310(N/A\|-11.40510\|0.05076 | | | 0 |
| 0,96619257 | 0,02664766 | vca1037(N/A\| | vs_rs17335(gl | vc0395_rs010 | amino acid ABC transporter | | 0 |
| 0,96619257 | 0,02703454 | vc0622(N/A\|- | vs_rs11795(N | vc0395_rs061 | sensory box sensor histidine | | 0 |
| 0,96619257 | 0,02708075 | vc1171(N/A\|- | vs_rs09225(N | vc0395_rs093 | bifunctional indole-3-glycer | | 0 |
| 0,96619257 | 0,02724277 | vca0475(N/A\|-8.43769\|0.2 | | vc0395_rs038 | doc protein | | 1 |
| 0,96619257 | 0,02736236 | vc2771(N/A\|- | vs_rs14780(N | vc0395_rs179 | F0F1 ATP synthase subunit I | | 0 |
| 0,96619257 | 0,02744216 | vc1581(N/A\|-11.29700\|0.0 | | vc0395_rs113 | NADH dehydrogenase subun | | 1 |
| 0,96941656 | 0,02823143 | vc1356(N/A\|- | vs_rs06560(N | vc0395_rs102 | sulfite reductase subunit gar | | 0 |
| 0,96941656 | 0,02845592 | vc1184(N/A\|- | vs_rs05585(N | vc0395_rs093 | NifS-like protein | | 0 |
| 0,96941656 | 0,02925497 | vc0046(def\|- | vs_rs14630(N | vc0395_rs176 | peptide deformylase | | 0 |
| 0,96941656 | 0,02941438 | vca0785(N/A\| | vs_rs18380(N | vc0395_rs034 | diguanylate cyclase | | 0 |

(2)

| Matrix | Position | Strand | Binding Seque | Score | Core Score | Locus (Gene) | Dist to ATG | Product |
|---|---|---|---|---|---|---|---|---|
| MX000366 | 18 | + | AGCGAGCCAA | 5.124725521 | 3.712140441 | undefined (un | undefined | undefined |
| MX000366 | 81 | - | TTAGATCCAGT | 4.889834628 | 3.712140441 | undefined (un | undefined | undefined |
| MX000366 | 48 | + | TGTGATCCTAC | 4.794736798 | 3.567173032 | undefined (un | undefined | undefined |
| MX000366 | 20 | + | CGAGCCAAAT | 4.359936236 | 2.852052152 | undefined (un | undefined | undefined |
| MX000366 | 43 | - | GTAGGATCAC | 4.211242630 | 3.307393011 | undefined (un | undefined | undefined |
| MX000366 | 168 | * | TACGATTTACC | 4.188143612 | 3.225760358 | undefined (un | undefined | undefined |

(3)

| For promoter at 64: |
|---|
| ompR: AAATCACA at position 27 Score - 7 |
| crp: ATCACAAA at position 29 Score - 10 |
| arcA: TCACAAAA at position 30 Score - 10 |
| ihf: ACAAAAAA at position 32 Score - 9 |

**Table S3:** Strains, plasmids and primers

| **Strain number** | **Phenotype of interest** |
|---|---|
| *Acinetobacter baumannii* ATCC 19606 | WT |
| *E. coli* MG1655 | WT |
| *Klebsiella pneumoniae* ML21 | WT |
| *Pseudomonas aeruginosa* ATCC 10145 | WT |
| *Vibrio cholerae* N16961 | WT |
| *V. tasmaniensis* | WT |
| *V. mimicus* | WT |
| *V. parahaemolyticus* | WT |
| *V. vulnificus* | WT |
| D908 | N16961 ΔctrR clean deletion (formerly known as Δ ncRNA586) |
| F862 | *V. cholerae N16961 with* PctrRVch plasmid that contains ctrR-pBAD24 |
| F863 | *V. cholerae N16961* with PctrRVta plasmid containing VsR_217-pBAD24 |
| F864 | V. cholerae N16961 with empty-pBAD24 |
| H114/H115 | V. cholerae N16961 with PctrRVch-CC plasmid that contains the mutated version ctrR-CC-pBAD24 |
| K830 | ΔVC1820-28 clean deletion using plasmid A200 and oligos 5VC181, 7VC181, 8VC181, 6VC181 that amplify 500 bp upstream and dowstream from VC1820-28 locus |
| L147 | ΔlamB, clean deletion using plasmid A200 and oligos 5lamB, 7lamB, 8lamB, 6lamB that amplify 500 bp upstream and dowstream from the maltoporin gene |
| L148 | Δmalop, clean deletion using plasmid A200 and oligos 5malop1, 7malop1, 8malop1, 6malop1 that amplify 500 bp upstream and dowstream from the maltose transport operon |
| Top 10 | F- mcrA Δ( mrr-hsdRMS-mcrBC) Φ80lacZΔM15 Δ lacX74 recA1 araD139 Δ(ara-leu)7697 galU galK rpsL (StrR) endA1 nupG |
| π3813 cells | To grow plasmids for mutant construction before allelic exchange |
| β3914 | β2163 [(F-) RP4-2-Tc::Mu ΔdapA::(erm-pir116)] gyrA462 zei-298::Tn10 |

| **Plasmid** | **Description** |
|---|---|
| pF585 | pBAD24 with ctrRVch cloned for Ara overexpression |
| pF587 | pBAD24 with ctrRVta cloned for Ara overexpression |
| pBAD24 | ori pBR322, carbenicillin resistance |
| A200 | Flipase continaing plasmid |
| pSW7848 | Plasmid contining frt sites surrounding the aph gene for allelic exchange of target genes |
| pSC1819 | Plasmid containing the mariner tranposon and a spec resistance gene for saturated mutant library generation |

| **Oligo name** | **Sequence** | **Use** |
|---|---|---|
| 5malop1 | CTATTATTTAAACTCTTTCCCAACCACTGGGATGGAGCG | Amplification of the 500 bp upstream of the maltose operon for clean deletion |
| 7malop1 | CTACACAATCGCTCAAGACGTGTGGCGTTACGCATAGCTGGC | Amplification of the 500 bp upstream of the maltose operon for clean deletion |
| 8malop1 | CTAATTCCCATGTCAGCCGTTCCTCCTAACCCCTACGCC | Amplification of the 500 bp downstream of the maltose operon for clean deletion |
| 6malop1 | TACGTAGAATGTATCAGACTATCAAGTGGTTTCGTTGGCAGCG | Amplification of the 500 bp downstream of the maltose operon for clean deletion |
| 5vc181 | CTATTATTTAAACTCTTTCCACCTCGCCTACCGTATGG | Amplification of the 500 bp upstream |

| | | |
|---|---|---|
| | | of the VC1820-27 locus for clean deletion |
| 7vc181 | CTACACAATCGCTCAAGACGTGTTCTTGAAGTGGCAGCGG | Amplification of the 500 bp upstream of the VC1820-27 locus for clean deletion |
| 8vc181 | CTAATTCCCATGTCAGCCGTGCACATATCGAGGAAATAGCCG | Amplification of the 500 bp downstream of the VC1820-27locus for clean deletion |
| 6vc181 | TACGTAGAATGTATCAGACTTATTCGTCATGCAGCGGC | Amplification of the 500 bp downstream of the VC1820-27 locus for clean deletion |
| 5lamB | CTATTATTTAAACTCTTTCCAGAAGGACCAGAGACATCCAGC | Amplification of the 500 bp upstream of the lamB gene for clean deletion |
| 7lamB | CTACACAATCGCTCAAGACGTGAGCGGGTTGTCAGAGTACAGC | Amplification of the 500 bp upstream of the lamB gene for clean deletion |
| 8lamB | CTAATTCCCATGTCAGCCGTAGGCAGAAAGTTGGCAGC | Amplification of the 500 bp downstream of the lamB gene for clean deletion |
| 6lamB | TACGTAGAATGTATCAGACTAATGCAATTCATCATTGAGCTGG | Amplification of the 500 bp downstream of the lamB gene for clean deletion |
| crpfix500F | TTGTAGGGCTCACCAAGAGG | Amplification of the ctrR promoter region for EMSA |
| crpfix500R | ACATCAAGTGGTTTCGTTGGC | Amplification of the ctrR promoter region for EMSA |
| micX500F | ACCAACCGCAAGCGTGTACG | Amplification of the micX promoter region for EMSA |
| micX500R | TGCTGATGGTTCAGTCACTCCACC | Amplification of the micX promoter region for EMSA |
| MV268 | CACGTCTTGAGCGATTGTGTAG | For amplification of construction verification of pSW7848 plasmid for clean deletion of different mutants constructed |
| MV269 | ACGGCTGACATGGGAATTAG | For amplification of construction verification of pSW7848 plasmid for clean deletion of different mutants constructed |
| MV450 | GGAAAGAGTTTAAATAATAGTGCAGGAATTCGATATCAAG | For amplification of construction verification of pSW7848 plasmid for clean deletion of different mutants constructed |
| MV451 | AGTCTGATACATTCTACGTAGGGATCCACTAGTTCTAGAG | For amplification of construction verification of pSW7848 plasmid for clean deletion of different mutants constructed |
| 4347 | TCTCCATATCCCAGAGAGGATGGTAAGAGCCTTTCC | Amplification of the 197 bp containing ctrR sequence from V. cholerae |
| 4348 | TATTTGATGAAAAGAAAGGGTGAACGCTCCGGC | Amplification of the 197 bp containing ctrR sequence from V. cholerae |
| 4343 | CTGGGATATGGAGAAACAGTAGAGAGTTGCGATAAAAAGC | Amplification of pBAD24 plasmid for cloning ctrR-Vch |
| 4344 | TTCTTTTCATCAAATAAAACGAAAGGCTCAGTCGAAAG | Amplification of pBAD24 plasmid for cloning ctrR-Vch |
| ML 435 | CGCGAATTCATCCAAGAGTGGATAGTAAGAACCC | Amplification Vsr217-Vta |
| ML 436 | CGCTCTAGATTAACCAAGCCCACCCGTCATAGCC | Amplification VsR217-Vta |
| ML466 | cgctctagaCACGGGACTGCCGCGAAAAGTAGG | Point mutation in ctrR |
| dRT-PCR | | |
| ML 455 | [FAM] CTGGTTCCACCGCCAAGTAGGTTGT [BHQ1] | Probe |
| ML 456 | GTTGTTTCCCCCTACACAAAG | Reverse |
| ML 457 | CCCAGAGAGGATGGTAAGAG | forward |
